# New potential antimicrobial peptides with amazing symmetrical structure in fungi and insects

**DOI:** 10.64898/2025.12.21.695455

**Authors:** Jiao Zhu, Wolfgang Knoll, Bing Wang, Paolo Pelosi

## Abstract

We have discovered a new family of genes encoding potential antimicrobial peptides with unique compact and elegant structure in the genomes of several Fungi and some arthropod species. Their expression products are constituted of about 85 amino acids, including a signal peptide, and are folded into two α-helical segments connected by a short unstructured coil. Three conserved disulphide bridges between cysteines located in symmetrically mirrored positions connect the two helical domains. The two ends of the chain are thus brought together and the elongated and compact shape suggests that of a nail, potentially able to penetrate the cell membrane. A high abundance of hydrophobic residues supports such hypothesis. These peptides, that we name Hairpin Loop Peptides (HLPs), have been found in the genomes of many Fungi species but only in selected clades. Orthologues have also been discovered in the genomes of some insects, notably Hemiptera, a few other arthropods and other organisms, but are absent in plants. They appear to have originated in Fungi and then migrated to insects through horizontal gene transfer. The antimicrobial activity of HLPs is predicted by several software programmes, although this still needs to be confirmed by experimental evidence. The occurrence of HLPs in several edible mushrooms supports potential uses of these peptides in food preservation and possibly also in medical applications. Their simple and nearly rigid structure can be easily modified to improve specificity, stability and solubility, thus making these molecular weapons suitable for a variety of different applications.

**Significance statement:** The increasing emergence of bacteria resistant to current antibiotics has stimulated a rapid search for alternative treatments. In recent years, antimicrobial peptides have attracted considerable interest for their potential applications in medicine and as food preservatives. We have identified a new class of peptides primarily expressed in fungi, including edible mushrooms, and also detected in some insects and other arthropods, likely as a result of horizontal gene-transfer events. These peptides are particularly noteworthy because of their compact, elongated, and highly symmetrical structures, which give them a nail-like shape capable of penetrating cellular membranes. Such structural features suggest potential antimicrobial activity, a prediction supported by computational analyses. Their widespread presence in edible mushrooms further indicates the potential safety of these peptides for human use.

## Introduction

The advent of the genome era has deeply changed the approach to study the biochemical mechanisms responsible for the functioning of cells and living organisms. Scientists often start with the amino acid sequence of a protein and by leveraging bioinformatic analyses and experimental data, they seek to deduce the protein’s function, chart its spatial and temporal expression, and uncover its interactions with other cellular components. Such approach has been recently applied to odorant-binding proteins to uncover their roles and those of their ligands in chemical communication along a strategy known as *Reverse Chemical Ecology* (Leal 2017; Zhu et al. 2017; Zaremska et al. 2021; Zaremska et al. 2022). The availability of the Alpha-Fold programme, able to predict the three-dimensional structure of a protein *ab-initio* with unprecedented accuracy and reliability, has added an important and very precious tool for investigating the functions of unknown proteins (Jumper and Hassabis 2022).

In this context, the search for novel antimicrobial agents has seen rapid advancement. Simultaneously, the emergence of microbial resistance to conventional antibiotics and the rise of new pathogens unresponsive to existing treatments have intensified the quest for alternative solutions. Among these, antimicrobial peptides (AMPs) have emerged as especially promising candidates (Islam et al. 2024; Ji et al. 2024; Alzain et al. 2025; Oliveira Júnior et al. 2025; Pérez de la Lastra et al. 2025; Roque-Borda et al. 2025; Zhang 2025). The AMP database (APD3, http://aps.unmc.edu/AP/) as of December 2024 contains 5099 peptides, including 3306 natural (410 from bacteria, 5 from archaea, 8 from protists, 29 from fungi, 268 from plants, and 2580 from animals), 1299 synthetic, and 231 predicted AMPs (Wang and Wang 2004; Wang et al. 2009; Wang et al. 2016). A variety of biological activities have been associated with these peptides, including antibacterial, antiviral, anticancer, immune regulation and prevention of biofilm formation (Mookherjee et al. 2020; Qu et al. 2023; Loffredo et al. 2024; Polat et al. 2024; Varela-Quitián et al. 2025). Compared to traditional antibiotics, they present the advantages of low cytotoxicity to eukaryotic cells, high thermal stability and good solubility (Luo and Song 2021). Moreover, thanks to their manyfold mechanisms of action, it is difficult for microbes to develop resistance against AMPs (Gao et al. 2021). These advantages may be further optimised through the design and synthesis of novel peptides inspired by natural counterparts, employing molecular biology techniques and leveraging artificial intelligence programmes for guidance.

AMPs are generally composed of 10–50 amino acids, but also include longer members, although with molecular weights lower than 10 kDa. Most AMPs are positively charged (2–13 Lys/Arg residues). Hydrophobic amino acids play an important role in their antibacterial activity, providing regions for easy interactions with the cellular membrane. In fact, AMPs lacking hydrophobic residues usually show weaker affinity to the membrane lipids, while the members with high hydrophobicity, such as brevipeptides, often remain attached to the membrane for longer times (Bin Hafeez et al. 2021). The term AMPs includes several peptide families, different in size, three-dimensional folding, hydrophobicity and isoelectric point. Defensins represent perhaps the most important and certainly the best studied group of AMPs. They are present in all living organisms, mainly in plants, insects, vertebrates and, to a minor extent, in fungi and other organisms (van der Weerden and Anderson 2013; Kovaleva et al. 2020; Gao et al. 2021; Di Somma et al. 2022; Gao and Zhu 2024; Scieuzo et al. 2024; Zhao et al. 2024; Nagib et al. 2025; Defensin Database: http://defensins.bii.a-star.edu.sg/). Despite their common name, defensins can be very different in structure depending on the sub-type and on the phylum in whose species are they expressed.

In this work we report on the discovery of a new family of genes encoding small peptides, that we can tentatively classify as antimicrobial agents, in many fungal species, but also in some arthropods, where their presence is likely the result of horizontal gene transfer (HGT). Based on their predicted structure, we name these molecules “Hairpin Loop Peptides” (HLPs).

## Results and Discussion

### Discovery of hairpin loop peptides (HLPs)

While screening the results of a transcriptome project on the antennae of the pea aphid *Acyrthosiphon pisum*, we came across some sequences encoding small proteins of 86 amino acids, including a signal peptide. The unique specular symmetry of the six conserved cysteines attracted our attention and immediately suggested a very compact structure stabilised by three disulphide bridges which could easily form between facing cysteines when the polypeptide chain was half-folded. A BLAST search through the protein database returned hundreds of entries in Fungi, a significant number in some insect species, only a few in other arthropods and in other phyla. Notably, they are absent in plants. Several classes of antimicrobial peptides have been described in Fungi and other organisms, as reported in the Introduction, but none of them resembles the elegant symmetrical and highly compact structure of HLPs.

### Phylogenetic distribution of HLPs in Fungi

To shed light on the function of these peptides we first explored their occurrence across the fungal kingdom. Therefore, we performed a BLAST search using one of the HLPs of *Aspergillus ruber* (XP_040642819.1) as a query against the protein database of NCBI and limiting our search to Fungi. While we found these peptides in a large number of species, they could only be detected, quite unexpectedly, in a few selected clades. In particular, most of the hits (about 800) came from species of the sub-phyla of Pezizomycotina and Agaricomycotina, which include several important edible mushrooms, about 150 from Glomeromycota, Mucoromycota, Mortierellomycota and a few from other phyla and sub-phyla. Figure 1A reports a partial and simplified phylogenetic tree of Fungi adapted from the classification of Strassert and Monaghan (2022), and limited to the clades where HLPs were found, together with the numbers of these peptides identified in each branch. The great majority of species express only one to five HLPs, generally 2-4, and only a few were endowed with larger numbers, up to 20 in exceptional cases. Outside these clades, we could not find HLPs when we performed a BLAST search limited to Fungi while excluding the phyla and sub-phyla reported in red font in Figure 1A. The numbers of entries were then confirmed by novel BLAST searches, each limited to a single clade. Supplementary Files S1-S6 report all the amino acid sequences found in different phyla and sub-phyla of Fungi. Particularly interesting is the presence of HLPs in the phyla of Chytridiomycota and Rozellomycota, regarded to contain some of the most primitive fungal species (South et al. 2024).

**Figure 1.**
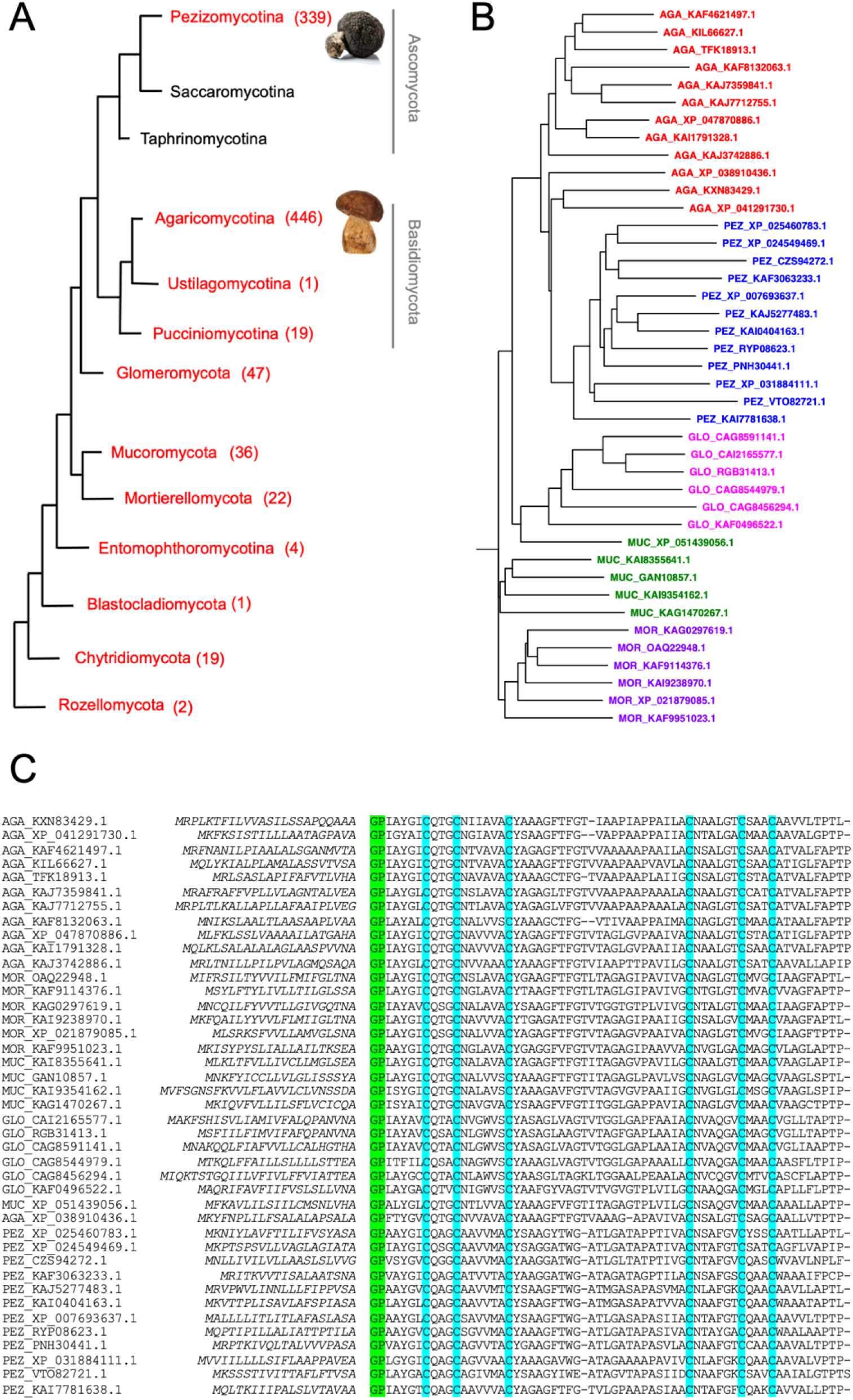
HLPs of Fungi. (**A**) Based on NCBI BLAST searches, HLPs were only found in the clades reported in red font with their numbers in brackets. The tree is a modified section from Strassert and Monaghan (2022), and reports only the branches were HLPs were found. (**C**) Alignment and (**B**) relative phylogenetic tree of selected HLPs representatives of two subphyla and three phyla where most of these peptides have been identified. Signal peptides are in italics. Conserved cysteines are highlighted in blue. All mature sequences start with GP (highlighted in green). AGA: Agaromycotina; MOR: Morteriellomycota; MUC: Mucoromycota; GLO: Glomeromycota; PEZ: Pezizomycotina. The selected sequences are those reported in bold font in Supplementary Files S1-S5. The sequences belonging to the five different phyla and sub-phyla segregate into separate clades of the phylogenetic tree.

The reason why such peptides are only expressed in some specific taxa cannot be simply explained by differences in genome sequencing information, nor on the basis of their phylogenetic distances. In fact, as an example, HLPs are absent from species of the subphylum of Saccharomycotina, whose genomes were among the first to be sequenced; moreover, such species share the phylum of Ascomycota with Pezizomycotina where HLPs are highly represented. Assuming a likely function as antimicrobial agents for these polypeptides, we could suggest that perhaps environmental factors could be in part responsible for such a patchy distribution. Under such perspective, it is well known that Fungi, particularly those interacting with plants through mycorrhizal associations host large amounts and variety of bacterial communities, thus establishing a symbiotic continuum between plant, fungus and bacteria in which carbon is transferred top-down, while mineral nutrients are transferred down-top (Duan et al. 2024). It is reasonable then to assume that soil Fungi interacting with bacterial communities might have developed weapons to fight threatening/competing microbial species, as part of their innate immune-like system (Venice et al. 2020; Daskalov 2023). On the other hand, the large number of different antimicrobial agents reported in the literature might indicate that species belonging to different phylogenetic clades may have developed different molecular tools after branching of such clades from their common origin.

In Figure 1C some representative sequences from each of the main five clades (AGA: Agaricomycotina, MOR: Morteriellomycota, MUC: Mucoromycota, GLO: Glomeromycota; PEZ: Pezizomycotina) are aligned, with the relative phylogenetic tree shown in Figure 1B. We can observe the conserved motif of six cysteines and a strong predominance of hydrophobic residues (on the average 35-45 in the mature polypeptide, corresponding to about 58-75%), while only one or two (or none) charged amino acids are present. Moreover, all HLPs contain a signal peptide of 15-20 residues, revealing their secretory nature. With only a couple of exceptions, calculated isoelectric points are very close to neutrality with values between 6.8 and 7.2, being different in such characteristics from other AMPs, particularly defensins, which are strongly positively charged. The phylogenetic tree (Figure 1B) shows that the selected sequences segregate into five clades, each relative to one of the phyla or sub-phyla considered. Similarities between sequences within each group are however not very high, with 40 to 60% of identical residues in most cases.

### Structure

The AlphaFold database contains several predicted three-dimensional structures of HLPs. Figure 2A shows the model of the *Valsa mali* (also known as *Cytospora mali*) member (sub-phylum: Pezizomycotina; GenBank: KUI64379.1, PDB: A0A194VJR5). The model, which is very similar to those of other HLPs, confirms the presence of three disulphide bridges between the specular symmetric cysteines which bring together two α-helical segments connected by a non-structured loop of 15 residues. A starting glycine and a proline at the two ends of the chain are also conserved in most HLPs.

**Figure 2.**
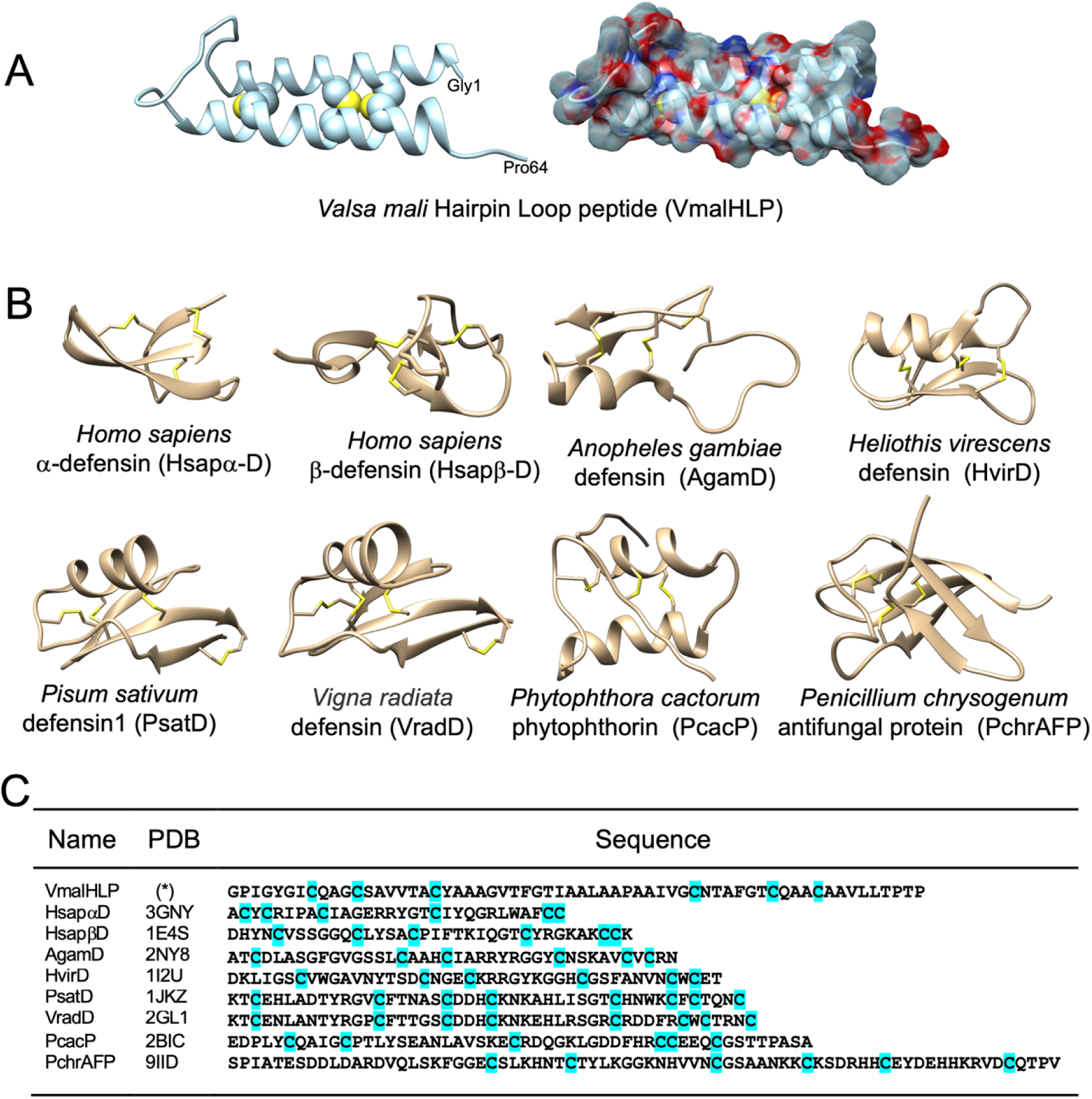
(**A**) AlphaFold three-dimensional model of a typical fungal HLP (*Valsa mali*, Agaricomycotina; GenBank: KUI62490.1, PDB: A0A194VJR5). Gly1 and Pro64 mark the origin and the end of the mature sequence. (**B)**. Predicted three-dimensional folding of typical members of AMPs, including six defensins, a member of the small phytophthorin family and a member of antifungal proteins (AFPs). They strongly differ in amino acid sequences (**C**) and in their cysteine patterns; (*) see above.

When observed in a space-filling model, the shape of HLP leaves no room for cavities and presents itself as a rigid rod with a larger head and tapering at the other end where the two termini of the peptide chain come close to each other. Its total length is about 44 Å, while its section at the middle of the rod is between 10 and 15 Å. The random-coil loop connecting the two helical domains is too small for a cavity or a binding site. Instead, the rigid and compact structure gives such peptides the appearance of molecular nails able to penetrate the cell membrane, also thanks to the hydrophobic nature of most of their amino acids.

### Comparison with other antimicrobial peptides

When compared with the structures of some AMPs from different organisms (Figure 2B), we can appreciate the striking difference between the elongated, rod-like, shape of HLPs, generated by the nested connection of the symmetrical cysteines, and the roughly round folding of all other AMP families produced by interlocked disulphide bridges between irregularly positioned cysteines. All sequences contain 6 or 8 cysteines paired in 3-4 disulphide bridges (Figure 2C). They include six defensins, the most abundant and best studied family of AMPs (two from humans, two from insects and two from plants), a member of a small group of AMPs called phytophthorins (PcacP) and a member of antifungal proteins (PchrAFPs). Phytophthorins have been reported only in species of the genus *Phytophthora*, plant pathogens belonging to the order of Peronosporales, phylum of Oomycota (Orsomando et al. 2001; Nicastro et al. 2009). A Blast search of the NCBI protein database returned only 22 hits, all from species of the same genus, with 10 of them from the same species (*P. infestans*), even when using highly flexible Blast parameters (threshold 10,000; word 2). Phytophthorins resemble other AMPs for their small size, the presence of several cysteines and their disulphide bridges that constrain the peptide chain into a double folded shape. Their isoelectric points vary across a wide range from 4 to 8 (Supplementary File S7). AFPs, instead, have been reported in several species of fungi (Hegedüs and Marx 2013; Tong et al. 2020; Wang et al. 2025). Our Blast search found 99 members of this family, all in Fungi and none in other living organism (Supplementary File S7). All sequences belong to species of the Pezizomycotina sub-phylum, most of them to the genera of *Aspergillus*, *Fusarium* and *Penicillium*.

The roughly round shapes of all other AMPs as compared with the elongated structure of HLPs are clearly due to the presence of one or more interlocked disulphide bridges. This appears clear when comparing the interlinked topology of cysteine pairing, although not conserved in different groups of AMPs, with the nested symmetrical arrangement of disulphide bridges of HLPs, as shown in Figure 3. Defensins of mammals and insects, as well as phytophthorins and antifungal proteins, generally contain six cysteines connected by three disulphide bridges in various fashions, while plant defensins present an additional bridge connecting the two ends of the polypeptide chain.

**Figure 3.**
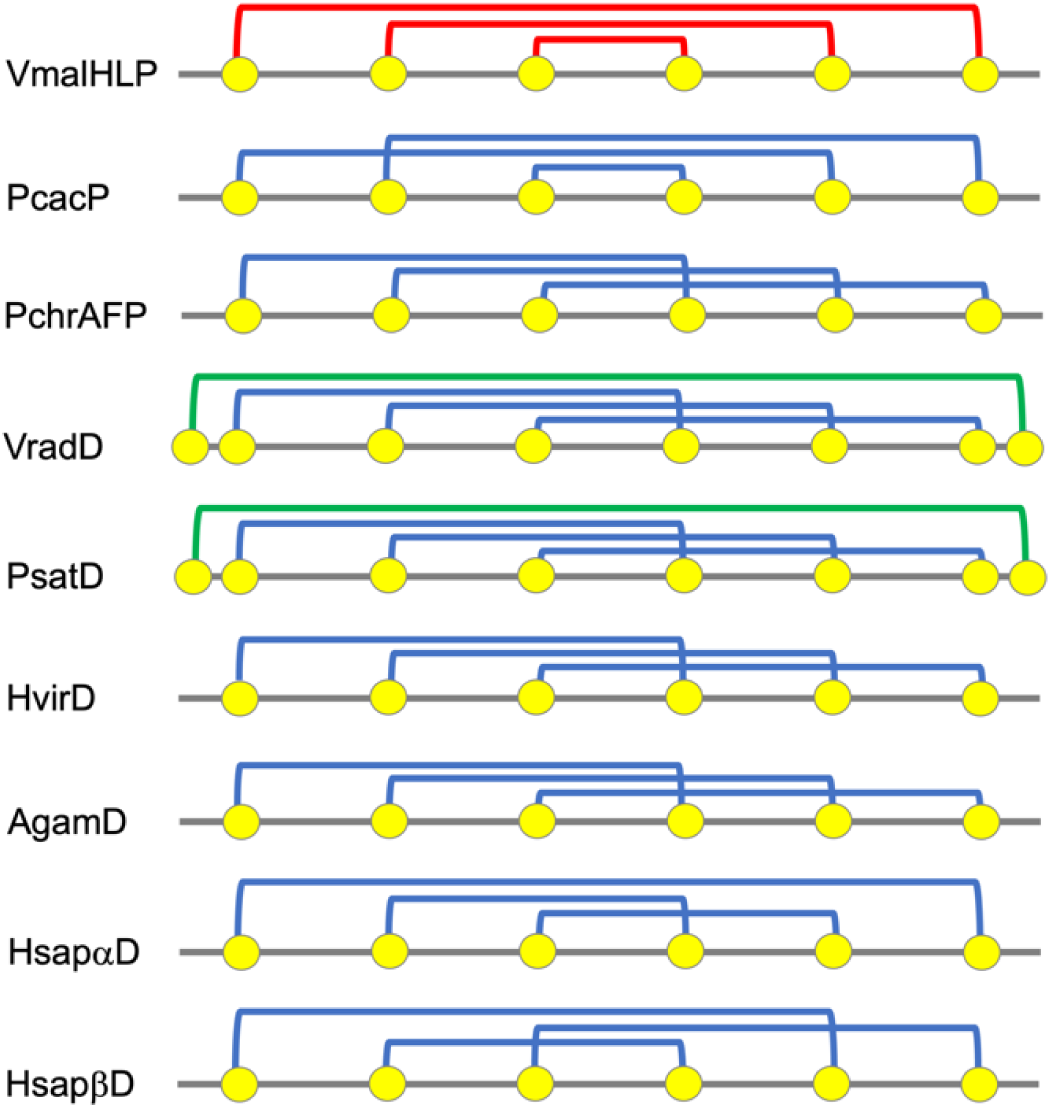
Topology of disulphide bridges in different types of AMPs. Generally, six cysteines are paired into three disulphide bridges (blue lines), except for the plant defensins, which present a fourth bridge connecting two additional cysteines located at both ends of the polypeptide chain (green lines). The topology of disulphide bridges of HLPs, exemplified by the *V. mali* member (red lines), is unique in being specular symmetric and responsible for the elongated shape of these peptides. Names are as in Figure 2.

### HLPs in arthropods

Having established the presence of HLPs in many fungi species, we returned to the starting point of our discovery asking whether aphids were endowed with genes encoding HLPs, or their presence in these insects could only be regarded as a consequence of fungal contamination. Therefore, we applied a BLAST search to the pea aphid genome using the HLP sequences originally detected in the antennal transcriptome as queries, and found the same five members, thus confirming that their genes are part of the pea aphid genome. Further evidence is given by the fact that four of them are located on chromosomes A1 and A3, as reported in Table 1.

**Table 1.** HLP sequences found in the genome of the pea aphid *Acyrthosiphon pisum*.

| Protein Seq | Genome Seq | Chromosome | from | to | Frame |
| --- | --- | --- | --- | --- | --- |
| XP_003248175.1 | NW_021763214.1 | Unplaced | 8747 | 9004 | +2 |
| XP_029343601 | NC_042494.1 | A1 | 90275747 | 90276004 | +2 |
| 016657431.1 | NC_042496.1 | A3 | 2770789 | 2771046 | +1 |
| 003247138.1 | NC_042494.1 | A1 | 90257858 | 90258115 | +2 |
| 016657429.1 | NC_042496.1 | A3 | 2985902 | 2985645 | -3 |

Then, to estimate how different the expressed peptides were from those of Fungi, we performed a BLAST search using the five aphid sequences against the Fungi protein database and collected only the three most similar hits for each of them, obtaining a total of eight fungal sequences. These are aligned in Figure 4, which also shows a phylogenetic tree where the two groups of HLPs are segregated into two different clades. In fact, similarities between sequences of the same group were 84-87% for pea aphid and 51-72% for Fungi, whereas these values dropped to 39-56% when sequences of different groups were compared. This suggests that HLPs have been part of the aphid genome long enough for undergoing duplication and differentiation.

**Figure 4.**
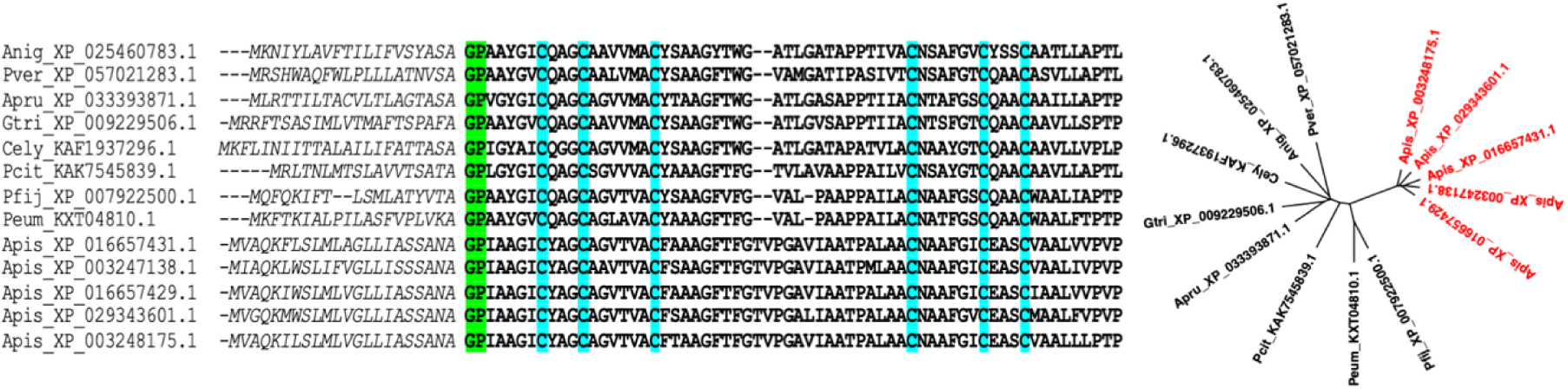
Alignment of the five HLPs of the pea aphid (Apis) with the eight most similar fungal HLPs. The phylogenetic tree shows clear segregation of the two groups of peptides into separate clades. Pfij: *Pseudocercospora fijiensis*, Pcit: *Phyllosticta citricarpa*, Apru: *Aplosporella prunicola,* Peum: *Pseudocercospora eumusae*, Cely: *Clathrospora elynae* Gtri: *Gaeumannomyces tritici,* Anig: *Aspergillus niger,* Pver: *Penicillium verhagenii*. The *A. pisum* members are shown in red in the phylogenetic tree, those of Fungi in black. The sequences used for this alignment are reported in Supplementary File S8.

Next, we extended our search to all Hexapoda. After refining the results of a BLAST search, we obtained 54 HLP members in several species of different orders. We then searched each of these sequences in the relative specie’s genome. The results are summarised in Table 2. For most of the 54 HLPs we could find the entire amino acid sequence which was 100% identical with that found in our first search. For 11 of them (indicated with a “p” in Table 2) we only detected at least half of the mature peptide, which however was 100% identical with the corresponding part of our query. We can also notice that most of the insect HLPs are found in species of the order of Hemiptera, which have lost the immune deficiency (IMD) pathway and lack the AMP genes (Gerardo et al. 2010; Ma et al. 2022).

**Table 2.** Species of Hexapoda containing in their genomes genes encoding HLPs. In some cases, the sequences have been localised to specific chromosomes (Chr), in all other cases they resulted from sequencing of the whole genome. Partial sequences, indicated by “p”, have been included only if more than half of the sequence was reported.

| Species | Seqs | Chr |
| --- | --- | --- |
| **Hemiptera** |  |  |
| *Apis Acyrthosiphon pisum* | 5 | A1,A3 |
| *Agos Aphis gossypii* | 1 | 2 |
| *Acra Aphis craccivora* | 1,1p | - |
| *Aluc Apolygus lucorum* | 2 | - |
| *Btab Bemisia tabaci* | 1 | 6 |
| *Cced Cinara cedri* | 1 | - |
| Dvit *Daktulosphaira vitifoliae* | 1 | - |
| Dcit *Diaphorina citri* | 1 | - |
| Dnox *Diuraphis noxia* | 1 | - |
| Meup *Macrosiphum euphorbiae* | 3 | 3,4 |
| Msac *Melanaphis sacchari* | 1 | - |
| Mdir *Metopolophium dirhodum* | 1 | 8 |
| Mper *Myzus persicae* | 2 | - |
| Pcor *Parthenolecanium corni* | 1p | - |
| Rmai *Rhopalosiphum maidis* | 1 | 1 |
| Rfus *Rhynocoris fuscipes* | 1 | 12 |
| Sher *Semiaphis heraclei* | 1 | 3 |
| Ufor *Uroleucon formosanum* | 1 | - |
| **Hymenoptera** |  |  |
| Agif *Aphidius gifuensis* | 1p | - |
| Bkin *Belonocnema kinseyi* | 1 | 6 |
| Dsim *Diprion similis* | 1 | 11 |
| Fari *Fopius arisanus* | 1 | - |
| Lbou *Leptopilina boulardi* | 1 | 4 |
| Lhet *Leptopilina heterotoma* | 1 | - |
| Nfab *Neodiprion fabricii* | 1 | 6 |
| Nlec *Neodiprion lecontei* | 1 | 6 |
| Tkay *Trichogramma kaykai* | 2 | - |
| **Diptera** |  |  |
| Crip *Chironomus riparius* | 1 | 3 |
| Cmar *Clunio marinus* | 1p | - |
| Cson *Culicoides sonorensis* | 1p | 2 |
| **Coleoptera** |  |  |
| Cass *Ceutorhynchus assimilis* | 3 | 4,4,4 |
| Csep *Coccinella septempunctata* | 1p | 4 |
| Cmon *Cryptolaemus montrouzieri* | 1 | - |
| Hvig *Henosepilachna vigintioctopunctata* | 1 | - |
| **Entomobryomorpha** |  |  |
| Fcan *Folsomia candida* | 1 | - |
| Odal *Orchesella dallai* | 1 | - |
| **Blattodea** |  |  |
| Bger *Blattella germanica* | 2 | - |
| **Thysanoptera** |  |  |
| Ffus *Frankliniella fusca* | 1p | - |
| Focc *Frankliniella occidentalis* | 1 | 2 |
| Musi *Megalurothrips usitatus* | 1p | 1 |
| Tpal *Thrips palmi* | 1 | - |

The 54 sequences were then used in a BLAST search against the Fungi protein database, downloading only the first hit for each hexapodan species. This search provided 26 fungi HLPs, which are reported together with the 54 hexapodan members in Supp. File S9. The alignment of all 80 sequences afforded the phylogenetic tree of Figure 5. We can observe that the proteins segregate into two distinct clades, one containing only insect HLPs, the other containing all the Fungi HLPs together with some insect members.

**Figure 5.**
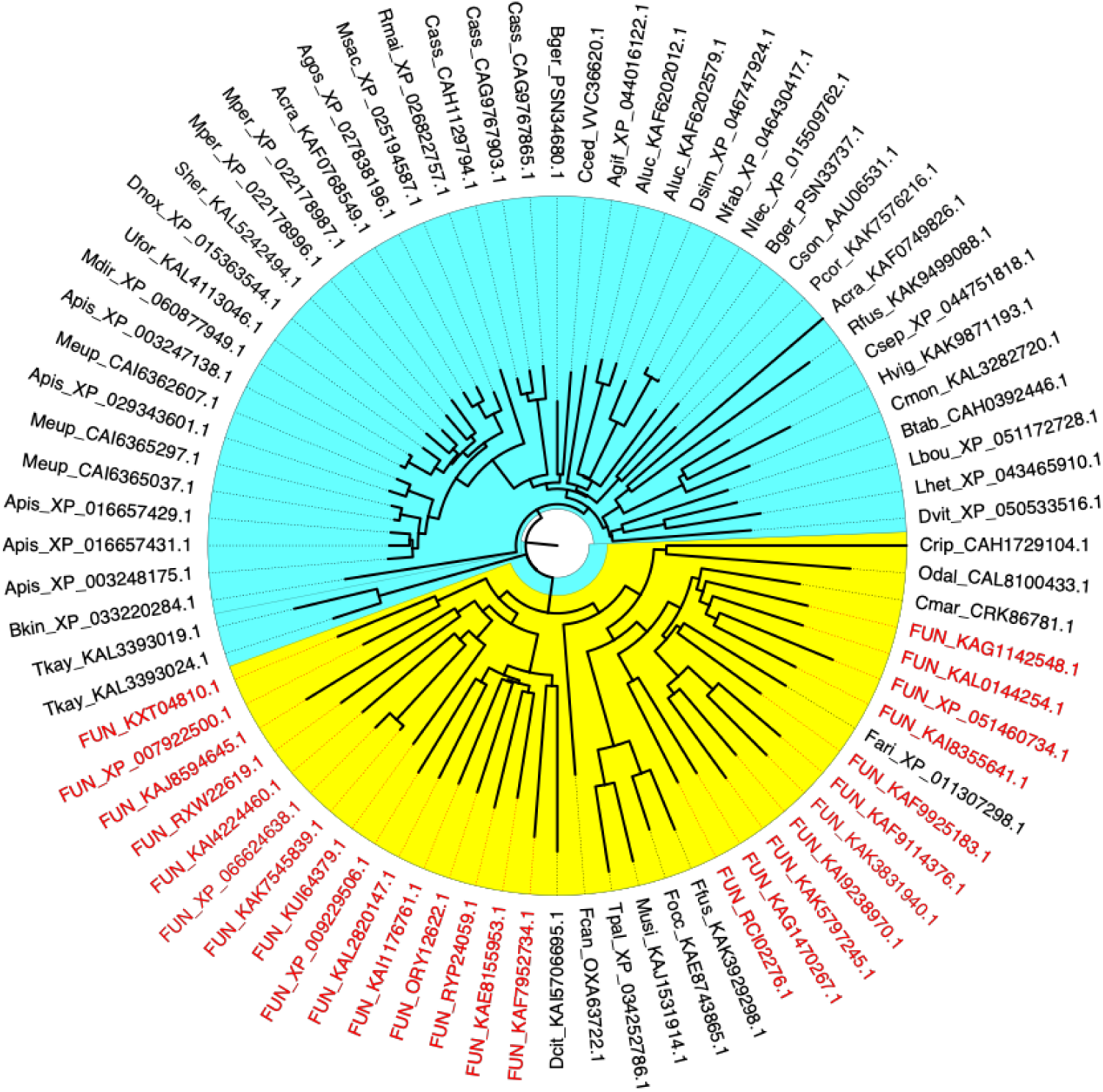
Alignment of the 54 HLPs found in hexapodan species (black; codes are those reported in Table 2) and their 26 most similar peptides in Fungi (red; FUN). We can observe two distinct clades, one containing only insect HLPs, the other with all the fungi peptides and some insect members. All sequences are reported in Suppl. File S9.

The insect HLPs clustering with the Fungi peptides do not present any special character which could differentiate them from those forming a separate cluster. It is true that most of them belong to primitive insects (Thysanoptera and Entomobryomorpha), but representative members of Hemiptera, Hymenoptera and Diptera are also included. We cannot exclude that in these species the gene transfer from Fungi may have occurred more recently, leaving shorter times for differentiation.

Next, we extended the quest for HLPs to all other arthropods. We performed two separate BLAST searches using as queries the fungal *A. ruber* HLP (XP_040642819.1) and the five aphid (*A. pisum*) members previously identified. We only collected 23 sequences, most of them in crustaceans of the genus *Daphnia*, including some members very similar to each other, possibly the results of gene duplication or sequencing errors (Supplementary File 10). Nevertheless, we confirmed at least for *Daphnia* species, the occurrence of HLPs in the relative genomes (Table 3).

**Table 3.**
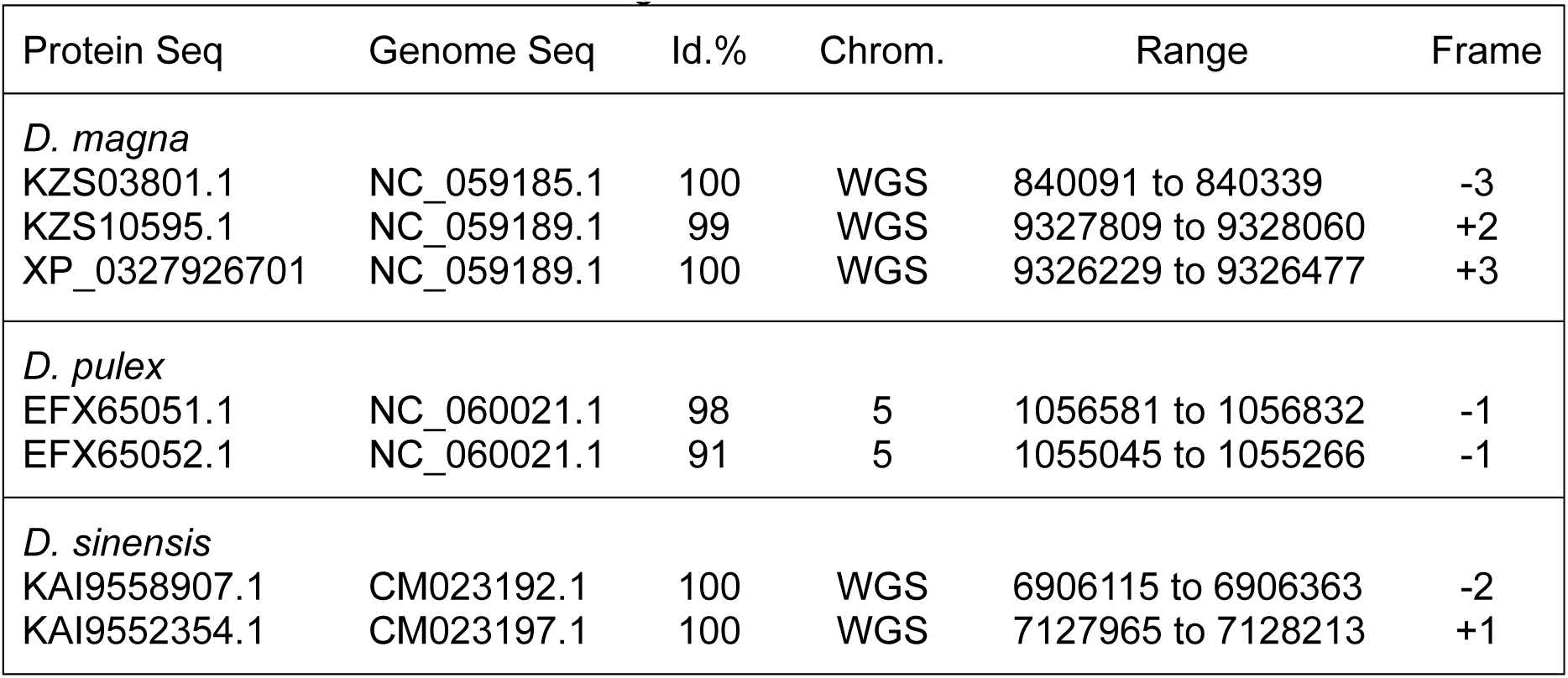
Sequences of HLPs found in the genomes of *Daphnia magna*, *D. pulex* and *D. sinensis*. WGS: Whole Genome Shotgun.

Finally, we looked at all other living organisms and performed a BLAST search excluding Fungi and arthropods and using the *A. ruber* HLP as a query. Our exploration only returned 17 sequences among nematodes and 102 sequences scattered in different *phyla* of invertebrates, including Gyrista, Chlorophyta, Rotifera, Cnidaria and others (Suppl. File S11). We also searched the genomes of each of the above species for the presence of the identified HLPs. In several cases, we could detect a sequence in the genome which was 100% identical (or for a couple of them 99% identical) with the query sequence. However, apart from only two cases, no indication on its position on the chromosome could be found, as the sequencing had always been performed on the whole genome shotgun. Table 4 summarizes these results.

**Table 4.** Species other than Fungi and Arthropoda containing in their genomes genes encoding HLPs. Only in two cases, the sequences have been localised to specific chromosomes, otherwise they were identified from the whole genome shotgun (WGS). Generally, the entire sequence was found in the genome, but we also included a few hits which were at least 99% identical to the mature sequence of the query.

| Species | Seqs | Chr. |
| --- | --- | --- |
| **Oomycota** |  |  |
| Pbel *Peronospora belbahrii* | 1 | WGS |
| Peff *Peronospora effusa* | 1 | 17 |
| Pale *Phytophthora aleatoria* | 2 | WGS |
| Pcac *Phytophthora cactorum* | 1 | WGS |
| Pcap *Phytophthora capsici* | 1 | 20 |
| Pcit *Phytophthora citrophthora* | 1 | WGS |
| Pfra *Phytophthora fragariae* | 1 | WGS |
| Pida *Phytophthora idaei* | 2 | WGS |
| Pmeg *Phytophthora megakarya* | 2 | WGS |
| Pole *Phytophthora oleae* | 2 | WGS |
| Prub *Phytophthora rubi* | 2 | WGS |
| Psoj *Phytophthora sojae* | 2 | WGS |
| **Nematoda** |  |  |
| Aave *Aphelenchoides avenae* | 1 | WGS |
| Hsch *Heterodera schachtii* | 3 | WGS |
| Htri *Heterodera trifolii* | 9 | WGS |
| **Cnidaria** |  |  |
| Dper *Desmophyllum pertusum* | 3 | WGS |
| Edia *Exaiptasia diaphana* | 1 | WGS |
| Nvec *Nematostella vectensis* | 1 | WGS |
| Plob *Porites lobata* | 1 | WGS |
| **Rotaria** |  |  |
| Aric *Adineta ricciae* | 2 | WGS |
| Aste *Adineta steineri* | 3 | WGS |
| Avag *Adineta vaga* | 1 | WGS |
| Bcal *Brachionus calyciflorus* | 2 | WGS |
| Rsp *Rotaria sp. Silwood2* | 1 | WGS |
| **Gyrista** |  |  |
| Spro *Scytosiphon promiscuus* | 1 | WGS |

To summarise these findings, Figure 6 illustrates the distribution of HLPs across various living organisms. Notably, these peptides have not been identified in plants, bacteria, or other procaryotes, as confirmed by targeted BLAST searches. The origins of these peptides remain ambiguous, and their patchy presence in only a limited number of species within each phylum raises challenging questions that may persist even when more sequence data will become available. In any case, the sporadic sequences found in insects and other animal species might reasonably be the result of horizontal gene transfer (HGT) events from Fungi that are known to be associated to many different organisms (Hixson et al. 2024; Michalik and Szklarzewicz 2025). Events of HGT, well described in bacteria (Nikolaidis et al. 2014; Acar Kirit et al. 2022; Wen and Herman 2024; Mishra and Lercher 2025; Zhaxybayeva and Nesbø 2025), have been more recently documented in increasing numbers of studies and with better evidence also in insects and other species (Chen et al. 2024; St Leger 2024; Wang et al. 2024; Colinet et al. 2025; García-Lozano and Salem 2025; Kirsch et al. 2025; Zhang et al. 2025).

**Figure 6.**
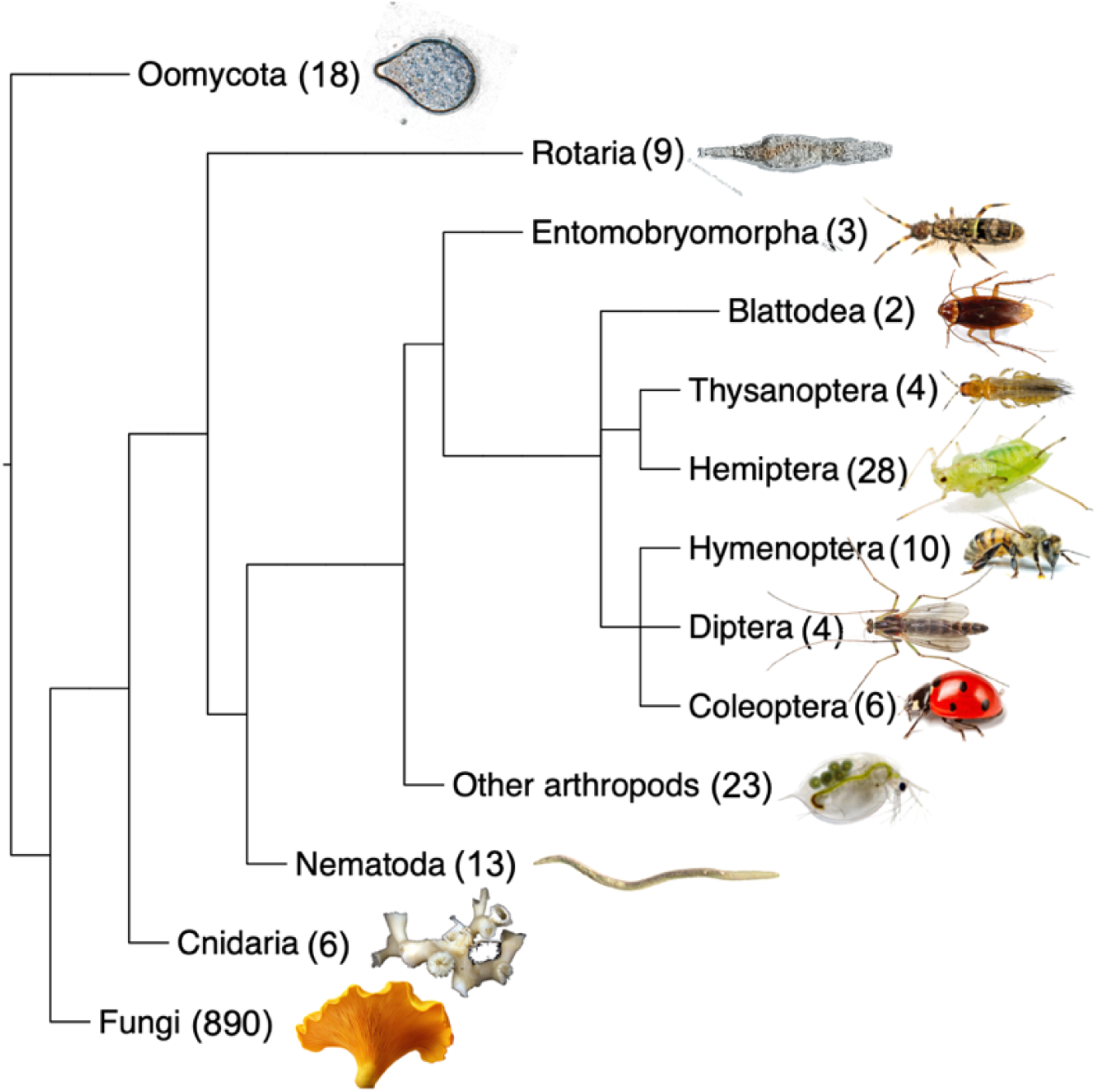
Occurrence of HLPs in different living organisms. These peptides have not been found in plants, bacteria and other procaryotes. Most of them are expressed in fungi, while the few scattered genes detected in the genomes of other species are likely the result of horizontal gene transfer events. In brackets the numbers of genes confirmed in the relative genomes.

In particular, the first transfer of genes from Fungi to aphids to be identified reported the occurrence of a red pigmentation in some populations of the pea aphid *A. pisum* and the potato aphid *M. persicae* due to acquisition of genes encoding a carotenoid desaturase from several Fungi (Moran and Jarvik 2010). Such genes became then duplicated within the aphid genome giving rise to individuals with stable colour in parthenogenetic clones. In particular, it was demonstrated that the transfer could not have occurred from the obligate bacterium symbiont *Buchnera aphidicola*, as the pea aphid genome does not contain any gene of *Buchnera* origin (Shigenobu et al. 2000; Consortium 2010; Nikoh et al. 2010; Shigenobu and Wilson 2011). Since this first work on HGT from fungi to aphids, several papers reported the acquisition of genes by insects from plants and microbes (Dai et al. 2021; Xia et al. 2021; Gilbert and Maumus 2022), but only a couple from Fungi (Roelofs et al. 2013; Wang et al. 2023).

### Prediction of antimicrobial properties

Several structural elements of HLPs indicate these polypeptides as potential antimicrobial agents. Their small size, the highly compact and needle-like shape, together with the presence of a large number of hydrophobic residues, make these molecules perfect weapons to penetrate and damage the cell membrane. Several software programmes have been developed to predict the activity of small peptides against different types of pathogens (Lira et al. 2013; Toropova et al. 2018; Wang et al. 2022; Bournez et al. 2023). However, these predictive approaches primarily rely on statistical evaluations based on similarities with known antimicrobial peptides (AMPs). Consequently, when these tools are applied to HLPs – whose three-dimensional structures differ significantly from those of established AMPs – the reliability of their predictions may be limited.

Nevertheless, we tested, as representative HLPs, the 41 fungal HLPs reported in Figure 1 and the 5 sequences from *A. pisum* (Figure 4 and Table 1) with established software tools for predicting antimicrobial activity. The evaluation considered parameters such as size, charge, and hydrophobicity, aiming to provide initial insights into the antimicrobial capabilities of these peptides. The antimicrobial activity was predicted using the on-line software CAMPR3 (http://www.camp3.bicnirrh.res.in/prediction.php), applying different classifier algorithms. The 41 fungi HLPs were all classified as antimicrobial by the Support Vector Machine (SVM) with scores higher than 0.988. The Random Forest programme rated all of them as active with scores between 0.52 and 0.83, except for sequence KAJ5277483.1 (score 0.473). Similar results were obtained with the Artificial Neural Network (ANN) classifier, where only the above sequence did not pass the test. Finally, the Discriminant Analysis Classifier (DAC) predicted antimicrobial activity for the same sequences with scores higher than 0.625, except for the above sequence (score 0.379) and for sequence KAF0496522.1 (score 0.473). The 5 *A. pisum* HLPs were also predicted as antimicrobial agents by Support Vector Machine (SVM) all with scores of 1.000 and by the Random Forest programme with scores between 0.546 and 0.635. With the other two software (ANN and DAC), only sequence XP_029343601.1 was rated as non-antimicrobial by the Discriminant Analysis classifier (score 0.495). The antifungal properties of HLPs were evaluated using the Antifp software (https://webs.iiitd.edu.in/raghava/antifp), applying a threshold of 0.5. Out of the 41 representative fungal HLP sequences, only 16 were predicted to possess antifungal properties, while none of the five *A. pisum* proteins met the criteria for antifungal activity. In summary, we can reasonably conclude that HLPs might be potential antimicrobial agents, but their specific targets can only be identified after detailed experimental investigation.

## Conclusions and perspectives

Using a bioinformatic approach, we have identified genes encoding peptides of about 85 amino acids in several species of Fungi, as well as in some arthropods and few other species. Most likely such molecules represent a new class of antimicrobial agents, although further speculation and conclusions should be supported by experimental evidence. With respect to other AMPs, they present some elements of interest and novelty:

1. Structure: their elegant and extremely stable folding is unique and different from all other AMPs. Due to the mirror symmetrical position of their six cysteines connected by three nested disulphide bridges, these peptides present the shape of molecular nails, ideal for penetrating the cellular membrane.
2. Phylogenesis: although they have likely originated in Fungi, being present also in some of the most primitive species, intriguingly, HLPs are only found in some clades without any phylogenetic relationship.
3. Evolution: we have also found some HLPs in certain insects, notably in Hemiptera, other arthropods and, to a lesser extent in other phyla. They are absent from plants. Their presence in those invertebrates is likely the result of Horizontal Gene Transfer (HGT) events. This fact is particularly interesting, as HGT events from Fungi to insects have only been reported in a very limited number of cases. The occurrence of HLPs in many hemipteran genera may be related to their less efficient immune system, compared to other insects, and the lack of AMPs.

Potential applications of HLPs as alternatives to antibiotics can be expected as for other classes of AMPs (Boparai and Sharma 2020; Li et al. 2022; Mabrouk 2022; Xuan et al. 2023; Jacobowski et al. 2025). In addition, HLPs, which are expressed in a variety of edible mushrooms, appear to be safe for human consumption, and could be employed without risk as food preservatives, as suggested for other AMPs (Rai et al. 2016; Xu et al. 2023; Singh et al. 2024; Bermúdez-Puga et al. 2025; Ramesh et al. 2025). Moreover, their small size and compact structure, suggesting refractiveness to proteolytic enzymes, makes them stable over long shelf periods.

The predicted antimicrobial activity of HLPs is likely attributed to their compact, nail-like structure. As such, it is plausible that amino acid substitutions within their sequences could be introduced without compromising their antimicrobial properties, and may even enhance their activity or improve selectivity against specific pathogens. Several strategies could be considered:

- Increasing Hydrophobicity: Replacing polar residues with hydrophobic ones could enhance their ability to disrupt lipid membranes.
- Altering Proteolytic Sites: Adjusting sites targeted by proteolytic enzymes could improve the stability and longevity of HLPs.
- Introducing Binding Sites: Incorporating binding sites for metals or other chemicals with potential toxicity for target pathogens could further augment their antimicrobial effect.

These approaches are expected to become better targeted and effective as experimental validation of HLP activity and clarification of their mechanism of action will be achieved.

## Methods

BlastP searches were performed using as queries the five HLPs identified in the transcriptome of *A. pisum* antennae or representative fungal HLPs or other members of the same family, as indicated in the Results section. Default parameters (Expected threshold: 0.05; Word size: 5) were adopted unless otherwise reported. Signal peptides were predicted using the on-line SignalP 6.0 software (https://services.healthtech.dtu.dk/services/SignalP-6.0/). Alignment of amino acid sequences was performed with the ClustalW programme (https://www.genome.jp/tools-bin/clustalw), using default parameters, and trees were generated with Neighbor-Joining method. Visualization of the trees was done with the programme FigTree, version 1.4.4 (https://github.com/rambaut/figtree/releases). The AlphaFold three-dimensional model of the *Valsa mali* (also known as *Cytospora mali*) HLP, Pezizomycotina, was available from the database (GenBank: KUI64379.1, PDB: A0A194VJR5). The three-dimensional structures of the other AMPs (Figure 2) were also all available in the databases. Prediction for antimicrobial activity was performed on line, using the software CAMPR3 (http://www.camp3.bicnirrh.res.in/prediction.php), as described in the Results section. Antifungal activity was predicted by the software Antifp (https://webs.iiitd.edu.in/raghava/antifp) setting the threshold at 0.5.

## Acknowledgements

We thank Paola Bonfante, University of Torino, Italy, and Krishna Persaud, University of Manchester, UK, for precious suggestions and critical reading of the manuscript.

## Author Contributions

P.P. designed the study and drafted the manuscript. B.W., J.Z. and P.P. performed sequence searches, data analysis and processing. All authors contributed to the final version of the manuscript.

## Data Availability

All the sequences used in this work are available as Supplementary Files.

