## Supplemental Files for "New potential antimicrobial peptides with amazing symmetrical structure in fungi and insects"

#### Supplementary Files

#### Supplementary File S1. Agaricomycotina HLPs: 446 sequences

>KAI0785557.1 hypothetical protein C8Q75DRAFT_775386 [Abortiporus biennis]
MNPRSMLFAGLAAMSAAPAAHAGLIAYGICQTGCNTLAVACYAGAGFTFGTVVASALAPPAILACNAALGTCSAACASVALLAPTP
>KAF9554637.1 hypothetical protein CPC08DRAFT_712746 [Agrocybe pediades]
MRINANLLSTVALALSGASMVSAGPIAYGLCQTDCNTVAVACYADAGFTFGTVAAADAPAAVLACNSALGTCSAKCASVTLLAPTS
>KAF9554638.1 hypothetical protein CPC08DRAFT_712747 [Agrocybe pediades]
MRINTNLLSTVALALSGASMVSAGPIAYGLCQTGCNTVAVACYAGAGFTFGTVVAAAAAPAAVLACNTALGTCSAMCASVALLAPIP
>KAF4620962.1 hypothetical protein D9613_001254 [Agrocybe pediades]
MRINANLLSTVALALSGASIVSAGPIAYGLCQTDCNTVAVACYAGAGFTFGTVAAADAPAAVLACNSALGTCSEKCASVTLLAPTTS
**>KAF4621497.1 hypothetical protein D9613_001246 [Agrocybe pediades]**
**MRFNANILPIAALALSGANMVTAGPIAYGICQTGCNTVAVACYAGAGFTFGTVVAAAAAPAAILACNSALGTCSAACATVALFAPTP**
>KAF8661385.1 hypothetical protein AX14_007266 [Amanita brunnescens Koide BX004]
MKLFSFVVPVAAVLLSATSVVQAGPISYAICQTGCNIVAVACYGAAGATFGTVAAPLAPPAILTCNAALGSCMALCAPLLIAPIP
>KAF8735275.1 hypothetical protein AX14_002369 [Amanita brunnescens Koide BX004]
MQLFLFNFILPFAAVLCVTTVVQAGPILYAMCKLGCDVVANACAASSSGGTFGTTGAPPASPAIAACKTALALCALLFPPGP
**>KIL66627.1 hypothetical protein M378DRAFT_23285 [Amanita muscaria Koide BX008]**
**MQLYKIALPLAMALASSVTVSAGPIAYGICQTGCNTVAVACYAAAGFTFGTVVAAPAAPVAVLACNAALGTCSAACATIGLFAPTP**
>KAF8335236.1 hypothetical protein F5887DRAFT_614114 [Amanita rubescens]
MRPSKLLLPIAVALSTTGIVTAGPIAYGVCQTACNAGAVTCYTGVGFTFGVTLVAAPPAILACNSILGACMALCTPFLIAPTP
>KAF8351847.1 hypothetical protein F5887DRAFT_4392 [Amanita rubescens]
MRLSKLLLPIAVALSSTGIVNAGPIAYGLCQTVCNFGAVACYAAAGFTFGTVAAPAAPPIILACNAAQGACMTLCAPLLVAPTF
>PFH45591.1 hypothetical protein AMATHDRAFT_158614 [Amanita thiersii Skay4041]
MRLTRVFAPLGIIVALSTVPQIVQAGPILYGICQTGCNSLAVICYAAGGVVFGTVLAATAPAAILACNAAQGSCMALCAVTVLPLPTP
>KAH9948988.1 hypothetical protein B0H21DRAFT_689192 [Amylocystis lapponica]
MKFSLLVPFALLAASANAGPIAYGICQTGCNTVAVACYAAAGVQFGTIAAPLAPATVLGCNTALGTCSAACATVCLLAPTP
>KAI0323098.1 hypothetical protein OF83DRAFT_1167113 [Amylostereum chailletii]
MKLSLIVALLAATAPTVFAGPIAYGICQTGCNVVAVACYAAAGATFGTVVAAAAAPAAILGCNAALGTCSSMCAGVALLAPTP
>KAI0323099.1 hypothetical protein OF83DRAFT_1090726 [Amylostereum chailletii]
MKLFSILTLLTLSTPAVIAGPIAYGICQTGCNVVAVACYAAAGATFGTVAAPAAPAAILGCNAALGTCSTMCASVALFAPTP
>KAI0930540.1 hypothetical protein AcV5_007225 [Antrodia cinnamomea]
MKFFLLSSLALAGVSLVNAGPIAYGLCQTGCNTVAVACYAAAGFQFGTVVAGPLAPATILACNAALGTCSAACAGVTLLAPTP
>THH27703.1 hypothetical protein EUX98_g6480 [Antrodiella citrinella]
MKFSALSALAVLATTPFVAGGPIAYGICQTGCNTLAVACYAAAGFTFGTVIAAPAAPAAILACNAGLGTCSAACATIGLFAPTP
>XP_028477985.1 hypothetical protein EHS24_006059 [Apiotrichum porosum]
MKAAIPLALAILASPVAAGPVAWGLCYTACNASYGVCLGALGLVAGTFTLGAGTPVAVVTCSVAQGACMSACSPILMAPTP
>KAK0445166.1 hypothetical protein EV421DRAFT_343310 [Armillaria borealis]
MRLSPILTFLVTSLALAPQAHAGPIAYGICQTGCNVLAVACYAAAGFTFGTVAAPAAPAAIIGCNSALGTCSAACASVALLAPTP
>KAK0435962.1 hypothetical protein EV421DRAFT_1145183 [Armillaria borealis]
MRLSPILTSLVMSLALAPQAHAGPIAYGICQTGCNVLAVACYAAAGFTFGTVAAPAAPAAIIGCNSGLGTCSAACASVALLAPTP
>KAK0445201.1 hypothetical protein EV421DRAFT_1902652 [Armillaria borealis]
MRLSRAFACLATSLVLAPQAYAGPIAYGLCQTGCNTMAVACYAAAGATFGTVVAAAATPAVILGCNVALGTCSATCATVGLFAPTP
>KAK0211506.1 hypothetical protein IW262DRAFT_1468047 [Armillaria fumosa]
MRLSPIFTFLVTSLTLAPQAYAGPIAYGLCQTGCNVVVVACYAAAGFTFGTVAAPAAPAAILGCNSALGTCSAVCASVALLAPTP
>KAK0211537.1 hypothetical protein IW262DRAFT_371639 [Armillaria fumosa]
MRLSRAFACLATFLVLVPQAYAGPIAYGICQTATFGTVVAAPAAPAVILACNAALGTCSATCATVALFAPTP
>PBK78949.1 hypothetical protein ARMGADRAFT_1093628 [Armillaria gallica]
MRLSRAFACLATSLVLAPQAYAGPIAYGICQTGCNTVAVACYAAAGFTFGTVIAAPAVPAVILTCNAALGTCSAACATVALFAPTP
>PBK94666.1 hypothetical protein ARMGADRAFT_61866 [Armillaria gallica]
MRLSPILTFLVTSLAIAPQVHAGPIAYGICQTGCNVVAVACYAAAGFTFGTVAAPAAPAAIIGCNTALGTCSAACASVALLAPTP
>PBK94627.1 hypothetical protein ARMGADRAFT_59951 [Armillaria gallica]
MRLSRAFACLATSLVLAPQAYAGPIAYGICQTGCNTMAVACYAAAGATFGTVVAAAAAPAAILACNAALGTCSTACATVGLFAPTP
>KAK0496260.1 hypothetical protein EDD18DRAFT_200744 [Armillaria luteobubalina]
MRLSPIFTFLATSLALAPQAYAGPIAYGICQTGCNVLAVACYAAAGFTFGTVAAPAAPAAIVACNSGLGTCSAACASVALLAPTP
>KAK0496299.1 hypothetical protein EDD18DRAFT_202440 [Armillaria luteobubalina]
MRLSRALACLATSLVLAPQAYAGPIAYGLCQTGCNTMAVACYAAAGVTFGTVVAAAATPAVILTCNASLGVCSATCATVALLAPTP
>KAK0192253.1 hypothetical protein F5146DRAFT_1039048 [Armillaria mellea]
MRLSRAFACLATSLVLAPQAYAGPIAYGICQTGCNTMAVACYAAAGATFGTIVAAAAAPVAILGCNAALGTCSATCATVALLAPTP
>KAK0192295.1 hypothetical protein F5146DRAFT_1136064 [Armillaria mellea]
MRLSPIFTFLVTSLALAPQAYAGPIAYGICQTGCNVLAVACYAAAGFTFGTVAAPAAPAAIIGCNSALGTCSAACASVALLAPTP
>KAK0232397.1 hypothetical protein EDD85DRAFT_956241 [Armillaria nabsnona]
MRLSPILTFLVTSLAIAPQARAGPIAYGICQTGCNVLAVACYAAAGFTFGTVAAPAAPAAIIGCNSALGTCSAACASVALLAPTP
>KAK0232409.1 hypothetical protein EDD85DRAFT_108986 [Armillaria nabsnona]

MRLSPILTFLVTSVALAPQAHAGPIAYGICQTGCNVVAVACYAAAGFTFGTVAAPVAPVAIIGCNSALGTCSAMCAGVALLAPTP
>KAK0232361.1 hypothetical protein EDD85DRAFT_956211 [Armillaria nabsnona]
MRLSRAFACLATSLVLAPQAYAGPIAYGICQTGCNTMAVACYAAAGATFGTVVAAAAAPAVILACNASLGTCSTACATVALFAPTP
>KAK0477555.1 hypothetical protein IW261DRAFT_265418 [Armillaria novae-zelandiae]
MRLQSPIFAFLFTALALAPQAYAGLIAYGICQTGCNVLAVACYSAAGFTFGTIAAPAAPAAIVGCNAALGSCSAICASVALLAPTP
>KAK0477562.1 hypothetical protein IW261DRAFT_1565876 [Armillaria novae-zelandiae]
MRLQSPIFAFLFTALALAPQAYAGPIAYGICQTGCNVLAVACYSAAGFTFGTVAAPAAPAAIVGCNAALGSCSTMCATVALLAPTP
>KAK0480870.1 hypothetical protein IW261DRAFT_1475037 [Armillaria novae-zelandiae]
MRLSPIFTFLVTSLALAPQAYAGPIAYGICQTGCNVVAVACYAAAGFTFGTIAAPVAPVAIIGCNTALGTCSAACATVALFAPTP
>KAK0480837.1 hypothetical protein IW261DRAFT_1474858 [Armillaria novae-zelandiae]
MRLSRAFACLATSLVLAPQAYAGPIAYGICQTGCNTMAVACYAAAGATFGTVIAAAAAPAAILGCNAALGTCSATCATVGLLAPTP
>SJL12908.1 uncharacterized protein ARMOST_16341 [Armillaria ostoyae]
MRLSRTFVCLATSLVFAPQAYAGPIAYGLCQTGCNAMAVACYAAAGATFGTVVAAAATPAVILGCNVALGTCSATCATVGLFAPTP
>SJL10225.1 uncharacterized protein ARMOST_13609 [Armillaria ostoyae]
MRLSPILTSLVTSLALAHQAHAGPIAYGICQTGCNVLAVACYAAAGFTFGTVAAPAAPAAIVGCNSGLGTCSAACATVALLAPTP
>PBK70936.1 hypothetical protein ARMSODRAFT_934503 [Armillaria solidipes]
MRLSPILTFLVTSLALAPQAHAGPIAYGICQTGCNVLAVACYAAAGFTFGTVAAPAAPAAIIGCNSGLGTCSAACATVALLAPTP
>PBK70903.1 hypothetical protein ARMSODRAFT_1017680 [Armillaria solidipes]
MRLSRAFACLATSLVLAPQAYAGPIAYGLCQTGCNTMAVACYAAAGVTFGTVVAAAAAPAVILGCNAALGTCSATCATVALLAPTP
>KAG6331624.1 hypothetical protein ID866_7469 [Astraeus odoratus]
MNLRCIAAYSLISLPLAMAGPLAYAACQTGCNAIVVACYAGAGFTFGVALPVAPPAILACNAALGTCMATCATIALAPTP
>EJD44225.1 hypothetical protein AURDEDRAFT_65424 [Auricularia subglabra TFB-10046 SS5]
MTLITSALLLAFAAPASASLILYGICQTGCNMGAVSCYGVACATFGTVVATPLTPAIILWCNAALGTCMATLCAPLLLIPLP
>TRM64791.1 hypothetical protein BD626DRAFT_489882 [Auriculariopsis ampla]
MRVTAILAPVALATAVAAGPIAYGICQTGCNTLAVACYAAAGFTFGVALPAAPPVILACNAGLGTCSAACATVALLAPTP
>KAH7096490.1 hypothetical protein BKA62DRAFT_662757 [Auriculariales sp. MPI-PUGE-AT-0066]
MKLIRPTRLIATTLVLLAPTQVRASLIAYGICQTGCNIGAVTCYAAAGFTFGTIAAPVAPLAILGCNAALGTCMAAMCAPLLLIPFF
>KAH7096489.1 hypothetical protein BKA62DRAFT_719469 [Auriculariales sp. MPI-PUGE-AT-0066]
MKVIRPTRLIASTLVLLAPTQVHAGLIAYGICQTSCNILAVACYGAAGAVFGTIAAPVAPPAILACNAALGTCMGAICAPLLLIPFI
>KAH7090935.1 hypothetical protein BKA62DRAFT_645562 [Auriculariales sp. MPI-PUGE-AT-0066]
MKFLHPITRLATALALLAPTQVRAGIIAYGLCRTGCNVIVMGCYGAAGAVFGTVLAVTASPTILACNGAQGLCMTTLFAPLLLTPVP
>KAI9568904.1 hypothetical protein HD554DRAFT_2021511 [Boletus coccyginus]
MNFKSLAALTLTASAAPFAAAGPLAYAICQTGCNVLAVSCYGAAGFTFGVTIVAAPPAIMACNAGLGTCMAACATAALFAPTP
**>KAF8132063.1 hypothetical protein EV363DRAFT_1329745 [Boletus edulis]**
**MNIKSLAALTLAASAAPLVAAGPLAYALCQTGCNALVVSCYAAAGCTFGVTIVAAPPAIMACNAGLGTCMAACATAALFAPTP**
>KAF8415388.1 hypothetical protein L210DRAFT_3464449 [Boletus edulis BED1]
MNFKSLAALTLAASAAPLAAAGPLAYALCQTGCNTLAVSCYAAAGFIFGVTIIGVPPAIMGCNAGLGTCMATCATVALFAPTP
>KAF8131977.1 hypothetical protein EV363DRAFT_1329574 [Boletus edulis]
MNFKSLAALTLAASAAPLAAAGPLAYALCQTGCNTLAVSCYAAAGFTFGVTIIGVPPAIMGCNAGLGTCMAACAVVGLFAPTP
>KAG6382117.1 hypothetical protein JVT61DRAFT_760 [Boletus reticuloceps]
MNFKSLAALTLAASAAPLAAAGPLAYAVCQTGCNTLAVSCYAAAGFTFGVTIVAVPPAIIGCNVGLGTCMAACAVVGLFAPTP
>KDQ08146.1 hypothetical protein BOTBODRAFT_139194 [Botryobasidium botryosum FD-172 SS1]
MRVFSLAAFVIAPFYLATGAYAGPIAYGLCQTGCNTLAVACYAAAGFQFGTVVAAAATPATILACNAGLGTCSATCATVALLAPTP
>KAG8220444.1 hypothetical protein J3R82DRAFT_3138 [Butyriboletus roseoflavus]
MNLKSLAALTLAASVPLASAGPLAYGLCQTASPRRPGCNALVVSCYAGAGFTFGVTVVGAPAAIIACNAGLGTCMAACAATALIAPIP
>RXW13811.1 hypothetical protein EST38_g12044 [Candolleomyces aberdarensis]
MRPSLLFVPVLAASVAQAGPIAYGICQTGCNAVVVACYAAAGFTFGTVAAPLAPPAIVACNTALGTCSAACATVALLAPIP
>RXW22619.1 hypothetical protein EST38_g3243 [Candolleomyces aberdarensis]
MRPSLLLIPVLAASTAQAGLIAYGICQTGCNAVTVACYAAAGFTFGTIAAPLAPPAIVACNAGLGTCSTACATVALLAPTP
**>XP_038910436.1 uncharacterized protein EI90DRAFT_3079712 [Cantharellus anzutake]**
**MKYFNPLILFSALALAPSALAGPFTYGVCQTGCNVVAVACYAAAGFTFGTVAAAGAPAVIVACNSALGTCSAGCAALLVTPTP**
>XP_038910438.1 uncharacterized protein EI90DRAFT_3079715 [Cantharellus anzutake]
MRFSIASTFAFVAMALNVTHVQAGPVAMGLCYSACNAGYVTCCTAAGVTAGTFTLGLGAPVALIACSLVQGACMSACTPLLAAPTP
>XP_038921664.1 uncharacterized protein EI90DRAFT_2906664 [Cantharellus anzutake]
MKYFNPLILFSALTLAPSALAGPFTYGVCQTGCNVVTVACYAAAGFTFGTVAAAGAPAVIVACNSALGTCSAGCAALLVTPTP
>XP_038922423.1 uncharacterized protein EI90DRAFT_2989286 [Cantharellus anzutake]
MKYFNPLILFSTLALAPSALAGPFTYGVCQTGCNVVAVACYAAAGFTFGTVAAVGAPSAIVACNSALGTCSAGCAALLIAPAP
>KAG9075620.1 hypothetical protein FS749_012700 [Ceratobasidium sp. UAMH 11750]
MKSSLTQLSVIAFALATGRSVQAGPIAMGLCYSACNAGYVACCAGAGATAGTFTLGLGAPAALMACSVVQGTCMAACTPFLAAPSP
>QRV76181.1 transmembrane protein [Ceratobasidium sp. AG-Ba]
MKLSVTSLVAFAAVTMNIQQVQAGPVAMGLCYSACNTGYVTCCTAAGVTAGTFTLGLGVPAALLGCSAVQGACMAACTPLLAAPSP
>KAF8604944.1 hypothetical protein BDV93DRAFT_521837 [Ceratobasidium sp. AG-I]
MKLSIRSLVAAVVVSLSAPQPALAGPIAMGLCYSACNAGYTTCCTVAGVTAGTFTFGLGVPAAIAACSVIQGTCMATCTPLLTAPSP
>KAF8604943.1 hypothetical protein BDV93DRAFT_491014 [Ceratobasidium sp. AG-I]
MKFSFTSIVAVVAVALNASQPVQAGPIAMGLCYSACNAGYVACCASAGTTAGTFTLGLGVPAAVAGCSGAQGACMAACTTLIVTPTP
>KAG9082513.1 hypothetical protein FS749_006798 [Ceratobasidium sp. UAMH 11750]
MKFSFTSLVAVVAIALSAERVQAGPVAMGLCYSACNAGYVTCCAAAGAVAGTFTLGLGVPAALFVCSAVQGTCMAACTPLLAAPTP
>QRV90993.1 transmembrane protein [Ceratobasidium sp. AG-Ba]
MKFSVTSLVALIAVTMNVQQAQAGPVAMGLCYSACNAGYVTCCTTAGAVAGTFTLGLGVPAALFACSAVQGACMAACTPLLAAPTP
>KAG9125829.1 hypothetical protein FRC07_006057 [Ceratobasidium sp. 392]
MKLSLTSIVAIMTIALSAERAQAGPVAMGLCYSACNAGYVTCCATAGAIAGTFTLGLGVPAALFTCSVVQGTCMAACTPLLAAPTP
>KAG8697657.1 hypothetical protein FRC08_006389 [Ceratobasidium sp. 394]
MKLSIAFFSVITAVALHTGNVRAGPMALALCTATCQAGYTTCCAAAGTAIGIFTFGLGTPVAVAGCSLARGACVAACAPLLAAQGP
>KAI0693946.1 hypothetical protein C8T65DRAFT_744563 [Cerioporus squamosus]
MRFAILAALAAIVAVPTAEAGPLAYAICQTGCNSLVVACYANAGAVFGTVTAGVGVAPAILACNAALGTCFTACATAALCAPTP
>KAG9312233.1 hypothetical protein JVU11DRAFT_7532 [Chiua virens]
MNFKSLTAITLAAAAVPLASAGPLAYAACQTGCNGLAVACYTGAGFVFGVTVVGGPPAILACNAALGTCMATCATVALFAPTP

>KAG9311090.1 hypothetical protein JVU11DRAFT_8998 [Chiua virens]
MNPKSLAALTLAAAAVPLVSAGPIAYAICQTGCNSLAVVCYSAAGFTFGTVAAPAAPAAILACNAGLGLCMTACAATALIAPIP
>KAF5390836.1 hypothetical protein D9757_004472 [Collybiopsis confluens]
MRFTKASISVLAVFTGLQTAQAGPIAYGICQTGCNTVTVACYAAAGFTFGTVVAAAAAPPMILACNAALGTCSAACATVALFAPTP
>XP_007768952.1 hypothetical protein CONPUDRAFT_20752, [Coniophora puteana RWD-64-598 SS2]
VAASTAPAAFGGPLAYAACQTGCNTLAVACYAAAGFTFGTTIVAGPPAIVACNAALGTCMTACATVALFAPTP
>XP_007769248.1 hypothetical protein CONPUDRAFT_105217 [Coniophora puteana RWD-64-598 SS2]
MNLKLAGALLVAASAAPAAVGGPIAYGICQTGCNGLAVACYAGAGFTFGVTIIGGPPAVIACNVALGTCMAGCATVALFAPTP
**>TFK18913.1 hypothetical protein FA15DRAFT_602445 [Coprinopsis marcescibilis]**
**MRLSASLAPIFAFVTLVHAGPIAYGICQTGCNAVAVACYAAAGCTFGTVAAPAAPLAIIGCNSALGTCSTACATVALFAPTP**
>KAH6901375.1 hypothetical protein BKA70DRAFT_1310477 [Coprinopsis sp. MPI-PUGE-AT-0042]
MRLTLAAASLIAFVGQVNAGLIMYGICQTGCNTVAVACYAAAGFTFGTIAAPVAPAAIVACNGALGTCSAACATVGLFAPTP
>KAH6902114.1 hypothetical protein BKA70DRAFT_1307299 [Coprinopsis sp. MPI-PUGE-AT-0042]
MRLTFTLASLALISQVNAGLIAYGICQTGCNALAVACYAAAGFTFGTVAAPAAPAAIVACNSALGTCSAACAAVALCAPTP
>KAH6901377.1 hypothetical protein BKA70DRAFT_1310482, partial [Coprinopsis sp. MPI-PUGE-AT-0042]
MRLALAVASLIVFIGQVNAGPIMYGICLAGCNATAATCYAAAGTTFATIAALLAPATIFVCNSALATCSASCTAAFFYPWSM
>KAH6901376.1 hypothetical protein BKA70DRAFT_1310479 [Coprinopsis sp. MPI-PUGE-AT-0042]
MRLTLAVASLVAFVAQVHAGPIMYGICQTGCNAVAVACYAAAGATFGTIAAPAAPAAIVACNGALGTCSAACATVGLFAPTP
>KAH6902112.1 hypothetical protein BKA70DRAFT_1157425 [Coprinopsis sp. MPI-PUGE-AT-0042]
MRFTTSIAVASLVAFAGQVSAGPIMYGICQTGCNVVAVACYAAAGATFGTIAAPAAPAAILGCNSALGSCSAACATVALFAPTP
>KAH6902116.1 hypothetical protein BKA70DRAFT_1307301 [Coprinopsis sp. MPI-PUGE-AT-0042]
MMRFTIALASLIAFVGQVQAGPIAYGICQTGCNAVVVACYAAAGATFGTVAAPAAPAAIVACNTALGTCSAACATVALLAPTP
>KAH6902106.1 hypothetical protein BKA70DRAFT_674323 [Coprinopsis sp. MPI-PUGE-AT-0042]
MRLTLGAAASLVAFIGQVNAGPTSEPYGDCQTGCNAIAAKCYTAAGFTFGTVAAADAPPAILACNSALASCSAVCADNHPSGNGKARMG
>KAF8153443.1 hypothetical protein B0H34DRAFT_800644 [Crassisporium funariophilum]
MSTAVLAGPIAYGICQTGCNALVVSCYAAAGFTFGTVVAAPAVPAVILGCNAGLGTCSAACATVALFAPTP
>KAF9528339.1 hypothetical protein CPB83DRAFT_894379 [Crepidotus variabilis]
MRFNTAATLAILAATTSSVMGGPLSYGLCQTGCNTVAVACYAAAGCVFGTVVAAAAAPAAVLGCNAALGTCSATCATLVLFAPIP
>KAH8107856.1 hypothetical protein BXZ70DRAFT_1003281 [Cristinia sonorae]
MKLSILTPLAVLAAAPTALGGPIAYGICQTGCNTVAVACYAAAGFTFGTVVAAAAAPAVIIGCNSALGTCSAACATVALFAPTP
>TFK33514.1 cysteine-rich protein [Crucibulum laeve]
MRLSTLTATLAMALVYVPTAEAGIISYGICQTGCNVLAVACYAAAGFTFGTVVAAAAAPPAILACNAGLGTCSSACAVVALTPTP
>KAH9894858.1 cysteine-rich protein [Cubamyces lactineus]
MKLSTLFIPVALTLGALPSADAGLLGYGVCQTGCNALAVACYAAAGYTFGTVTAGLGTPAVIVGCNAALGKCSAACAIVALAPTP
>KAI0326571.1 hypothetical protein GY45DRAFT_1328722 [Cubamyces sp. BRFM 1775]
MKLSTFFIPVALTLGALPSANAGLLGYGICQTGCNAVAVACYAAAGFTFGTVTAGLGTPAVILGCNAALGKCSAACAIVALTPTP
>KAI0656638.1 cysteine-rich protein [Cubamyces menziesii]
MKLSAFFIPIALGLGALPSANAGIIGYGICQTGCNVVAVACYAAAGYTFGTVTAGLGTPAVILGCNAALGKCSAACAIVALTPTP
>KAI0656637.1 hypothetical protein C8Q70DRAFT_1056539 [Cubamyces menziesii]
MNFISFAVLLTLVACAATADAGPIAYGLCQTGCNAVVVACYAAAGATFGTVTAGVGTPAAIIACNLALGQCSAACATIALLAPTP
>KAH9894857.1 hypothetical protein C8Q73DRAFT_790064 [Cubamyces lactineus]
MNFKSFPALLLTFIACAATVDAGPIAYGLCQTGCNAVVVACYAAAGATFGTITAGVGTPAAIIACNVALGQCSAACATIALFAPTP
>KAF9014159.1 hypothetical protein BDQ17DRAFT_1270363 [Cyathus striatus]
MRFTPILASLLIAPVVLSGPISYGICQSGCNAVAVACYAAAGFTFGTVAAAAAPAALVGCNSALGTCSAACASTTLIAPIP
>KAF8980957.1 hypothetical protein BDQ17DRAFT_1263431 [Cyathus striatus]
MRLSAVVFPLAFAPLVLGGPIAYGICQAGCTAAAATCYSAAGFIFGIYVPPLAPAAITACNTALATCSAACYMSWFAPTP
>KAF8980955.1 hypothetical protein BDQ17DRAFT_1438695 [Cyathus striatus]
MRLSAVVSPLAFAPLVLGGPIAYGICQTGCNTLAVACYAAAGFTFGTIAAPLAPVAIIGCNTALGTCSAACAVTALIAPIP
>KAF8977750.1 hypothetical protein BDQ17DRAFT_1293201 [Cyathus striatus]
MRLSAIVAPLAFAPLVLGGPIAYGICQTGCNTVAVACYAAAGFTFGTIAAPAAPAAIVGCNTALGSCSAMCAGIALLAPTP
>KAI0705164.1 hypothetical protein BC835DRAFT_1230698, partial [Cytidiella melzeri]
LFTTLAAAATVNGGPIAYGICQTGCNTVAVACYAGAGFVFGVALPAAPPAIMACNAALGTCSAACATVALFAPTP
>KZT72154.1 hypothetical protein DAEQUDRAFT_723321 [Daedalea quercina L-15889]
MKTPFALIGAALAMAAPAFAGPIAYGICQTGCNTVAVACYAAAGFTFGTVIAAPAAPAAIIACNAALGTCSAVCASVALFAPTP
>KZT72155.1 hypothetical protein DAEQUDRAFT_723324 [Daedalea quercina L-15889]
MKIPFALTGAALAMAAPVSADPIALVICLIGCNTVAVACYAAAGFTFGTVIAAPATPVAIIACNAALRTCSAACAGT
>KAI0737527.1 hypothetical protein C8Q80DRAFT_1348242 [ç nitida]
MSLFRRAIVVAAVTLVALPSTEAGLIAYGICQTGCNALAVACYAGAGAVFGTVTAGVGTPAAILACNAALGQCSAACAVVALTPTP
>KAI0737528.1 hypothetical protein C8Q80DRAFT_1114935 [Daedaleopsis nitida]
MNFARLSLLSAAALYMTVPAVQAGPIAYGICQTGCNSVAVACYAAAGVVFGTVTAGVGVPPAILACNMALGVCSAACATVGLFAPTP
>KAA1466913.1 hypothetical protein DENSPDRAFT_876911 [Dentipellis sp. KUC8613]
MRLSLLPLAAAAALVPSVLGGPISYAICQTGCNTVAVACYAAAGFQFGTVLAVAAPATILACNSALGTCSATCAGITLLAPIP
>TFY71517.1 hypothetical protein EVG20_g1493 [Dentipellis fragilis]
MRFSYLTIVATMALLPTAMGGPISYAICQTGCNTVAVACYAAAGFQFGTILAVAAPATIIVCNSALGTCSAACAGITLLAPIP
>KAA1466928.1 hypothetical protein DENSPDRAFT_926240 [Dentipellis sp. KUC8613]
MRFSYLAGAVVFALSPAVMGGPISYAICQTGCNAVVVACYAAAGFQFGTVLAAAAPASIVACNSALGTCSAACAGITLLAPIP
>THV03464.1 hypothetical protein K435DRAFT_651135 [Dendrothele bispora CBS 962.96]
MLLLTPTSVVLLIGLAILQSTQADLIAYGICQTTCNSAAAACYAAAGFMFGTMIAVHNVPPVVLACNTGLGTCSATCVVALLTPD
>THU93403.1 hypothetical protein K435DRAFT_670303 [Dendrothele bispora CBS 962.96]
MRLSTVFAPVLVGLGALQSVQAGPIAYGICQTGCNAVAVACYAGAGFTFGTVVAAAAAPPVILACNAALGTCSTACATVALFAPTP
>KAK0204073.1 hypothetical protein DFS33DRAFT_1384333 [Desarmillaria ectypa]
MRLSRAFAFLATSLALAPQVHAGPIAYGICQTGCNTVVVACYAAAGFTFGTVIAAPAAPAVVIACNAALGTCSAACATVALFAPTP
>KAK0204044.1 hypothetical protein DFS33DRAFT_1336384 [Desarmillaria ectypa]
MRLSPIFAFLVTSLALAPQAYAGPIAYGICQTGCNVVAVACYAAAGFTFGTVAAPAAPPAIVGCNTALGTCSAACASVVLLAPIP
>XP_060325687.1 uncharacterized protein EV420DRAFT_914577 [Desarmillaria tabescens]
MRLSPVLAFLVTSLALAPQAHAGPIAYGICQTGCNVLAVACYAAAGFTFGTVAAPAAPAAIVACNSGLGTCSAACASVALLAPTP
>XP_060325653.1 uncharacterized protein EV420DRAFT_911665 [Desarmillaria tabescens]

MRLSRAFAFLATSLALVPQAHAGPIAYGICQTGCNTVVVACYAAAGFTFGTVIAAPAAPAAVLACNAALGTCSAACATVALLAPTP
>XP_007366826.1 uncharacterized protein DICSQDRAFT_107485 [Dichomitus squalens LYAD-421 SS1]
MNLRLSTLVIVATGLLAASPIVNAGPVAYGICQTGCNAVVVACYAGAGFTFGTVTAGLGVPAAIVACNAALGTCSAACATVALFAPTP
>XP_007370845.1 uncharacterized protein DICSQDRAFT_174921 [Dichomitus squalens LYAD-421 SS1]
MRFHLSTLAIAATSLLAVFPTVTAGPIAYGLCQTGCNTVTVACYAGAGFTFGTVTAGAGVPAAILACNAALGVCSSTCATVALFAPTP
>KAI0744947.1 hypothetical protein C8Q76DRAFT_789409 [Earliella scabrosa]
MKLTLPVISTLAISLSAFPSVHAGLIAYGICQTGCNTVAVACYAAAGAVFGTVTAGVGTPAAILGCNAALGQCSAACAVVALTPTP
>KAI0744948.1 hypothetical protein C8Q76DRAFT_789410 [Earliella scabrosa]
MNVKLLSIAVVLSTLPALPVYAGPLAYALCQTGCNAVVVACYGAAGAVFGTVTAGVAVAPAILACNAALGTCSAACAATALIAPTP
>XP_047873742.1 uncharacterized protein BXZ73DRAFT_105425 [Epithele typhae]
MHFTPSSLLAAAVLLATGAHAGPVAPYGVCQTGCVLVAVACYSAAGFTFGTVKADDPHVPAAVLNCNAALGTCQAACAKVTLPAATPH
>XP_047873745.1 uncharacterized protein BXZ73DRAFT_105428 [Epithele typhae]
MQLKPSLLAAAALLATGARASSVAYDVCQTAGCNTVAVACYAGASFAFGTFTAGLGVPPALVACQTTLEKCSSACASLFPSTP
**>XP_047870886.1 uncharacterized protein BXZ73DRAFT_93778 [Epithele typhae]**
**MLFKLSSLVAAAAILATGAHAGPIAYGICQTGCNAVVVACYAGAGFTFGTVTAGLGVPAAIVACNAALGTCSTACATIGLFAPTP**
>XP_047873747.1 uncharacterized protein BXZ73DRAFT_53238 [Epithele typhae]
MLFKLSSLVAAAAILATGARAGPIAYGICQTGCNTVAVACYAGAGFTFGTVTAGLGVPAAIVACNAALGTCSAACATIGLFAPTP
>XP_047873743.1 uncharacterized protein BXZ73DRAFT_105426 [Epithele typhae]
MQFKLSSLLAAALLATGAQAGPALYGICQTGCNTLAFACYAGAGFTFGTVTAGLGIPAVIVGCNTALGTCSAACAAVTLLAPTP
>XP_047873744.1 uncharacterized protein BXZ73DRAFT_105427 [Epithele typhae]
MQFKLSSLLAAALLATGASAGPAFYGICQTGCNTLAVACYAGAGFTFGTVTAGVGIPAAIAACNSALGTCSAACAAVTLLAPTP
>XP_047873746.1 glutathione S-transferase [Epithele typhae]
GELKLSSLLAATALITTGVHAGPALYGVCQTGCNAVTVACYAGAGFTFGTVVGAGAPQAVAACSAAQGKCSSACAAVTLFAPTP
>XP_047873750.1 uncharacterized protein BXZ73DRAFT_105434 [Epithele typhae]
MHFKLSSFLAAAALRATCGSQAGPMAYGICQTGCNKGVVACYAGAGFTFGSISTAGVDVPAAITTCNATLGVCLAACAAVTFFPST
>KZV97636.1 hypothetical protein EXIGLDRAFT_730290 [Exidia glandulosa HHB12029]
MKPSRIVLPLTLVLSANAGLIAYGICQTGCNMGAVACYAVAGAVFGTVAAPTAPAAILACNAAQGMCMATLCAPLLLIPFP
>KAI0779875.1 hypothetical protein C8Q74DRAFT_1367694 [Fomes fomentarius]
MNFKLSALSALAVLYVVPTVEAGPLAYGLCQTGCNAVVVACYGAAGAVFGTVTAGVAVAPAIVACNAALGVCSAACAATALIAPTP
>KZP29597.1 hypothetical protein FIBSPDRAFT_851544 [Fibularhizoctonia sp. CBS 109695]
MRFTPVALLAIVAATPVLGGPIAYALCQTGCNGLAVACYAGAGFTMGVAIVAAPPAIMACNAGLGGCMAICATVGLFAPTP
>KZP33058.1 hypothetical protein FIBSPDRAFT_943490 [Fibularhizoctonia sp. CBS 109695]
MRFTLIALLAVIAAATPALGGPLAALACQTGCISLTATCYAAAGFIFAPTVIGVPPAIITCNVALGTCTAGATVVLFAPTP
>KZP06955.1 hypothetical protein FIBSPDRAFT_1053254 [Fibularhizoctonia sp. CBS 109695]
MRFTPVALLAIVAVATPALGGPLAYAACQTGCNGLAVACYAGAGFVMGVTIVGAPPAVMACNAGLGGCMAICATVGLFAPTP
>KZP33060.1 hypothetical protein FIBSPDRAFT_847674 [Fibularhizoctonia sp. CBS 109695]
MRITPVTILAVAAVATPALGGPLAYALCQTGCNGLAVACYAGAGFTMGVAIVAAPPAIIACNLALGTCMATCATVALLAPTP
>KZP05526.1 hypothetical protein FIBSPDRAFT_765712 [Fibularhizoctonia sp. CBS 109695]
MRLTHVTLLAIAGAATPAMGGPLAYAACQTGCNGLAVACWAAAGFTFGVTIVLVPPAILACNVGLGTCMATCATVALFAPTL
>KZP03955.1 hypothetical protein FIBSPDRAFT_878987 [Fibularhizoctonia sp. CBS 109695]
MRFTPVALLAIAAAATPVLGGPLAYAMCQTGCNGLAVACYSGAGFIMGTTIVGGPPAIIACNLGLGTCMATCATVALFAPTP
>KIY47514.1 hypothetical protein FISHEDRAFT_45305 [Fistulina hepatica ATCC 64428]
MQITKPCLALLAACGLAQAGPIAYGICQTGCNVVAVACYAATGFTFGTVVASAATPAVILGCNSALGTCSAACASVALLAPTP
>KAH8831031.1 hypothetical protein DL96DRAFT_1586273 [Flagelloscypha sp. PMI_526]
MRLSHLFMAFAGMALAPTGAAGPLAYAVCQTGCNTIAVACYAAAGVQFGTVVAAAGAPATVIGCNVALGTCMTACAGTALIAPIP
>KAF8957624.1 hypothetical protein BDZ97DRAFT_1669783 [Flammula alnicola]
MRFSLIAAPILYVLASTSIAQAGPIAYGLCQTGCNVMAVACYAGAGFTFGTVIAAPAAPAAVLACNAALGTCSATCATVALLAPTP
>KDR81157.1 hypothetical protein GALMADRAFT_136196 [Galerina marginata CBS 339.88]
MRFSIAVAPFLIALCTTTSFVSAGPIFYGICQTGCNAVAVACYAGAGATFGTVVAAAAAPAAILACNSALGTCSAACAAVTLLAPTP
>KDR81156.1 hypothetical protein GALMADRAFT_241716 [Galerina marginata CBS 339.88]
MRFPTVIAPFLIALSTTVFLVSAGPIEYGICQTACNDGAVACYRGAGATFGTVVTDADTPAAILACNAGLGACSAACPAVALPGPTS
**>KAI1791328.1 hypothetical protein LXA43DRAFT_1094705 [Ganoderma leucocontextum]**
**MQLKLSALALALAGLAASPVVNAGPIAYGICQTGCNAVVVACYAGAGFTFGTVTAGLGVPAAIIACNAALGTCSAACATVALFAPTP**
>PIL23384.1 hypothetical protein GSI_14695 [Ganoderma sinense ZZ0214-1]
MHLKLSALTLALVGLAASPVANAGPIAYGICQTGCNTVAVACYAGAGFTFGTVTAGIGVPAAILACNAALGTCSSACATVALFAPTP
>KAI1785556.1 hypothetical protein LXA43DRAFT_123492 [Ganoderma leucocontextum]
MDFRFKAISLLVGVTATIGIVLAVVNGNPSAHEVCQTGCNAVLVACYGTGGFSLGRAQGVSLGPPPSMLGAICNTAIGSCLAACTAAICQ
>KAI1794409.1 hypothetical protein LXA43DRAFT_138977 [Ganoderma leucocontextum]
MHFKLSALALALVGLAASPVANAGPIAYGICQTGCNTVTVACYAAAGFTFGTVTAGVGVPAVILGCNTALGICSSACATVALFAPTP
>PIL33906.1 hypothetical protein GSI_03612 [Ganoderma sinense ZZ0214-1]
MQLKLSFKLSAALAVTLSLPQVANAGPIAYGICQTGCNVVAVACYAGAGFTFGTVTAGLGVPAAVLACNAALGTCSSACATVALFAPTP
>KAF8500117.1 hypothetical protein JB92DRAFT_2979463 [Gautieria morchelliformis]
MNLKSHALVLLAALIPAVNGGPVAYALCQTGCNAVAVACYGAAGFTFGVTIVGAPAAILGCNAALGTCMATCATVALLAPTP
>XP_007868697.1 hypothetical protein GLOTRDRAFT_46665 [Gloeophyllum trabeum ATCC 11539]
MRFYTIALPLLAAMASIPSTIAGPIAYGICQTGCNALAVACYAGAGFTFGTVVAAPAGPAAVLACNAALGTCSAACATTALIAPIP
>XP_043040323.1 uncharacterized protein BT62DRAFT_931391 [Guyanagaster necrorhizus MCA 3950]
MRLSRVFAFLATSLALAPQAHAGPIAYGICQTGCNGMAVACYAAAGFTFGTVIAAPAAPAAILACNAALGTCSTACATVALLAPTP
>XP_043040322.1 uncharacterized protein BT62DRAFT_931390 [Guyanagaster necrorhizus MCA 3950]
MRLSPIFVFFVTSLALVHQTHAGPISYGICQTGCNALAVACYAAAGFTFGTIAAPAAPAAIVSCNSALGACSASCASVALFSPTP
>PPR07183.1 hypothetical protein CVT26_012613 [Gymnopilus dilepis]
MRFNALVSALAVVIPMASAGPIAYGICQTGCNTVAVACYAAAGFTFGTIAAPVAPAAVVACNAALGTCSAACATVALLAPTP
>KAF8898031.1 hypothetical protein CPB84DRAFT_1781195 [Gymnopilus junonius]
MRSALLIALPFISMAAAGPIAYGICQTGCNTVAVACYAAAGLTFGTIAAPAAPAAILGCNAALGTCSATCATVALLAPIP
>KAF9222454.1 hypothetical protein BS17DRAFT_783729 [Gyrodon lividus]
MNLKSLAALTLAVSATPAVMAGPFAYGLCQTGCNVLVGACYAGAGFTFGVTIVAAPPAIIACNTGLGACMAACAATALIAPTP
>KAF9222481.1 hypothetical protein BS17DRAFT_783760 [Gyrodon lividus]
MKLKFVTALAVAASIPPLTIAGPIAYAICQTGCNTLAVACYAGAGFTFGVTVVGAPAAIVACNFALGKCMTACALTALPAPIP

>KIM37591.1 hypothetical protein M413DRAFT_448389 [Hebeloma cylindrosporum h7]
MHFSKLFAPVAIAIASASVVQGGPIAYGICQTGCNAVVVACYSAAGATFGTVVAAIAAPPALLACNAALGTCSAACATVALLAPTP
>THG95442.1 hypothetical protein EW026_g6218 [Hermanssonia centrifuga]
MNFKVLAAAVLAAVAPVANAGPIAYGICQTGCNSLAVACYAGAGLTFGTIVAAPLAPAAALACNVALGTCSAACATVALFAPTP
>XP_009549554.1 cep2 cellulose medium expressed protein 2 [Heterobasidion irregulare TC 32-1]
MVRITPLAAVSLLSAIPLVAGGPISYGLCQTGCNTVAVACYAAAGFQFGTVVAAAATPATILACNAALGTCSATCATLVLFAPIP
>KIJ66076.1 hypothetical protein HYDPIDRAFT_87212 [Hydnomerulius pinastri MD-312]
MNFKALAALTLAASAAPLTMAGPLAYAACQTGCNVLAVACYAGAGFTFGVTIVAVPPAIMACNAGLGTCMAACATVALFAPTP
>KAH7911314.1 hypothetical protein BJ138DRAFT_1150936 [Hygrophoropsis aurantiaca]
MNLKSTAALILVAASAPAVLGGPLAYAACQTGCNGLAVACYAAAGFTFGVTIVGAPPAIMACNLGLGTCMATCATIGLFAPTP
>KAF9018149.1 hypothetical protein BDZ89DRAFT_357746 [Hymenopellis radicata]
MVRFQRLALLALPFIPAINAGPIAYGICQTGCNAVAVACYAGAGFTFGTITAGVGIPAAIVACNAALGTCSAACASVTLLAPTP
>KJA16367.1 hypothetical protein HYPSUDRAFT_147795 [Hypholoma sublateritium FD-334 SS-4]
MRFSTLAIALASAASVSAGPIAYGLCQTGCNVVAVACYGAAGATFGTVVAAAAAPAAILACNSALGTCSAMCASVALLAPTP
>KJA27646.1 hypothetical protein HYPSUDRAFT_130814 [Hypholoma sublateritium FD-334 SS-4]
MRLAVLTTLAVGAATATAGPIAYGVCQTECNTVAEACYTAAGFTFGTVVAGPETPAVVLRCNAALGTCAHACATSALRAPTP
>KJA16363.1 hypothetical protein HYPSUDRAFT_115057, [Hypholoma sublateritium FD-334 SS-4]
AVALASIGSANAGLITYGICQTGCNTVAVACYAAAGFTFGTVIAAPATPAVILACNAALGTCSTMCATVALLAPTP
>KJA16366.1 hypothetical protein HYPSUDRAFT_47387 [Hypholoma sublateritium FD-334 SS-4]
MRLSILAPLAVALASIGSANAGLITYGICQTGCNTVAVACYAAAGFTFGTVIAAPATPAVILACNAALGTCSTMCATVALLAPTP
>RDB15947.1 hypothetical protein Hypma_003602 [Hypsizygus marmoreus]
MRLSILAPLIFALSATQGVKGGPIAYGICQTGCNALVVSCYAAAGFTFGTVVAAPATPLVILGCNTGLGTCSAACATVALFAPTP
>KAF8554845.1 hypothetical protein OG21DRAFT_1508491 [Imleria badia]
MNLKSLAALTLAASAAPLAAAGPLAYALCQTGCNGLAVACYTAAGFTFGVTIVAAPPAIIGCNVGLGACMAACAGTALIAPTP
>KAF8869089.1 hypothetical protein BD779DRAFT_1682623 [Infundibulicybe gibba]
MRFSSALLVAASMAPVVLGGPISYGICQSGCYAAAVICYAAAGFTFSVIVATPAIPPALLVCNAGLAACSAVCNTTFFASTP
>KAF8872941.1 hypothetical protein BD779DRAFT_1613733 [Infundibulicybe gibba]
MRFSTAFLVTLGMAPVALGGPIAYALCQTGCNSLAVACYAAAGFTFGTVVATVATPAVIVGCNAGLGTCSAACATVALFAPTP
>KAF8869829.1 hypothetical protein BD779DRAFT_1730859 [Infundibulicybe gibba]
MRFSTALLIAASMAPVALGGPISYGICQSGCNAVVVACYAAAGFTFGTVVAAPATPAVLLACNAGLGTCSAACATVALFAPTP
>KAF8872939.1 hypothetical protein BD779DRAFT_1452367 [Infundibulicybe gibba]
MRFSTTFLVALGMAPVALGGPIAYGICQTGCNSLAVACYAAAGFTFGTVVAAAATPAVIVACNAGLGTCSAACATVALFAPTP
>KAI0089603.1 hypothetical protein BDY19DRAFT_872480, partial [Irpex rosettiformis]
LLALVAVAGTANAGPIAYGICQTGCNAVAVACYAAAGFTFGTVVAAPATPAVIIACNAALGTCSAACATVALFAPTP
>KDQ57059.1 hypothetical protein JAAARDRAFT_207405 [Jaapia argillacea MUCL 33604]
MRPYTLFLPILAAVATVLPTASAGPIAYGLCQTGCNALAVACYAGAGFTFGTVVAAAAAPAAVLACNAALGTCSAACAATALLAPIP
>KIK03653.1 hypothetical protein K443DRAFT_94727 [Laccaria amethystina LaAM-08-1]
MRISTTLLLSPLLMVANVNAGPLAYGLCQTGCNTVAVACYAAAGFTFGTVIAAAATPAAILGCNAALGTCSATCATLVLFAPTP
>KIJ99165.1 hypothetical protein K443DRAFT_102635 [Laccaria amethystina LaAM-08-1]
MRISSAILLSPFLMVANAGPIAYGLCQTGCNTVVVACYAAAGFTFGTVIAAPAAPAAILGCNAALGTCSATCATLVLLAPTP
>XP_040768338.1 uncharacterized protein LAESUDRAFT_644095 [Laetiporus sulphureus 93-53]
MKFTTTLTTLALALATPAAAGPIAYGLCQTGCNTVAVACYAAAGFQFGTVVASPLVPATILACNAALGTCSATCATVVLLAPIP
>XP_040768339.1 uncharacterized protein LAESUDRAFT_644175 [Laetiporus sulphureus 93-53]
MKFTTTLTTIALALATPAAAGPIAYGLCQTGCNAVAVACYAAAGFQFGTVVATPLAPATVLACNAALGTCSATCATVVLFAPIP
>KAJ4480927.1 hypothetical protein J3R30DRAFT_2380869 [Lentinula aciculospora]
MRLTNTLLPFLSVLAGLQSAQAGIIAYGICQTGCNVAAGACYTAAGFTFGTVVAAPATPAVILGCNAALGTCSAACATVALLAPTP
>KAJ3998398.1 hypothetical protein F5050DRAFT_1805941 [Lentinula boryana]
MRLTNILLPILPVLAGMQSAQAGPIAYGLCQTGCNVVAVACYAAAGFTFGTVVAAPATPAVILGCNAALGTCSATCAAVALLAPIP
**>KAJ3742886.1 hypothetical protein DFH05DRAFT_1262705 [Lentinula detonsa]**
**MRLTNILLPILPVLAGMQSAQAGPIAYGLCQTGCNVVAAACYAAAGFTFGTVIAAPTTPAVILGCNAALGTCSATCAAVALLAPIP**
>KAJ3793355.1 hypothetical protein GGU11DRAFT_800070 [Lentinula aff. detonsa]
MRLTNILLPILPVLAGMQSAQAGPIAYGLCQTGCNTVVVACYAAAGFTFGTVVAAPATPAVILGCNAALATCSATCATVALLAPTP
>KAJ3870732.1 hypothetical protein F5051DRAFT_423777 [Lentinula edodes]
MRLTNVLVPILSVLAGLQGAQAGPIAYGLCQTGCNIVAGACYAAAGFTFGTVVAAPATPAVILGCNAALGTCSAMCATVALLAPTP
>KAJ3874754.1 hypothetical protein F5051DRAFT_416033 [Lentinula edodes]
MRLTNVLVPVLSVLAGLQGAKAGLIAYGLCQTGCNTVAVACYAAAGFTFGTVIAAPATPAVILGCNAALGSCSAMCATVALLAPTP
>KAJ3729090.1 hypothetical protein DFJ43DRAFT_1225335 [Lentinula guzmanii]
MRLTNILLPILPVLAGMQSAQAGPIAYGLCQTGCNVVAAACYAAAGFTFGTVTAAPTTPVVLLGCNAALGTCSATCATVALLAPTP
>KAJ3868585.1 hypothetical protein EV359DRAFT_31918 [Lentinula novae-zelandiae]
MRLTNVLVPVLSVLAGLQGAKAGLIAYGLCQTGCNSVAVACYAAAGFTFGTVIAAPATPAVILGCNAALGTCSAMCATVALLAPTP
>KAJ3727548.1 hypothetical protein C8R42DRAFT_573285 [Lentinula raphanica]
MRFTNILLPVLSVLAGMHNVQAGPIAYGLCQTGCNVVAVACYAAAGFTFGTVVAAPATPAVILGCNAALGTCSATCATVALLAPTP
>RPD73715.1 hypothetical protein L226DRAFT_571991 [Lentinus tigrinus ALCF2SS1-7]
MRFVLATVLVALVAVPTAVDAGPLAYGICQTGCNAVVVACYGAAGAVFGTVTAGVGVPPAILACNAALGICSAACAAAVLAPTP
>KAH9851373.1 hypothetical protein C2E23DRAFT_886554 [Lenzites betulinus]
MFFKPSPTVFLTFLAALSAAPAAHAGPLAYGICQTGCNALVVACYAGAGAVFGTVTAGVGTPAAIVACNVALGQCSAACALIVLAPTP
>KAH9851372.1 hypothetical protein C2E23DRAFT_733103 [Lenzites betulinus]
MKFSTIVSVALGLAAVPSAKAGLLAYGICQTGCNTMAVACYAAAGATFGTVTAGAATPAIILGCNAALGKCSASCALVTLLPTP
>KAF9461003.1 hypothetical protein BDZ94DRAFT_1168475 [Lepista nuda]
MRFSKILLSAALALPTVQAGLISYGLCQTGCNTVAVACYAAAGFTFGTVIASAATPAVIVGCNAALGTCSATCASLVLLAPIP
>KAF9461010.1 hypothetical protein BDZ94DRAFT_1168462, partial [Lepista nuda]
ALICAFLAIPSAQAGPLLYGVCQTDCNALAVSCYAATGFTFGTVIATPAVPAVILACNASLGTCSAACVAVTLLAPTL
>KXN83432.1 hypothetical protein AN958_01446 [Leucoagaricus sp. SymC.cos]
MRPLKVFILVVASILSSSPQQAAAGPIAYGICQTGCNVVAVACYAAAGFTFGTIAAPVAPPAILACNAALGTCSAACAAVVLTPTL
>KXN81170.1 hypothetical protein AN958_05941 [Leucoagaricus sp. SymC.cos]
MRPFRTLTLVVAAILSSAPQQAMAGLIAYGICQTGCNTVAVACYGAAGVTFGTILAVAAPPAILACNAALGTCSAACAAVALTPTP
>KXN83427.1 hypothetical protein AN958_01441 [Leucoagaricus sp. SymC.cos]

MRPLKAFILVVASILSSAPQQEAAGPIAYGICQTGYNVVVVACYAAVGFTFGTIVAPVAPPAILAYNTALGTCSTARSAVASTSTP
>KXN91089.1 hypothetical protein AN958_02956 [Leucoagaricus sp. SymC.cos]
MRPFRTLILVVAAVLSSAPQQTMAGPIAYGICQTGCNVLAVACYAAAGFTFGTVAAAVAPPAILACNAGLGTCSAACAAVALTPTP
**>KXN83429.1 hypothetical protein AN958_01443 [Leucoagaricus sp. SymC.cos]**
**MRPLKTFILVVASILSSAPQQAAAGPIAYGICQTGCNIIAVACYAAAGFTFGTIAAPIAPPAILACNAALGTCSAACAAVVLTPTL**
>KAF5347592.1 hypothetical protein D9756_010704 [Leucoagaricus leucothites]
MRLQFKTCSTIAATILLIQPASAGLIAYGICQTGCNTLAVACYAAAGFTFGTIAAAAAPPAIVACNSALGTCSAACATIALTPTP
>KXN83428.1 hypothetical protein AN958_01442 [Leucoagaricus sp. SymC.cos]
MAGLIGYGICQTGCNAVAGACYAAAGFTFGTVLAVAAPPAILACNAALGTCSAACAVVALTPTP
>KXN93170.1 hypothetical protein AN958_00094 [Leucoagaricus sp. SymC.cos]
MRPLKAFILVVASILSSAPQQAAAGPIVYGMCQTGCNVVVAARYAAAGFTSGTIAAPVAVPAISTRNAAFGICSAACAAVASTPTP
>XP_040768337.1 uncharacterized protein LAESUDRAFT_636027, [Laetiporus sulphureus 93-53]
GPIAYALCQTGCNTVAAACYSAAGFQFGTVVASLLAPATILACNTALGTCSATCATVALFAPTP
>KAH7920598.1 hypothetical protein BV22DRAFT_1073654 [Leucogyrophana mollusca]
MNLRSTIALFLAAASAPVAIGGPLAYAACQTGCNGLAVACYTAAGFTFGVTIVAAPPAIMACNAGLGACMATCATIGLFAPTP
>KAK1228354.1 hypothetical protein PQX77_008607 [Marasmius sp. AFHP31]
MRFTTLTAFAVALFATLQGVNGGPIMYGVCQTGCNCVAVACYSAAGFTFGTVVAAAAAPAAILACNSALGTCSAACAATALIAPTP
>KAF9256214.1 hypothetical protein L218DRAFT_882284 [Marasmius fiardii PR-910]
MRFASLTTAVLLLVGLQEVKAGPIAYGICQTGCNIVAVACYAAAGATFGTVAAPAAPAAILGCNSALGSCSAACAATALIAPIP
>KAF9255919.1 hypothetical protein L218DRAFT_966834 [Marasmius fiardii PR-910]
MHFSFSTVAVLLSMVPWEVNAGLLAYGLCQTGCNCLAVACYSAAGATFGTVVASPAAPVAILACNAALGKCSAACATTALIAPTP
>XP_043004193.1 uncharacterized protein E1B28_013668 [Marasmius oreades]
MRSTSLATVLLLIGLQEVNAGPIAYGICQTGCNVVAVACYAAAGFTFATVAAPAAPAALIGCNSALGTCSAACAATALIAPIP
>KAJ8072958.1 hypothetical protein PM082_019821 [Marasmius tenuissimus]
MRFTALTAGTVALLAALQGVNGGPIAYGICQTGCNSVAVACYAAAGFTFGTVIAAPATPAVILACNAALGTCSAACATVGLFAPTP
>KAF9237865.1 hypothetical protein BU15DRAFT_75666 [Melanogaster broomeanus]
MNFKSLAALTLVASAAPLTMAGPLAYGLCQTGCNVLVGSCYAAAGFTFGVTIVAAPPAIMACNAGLGTCMATCATVALLAPTP
>ESK93305.1 proteophosphoglycan 5 [Moniliophthora roreri MCA 2997]
MRFTNILAPSALALLTGIQGVNGGLIAYGLCQTGCNTVAVACYAAAGFTFGTVVAAIAAPPVILACNAALGTCSAACAATALIAPIP
>KAJ6579297.1 hypothetical protein B0H10DRAFT_1835963 [Mycena sp. CBHHK59/15]
MRPTISTLAPLLLALAAVPAVQGGLISYGLCQTGCNTLAVACYAGAGLVFGTVAASPAAPAAALACNAALGQCSAICATVGLFAPTP
>KAJ6621514.1 hypothetical protein B0H10DRAFT_1789801, partial [Mycena sp. CBHHK59/15]
LLALATVPLVQAGPIAYGLCQTGCNTIAVVCYAGAGLVFGTVVAAPAAPAAALACNVALGTCSATCATVALLAPTP
>KAJ6621484.1 hypothetical protein B0H10DRAFT_1789821 [Mycena sp. CBHHK59/15]
MRAFKIIFAPLLSLATVPLVQAGPIAYGLCQTGCNTIAVVCYAGAGLVFGTVVAAPAAPAAALACNVALGTCSATCATVALFAPTP
**>KAJ7359841.1 hypothetical protein DFH08DRAFT_952947 [Mycena albidolilacea]**
**MRAFRAFFVPLLVLAGNTALVEAGPLAYGLCQTGCNSLAVACYAGAGLTFGTVVAAPAAPAAALACNAALGTCCATCATVALFAPTP**
>KAJ7330443.1 hypothetical protein DFH08DRAFT_751195 [Mycena albidolilacea]
MRAFKILIVPLLALSGITLVEAGPIAYALCQTGCNTVAVACYAGAGLVFGTVVAAPAAPAAAIACNVALGTCSATYATVALFAPTP
>KAJ7330535.1 hypothetical protein DFH08DRAFT_786255 [Mycena albidolilacea]
MRVFKTIAVPLLALSGITLVQAGLIAYAICQTAGCNTFAVACYAGAGLTFGMVVAVAEAPAAAIPCNSALGICSANCAAVALVAPTP
>KAJ7036623.1 hypothetical protein C8F04DRAFT_1210055 [Mycena alexandri]
MRTFNVLFAPLLALAAIPLVQGGPLAYALCQTGCNTVVVACYAGAGLVFGTVVAAPAAPAAALACNVALGTCSATCATVALLAPTP
>KAJ7036624.1 hypothetical protein C8F04DRAFT_1394200 [Mycena alexandri]
MRTFEILFAILLAVRLPLVQGDKVKDEVAYGLCQTGCNNITVACYSAAGLVFGTVIADADAPVAALACNKALSECSSNCT
>KAJ7080818.1 hypothetical protein B0H15DRAFT_786982 [Mycena belliae]
MRTANALVLPLAALSAGVSLVEAGPIAYGICQTGCNTVTVACYAGAGLVFGTVVAAPAAPAAAIACNVALGTCSAACATVALFAPTP
>KAJ6580769.1 hypothetical protein B0H19DRAFT_484617 [Mycena capillaripes]
MRASKTLFVPLLALAGIPFVQGGPLAYGLCQTGCNTLAVACYAGAGLVFGTVVAAPAAPAAALACNAALGTCSATCATVALLAPTP
>GAT42696.1 predicted protein [Mycena chlorophos]
MNPTKALTALVVAALASTVQAGPLAYGICQTGCNTVAVACYAGAGLTFGTVVAAAAAPAAALACNTALGTCSAA
>KAF7289020.1 hypothetical protein HMN09_01349900 [Mycena chlorophos]
MNPTKALTALVFAAVASTVQAGPLAYGICQTGCNTVAVACYAGAGLTFGTVVAAAAAPAAALACNAALGTCSAACATVALLAPTP
>KAJ7147695.1 hypothetical protein C8R43DRAFT_1129611 [Mycena crocata]
MRAFNALVLPLLALTGIPLVQGGPLAYGLCQTGCNTVAVACYAGAGLVFGTVVAAPAAPAAAVACNVALGTCSATCATVALFAPTP
>KAJ7114020.1 hypothetical protein C8R44DRAFT_710420 [Mycena epipterygia]
MRTFNTLFVPILALAGIPLVQGGPISYGLCQTGCNTLAVACYAGAGLVFGTVVAAPAAPAAALACNAGLGTCSATCATVALFAPTP
>KAJ7187815.1 hypothetical protein C8R46DRAFT_1052674 [Mycena filopes]
MHAFKVLFAPLLALAAVPLVQAGPLAYALCQTGCNTVAVACYAGAGLVFGTVIAAPAAPAAALACNVALGTCSATCATVALFAPTP
>KAJ7586376.1 hypothetical protein C8J56DRAFT_828258 [Mycena floridula]
MVGITKISALLVTSMAFLAIPVTAGPIAYGLCQTGCNTLAVACYAAAGFTFGTVVAAPAAPAVILACNAGLGTCSATCATVALFAPTP
>KAJ7498165.1 hypothetical protein B0H11DRAFT_833967 [Mycena galericulata]
MRAFNALHILALAAAPLVHGGVIPYVECQTGCNSVAVACYSAAGLVFGTVVATPDAPPAALACNNNLSTCATNCSTTALLAPTL
>KAJ7498176.1 hypothetical protein B0H11DRAFT_1998998 [Mycena galericulata]
MRAFNSAPLLALALAAVPLVQGGPIAYALCQTGCNTLAVACYAGAGLVFGTVIAAPAAPAAALACNAGLGTCSATCATVALLAPTP
>KAF8211017.1 hypothetical protein K438DRAFT_2011435 [Mycena galopus ATCC 62051]
MRAFSKLLLALSGVSLVHAGPIAYGLCQTGCNTVAVACYAGAGLVFGTVVAAPAAPAAALACNAALGTCSATCATVALLAPTP
>KAJ7208767.1 hypothetical protein B0H12DRAFT_1034116 [Mycena haematopus]
MRAFKILAVPFIALSGISLVQAGPLAYGLCQTGCNTVAVACYAGAGLIFGTIIAAPLAPPAAIACNVALGTCSATCATVALFAPTP
>XP_037222833.1 uncharacterized protein MIND_00309100 [Mycena indigotica]
MKLTKAISFILVALSPVLVEAGPIAYGICQTGCNTVAVACYAAAGFTFGTVVAAAAAPAAILACNSALGTCSTACATVALLAPTP
>KAJ7483446.1 hypothetical protein FB451DRAFT_1393550 [Mycena latifolia]
MRAFTAFVVPFVALAGIPLAHAGPIAYALCQTGCNTVAVACYAGAGLVFGTVVAAPAAPAAALACNVALGTCSATCATVALFAPTP
>KAJ7830972.1 hypothetical protein B0H13DRAFT_2371685 [Mycena leptocephala]
MRPTLVILAPLILALTAVPTVQGGLVSYGLCQTGCNTLAVACYASAGLVFGTVVASPAAPAAALVCNAALGKCSAICATVGLFAPTP
>KAJ7780067.1 hypothetical protein DFH07DRAFT_462579 [Mycena maculata]
MPTFKALLSVIIVAAILLVQGGPLAYGECQTGCNNLTVACYSRAGLVFGTVVAEPAAPAAALVCNKALSTCSSVCAKETLSAPTQ

>KAJ7780066.1 hypothetical protein DFH07DRAFT_1025822 [Mycena maculata]
MRANALSAPVLALALTALPLAHAGPVAYALCQTGCNTVAVACYAGAGLIFGTVVAAPAAPVAALACNAALGTCSATCATVALFAPTP

**>KAJ7712755.1 hypothetical protein B0H16DRAFT_1743832 [Mycena metata]**
**MRPLTLKALLAPLLAFAAIPLVEGGPIAYGLCQTGCNTLAVACYAGAGLVFGTVVAAPAAPAAALACNAGLGTCSATCATVALFAPTP**

>KAJ7724508.1 hypothetical protein B0H16DRAFT_1736697 [Mycena metata]
MRPLALKALLAPLLAFAAIPLVEGGPIAYGLCQTGCNTLAVACYAGAGLVFGTVIAAPAAPVAALACNAALGTCSATCATVALFAPTP

>KAJ7777091.1 hypothetical protein B0H16DRAFT_1504302 [Mycena metata]
MRAFNVLFAPLLALAAIPLVQGGPLAYALCQTGCNTVVVACYAGAGLVFGTVVAAPAAPAAALACNIALGTCSATCATVALLAPTP

>KAJ7743748.1 hypothetical protein B0H14DRAFT_3897621 [Mycena olivaceomarginata]
MRAFKILVVPLLALSGITLVEAGPIAYALCQTGCNTVAVACYAGAGLVFGTVVAAPAAPAAAIACNVALGTCSATCATVALFAPTP

>KAJ7772818.1 hypothetical protein B0H14DRAFT_2965923 [Mycena olivaceomarginata]
MRAFRALFVPLLGLAGNTTLVEAGPLAYGLCQTGCNTLAVACYAGAGLTFGTVVAAPAAPAAALACNAALGTCCATCATVALFAPTP

>KAJ7832735.1 hypothetical protein B0H14DRAFT_2802403 [Mycena olivaceomarginata]
MHAQLFLPLLALGGMVAARPNATPAYKLCQTACNIRAVACYSAAGLVFGTVVADAAALPVALRCNAALGKCAADCALDTDTTKYV

>KAJ7811222.1 hypothetical protein B0H14DRAFT_3150849 [Mycena olivaceomarginata]
MHAQLFLPLLALSGMVAARPNATPAYKLCQTACNTRAVACYSAAGLVFGTVVADAAAPPVALRCNAALGKCAADCALDTDTITSK

>KAJ7664006.1 hypothetical protein DFH06DRAFT_1189181 [Mycena polygramma]
MKAPFIRTTMRAFKTLFVPLITLAVIPLVQGGPISYALCQTGCNTVAVACYAGAGLVFGTVVAAPAAPAAALACNAALGTCSGVCATVALLAPTP

>KAJ7230373.1 hypothetical protein GGX14DRAFT_553857 [Mycena pura]
MRSSKVNALAIAVSSSASSLVMGGPISYGICQTSCNTVAGACHAAAGVAFEFSVFDAAAAPDVVLKCNEALGICSKTCAGMELFAPTP

>KAJ7230375.1 hypothetical protein GGX14DRAFT_583007 [Mycena pura]
MRAFKVLAIAVSAFPVVMGGPIAYGLCQTGCNVMAVACYAAAGFTFGTVVAAPAAPVAVLGCNAALGTCAATCATVALLAPTP

>KAJ7264656.1 hypothetical protein C8J57DRAFT_1331646 [Mycena rebaudengoi]
MRAFKALFVPLLALVGSTTLVEAGPLAYGLCQTGCNTLAVACYAGAGLTFGTVVAAAAAPAAALACNAALGTCCATCATVALFAPTP

>KAJ7669782.1 hypothetical protein B0H17DRAFT_1209537 [Mycena rosella]
MRPTLVILAPLIVALTAVPAVQGGLISYGLCQTGCNALAVACYAGAGLVFGTVVASPAAPAAALACNVALGQCSAMCATVALLAPTP

>KAJ7691962.1 hypothetical protein B0H17DRAFT_1062265 [Mycena rosella]
MRFFNALLVPIVALPLTQAGPIAYALCQTGCNTVAVACYAGAGLVFGTVVAAPAAPAAALACNLALGTCSATCATIGLFAPTP

>KAF7371167.1 Proteophosphoglycan 5 [Mycena sanguinolenta]
MRFSKIALPLFALSGISLVYAGPIAYGLCQTGCNTVAVACYAGAGLTFGTVVAAAAAPAAAIACNVALGTCSATCATVALLAPTP

>KAF7371160.1 hypothetical protein MSAN_00751400 [Mycena sanguinolenta]
MRLSKFLALPLVALSGISLVQAGPIAYGLCQTGCNTLAVACYAAAGLTFGTVVAAPAAPAAALACNAALGTCSATCATVGLFAPTP

>KAJ6471872.1 hypothetical protein C8R45DRAFT_836297 [Mycena sanguinolenta]
MRFSKILALPLFALSAISFVQAGPLAYGLCQTGCNTVAVACYAGAGLTFGTVIAAAAAPAAAIACNIALGTCSATCATVALLAPTP

>KAF7345529.1 hypothetical protein MVEN_01571500 [Mycena venus]
MRAFNIVFVPLALAGITLVHAGPIAYGLCQTGCNAVAVACYAGAGLVFGTVVAAPAAPAAAITCNVALGTCSATCATVALLAPTP

>KAJ6462394.1 hypothetical protein C8R47DRAFT_1225541 [Mycena vitilis]
MHAFKTLFVPLIALAAIPLVQGGPIAYALCQTGCNTVAVACYAGAGLVFGTVVAAPAAPAAALACNAALGTCSGVCATVGLLAPTP

>KAJ6517837.1 hypothetical protein DFH09DRAFT_1048583 [Mycena vulgaris]
MRPTLVILAPLIVALTAVPAVQGGLISYGLCQTGCNVLAVACYAGAGLVFGTVVASPAAPAAALACNAALGQCSTMCATVALLAPTP

>KZT19035.1 hypothetical protein NEOLEDRAFT_1183737 [Neolentinus lepideus HHB14362 ss-1]
MRFYITVLPLLAALAVVPSTNAGIIAYGICQTGCNTVAVACYAAAGFTFGTVVAAAAAPAAILGCNSALGTCSAACAATALIAPIP

>XP_047899200.1 uncharacterized protein B0H18DRAFT_976291 [Neoantrodia serialis]
*MKIPAALIAATALALATPAFA*GPIAYGICQTGCNTVVVACYAAAGVTFGTVIAAPATPAVILGCNAALGTCSAACATIALFAPTP

>PPQ74617.1 hypothetical protein CVT24_004163 [Panaeolus cyanescens]
MQFKLNTLATSAMLAMTLRPSTVSAGPIAYGICQTGCNVVAVACYAAAGFTFGTIAAPLAPPAILGCNAALGTCSATCATVALLAPTP

>PPQ74615.1 hypothetical protein CVT24_004161 [Panaeolus cyanescens]
MQFKLSALATSTVVAMTLLPATVSAGPIAYGICQTGCNAVVVACYAAAGVTFGTVAAALAPPAIVACNAALGTCSATCATVALLAPTP

>KAF9042497.1 hypothetical protein BJ165DRAFT_1349092 [Panaeolus papilionaceus]
MQLKLNRTFTSAAVLAVALLPTVNAGFITYGICQTGCDSLAVACYAAAGFTFGAVAAPLALPAIVPCNIALGTCSATCAAITFFSPI

>KAF9042500.1 hypothetical protein BJ165DRAFT_268954 [Panaeolus papilionaceus]
MQFKLNRTFTSAALLAMALLPTVNAGLITYGLCQTGCNTLAVACYAAAGFTFGTIAAPLAPPAIVACNGALGTCSATCAAITLLSPI

>KAF9042504.1 hypothetical protein BJ165DRAFT_1529892 [Panaeolus papilionaceus]
MQFKLNRTFTTAALLAMALLPTVNAGLITYGICQTGCNTVVVACYAAAGFTFGTVAAPLAPPAIVACNGALGTCSATCAAITLLSPI

>KAF9042501.1 hypothetical protein BJ165DRAFT_268894 [Panaeolus papilionaceus]
MQFKLNRTFTSTAVLVMVLLPSVNAGNINYGICQTECNTVVVACYTAAGFTFGTVVAPLAPPAIVACNNALGTCSAACAAITLGSPI

>KAI0071738.1 hypothetical protein K474DRAFT_1668679 [Panus rudis PR-1116 ss-1]
MKILAVAVTSFSLIGQASAGLVAYGLCQTGCNALVMACYGAAGAVFGTVAAPAAPPAILACNAALGTCSSACAATALIAPIP

>KAI0071739.1 hypothetical protein K474DRAFT_1668680 [Panus rudis PR-1116 ss-1]
MMFKWTVLLASVVALAAPAKGGPIAYGICQTGCNTVAVACYAGAGFTFGTVVAAAAAPAAVLACNAALGTCSAACATVALFAPTP

>KAF8845957.1 hypothetical protein BDN67DRAFT_960614 [Paxillus ammoniavirescens]
MNLKSLAALTVAASAAPLVMAGPLAYGLCQTGCNVLVGACYAGAGFTFGVTIVGAPPAILACNTGLGACMAACAATALIAPTP

>KAF8835601.1 hypothetical protein BDN67DRAFT_975095 [Paxillus ammoniavirescens]
MNLKSLATLTLAAPLVIASPLAYYNYGACQTGCNTRAGACYTDAGFVFGAVTAAGAPPAILACNDNLGTCMAACAVTARPVRVR

>KIJ15618.1 hypothetical protein PAXINDRAFT_11736 [Paxillus involutus ATCC 200175]
MNLKSLAVLTAVASAAPLVMAGPLAYGLCQTGCNALVGVCYAGAGFVFGATIVAAPPAILACNGGLGTCMAACAVTALLAPTP

>KIJ13640.1 hypothetical protein PAXINDRAFT_80856 [Paxillus involutus ATCC 200175]
MNLKSPAALALAAPLVQVIASPLAYYNYGACQTGCNTGAGACYTAAGFVFGNVTTAEAPPAILTCNASQGTCMAACARASSP

>KIJ15617.1 hypothetical protein PAXINDRAFT_169061 [Paxillus involutus ATCC 200175]
MNLKSLVVLTVVASAAPLVTAYYVICQTGCNVLACACYVGVGFTFGVTIVAAPPAILACNAGLGTCMAACAATALIGPIP

>KIJ15626.1 hypothetical protein PAXINDRAFT_76972 [Paxillus involutus ATCC 200175]
MNLKSLVVLTVVASAAPLVTAGPLAYAICQTGCNVLACACYGGAGFVFGVTIVGVPPAILACNAGLGTCMAACAATALIAPIP

>KIJ15625.1 hypothetical protein PAXINDRAFT_169066 [Paxillus involutus ATCC 200175]
MNLKSLVALTVAASATPLVMAGPIAYGLCQTGCNSLLGACYAGVGFTVGVTIVGAPPAILACNAGLGTCMAACAVTCLFAPTP

>KIK73891.1 hypothetical protein PAXRUDRAFT_836129 [Paxillus rubicundulus Ve08.2h10]
MNFKALAALTVAASAAPLVTAGPLAYGLCQTGCNVLVGACYAGAGFTFGVTIVGAPPAILACNAGLGTCMAACAATALLAPTP

>KIK77344.1 hypothetical protein PAXRUDRAFT_167182 [Paxillus rubicundulus Ve08.2h10]

MNLKRLVALTITVSAAPLVMAGPIAYALCQMGCNVLAYACYAGVGFTFGMTIIGAPPTILACNASLGTCMAACAATAPIP
>VDB91597.1 unnamed protein product [Peniophora sp. CBMAI 1063]
MKLILPLPVLLAPAAVSAGPIAYGICQTGCNAVAVACYAAAGATFGTVIAAAAAPAAIIACNSALGTCSAACATVTLLAPTP
>KZV67638.1 hypothetical protein PENSPDRAFT_47648 [Peniophora sp. CONT]
MKLTLPLILLAPAVVSAGPIAYGICQTGCNAVVVACYAGAGAVFGTVVAAPAAPAAILACNAALGTCSAACATVALFAPTP
>THH07108.1 hypothetical protein EW145_g3608 [Phellinidium pouzarii]
MRASLIALAAVSAAFPAVLGGPLAYGVCQTGCNVAAGTCYAAAGFTFGTVLVATAPASIMACNALLGSCMVACAGMAIAPTL
>KAH8106389.1 hypothetical protein DFH11DRAFT_1640741 [Phellopilus nigrolimitatus]
MRFSLRVFALAAAAASLPGALGEPFAPAFCQSSCYAGIAACYGAAGLTFGTVPVAGPPAITVCNALFGTCMAGCEALG
>KAH8106391.1 hypothetical protein DFH11DRAFT_1518269 [Phellopilus nigrolimitatus]
MRLSPLALLAAALPGALGGPLAYGLCQTGCNAGVVACYGGAGFTFGTVPVIGAPAAILGCNALLGTCMAACAGLVLAPTP
>KAH7883341.1 hypothetical protein F5I97DRAFT_1815740 [Phlebopus sp. FC_14]
MNLKSLAAIALAAAAAPIAMGGPIAYGLCQTGCNVVAVACYAGAGFTFGVTIVGAPPAIMACNAALGTCMATCATVALFAPTP
>KAF8808935.1 hypothetical protein BYT27DRAFT_7136999 [Phlegmacium glaucopus]
MRLSSLSKSAIALIIMTTSVSAGPIAYGICQTGCNAVVVACYAAAGFTFGTVVASPAAPAVLLACNAGLGVCSASCATVALFAPTP
>KAF8808937.1 cysteine-rich protein [Phlegmacium glaucopus]
MHFSSLSKSALTIMIISMMPTSVFAGPIAYGICQTGCNALAVTCYAAAGYTFGTVVASPAAPAVVQSCNAGLAACSTACATVALLAPTP
>KAF9472595.1 hypothetical protein BDN70DRAFT_915839 [Pholiota conissans]
MRFSNVVAPIICALAATSTVMAGPIAYGLCQTGCNTVAVACYASAGMTFGTVVAAAAAPPLILGCNAALGTCSAMCATVALLAPTP
>KAF8176973.1 hypothetical protein BJ912DRAFT_986571 [Pholiota molesta]
MRFSATFATPLLFALASTSIVQAGPIAYGICQTGCNGLAVACYAGAGFTFGTVVAAPAAPAAIMACNAALGSCSAMCASVALFAPTP
>KAF8176972.1 hypothetical protein BJ912DRAFT_986570 [Pholiota molesta]
SPLLTFLPPLVFGLANTSIVQAGPIAYGLCQTGCNTVAVACYAAAGFTFGTVIAAPAAPAAIMACNAALGSCSAMCASVALFAPTP
>KAJ3487682.1 hypothetical protein NLI96_g3372 [Physisporinus lineatus]
MNSKITALVILATGISQVIAGPVAYGICQTGCNVLAGACYAAAGFTFGTVAAPLAPPAILACNTALGSCSAACAAAILLPTP
>KAI0373461.1 hypothetical protein BV20DRAFT_1049871 [Pilatotrama ljubarskyi]
MKLASVVAHIALVFAAVPYAQAGLLSYGICQTGCNTMAVACYAAAGAVFGTVTAGVGTPAVILGCNAALGKCSAACAIVALTPTP
>KAI0373460.1 hypothetical protein BV20DRAFT_1033802 [Pilatotrama ljubarskyi]
MNFKLSALLAALGTLTTVPTVTAGPIAYGICQTGCNAVVVACYAGAGAVFGTITAGVGTPAAIIACNVALGQCSAACAVVAFTPTP
>KIM87109.1 hypothetical protein PILCRDRAFT_815570 [Piloderma croceum F 1598]
MRLTTPLTLLAAAAAATPVAGGPLAYAACQTGCNGLAVACYAAAGFTFGVTIVAVPPAIMGCNIGLGTCMATCATVGLFAPTP
>KAI6004897.1 hypothetical protein EDD15DRAFT_1023739 [Pisolithus albus]
MNVKLPLILALGSLPAAMAGPIAYAICQTGCNVLAGSCYAAAGFTFGTVAAPAAPPMIVACNAGLGTCMAACAATALLAPIP
>KAI6102071.1 hypothetical protein EDD16DRAFT_1647392 [Pisolithus croceorrhizus]
MNFKLLSLLALSSLPVAMAGPIAYGICQTGCNVVAGACYAAAGVTFGTVAAPAAPPLVVACNAALGTCMAACAATALLAPIP
>KAI6098867.1 hypothetical protein EV401DRAFT_992401 [Pisolithus croceorrhizus]
MNLKLPIILALSSLPVATAGPLAYALCQTGCNMLAVGCYSAAGFTFGTVAAAAAPPLILACNAAQGTCMAACAATALLAPIP
>KAI6111997.1 hypothetical protein EDD16DRAFT_1605525 [Pisolithus croceorrhizus]
MNFKLLSLLALSSLPVAMAGPIAYGICQTGCNVLAGACYAAAGVAFGTVAAPAAPPMIVTCNAGLGTCMAACAATALLAPIP
>KAI6131194.1 hypothetical protein EV401DRAFT_1920563 [Pisolithus croceorrhizus]
MNPKLLSLLALCSLPVAMAGPIAYGICQTGCNVVAGACYAAGGVTFGTVAAPAAPLLIASCNAALGTCMAACATTALLAPTP
>KAI6009872.1 hypothetical protein EDC04DRAFT_2772360 [Pisolithus marmoratus]
MNFKLLSLLALGSLPIAMAGPLAYAVCQTGCNMVAVTCYSVAGFTFGAVAAPAAPPLILACNAAQGACMATCAATALLAPIP
>KAI6017553.1 hypothetical protein BKA83DRAFT_4321045 [Pisolithus microcarpus]
MNFKLPLILALSSLPIAMAGPIAYAVCQTGCNALVSTCYAAAGFTFGTVIAGVAPPMIVACNAGLGTCMAACAATALLAPIP
>KAI6037129.1 hypothetical protein BKA83DRAFT_4172100 [Pisolithus microcarpus]
MNFKLPLILALSSLPVAMAGPIAYAVCQSGCNALVGTCYAAAGFTFGTVIAGAAPAMIVACNAGLGTCMTGCATTALLAPIP
>KAI6017551.1 hypothetical protein BKA83DRAFT_678298 [Pisolithus microcarpus]
MNFRLPLILALSSLPLAMAGPFAYAVCQTGCNVLAGSCYAAAGFTFGTVAAPAAPPMIVACNAGLGTCMAACAATALLAPIP
>XP_051595875.1 uncharacterized protein F5J12DRAFT_858457 [Pisolithus orientalis]
MNLKLLGLLALSSVPVAMAGPFAYALCQTGCNMVAVGCYAASGFTFGTVAAPAAPQMILACNAAQGACMAACAATALLAPIP
>XP_051595080.1 uncharacterized protein F5J12DRAFT_864180 [Pisolithus orientalis]
MNLKLLSIITLSSTPVAMAGPLAYAACQTGCNMIAVGCYSVAGFTFGTVAAAVAPPMILACNAAQGTCMAACAATALWAPIP
>KAI6155686.1 hypothetical protein BKA82DRAFT_991315 [Pisolithus tinctorius]
MNLKLLSIVTLSSLPVAMAGPLAYAACQTGCNMIAVGCYSVAGFTFGTVAAAAAPPMILACNAAQGTCMAACAATALWAPIP
>KAI6148428.1 hypothetical protein BKA82DRAFT_992208 [Pisolithus tinctorius]
MAGPLAYAACQTGCNMLTVGCYSLAGFTFGTVAAPAAPPLILACNAAQGTCMAACAATALWAPIP
>KAI6147894.1 hypothetical protein BKA82DRAFT_1006901 [Pisolithus tinctorius]
MNLKLLSLLALSSVPVAMAGPFAYALCQTGCNTVTVACYAAAGFTFGSVAAAAAPSLLVGCNTAQGACMAACAATALLAPVP
>KAG9220390.1 hypothetical protein CCMSSC00406_0006655 [Pleurotus cornucopiae]
MRFSKLAPVSLVLAALSRVEAGPIAYGLCQTGCNTVTVACYAAAGFTFGTVIAAPATPAVLLACNAALGVCSATCATVALFAPTP
>KAF9497993.1 hypothetical protein BDN71DRAFT_1386764 [Pleurotus eryngii]
MHFSKLAPVAVVLAALSGVEAGPIAYGLCQTGCNAVAVACYAAAGFTFGTVIAAPAAPAAILACNAALGACSATCATIGLFAPTP
>XP_036630286.1 uncharacterized protein PC9H_007131 [Pleurotus ostreatus]
MRFSKLAAAAVLAALSGVEAGPIAYGLCQTGCNTVAVACYAAAGFTFGTVIAAPAAPAAVLACNAALGACSATCATIGLFAPTP
>KAF7793897.1 hypothetical protein EIP86_005019 [Pleurotus ostreatoroseus]
MNFKAVVIVSFALAAAHQVSAGLIAYGICQTVACYAGAGFTFGTVVAAALAPPAILACNAALGTCSAACATVALLAPTP
>XP_036630288.1 uncharacterized protein PC9H_007133 [Pleurotus ostreatus]
STTLSSVKASSTATALSRVEADPIAYGLCQTGCNAVAVSCYAAAGFTFGTVIATPEAPAAVLACNAALGACSATCATGLVAPTS
>KAF4567255.1 hypothetical protein EYR36_010873 [Pleurotus pulmonarius]
MHFSKLAPAAVVLTALSGVQAGIIAYGICQTGCNVVAVACYAAAGFTFGTVAAPAAPAAILACNAALGTCSSACATVGLLAPTP
>KAF4597889.1 hypothetical protein EYR38_006281 [Pleurotus pulmonarius]
MHFSKLAPVAVLAALSGVQAGPIAYGLCQTGCNTVAVACYAGAGFTFGTVIAAPAAPAAVLACNAALGACSATCATIGLFAPTP
>KII83275.1 hypothetical protein PLICRDRAFT_148229, partial [Plicaturopsis crispa FD-325 SS-3]
FSTALIVAALPAVTSAGPLAYGLCQTGCNTLAVACYAGAGFTFGTVIAAAATPAALVACNAALGTCSATCATVALFAPTP
>TFK72753.1 hypothetical protein BDN72DRAFT_835845 [Pluteus cervinus]
MRFTTLASALVAMAIPAVQAGPLAYAICQTGCNALVVACYAGAGATFGTIAAPAAPAAIIACNSALGTCSAACAATALIAPTP

>RDX45665.1 hypothetical protein OH76DRAFT_1558969 [Polyporus brumalis]
MRFALVAALVALVAVQTAEAGPLAYGLCQSGCNALAVACYGAAGAVFGTVTAGVGVAPAIIGCNAALGTCTAACAATALIAPTP
>XP_024343044.1 hypothetical protein POSPLADRAFT_1132910 [Postia placenta MAD-698-R-SB12]
MKCTAVLAALAAIAITPVNGGPIAYGICQTGCNAVVVACYAAAGFQFGTVIAAAAAPATILACNAALGTCSATCATVALFAPIP
>XP_024343832.1 hypothetical protein POSPLADRAFT_1037841 [Postia placenta MAD-698-R-SB12]
MKFTAAAAFAALALMTASPVGAGPIAYGVCQTGCNAVAVACYAAAGFQFGTVVAAVAAPATILACNAALGTCSTMCATVALFAPTP
>EED79690.1 predicted protein [Postia placenta Mad-698-R]
MKFTAAATFAALALMTASPVGAGPIAYGICQTGCNTVAVACYAAAGFQFGTVVAAAAAPATILACNAALGSCSAMCATVALFAPTP
>TFL02353.1 hypothetical protein BDV98DRAFT_565686 [Pterula gracilis]
MRLNVTLPLIAFTGSAHAGLIAYGLCQTGCNTLAVACYSAAGFTFGTVVAAAATPAVLVACNTGLGTCSAACAATALIAPIP
>XP_047747560.1 hypothetical protein JR316_0008532 [Psilocybe cubensis]
MRFLQLFALSGIGLLPIASAGPIAYGICQTGCNTVAVACYAAAGLTFGTVVAAPAAPAAAIAYNAALGTCSAACATVALLAPTP
>KAF5322938.1 hypothetical protein D9619_002267 [Psilocybe cf. subviscida]
MRASILVGPIALALASVTSVQAGPIAYGICQTGCNVVAVACYAGAGLTFGTVVAAAAAPAAALACNAALGTCSTACATVALSAPTP
>KAF5322937.1 hypothetical protein D9619_002266 [Psilocybe cf. subviscida]
MRVSNLIAPIALALASTTSVAAGPIAYGICQTGCNTLVVACYAAAGLTFGTIVAAPAAPAAALACNAGLGTCSAACATVALFAPTP
>KAF5322939.1 hypothetical protein D9619_002268 [Psilocybe cf. subviscida]
MRISNLIALIALALVSTTSVAAGPIAYGICQTGCNTLAVACYTAAGLTFGTIVAAPAAPAVALACNAALGTCSAACATVALFAPTP
>XP_007380194.1 hypothetical protein PUNSTDRAFT_18994, [Punctularia strigosozonata HHB-11173 SS5]
SIASFLTASAALVSAGPISYGICQTGCNIVAVACYAAAGFTFGTVAAPLAPPAILGCNAALGTCSAGCAAFLVAPIP
>OCB91128.1 hypothetical protein A7U60_g1610 [Sanghuangporus baumii]
MRLPIFPILATAASLSTVLGGPLAYCACQTACNAGVVTCYAAAGLTFGTVIAAPAAPAAAIACNSVLGVCMAACAASFLAPIP
>KAI4520142.1 hypothetical protein K525DRAFT_204403 [Schizophyllum commune Loenen D]
MRLSILFAPLALAATVAGGPIAYGICQTGCNTVAVACYAAAGFTMGVALPAAPPAILACNAALGTCSAACATIGLFAPTP
>KLO07918.1 hypothetical protein SCHPADRAFT_944882 [Schizopora paradoxa]
MRFNVILPVVVVVASAQNVLGGLLAYGLCQTGCNVLAAACYAAAGATFGTIAAPAAPAAVVGCNGALGTCMAACSVAVIAPTP
>KLO14569.1 hypothetical protein SCHPADRAFT_939493 [Schizopora paradoxa]
MRFSVVLPVVVLASAQNVLGGLLAYGICQTGCNVVAAACYAAAGATFGTIAAPVAPAAILGCNSALGTCMAACSVAVFAPTP
>KIM63914.1 hypothetical protein SCLCIDRAFT_116417 [Scleroderma citrinum Foug A]
MNFKAIATVVLLATPVVMAGPIAYGLCQTGCNILAGACYATAGCVFGTVAAPTAPAAILACNSAQGTCMAICAVVALAAPVP
>KIM55694.1 hypothetical protein SCLCIDRAFT_1220978 [Scleroderma citrinum Foug A]
MNFKALAALTALAAAPVVTAGPLAYALCQTGCNGLAVACYTAAGFTFGVALPAAPPMILGCNVALGTCMGACAVTALIAPIP
>EGO00833.1 hypothetical protein SERLA73DRAFT_133897 [Serpula lacrymans var. lacrymans S7.3]
MNLKSTAALLVVAASAPAALGGPLAYAMCQTGCNGLAVACYAAAGFTFGVTIVAAPPAIMACNVGLGTCMATCATVGLFAPTP
>XP_007316625.1 hypothetical protein SERLADRAFT_436264 [Serpula lacrymans var. lacrymans S7.9]
MNLKSTAALLVVAASAPAALGGPLAYAMCQTDCNRLAVECYAAAGFTFGVTIVAAPPAIMACNVGLGTCMATCATVGLFAPTP
>XP_007316624.1 hypothetical protein SERLADRAFT_436263 [Serpula lacrymans var. lacrymans S7.9]
MNFKSIAALVLVAATAPTVRGGPLAYAACQTGCNVLAVACYGAAGFTFGVTVVAGPPALIACNVGLGTCMATCATVALFAPTP
>KZT35398.1 hypothetical protein SISSUDRAFT_1051499 [Sistotremastrum suecicum HHB10207 ss-3]
MRLPFIALVALVPAMTGSGVAAGPLAYAACQAGCASLVMACYSAAGFVWGATLGVAAPPTIIACNVGYGTCQAACAGAALVAPTP
>KZS96226.1 hypothetical protein SISNIDRAFT_450866 [Sistotremastrum niveocremeum HHB9708]
MRLPFIALVALVPAMTGSGVAAGPLAYAACQAGCASLVMACYSAAGFVWGATLGVAAPPAIIACNVGYGTCQAACAGAALVAPTP
>KIJ22769.1 hypothetical protein M422DRAFT_276754 [Sphaerobolus stellatus SS14]
MRFSKLILITLPITLVQGGIIAYGICQTGCNTVAVACYAGAGFVFGTVVAAPLAPPAIIACNSALGVCSAACAATALIAPIP
>XP_007306302.1 hypothetical protein STEHIDRAFT_61372 [Stereum hirsutum FP-91666 SS1]
MVRVVPVALLAVLSSIPFVTGGPIAYGICQTGCNTVAVACYAAAGFQFGTVVAAVAAPATILACNAALGTCSTACATVALFAPTP
>KAG2033209.1 hypothetical protein BDR03DRAFT_925875 [Suillus americanus]
MNFKSLALFLTAAAVPQVVVAGPLAYGICQTGCNALVVACYAGAGFTFGVALPAAPPVLIACNVGLGTCMAACAAAVFAPTP
>KAG2033206.1 hypothetical protein BDR03DRAFT_872495 [Suillus americanus]
MNFKFLAVLLTAAAVPQAVVAGPLAYAICQTAGCNGLAVACYAGAGFTFGVAVPLAPPALLACNAGLGGCMAACAVVALTPTP
>KAG0702442.1 hypothetical protein DFH29DRAFT_921581 [Suillus ampliporus]
MNFKSLAVLLTAAAAPVAAGPLAYGICQTGCNGMAVACYAAAGCTFGVALPLAPAAIVGCNASLGACMVACAAVALAPTP
>KAG0695044.1 hypothetical protein DFH29DRAFT_957234 [Suillus ampliporus]
MNSKSTTVTLLCTAAAAPAVAGPLGYAICQMGCNGISVACYSAAGFTFGVALPAAPPAIVAYNVALGACMAACAAVALGPTP
>KAG0702443.1 hypothetical protein DFH29DRAFT_921586 [Suillus ampliporus]
MNFKSLAVLLTAAAAPVAAGPLAYAICQTGCNGMAVACYAAAGFTFGVAMPLAPAVIIGCNTSLGTCMVACAAVALAPTL
>KAG0701671.1 hypothetical protein DFH29DRAFT_925384 [Suillus ampliporus]
MNFRFTTVLLLTAAAAPAVAGPLGYAICQTGCNGIAVACYAGAGFTFGVALPAAPPAIIACNAALGACMAACALVALGPTP
>XP_041312473.1 uncharacterized protein EDB93DRAFT_1076476 [Suillus bovinus]
MNLKSLALLVTAAVVPHAVVAGPLASDYGICQTGCNGVAVACYTRAGFTFGVALPTVPPALAECNLALGRCMADCAAVTAPIL
>XP_041312481.1 cysteine-rich protein [Suillus bovinus]
MNFKSLAALLTAAAVPQVVVAGPLSYAICQTGCNGVAVACYAGAGFVFGVALPLAPPAVLACNTALGGCMAACAVVAFMPIP
>XP_041312474.1 uncharacterized protein EDB93DRAFT_1119984 [Suillus bovinus]
MNLKSLALLLAATAVPQVVVAGPLGYAICQTGCNGLAVACYAGAGFTFGVALPAAPPVLIACNTALGGCMAACAAVALAPTL
>KAG2752972.1 hypothetical protein P692DRAFT_20910151 [Suillus brevipes Sb2]
MNLKSITVLLTAAAVPVFAGPLGYALCQTGCNGLAVACYAGAGFTFGVALPAAPPVLLACNSALGGCMAACAVVALTPTL
>KAG2752971.1 hypothetical protein P692DRAFT_20798458 [Suillus brevipes Sb2]
MNFKSLALLLTAAAVPQAVVAGPLAYAICQTGCNGLAVACYAGAGFTFGVALPAAPPVLIACNAALGGCMAACAVVALTPTL
>KAG2747675.1 hypothetical protein P692DRAFT_20849092 [Suillus brevipes Sb2]
MKFKSITVAILLLTTGPTVAGPIGYAICQTGCNGIAVACYSAAGFTFGVAPPAAPPAIIACNAALGACMAACAAVALGPTP
>XP_041202747.1 cysteine-rich protein [Suillus clintonianus]
MNFKSIAVLLTAAAAPAYAGPLAYGICQTGCNGLAVACYAGAGFTFGVALPVAPAAVVGCNVALGGCMAACAVVALTPTL
>KAG2087744.1 hypothetical protein BD769DRAFT_468671 [Suillus cothurnatus]
MNFKSLALLLTAAAVPQAVVAGPLAYGICQTGCNGLAVACYAGAGFTFGVALPAAPPVLIACNVALGGCMATCAAIALAPTL
>KAG2117668.1 hypothetical protein BD769DRAFT_1049885 [Suillus cothurnatus]
MNFKSLALLLTAAVVPQTVVAGPLAYAMCQTGCNGVAVACYSAAGFTFGVALPLAPPVLIACNVALGGCMATCAAIALAPTL
>KAG2087746.1 hypothetical protein BD769DRAFT_1680666 [Suillus cothurnatus]

MNFKSLALLLTAAAVPQTVVAGPLAYAMCQTGCNGVAVACYSAAGFTFGVALPLAPPVLIACNVALGGCMATCAAIALAPTL
>KAG2117666.1 hypothetical protein BD769DRAFT_1049810 [Suillus cothurnatus]
MNFKSLALLLTAAAVPQAVVAGPLAYGICQTGCNGLAVACYAGAGFTFGVALPAAPPVLIACNVALGGCMAGCAAVALAPTL
>KAG2087743.1 hypothetical protein BD769DRAFT_1680663 [Suillus cothurnatus]
MNFKSLALLLTAAAVPQAVVAGPLAYGICQTGCNGLAVACYAAAGFTFGVAVPAAPPVLLACNAGLGGCMAACAAVALTPTP
>KAG2062618.1 cysteine-rich protein, partial [Suillus decipiens]
MNFKSLALLLTAAAIPQAVVAGPIGYAICQTGCNCLAVACYTGAGFTFGVPIPGSTPAAIMACNAGLGTCMAACAA
>KAG2074957.1 hypothetical protein BDR04DRAFT_1070862 [Suillus decipiens]
MNFRSLALFLAAAAVPQAVVAGPLAYGICQTGCNCLAVACYAGAGFTFGVPIPGSTPAAIMACNAGLGTCMAACAAIALTPTP
**>XP_041291730.1 uncharacterized protein F5147DRAFT_699559 [Suillus discolor]**
**MKFKSISTILLLAATAGPAVAGPIGYAICQTGCNGIAVACYSAAGFTFGVAPPAAPPAIIACNTALGACMAACAAVALGPTP**
>XP_041290277.1 uncharacterized protein F5147DRAFT_776163 [Suillus discolor]
MNLKSLALLLTAAAAPQVVVAGPLAYAICQTGCNGLAVACYAGAGFTFGVAVPLAPPALIACNVGLGGCMAACAAVALAPTL
>XP_041290278.1 uncharacterized protein F5147DRAFT_706829 [Suillus discolor]
MNFKSLALLLTAAAVPQVAVAGPLAYGICQTGCNGLAVACYAGAGFTFGVALPLAPPVLIACNVALGSCMVACATVVFAPTL
>XP_041290280.1 uncharacterized protein F5147DRAFT_639062 [Suillus discolor]
MNFKSLALLLTATAIPQVVVAGPLAYAICQTGCNGLAVACYAGAGFTFGVALPAAPPVLLACNAALGGCMAACAVVALTPTP
>XP_041224654.1 uncharacterized protein F5891DRAFT_1111359 [Suillus fuscotomentosus]
MNFKSLALLLTAAAVPQVVVAGPLAYGICQTGCNGLAVACYAGAGFTFGVALPAVPPALMACNVGLGGCMAACAAIALAPTLE
>XP_041224652.1 uncharacterized protein F5891DRAFT_954595 [Suillus fuscotomentosus]
MNLRSLALLLTATAIPQVVVAGPLAYAICQTGCNGLAVACYAGAGFTFGVALPAAPPVLLACNAALGGCMAACAVVALTPTP
>XP_041224653.1 uncharacterized protein F5891DRAFT_1279217 [Suillus fuscotomentosus]
MNLKSLTLLLTAAAVPQVVVAGPCLYAICQAGCIGLVVTCYAGAGFTLVVAPPLAGPAVIACNVAFGGCTLACAALMFAPTP
>KAG2048022.1 hypothetical protein BDR06DRAFT_896315 [Suillus hirtellus]
MNFKSLALLLTATAIPQVVVAGPLAYAICQTGCNGLAVACYAGAGFTFGVALPAAPPALLACNAALGGCMSACAVVALTPTP
>KAG2048018.1 hypothetical protein BDR06DRAFT_943444 [Suillus hirtellus]
MNFKSLALLLTAAAAPQVVVAGPLAYGVCQTGCNALVVACYAGAGFTFGVALPLAPPAIIACNVGLGTCMAGCAAVALAPTL
>KAG1742009.1 hypothetical protein EDB19DRAFT_691958 [Suillus lakei]
MKYKSIAVFLLTATAGPAVAGPIGYAICQTGCNGIAVACYSAAGFTFGVAAPAAPPAIIACNAALGACMAACALVALGPTP
>KAG1734527.1 hypothetical protein EDB19DRAFT_1156156 [Suillus lakei]
MNFKSIALLLTAAAVPRAVIAGPLAYAICQTGCNGLAVACYAGAGFTFGVALPAAPPVLIACNVALGGCMTACAAVALAPTL
>KIK35201.1 hypothetical protein CY34DRAFT_96534 [Suillus luteus UH-Slu-Lm8-n1]
MKFKSITVAILLLATGPAVAGPIGYAICQTAGCNGIAVACYSAAGFTFGVAPPAAPPAIIACNAALGACMAACAAVALGPTP
>KIK36068.1 hypothetical protein CY34DRAFT_811597 [Suillus luteus UH-Slu-Lm8-n1]
MNLKSITLLLTAAAVPVFAGPLGYALCQTGCNGLAVACYAGAGFTFGVALPAAPPVLLACNSALGGCMAACAVVALTPTL
>KAG1759280.1 hypothetical protein EDD22DRAFT_906775 [Suillus occidentalis]
MKFKSITVAILLLATGPAVAGPIGYAICQTGCNGIAVACYSAAGFTFGVAPPAAPPEIIACNAALGAYMAACAAVALGPTP
>KAG1762396.1 hypothetical protein EDD22DRAFT_778446 [Suillus occidentalis]
MNFKSLALLLTAAAVPQAVVAGPLAYAICQTGCNGLAVACYAGAGFTFGVALPAAPPVLIACNVGLGGCMAACAAVALTPTP
>XP_041171419.1 uncharacterized protein EDB91DRAFT_1061537 [Suillus paluster]
MNLKSIVVLLTAVAAPAVAGPLGYAICQTGCNSLAVACYGAAGFTMGVALPAVPAVLVSCNVGLGTCMAACAVVALSPTL
>XP_041169262.1 uncharacterized protein EDB91DRAFT_248283 [Suillus paluster]
MNLKSIAVLLTAVAAHAVTGPIEYASCQTGCNGHAVSCYGAAGFTFRVALPSVPPVLATCNTGLGTCMAACADLLSERVQL
>XP_041171032.1 uncharacterized protein EDB91DRAFT_1254637 [Suillus paluster]
MNFKYITVLLAATAAPAVAGPLGYAICQTGCNGIAVACYAGAGFTFGVALPAAPPAIIACNAALGACMAACAVIALGPTP
>KAG1774592.1 hypothetical protein EV702DRAFT_1200173 [Suillus placidus]
MNFKSLALLLTAAAVPQAVVAGPLAYAICQTGCNGLAVACYAGAGFTFGVALPAAPPVLIACNVALGGCMTACAAVALAPTL
>KAG1774591.1 hypothetical protein EV702DRAFT_1031834 [Suillus placidus]
MNLKSIALLLTAAAVPQAVVAGPLAYGICQTGCNGLAVACYAGAGFTFGVALPAAPPALIACNAALGGCMAACAVVALAPTL
>XP_041155836.1 uncharacterized protein HD556DRAFT_847530 [Suillus plorans]
MNFKSLALLLTAAAVPQVAVAGPLAYGVCQTGCNALVVACYAGAGFTFGVALPLAPPALIACNVGLGTCMAACAAVALAPTP
>XP_041155838.1 uncharacterized protein HD556DRAFT_1244954 [Suillus plorans]
MNFKSLALLLTAAAAPQVAVAGPLAYAICQTGCNGLVVACYAGAGFTFGVALPLAPPALLACNVGLGTCMAACAVVALTPTL
>XP_041160953.1 uncharacterized protein HD556DRAFT_1269841 [Suillus plorans]
MKFKSISTILLLTATAVPAIAGPIGYAICQTGCNGIAVACYSAAGFTFGVAPPAAPPAIIACNTALGACMAACAVVAFGPTP
>XP_041160366.1 uncharacterized protein HD556DRAFT_1443095 [Suillus plorans]
MNFKSLALLLTAAAVPQVVVAGPLAYGICQTGCNGLAVACYAGAGFTFGVALPAVPPVLMACNVGLGGCMAACAAIALAPTL
>XP_041160372.1 cysteine-rich protein [Suillus plorans]
MNFKSLALLLTATAIPQVVVAGPLAYAICQTGCNGLAVACYAGAGFTFGVALPAAPPVLLACNAALGGCMAACAVVALTPTL
>KAG2353321.1 cysteine-rich protein [Suillus spraguei]
MNFKSLALFLTAAAVPQAVVAGPLAYTICQTGCNCLAVACYSAAGFTFGVLIPGVTPAAIMACNVGLGTCMAACAAIAFGPTP
>KAG2353322.1 cysteine-rich protein [Suillus spraguei]
MNFKSLALLLTVAAAPQAVMGGPLAYGICQTGCNCLAVACYAGAGFTFGVPIPGSTPAAIMACNAGLGTCMAACAAIALTPTP
>KAG2360710.1 hypothetical protein BDR07DRAFT_1359757 [Suillus spraguei]
MNFKSLALFLTAAAVPQAVVAGPIAYAICQTGCNCLAVACYSAAGFTFGVPIPGVTPAAIMACNAGLGTCMAACAAIAFGPTP
>XP_041237613.1 uncharacterized protein DFJ58DRAFT_749481 [Suillus subalutaceus]
MNFKSLALLLTAAAVPQAVVAGPLAYAICQTGCNGLAVACYAGAGFTFGVAVPLAPPALLACNAGLGGCMAACAVVALTPTP
>XP_041237617.1 uncharacterized protein DFJ58DRAFT_815892 [Suillus subalutaceus]
MNFKSLALFLTAAAVPQVVVAGPLAYGICQTGCNGLAVACYAAAGFTFGVALPAAPPVLIACNVGLGGCMAACAVVALTPTP
>XP_041237615.1 uncharacterized protein DFJ58DRAFT_815877 [Suillus subalutaceus]
MNFKSLALLLTAAVVPQAVVAGPLAYGICQTGCNGLAVACYAGAGFTFGVALPAAPPVLIACNVGLGGCMAACAAVALAPTL
>XP_041237616.1 uncharacterized protein DFJ58DRAFT_668908, partial [Suillus subalutaceus]
CSTGCNGLAVACYAGAGFTFGVALPLAPPAVIACNVALGGCMAACAVIALAPTL
>XP_041185215.1 uncharacterized protein BJ212DRAFT_1489758 [Suillus subaureus]
MNFKSLALFLTAAAVPQAVVAGPLAYGICQTGCNGLAVACYAGAGFTFGVALPLAPPALIACNVALGGCMATCAVVALAPTL
>XP_041185216.1 uncharacterized protein BJ212DRAFT_1291350 [Suillus subaureus]
MNFKSLALFLTAAAVPQAVVAGPLAYGICQTGCNGLAVACYAAAGFTFGVALPAAPPALVACNVGLGGCMAACAAVALAPTP

>KAG1882508.1 hypothetical protein F4604DRAFT_1619034 [Suillus subluteus]
MNFKSFALFLTAAAVPQLVVAGPLAYGICQTGCNGLAVACYAAARFTFGVALPAAPPVLIACNVGLGGCMAACAVVALTPTP
>KAG1839191.1 hypothetical protein C8R48DRAFT_91879 [Suillus tomentosus]
MKFKSITTILILAATAGPVVAGPIAYAICQTGCNGIAVACYSAAGFTFGVALPAAPPAIMACNAALGACMAACAAIALGPTP
>KAG1855037.1 hypothetical protein C8R48DRAFT_660633 [Suillus tomentosus]
MKFKSIGIILVLLAATAGPAVAGPIGHAICQTAGCNGIAVACYSAAGFTFGVAPPAAPPAIIACNTALGACMAACAAVAFGPTP
>KAG1839194.1 hypothetical protein C8R48DRAFT_621836 [Suillus tomentosus]
MKFKSTTTILILAATAGPVVAGPIAYAICQTAGCNGIAVACYSAASFTFGVALPAAPPAIMACNAALGACVAACAAIALGPTP
>KAG1875880.1 hypothetical protein C8R48DRAFT_591628 [Suillus tomentosus]
MNFKSLALLLTAAAAPQVVVAGPLAYGICQTGCNALVVACYAGAGFTFGVALPLAPPAIIACNVGLGTCMAACAAVALAPTL
>KAG2338307.1 hypothetical protein BDR05DRAFT_704677 [Suillus weaverae]
MNLKSVALLLTAAAIPQAVVAGPLAYGICQTGCNGLAVACYAGAGFTFGVALPAAPPALIACNSALGGCMAACAVVALAPTV
>KAG2338305.1 hypothetical protein BDR05DRAFT_893898 [Suillus weaverae]
MNFKSLALLLTAAAVPQAVVAGPLAYAICQTGCNGLAVACYAGAGFTFGVALPAAPPVLVACNVALGGCMTACAAVALAPTL
>CEL57659.1 hypothetical protein RSOLAG1IB_02402 [Rhizoctonia solani AG-1 IB]
MKLSITSVVAFVTVALNAGQVQAGPIAMGLCYSACNAGYVTCCISAGAVAGTFTLGLGTPVALAACSVVQGACMSACTPLLLAPTP
>KAF8707836.1 hypothetical protein RHS03_04129, partial [Rhizoctonia solani]
MKFSIASAAALAALSLNIGQVEAGPIAMGLCYTACNAGYVTCCTVAGVTAGTFTLGLGIPAAVAACSVVQGACMAACTPLLVAPTP
>CAE6474563.1 unnamed protein product [Rhizoctonia solani]
MKFSFASLVAFVAVALNAGQVQAGPVAMGLCYTACNAGYVTCCASAGAVAGTFTLGLGVPAALAVCSVVQGTCMAACTPLLAAPTP
>CAE6474554.1 unnamed protein product [Rhizoctonia solani]
MKFSFAPIVALATLALNAGQVQAGPIAMGMCYSACNAGYVTCCASAGAVAGTFTAGLGIPAALAACSVVQGTCMAACTPLLAAPTP
>CAE6506335.1 unnamed protein product [Rhizoctonia solani]
MKFSIAPVVALATLALSAGQVQAGPIAMGMCYSACNAGYVTCCTAAGVTAGTFTLGLGIPAAVAACSVVQGTCMAACTPLLAAPTP
>CAE6512876.1 unnamed protein product [Rhizoctonia solani]
MKFSFASVVAFAAVALNAGQVQAGPIAMGMCYSACNAGYVTCCTTAGITAGTFTLGLGIPAAIAACSVVQGTCMAACTPLLVAPTP
>CAE7122953.1 unnamed protein product [Rhizoctonia solani]
MKFSVAPIVALATVALNAGQVQAGPIAMGLCYSACNAGYVTCCTSAGAIAGTFTLGLGIPAALAGCSVIQGTCMAACTPLLAAPSP
>CUA77418.1 hypothetical protein RSOLAG22IIIB_02431 [Rhizoctonia solani]
MKFSFTSVVAFAAVALNAGQVQAGPIAMGLCYSACNAGYVTCCASAGAIAGTFTLGLGVPAALAVCSVVQGTCMAACTPLLAAPTP
>CEL57606.1 hypothetical protein RSOLAG1IB_02349 [Rhizoctonia solani AG-1 IB]
MKLSVASAVAFAVLALNADQVQAGPIAMGLCYTACNAGYVTCCVTAGAIAGTFTLGLGIPAAVAACSVIQGACMAACTPLLVVPTP
>CAE6449415.1 unnamed protein product [Rhizoctonia solani]
MKFSIAPIATLATLALNAGQVQAGPIAMGMCYSACNARYVTCCASAGTVAGTFTLGLGVPAALAACSVVQGTCMVACTPLLAAPSP
>CAE6512885.1 unnamed protein product [Rhizoctonia solani]
MKFSLASVFAFVAVALNASQVQAGPIAMGMCYSACNAGYVTCCTAAGTTAGTFTLGLGVPAAVAACSVVQGACMAACVPLGVAPIP
>CAE6470859.1 unnamed protein product [Rhizoctonia solani]
MKLTLAPALALVAFTLNARHVHAGPVAMGACYTACNAGYVACCATAGITAGIFTLGAGVPAALGACSAVQGVCMAACVPLGLAPTP
>EUC53909.1 transmembrane protein, putative [Rhizoctonia solani AG-3 Rhs1AP]
MKFSFTSVVAFAAVALNAGQVQAGPVAMGLCYSACNAGYVTCCTTAGAVAGTFTLGLGVPVALAVCSVVQGTCMAACTPLLAAPTP
>CAE6407853.1 unnamed protein product [Rhizoctonia solani]
MKFSITSVFAFAALALNAGQVQAGPVAMGLCYSACNAGYVTCCASAGAVAGTFTLGLGVPVALAACSVVQGACMSACTPLLLAPTP
>KAH7345108.1 hypothetical protein B0J17DRAFT_763814 [Rhizoctonia solani]
MKFSFASVVAFAAVALNAGQVQAGPVAMGLCYSACNAGYVTCCASAGAVAGTFTLGLGIPAALAVCSVVQGTCMAACTPLLAAPTP
>KAJ1311556.1 hypothetical protein OPQ81_010040 [Rhizoctonia solani]
MKLSIASALACVIPTLNARHVHAGPIFMAACYSACNAGYVTCCAAAGATIGVFTLGLGVPATLAACSAVQGTCMAACVPLGVAPTP
>CAE6407261.1 unnamed protein product [Rhizoctonia solani]
MKFSLASLAAFAALAFNAGQVQAGPIAMGLCYSACNAGYVTCCASAGATAGTFTLGLGVPAALAGCSVIQGACMSACTPLLAAPTP
>CAE6520723.1 unnamed protein product [Rhizoctonia solani]
MKLTITSTLAFVVLTLNARHVHAGPVAMGACYTACNVGYVTCCAGAGATVGLFTLGLGVPAALAACSVVQGACMAACVPLGLAPTP
>KAH7345111.1 cysteine-rich protein [Rhizoctonia solani]
MKLTITSAFALVVLALNTRHVHAGPATMGACYTACNVGYVTCCATAGVTAGVFTLGLGVPAALGACSVVQGACMAACVPLGFAPTP
>CAE6449405.1 unnamed protein product [Rhizoctonia solani]
MKLTVSSVFAFVLLALNARHAHAGPISMGAYYTACNVGYATCCATAGTTAGIFTLGLGVPAALAACSAVQGACMAACVPLGAAPTP
>KDN50300.1 hypothetical protein RSAG8_01636, partial [Rhizoctonia solani AG-8 WAC10335]
MKLTITSTLAFVVLTLNARHVHAGPVAMGACYTACNAGYVTCCAGAGATVGLFTLGLGVPAALAALAACSVVQGACMAACVPLGLAPTP
>EUC53908.1 zygote-specific protein, putative [Rhizoctonia solani AG-3 Rhs1AP]
MKFSIAPLVAFATLALNAGQVQAGPIAMGMCYSACNAGYVTCCVSAGTVAGTFTLGLGVPAAIAACSAVQGTCMAACTPLLLAPTP
>CUA77419.1 hypothetical protein RSOLAG22IIIB_02432 [Rhizoctonia solani]
MKFSIAPMVTLATLALNAGQVQAGPIAMGMCYSACNAGYVTCCASAGTVAGTFTLGLGIPAALAACSVVQGTCMAACTPLLAAPSP
>KAH7345109.1 hypothetical protein B0J17DRAFT_28587 [Rhizoctonia solani]
MKFSVAPIVALATLALNVGQVQAGPIAMGLCYSACNAGYVTCCAAAGVTAGTFTLGLGVPAALAGCSIVQGACMAACTPLLAAPTP
>KAJ8594645.1 hypothetical protein M405DRAFT_808728 [Rhizopogon salebrosus TDB-379]
MNYKHTAILLVAAIASPAVVAGPLGYAICQTGCNALAVACYAGAGFTFGVALPAAPPVIIACNAGLGTCMAACAVVALGPTP
>OJA16009.1 hypothetical protein AZE42_10625 [Rhizopogon vesiculosus]
MNIKSAISLITAAAAAPAVVAGPLGYAICQTGCNALVVSCYAGAGFTFGVALPAAPPVIMACNAGLGTCMAACAVVALGQPI
>OJA20466.1 hypothetical protein AZE42_04909 [Rhizopogon vesiculosus]
MNIKSTISLIIAAAAAPAVVAGPLGYAICQTGCNALVVSCYAGAGFTFGVALPVAPAVIVACNAGLGTCMAACAVVAFSPTL
>OAX33574.1 hypothetical protein K503DRAFT_514430 [Rhizopogon vinicolor AM-OR11-026]
MNIKSTISLIIAAAAAPAVVAGPIGYAICQTGCNALVVSCYAGAGFTFGVALPAAPPVIMACNAGLGTCMAACAVVALGQPL
>KAF9074370.1 hypothetical protein BDP27DRAFT_1214363 [Rhodocollybia butyracea]
MPVLSVLVGLQGAVAGPIAYGLCQTGCNTMAVACYAAAGFTFGTVIAAAAAPVAVLGCNAALGTCSATCATVALFAPTP
>TFY54996.1 hypothetical protein EVJ58_g8527 [Rhodofomes roseus]
MKTPYALIAALALAVATPASAGPIAYGLCQTGCNTVVVACYAAAGFTFGTVIAAPAVPAVILGCNAALGTCAATCATVALFAPTP
>TDL19155.1 hypothetical protein BD410DRAFT_792400 [Rickenella mellea]
MRFSSLLSSAIALAIAPGVLGGPIAYGICQTGCNTVAVACYAAAGFQFGTVVAAAAAPATILACNAALGTCSAMCATVALLAPTP
>KAJ7604390.1 hypothetical protein FB45DRAFT_1043797 [Roridomyces roridus]

```
MRFTTLLAPIALVAVPLVQGGLISYGLCQTGCNTVVVACYAGAGLVFGTVVAAPAAPAAALACNAALGTCSATCAAVVLLAPIP
>KAF5332937.1 hypothetical protein D9758_015962 [Tetrapyrgos nigripes]
MRFSLPSLLRITVAVATFQSVQAGLLAYGICQTGCNTLAVACYSAAGATFGTVVASAATPAAILACNAALGKCSAACAAVTLVAPTP
>KAI0360877.1 hypothetical protein OH77DRAFT_1517275 [Trametes cingulata]
MHFKLSALLVALGALTTAPAALAGPAAYGICQTGCNVVAVACYAAAGAVFGTVTAGVGTPAAILACNVALGQCSAACALVVLAPTP
>KAI0360878.1 hypothetical protein OH77DRAFT_1586382 [Trametes cingulata]
MKLTSIIAPIAFVIAAVPYAQAGLLSYGLCQTGCNTVAVACYAAAGATFGTITAGAGTPAVILGCNAALGKCSATCAALTLLAPIP
>CDO70453.1 hypothetical protein BN946_scf184496.g2 [Trametes cinnabarina]
MKLSTILVPVTLALGSFQSAKAGILSYGLCQTGCNSLAVACYAAGGFTFGTVTAGAGVPAVVLGCNAALGTCMAACAAVALAPIP
>OSC99408.1 cysteine-rich protein [Trametes coccinea BRFM310]
MKLSVVVVPLAIAFSCLPSARAGLLSYGVCQTGCNALAVACYAAAGYTFGTVTAGLGTPAVVLGCNAALGKCSAACAVVALTPIP
>OSC99407.1 hypothetical protein PYCCODRAFT_842717 [Trametes coccinea BRFM310]
MNVKSSALAAVLVIAAAVPATTAGPIAYGICQTGCNAVAVACYAGAGAVMGTVTAGVGTPVAVLACNVALGQCSAACATVALFAPTP
>KAJ8495237.1 hypothetical protein ONZ51_g1809 [Trametes cubensis]
MKLSAFFIPIALSLGALPSANAGIIGYGICQTGCNVVAVACYAAAGYTFGTVTAGLGTPAVILGCNAALGKCSAACAIVALTPTP
>KAI0824053.1 hypothetical protein BC628DRAFT_1323633 [Trametes gibbosa]
MFAKPSSVFLAFLAALAAVPTTQAGPLAYGVCQTGCNALVVACYASAGAVFGTVTAGVGTPAAIIACNVALGQCSAACALVALTPTL
>KAI0768850.1 hypothetical protein BD413DRAFT_614326 [Trametes elegans]
MQLRPIALLAALLSVSAMPGAYAGPIAYGICQTGCNAVAVACYAAAGAVFGTVTAGVGTPAAILGCNVALGQCSAACAVVAFTPTP
>KAI0666621.1 hypothetical protein C8Q78DRAFT_1058448 [Trametes maxima]
MQFKAPLGAVILTTLAIIPNAHAGPLAYGICQSGCNALAVTCYAAAGAVFGTVTAGVGTPAAILACNAALGTCATACVAAGFAPTL
>KAI0644389.1 hypothetical protein C8Q79DRAFT_912887 [Trametes meyenii]
MYSKIPLATFLTAVAIIPTAYAGPFSYGICQTGCNTVAVACYAAAGAVFGTVTAGVGTPVAILACNAALGTCSAACIAAGFAPIP
>KAI0633791.1 cysteine-rich protein [Trametes polyzona]
MKLSTVAASIALATSAVPAANAALIGYGICQTGCNTLAVACYAAAGYTFGTVVAGPAAPAVIMGCNAALGKCSAACAVTLLAPV
>OJT05103.1 hypothetical protein TRAPUB_4168 [Trametes pubescens]
MKFSSVVAPLALALATIAPSAYAGPLAYGICQTGCNALAVACYAGAGFTFGTVTAGVGVPAAIVGCNAGLGVCQAACAAAIFAPTL
>KAI8993820.1 cysteine-rich protein [Trametes punicea]
MKLSSIVAPLSVAISALPSAKAGLLSYGICQTGCNTMAVACYAAAGFTFGTVTAGAATPAVILGCNAALGKCSAACAVVALTPTP
>KAI8993819.1 hypothetical protein BD414DRAFT_411344 [Trametes punicea]
MIFKLSAFSVVPLVLAAIQGAEAGPIAYGVCQTGCNAVAVACYAGAGAVFGTVTAGIGTPAAIIACNVALGQCSAACATIALFAPIP
>KAI9059204.1 hypothetical protein FKP32DRAFT_1580281 [Trametes sanguinea]
MNTKLSALTTVLFVFAAVPVATAGPIAYGICQTGCNVVAVACYAGAGAVMGTVTAGVGTPVAVLACNVALGQCSAACATVALFAPTP
>KAI9059205.1 cysteine-rich protein [Trametes sanguinea]
MKLSTVVLPLVLAFSTLPSAKAGLISYGLCQTGCNALAVACYAAAGYTFGTVTAGLGTPAVILGCNSALGQCSAACAVVALSPIP
>XP_008038531.1 cysteine-rich protein, partial [Trametes versicolor FP-101664 SS1]
LAVASASAAPPRTGIQKLAYGVCQTGCNNFATICYNNAGRKFGTVTTDENTPTAILNCNAALGICQQACAKAV
>XP_008033715.1 uncharacterized protein TRAVEDRAFT_42882 [Trametes versicolor FP-101664 SS1]
MKFSVAAPLALVLATAAPSAYAGPLAYGICQTGCNALVVACYAGAGFTFGTVTAGAGVPAAVVACNAGLGVCMAGCAAAIFAPTP
>XP_008034073.1 cysteine-rich protein [Trametes versicolor FP-101664 SS1]
MQFKLSTSLRSLITLAAIVPTAHAGPLAYGICQTGCNIVAVACYAGAGFVFGTVTAGVGTPAAVLACNVALGQCSAACAVVALTPTP
>XP_008042174.1 uncharacterized protein TRAVEDRAFT_66328 [Trametes versicolor FP-101664 SS1]
MKLSTILASLAIAAVSSVHAGPLTDGVCQTGCKAAAAGCYSAAGMVFGAVTAGIATPVVALACNAVLDRCQAECAIQGADAV
>KAI0344501.1 hypothetical protein BDW22DRAFT_1354571 [Trametopsis cervina]
MNFKLAVLSVVAAAAAPAAYAGPLAYAACQTGCNVVAVACYAAAGATFGTVAAPAAPAAIVACNGALGTCSSMCATVGLFAPTP
>KAI0344500.1 hypothetical protein BDW22DRAFT_1427200 [Trametopsis cervina]
MYSKTALLVTFAVVAASLPAAHAGPIAYALCQTGCNAVAVACYAAAGATFGTIAAPAAPAAILGCNAALGSCSATCATIGLFAPTP
>KAF8219361.1 hypothetical protein L208DRAFT_1373477 [Tricholoma matsutake 945]
MRSLAVLIAALAATPSVYAGPIMYAICQTGCNTLAVSCYAAAGFTFGTVAAPAAPAAILACNSSLGVCSAACSTVALCAPTP
>KAG8993678.1 hypothetical protein FRB94_010464 [Tulasnella sp. JGI-2019a]
MRFNRVLATFALALAAVPAAEAGPISYGICQTGCNTVVVACYAAAGFTLGTVAAPTAPAAIVACNSALRTCSATCAFLLLAPIP
>KAG8993679.1 hypothetical protein FRB94_010465 [Tulasnella sp. JGI-2019a]
MRFNRVLATFALALAAVPAAEAGLISYGLCQTGCNTVAVACYAAAGFTFGTVAAPVAPAAIVACNGALGTCSATCASLVLLAPIP
>KAF6743528.1 hypothetical protein DFP72DRAFT_124234 [Tulosesus angulatus]
MRPSILLLPVLAFVSAAQAGPIAYGLCQTGCNTVAVACYAAAGATFGTVVASAAAPAVILACNAALGSCSAGCASFALFAPIP
>KAF6748547.1 hypothetical protein DFP72DRAFT_572925 [Tulosesus angulatus]
MRPSVLLLPVLAFVSAAQAGPIAYGLCQTGCNTVAVACYAAAGATFGTVVASAAAPAAILGCNAALGSCSAGCASFALLAPIP
>KAI0035134.1 hypothetical protein K488DRAFT_83339 [Vararia minispora EC-137]
MKLLALTLLAPALVAAGPLAYAACQTGCNAVAVACYAAAGLTFGTVVAAPAAPAAALACNAALGTCSAACATVALFAPTP
>KAF8651157.1 hypothetical protein AX16_004836 [Volvariella volvacea WC 439]
MRFSALVATLFTVSAPSLVQAGLISYGICQTGCNTVAVACYAAAGFTFGTVAAPLAPPAIVACNSALGTCSAACAVVALPAPVP
```

```
KAF8735275.1        -------MQLFLFNFILPFAAVLCVTTVVQAG-PILYAMCKLG----CDVVANACAAS--
KAI1785556.1        -----MDFRFKAISLLVGVTATIGIVLAVVNGNPSAHEVCQTG----CNAVLVACYGTGG
KAF8661385.1        -------MKLFSFVVPVAAVLLSATSVVQAGP--ISYAICQTG----CNIVAVACYGAAG
XP_007380194.1      ---------------SIASFLTASAALVSAGP--ISYGICQTG----CNIVAVACYAAAG
KAH6901377.1        ----------MRLAL-AVASLIVFIGQVNAGP--IMYGICLAG----CNATAATCYAAAG
KAH6902106.1        ----------MRLTLGAAASLVAFIGQVNAGPTSEPYGDCQTG----CNAIAAKCYTAAG
KAJ3487682.1        ----------MNSKITALVILATGISQVIAGP--VAYGICQTG----CNVLAGACYAAAG
KAF8980955.1        ------------MRLSAVVSPLAFAPLVLGGP--IAYGICQTG----CNTLAVACYAAAG
KAF8977750.1        ------------MRLSAIVAPLAFAPLVLGGP--IAYGICQTG----CNTVAVACYAAAG
KAF8980957.1        ------------MRLSAVVFPLAFAPLVLGGP--IAYGICQAG----CTAAAATCYSAAG
KLO07918.1          ---------MRFNVILPVVVVVASAQNVLGGL--LAYGLCQTG----CNVLAAACYAAAG
KLO14569.1          ---------MRFSVVLPVVVLAS-AQNVLGGL--LAYGICQTG----CNVVAAACYAAAG
XP_038910436.1      --------MKYFNPLILFS-ALALAPSALAGP--FTYGVCQTG----CNVVAVACYAAAG
XP_038921664.1      --------MKYFNPLILFS-ALTLAPSALAGP--FTYGVCQTG----CNVVTVACYAAAG
XP_038922423.1      --------MKYFNPLILFS-TLALAPSALAGP--FTYGVCQTG----CNVVAVACYAAAG
KAG8993678.1        --------MRFNRVLATFALALAAVPAAEAGP--ISYGICQTG----CNTVVVACYAAAG
KAG8993679.1        --------MRFNRVLATFALALAAVPAAEAGL--ISYGLCQTG----CNTVAVACYAAAG
RXW13811.1          ------------MRPSLLFVPVLAASVAQAGP--IAYGICQTG----CNAVVVACYAAAG
RXW22619.1          ------------MRPSLLLIPVLAASTAQAGL--IAYGICQTG----CNAVTVACYAAAG
KAF8651157.1        ---------MRFSALVATLFTVSAPSLVQAGL--ISYGICQTG----CNTVAVACYAAAG
PPQ74617.1          -----MQFKLNTLATSAMLAMTLRPSTVSAGP--IAYGICQTG----CNVVAVACYAAAG
PPQ74615.1          -----MQFKLSALATSTVVAMTLLPATVSAGP--IAYGICQTG----CNAVVVACYAAAG
KAF9042500.1        -----MQFKLNRTFTSAALLAMALLPTVNAGL--ITYGLCQTG----CNTLAVACYAAAG
KAF9042504.1        -----MQFKLNRTFTTAALLAMALLPTVNAGL--ITYGICQTG----CNTVVVACYAAAG
KAF9042497.1        -----MQLKLNRTFTSAAVLAVALLPTVNAGF--ITYGICQTG----CDSLAVACYAAAG
KAF9042501.1        -----MQFKLNRTFTSTAVLVMVLLPSVNAGN--INYGICQTE----CNTVVVACYTAAG
KAK0445166.1        --------MRLSPILTFLVTSLALAPQAHAGP--IAYGICQTG----CNVLAVACYAAAG
KAK0232397.1        --------MRLSPILTFLVTSLAIAPQARAGP--IAYGICQTG----CNVLAVACYAAAG
KAK0192295.1        --------MRLSPIFTFLVTSLALAPQAYAGP--IAYGICQTG----CNVLAVACYAAAG
KAK0435962.1        --------MRLSPILTSLVMSLALAPQAHAGP--IAYGICQTG----CNVLAVACYAAAG
PBK70936.1          --------MRLSPILTFLVTSLALAPQAHAGP--IAYGICQTG----CNVLAVACYAAAG
SJL10225.1          --------MRLSPILTSLVTSLALAHQAHAGP--IAYGICQTG----CNVLAVACYAAAG
KAK0496260.1        --------MRLSPIFTFLATSLALAPQAYAGP--IAYGICQTG----CNVLAVACYAAAG
XP_060325687.1      --------MRLSPVLAFLVTSLALAPQAHAGP--IAYGICQTG----CNVLAVACYAAAG
KAK0211506.1        --------MRLSPIFTFLVTSLTLAPQAYAGP--IAYGLCQTG----CNVVVVACYAAAG
KAK0204044.1        --------MRLSPIFAFLVTSLALAPQAYAGP--IAYGICQTG----CNVVAVACYAAAG
KAK0232409.1        --------MRLSPILTFLVTSVALAPQAHAGP--IAYGICQTG----CNVVAVACYAAAG
KAK0480870.1        --------MRLSPIFTFLVTSLALAPQAYAGP--IAYGICQTG----CNVVAVACYAAAG
PBK94666.1          --------MRLSPILTFLVTSLAIAPQVHAGP--IAYGICQTG----CNVVAVACYAAAG
XP_043040322.1      --------MRLSPIFVFFVTSLALVHQTHAGP--ISYGICQTG----CNALAVACYAAAG
KAK0477555.1        -------MRLQSPIFAFLFTALALAPQAYAGL--IAYGICQTG----CNVLAVACYSAAG
KAK0477562.1        -------MRLQSPIFAFLFTALALAPQAYAGP--IAYGICQTG----CNVLAVACYSAAG
KAH6902114.1        ------------MRLTFTLASLALISQVNAGL--IAYGICQTG----CNALAVACYAAAG
KAF8219361.1        -----------MRSLAVLIAALAATPSVYAGP--IMYAICQTG----CNTLAVSCYAAAG
KAH6901375.1        -----------MRLTLAAASLIAFVGQVNAGL--IMYGICQTG----CNTVAVACYAAAG
KAH6901376.1        -----------MRLTLAVASLVAFVAQVHAGP--IMYGICQTG----CNAVAVACYAAAG
KAH6902116.1        ----------MMRFTIALASLIAFVGQVQAGP--IAYGICQTG----CNAVVVACYAAAG
KAH6902112.1        ---------MRFTTSIAVASLVAFAGQVSAGP--IMYGICQTG----CNVVAVACYAAAG
TFK18913.1          -----------MRLSASLAPIFAFVTLVHAGP--IAYGICQTG----CNAVAVACYAAAG
KAI0344501.1        ---------MNFKLAVLSVVAAAAAPAAYAGP--LAYAACQTG----CNVVAVACYAAAG
KAI0344500.1        -------MYSKTALLVTFAVVAASLPAAHAGP--IAYALCQTG----CNAVAVACYAAAG
KAF9256214.1        ---------MRFASLTTAVLLLVGLQEVKAGP--IAYGICQTG----CNIVAVACYAAAG
XP_043004193.1      ----------MRSTSLATVLLLIGLQEVNAGP--IAYGICQTG----CNVVAVACYAAAG
KAF9014159.1        ------------MRFTPILASLLIAPVVLSGP--ISYGICQSG----CNAVAVACYAAAG
TFK72753.1          ----------MRFTTLASALVAMAIPAVQAGP--LAYAICQTG----CNALVVACYAGAG
KAH9948988.1        ------------MKFSLLVPFALLAASANAGP--IAYGICQTG----CNTVAVACYAAAG
PPR07183.1          -----------MRFNALVSALAVVIPMASAGP--IAYGICQTG----CNTVAVACYAAAG
KAF8898031.1        -------------MRSALLIALPFISMAAAGP--IAYGICQTG----CNTVAVACYAAAG
KAI0785557.1        ----------MNPRSMLFAGLAAMSAAPAAHAGLIAYGICQTG----CNTLAVACYAGAG
KAF7793897.1        -----------MNFKAVVIVSFALAAAHQVSAGLIAYGICQT----------VACYAGAG
KAI0071739.1        -----------MMFKWTVLLASVVALAAPAKGGPIAYGICQTG----CNTVAVACYAGAG
KAF9554637.1        ---------MRINANLLSTVALALSGASMVSAGPIAYGLCQTD----CNTVAVACYADAG
KAF4620962.1        ---------MRINANLLSTVALALSGASIVSAGPIAYGLCQTD----CNTVAVACYAGAG
KAF9554638.1        ---------MRINTNLLSTVALALSGASMVSAGPIAYGLCQTG----CNTVAVACYAGAG
KAF4621497.1        ---------MRFNANILPIAALALSGANMVTAGPIAYGICQTG----CNTVAVACYAGAG
KJA16367.1          -----------MRFS---TLAIALASAASVSAGPIAYGLCQTG----CNVVAVACYGAAG
TDL19155.1          -----------MRFSSLLSSAIALAIAPGVLGGPIAYGICQTG----CNTVAVACYAAAG
KAF5322937.1        ----------MRVSNLIAPIALALASTTSVAAGPIAYGICQTG----CNTLVVACYAAAG
KAF5322939.1        ----------MRISNLIALIALALVSTTSVAAGPIAYGICQTG----CNTLAVACYTAAG
KAF5322938.1        ----------MRASILVGPIALALASVTSVQAGPIAYGICQTG----CNVVAVACYAGAG
KIM37591.1          ----------MHFSKLFAPVAIAIASASVVQGGPIAYGICQTG----CNAVVVACYSAAG
THG95442.1          ------------MNFKVLAAAVLAAVAPVANAGPIAYGICQTG----CNSLAVACYAGAG
KAJ7080818.1        ---------MRTANALVLPLAALSAGVSLVEAGPIAYGICQTG----CNTVTVACYAGAG
GAT42696.1          -----------MNPTKALTALVVAALASTVQAGPLAYGICQTG----CNTVAVACYAGAG
KAF7289020.1        -----------MNPTKALTALVFAAVASTVQAGPLAYGICQTG----CNTVAVACYAGAG
XP_047747560.1      ------------MRFLQLFALSGIGLLPIASAGPIAYGICQTG----CNTVAVACYAAAG
KAJ7669782.1        ---------MRPTLVILAPLIVALTAVPAVQGGLISYGLCQTG----CNALAVACYAGAG
KAJ6517837.1        ---------MRPTLVILAPLIVALTAVPAVQGGLISYGLCQTG----CNVLAVACYAGAG
KAJ7830972.1        ---------MRPTLVILAPLILALTAVPTVQGGLVSYGLCQTG----CNTLAVACYASAG
KAJ6579297.1        ---------MRPTISTLAPLLLALAAVPAVQGGLISYGLCQTG----CNTLAVACYAGAG
KAJ7604390.1        ---------MRFTTLLAP---IALVAVPLVQGGLISYGLCQTG----CNTVVVACYAGAG
KAJ6621514.1        -------------------LLALAT-VPLVQAGPIAYGLCQTG----CNTIAVVCYAGAG
```

```
KAJ6621484.1        ---------MRAFKIIFAPLLSLAT-VPLVQAGPIAYGLCQTG----CNTIAVVCYAGAG
KAJ7036623.1        ---------MRTFNVLFAPLLALAA-IPLVQGGPLAYALCQTG----CNTVVVACYAGAG
KAJ7777091.1        ---------MRAFNVLFAPLLALAA-IPLVQGGPLAYALCQTG----CNTVVVACYAGAG
KAJ7187815.1        ---------MHAFKVLFAPLLALAA-VPLVQAGPLAYALCQTG----CNTVAVACYAGAG
KAJ7147695.1        ---------MRAFNALVLPLLALTG-IPLVQGGPLAYGLCQTG----CNTVAVACYAGAG
KAJ7664006.1        MKAPFIRTTMRAFKTLFVPLITLAV-IPLVQGGPISYALCQTG----CNTVAVACYAGAG
KAJ6462394.1        ---------MHAFKTLFVPLIALAA-IPLVQGGPIAYALCQTG----CNTVAVACYAGAG
KAJ6580769.1        ---------MRASKTLFVPLLALAG-IPFVQGGPLAYGLCQTG----CNTLAVACYAGAG
KAJ7114020.1        ---------MRTFNTLFVPILALAG-IPLVQGGPISYGLCQTG----CNTLAVACYAGAG
KAJ7498176.1        ---------MRAFNSAPLLALALAA-VPLVQGGPIAYALCQTG----CNTLAVACYAGAG
KAJ7712755.1        -------MRPLTLKALLAPLLAFAA-IPLVEGGPIAYGLCQTG----CNTLAVACYAGAG
KAJ7724508.1        -------MRPLALKALLAPLLAFAA-IPLVEGGPIAYGLCQTG----CNTLAVACYAGAG
KAJ7330443.1        ---------MRAFKILIVPLLALSG-ITLVEAGPIAYALCQTG----CNTVAVACYAGAG
KAJ7743748.1        ---------MRAFKILVVPLLALSG-ITLVEAGPIAYALCQTG----CNTVAVACYAGAG
KAJ7208767.1        ---------MRAFKILAVPFIALSG-ISLVQAGPLAYGLCQTG----CNTVAVACYAGAG
KAJ7483446.1        ---------MRAFTAFVVPFVALAG-IPLAHAGPIAYALCQTG----CNTVAVACYAGAG
KAF7345529.1        ---------MRAFNIVFVP-LALAG-ITLVHAGPIAYGLCQTG----CNAVAVACYAGAG
KAF8211017.1        ------------MRAFSKLLLALSG-VSLVHAGPIAYGLCQTG----CNTVAVACYAGAG
KAJ7780066.1        --------MRANALSAPVLALALTA-LPLAHAGPVAYALCQTG----CNTVAVACYAGAG
KAJ7691962.1        ------------MRFFNALLVPIVA-LPLTQAGPIAYALCQTG----CNTVAVACYAGAG
KAJ7359841.1        ---------MRAFRAFFVPLLVLAGNTALVEAGPLAYGLCQTG----CNSLAVACYAGAG
KAJ7772818.1        ---------MRAFRALFVPLLGLAGNTTLVEAGPLAYGLCQTG----CNTLAVACYAGAG
KAJ7264656.1        ---------MRAFKALFVPLLALVGSTTLVEAGPLAYGLCQTG----CNTLAVACYAGAG
KAF7371167.1        ----------MRFSKIALPLFALSG-ISLVYAGPIAYGLCQTG----CNTVAVACYAGAG
KAJ6471872.1        ---------MRFSKILALPLFALSA-ISFVQAGPLAYGLCQTG----CNTVAVACYAGAG
KAF7371160.1        ---------MRLSKFLALPLVALSG-ISLVQAGPIAYGLCQTG----CNTLAVACYAAAG
KAJ7330535.1        ---------MRVFKTIAVPLLALSG-ITLVQAGLIAYAICQTAG---CNTFAVACYAGAG
KZV67638.1          ---------------MKLTLPLILLAPAVVSAGPIAYGICQTG----CNAVVVACYAGAG
VDB91597.1          ---------------MKLILPLVLLAPAAVSAGPIAYGICQTG----CNAVAVACYAAAG
KAI0035134.1        ----------------MKLLALTLLAPALVAAGPLAYAACQTG----CNAVAVACYAAAG
XP_037222833.1      -----------MKLTKAISFILVALSPVLVEAGPIAYGICQTG----CNTVAVACYAAAG
KAF5390836.1        ----------MRFTKASISVLAVFTGLQTAQAGPIAYGICQTG----CNTVTVACYAAAG
THU93403.1          ----------MRLSTVFAPVLVGLGALQSVQAGPIAYGICQTG----CNAVAVACYAGAG
KAH8107856.1        ------------MKLSILTPLAVLAAAPTALGGPIAYGICQTG----CNTVAVACYAAAG
KAI0089603.1        -------------------LLALVAVAGTANAGPIAYGICQTG----CNAVAVACYAAAG
KIY47514.1          -------------MQITKPCLALLAACGLAQAGPIAYGICQTG----CNVVAVACYAATG
KXN83432.1          --------MRPLKVFILVVASILSSSPQQAAAGPIAYGICQTG----CNVVAVACYAAAG
KXN83429.1          --------MRPLKTFILVVASILSSAPQQAAAGPIAYGICQTG----CNIIAVACYAAAG
KXN83427.1          --------MRPLKAFILVVASILSSAPQQEAAGPIAYGICQTG----YNVVVVACYAAVG
KXN93170.1          --------MRPLKAFILVVASILSSAPQQAAAGPIVYGMCQTG----CNVVVAARYAAAG
KXN81170.1          --------MRPFRTLTLVVAAILSSAPQQAMAGLIAYGICQTG----CNTVAVACYGAAG
KXN91089.1          --------MRPFRTLILVVAAVLSSAPQQTMAGPIAYGICQTG----CNVLAVACYAAAG
KXN83428.1          ------------------------------MAGLIGYGICQTG----CNAVAGACYAAAG
KAF5347592.1        -------MRLQFKTCSTIAATILLIQP--ASAGLIAYGICQTG----CNTLAVACYAAAG
TFK33514.1          ----------MRLSTLTATLAMALVYVPTAEAGIISYGICQTG----CNVLAVACYAAAG
KAH9894858.1        ----------MKLSTLFIPVALTLGALPSADAGLLGYGVCQTG----CNALAVACYAAAG
KAI0326571.1        ----------MKLSTFFIPVALTLGALPSANAGLLGYGICQTG----CNAVAVACYAAAG
KAI0656638.1        ----------MKLSAFFIPIALGLGALPSANAGIIGYGICQTG----CNVVAVACYAAAG
KAJ8495237.1        ----------MKLSAFFIPIALSLGALPSANAGIIGYGICQTG----CNVVAVACYAAAG
OSC99408.1          ----------MKLSVVVVPLAIAFSCLPSARAGLLSYGVCQTG----CNALAVACYAAAG
KAI9059205.1        ----------MKLSTVVLPLVLAFSTLPSAKAGLISYGLCQTG----CNALAVACYAAAG
KAI8993820.1        ----------MKLSSIVAPLSVAISALPSAKAGLLSYGICQTG----CNTMAVACYAAAG
CDO70453.1          ----------MKLSTILVPVTLALGSFQSAKAGILSYGLCQTG----CNSLAVACYAAGG
KAI0373461.1        ----------MKLASVVAHIALVFAAVPYAQAGLLSYGICQTG----CNTMAVACYAAAG
KAI0360878.1        ----------MKLTSIIAPIAFVIAAVPYAQAGLLSYGLCQTG----CNTVAVACYAAAG
KAH9851372.1        -----------MKFSTIVSVALGLAAVPSAKAGLLAYGICQTG----CNTMAVACYAAAG
KAI0633791.1        ----------MKLSTVAASIALATSAVPAANAALIGYGICQTG----CNTLAVACYAAAG
KAI0737527.1        --------MSLFRRAIVVAAVTLVALP-STEAGLIAYGICQTG----CNALAVACYAGAG
KAI0744947.1        --------MKLTLPVISTLAISLSAFP-SVHAGLIAYGICQTG----CNTVAVACYAAAG
KAH9851373.1        ------MFFKPSPTVFLTFLAALSAAP-AAHAGPLAYGICQTG----CNALVVACYAGAG
KAI0824053.1        ------MFAKPS-SVFLAFLAALAAVP-TTQAGPLAYGVCQTG----CNALVVACYASAG
KAI0373460.1        --------MNFKLSALLAALGTLTTVP-TVTAGPIAYGICQTG----CNAVVVACYAGAG
KAI0360877.1        --------MHFKLSALLVALGALTTAP-AALAGPAAYGICQTG----CNVVAVACYAAAG
KAI0768850.1        --------MQLRPIALLAALLSVSAMP-GAYAGPIAYGICQTG----CNAVAVACYAAAG
XP_008034073.1      --------MQFKLSTSLRSLITLAAIVPTAHAGPLAYGICQTG----CNIVAVACYAGAG
KAI0666621.1        --------MQFKAPLGAVILTTLAIIP-NAHAGPLAYGICQSG----CNALAVTCYAAAG
KAI0644389.1        ---------MYSKIPLATFLTAVAIIP-TAYAGPFSYGICQTG----CNTVAVACYAAAG
OJT05103.1          --------MKFSSVVAPLALALATIAP-SAYAGPLAYGICQTG----CNALAVACYAGAG
XP_008033715.1      ---------MKFSVAAPLALVLATAAP-SAYAGPLAYGICQTG----CNALVVACYAGAG
THV03464.1          ---------MLLLTPTSVVLLIGLAILQSTQADLIAYGICQTT----CNSAAAACYAAAG
XP_036630288.1      -----------STTLSSVKASSTATALSRVEADPIAYGLCQTG----CNAVAVSCYAAAG
KAJ7036624.1        -------MRTFEILFAILLAVRLPLVQGDKVKDEVAYGLCQTG----CNNITVACYSAAG
KAJ7780067.1        -------MPTFKALLSVIIVAAILLVQG----GPLAYGECQTG----CNNLTVACYSRAG
KAJ7498165.1        -------MRAFNALHILALAAAPLVHGG-----VIPYVECQTG----CNSVAVACYSAAG
KAJ7832735.1        -------MHAQLFLPLLALGGMVAARPN----ATPAYKLCQTA----CNIRAVACYSAAG
KAJ7811222.1        -------MHAQLFLPLLALSGMVAARPN----ATPAYKLCQTA----CNTRAVACYSAAG
XP_008038531.1      ------------LAVASASAAPPRTGIQ-----KLAYGVCQTG----CNNFATICYNNAG
XP_008042174.1      -------MKLSTILASLAIAAVSSVHAG-----PLTDGVCQTG----CKAAAGCYSAAG
OCB91128.1          --------MRLPIFPILATAASLSTVLG----GPLAYCACQTA----CNAGVVTCYAAAG
KAF9461010.1        -------------ALICAFLAIPSAQAG-----PLLYGVCQTD----CNALAVSCYAATG
KAF9255919.1        -----------MHFSFSTVAVLLSMVPWEVNAGLLAYGLCQTG----CNCLAVACYSAAG
KAI0071738.1        -------------MKILAVAVTSFSLIGQASAGLVAYGLCQTG----CNALVMACYGAAG
```

```
TFL02353.1           --------------MRLNVTLPLIAFTGSAHAGLIAYGLCQTG----CNTLAVACYSAAG
KAF5332937.1         ---------MRFSLPSLLRITVAVATFQSVQAGLLAYGICQTG----CNTLAVACYSAAG
KDR81157.1           ---------MRFSIAVAPFLIALCTTTSFVSAGPIFYGICQTG----CNAVAVACYAGAG
KDR81156.1           ---------MRFPTVIAPFLIALSTTVFLVSAGPIEYGICQTA----CNDGAVACYRGAG
XP_007868697.1       ---------MRFYTIALPLLAAMASIP-STIAGPIAYGICQTG----CNALAVACYAGAG
KZT19035.1           ---------MRFYITVLPLLAALAVVP-STNAGIIAYGICQTG----CNTVAVACYAAAG
KDQ57059.1           ---------MRPYTLFLPILAAVATVLPTASAGPIAYGLCQTG----CNALAVACYAGAG
KAK1228354.1         ---------MRFT-TLTAFAVALFATLQGVNGGPIMYGVCQTG----CNCVAVACYSAAG
KAJ8072958.1         ---------MRFT-ALTAGTVALLAALQGVNGGPIAYGICQTG----CNSVAVACYAAAG
ESK93305.1           ---------MRFTNILAPSALALLTGIQGVNGGLIAYGLCQTG----CNTVAVACYAAAG
KIJ22769.1           ---------MRFS-----KLILITLPITLVQGGIIAYGICQTG----CNTVAVACYAGAG
KAF8176973.1         ---------MRFSATFATPLLFALASTSIVQAGPIAYGICQTG----CNGLAVACYAGAG
KAF8176972.1         ----------SPLLTFLPPLVFGLANTSIVQAGPIAYGLCQTG----CNTVAVACYAAAG
KAF6743528.1         -------------MRPSILLLPVLAFVSAAQAGPIAYGLCQTG----CNTVAVACYAAAG
KAF6748547.1         -------------MRPSVLLLPVLAFVSAAQAGPIAYGLCQTG----CNTVAVACYAAAG
XP_040768338.1       ------------MKFTTTLTTLALALATPAAAGPIAYGLCQTG----CNTVAVACYAAAG
XP_040768339.1       ------------MKFTTTLTTIALALATPAAAGPIAYGLCQTG----CNAVAVACYAAAG
XP_040768337.1       --------------------------------GPIAYALCQTG----CNTVAAACYSAAG
KAI0930540.1         -------------MKFFLLSSLALAGVSLVNAGPIAYGLCQTG----CNTVAVACYAAAG
XP_024343832.1       ----------MKFTAAAAFAALALMTASPVGAGPIAYGVCQTG----CNAVAVACYAAAG
EED79690.1           ----------MKFTAAATFAALALMTASPVGAGPIAYGICQTG----CNTVAVACYAAAG
XP_024343044.1       ------------MKCTAVLAALAAIAITPVNGGPIAYGICQTG----CNAVVVACYAAAG
KIK03653.1           ------------MRISTTLLLSPLLMVANVNAGPLAYGLCQTG----CNTVAVACYAAAG
KIJ99165.1           ------------MRISSAILLSPFLMVA--NAGPIAYGLCQTG----CNTVVVACYAAAG
KAF9461003.1         -------------MRFSKILLSAALALPTVQAGLISYGLCQTG----CNTVAVACYAAAG
KAF9528339.1         -----------MRFNTAATLAILAATTSSVMGGPLSYGLCQTG----CNTVAVACYAAAG
TFY71517.1           ------------MRFSYLTIVATMALLPTAMGGPISYAICQTG----CNTVAVACYAAAG
KAA1466928.1         ------------MRFSYLAGAVVFALSPAVMGGPISYAICQTG----CNAVVVACYAAAG
KAA1466913.1         ------------MRLSLLPLAAAAALVPSVLGGPISYAICQTG----CNTVAVACYAAAG
KAJ7230373.1         ---------MRSSKVNALAIAVSSSASSLVMGGPISYGICQTS----CNTVAGACHAAAG
XP_009549554.1       -----------MVRITPLAAVSLLSAIPLVAGGPISYGLCQTG----CNTVAVACYAAAG
XP_007306302.1       -----------MVRVVPVALLAVLSSIPFVTGGPIAYGICQTG----CNTVAVACYAAAG
KDQ08146.1           ----------MRVFSLAAFVIAPFYLATGAYAGPIAYGLCQTG----CNTLAVACYAAAG
KAJ7586376.1         --------MVGITKISALLVTSMAFLAIPVTAGPIAYGLCQTG----CNTLAVACYAAAG
KII83275.1           ----------------FSTALIVAALPAVTSAGPLAYGLCQTG----CNTLAVACYAGAG
KAF8153443.1         -------------------------MSTAVLAGPIAYGICQTG----CNALVVSCYAAAG
RDB15947.1           -----------MRLSILAPLIFALSATQGVKGGPIAYGICQTG----CNALVVSCYAAAG
KAF8808935.1         -------MRLSSLSKSAIALIIMTTS---VSAGPIAYGICQTG----CNAVVVACYAAAG
KAF8808937.1         -------MHFSSLSKSALTIMIISMMPTSVFAGPIAYGICQTG----CNALAVTCYAAAG
KAF8869089.1         -------------MRFSSALLVAASMAPVVLGGPISYGICQSG----CYAAAVICYAAAG
KAF8869829.1         -------------MRFSTALLIAASMAPVALGGPISYGICQSG----CNAVVVACYAAAG
KAF8872941.1         -------------MRFSTAFLVTLGMAPVALGGPIAYALCQTG----CNSLAVACYAAAG
KAF8872939.1         -------------MRFSTTFLVALGMAPVALGGPIAYGICQTG----CNSLAVACYAAAG
KAF8957624.1         ----------MRFSLIAAPILYVLASTSIAQAGPIAYGLCQTG----CNVMAVACYAGAG
KAF9074370.1         -----------------MPVLSVLVGLQGAVAGPIAYGLCQTG----CNTMAVACYAAAG
KAJ3874754.1         ----------MRLTNVLVPVLSVLAGLQGAKAGLIAYGLCQTG----CNTVAVACYAAAG
KAJ3868585.1         ----------MRLTNVLVPVLSVLAGLQGAKAGLIAYGLCQTG----CNSVAVACYAAAG
KAJ3870732.1         ----------MRLTNVLVPILSVLAGLQGAQAGPIAYGLCQTG----CNIVAGACYAAAG
KAJ4480927.1         ----------MRLTNTLLPFLSVLAGLQSAQAGIIAYGICQTG----CNVAAGACYTAAG
KAJ3742886.1         ----------MRLTNILLPILPVLAGMQSAQAGPIAYGLCQTG----CNVVAAACYAAAG
KAJ3729090.1         ----------MRLTNILLPILPVLAGMQSAQAGPIAYGLCQTG----CNVVAAACYAAAG
KAJ3998398.1         ----------MRLTNILLPILPVLAGMQSAQAGPIAYGLCQTG----CNVVAVACYAAAG
KAJ3793355.1         ----------MRLTNILLPILPVLAGMQSAQAGPIAYGLCQTG----CNTVVVACYAAAG
KAJ3727548.1         ----------MRFTNILLPVLSVLAGMHNVQAGPIAYGLCQTG----CNVVAVACYAAAG
KAF9472595.1         ----------MRFSNVVAPIICALAATSTVMAGPIAYGLCQTG----CNTVAVACYASAG
KAJ7230375.1         -------------MRAFKVLAIAVSAFPVVMGGPIAYGLCQTG----CNVMAVACYAAAG
KIL66627.1           ----------MQLYKIALPLAMALASSVTVSAGPIAYGICQTG----CNTVAVACYAAAG
THH27703.1           ------------MKFSALSALAVLATTPFVAGGPIAYGICQTG----CNTLAVACYAAAG
KAI0705164.1         -------------------LFTTLAAAATVNGGPIAYGICQTG----CNTVAVACYAGAG
KAI4520142.1         --------------MRLSILFAPLALAATVAGGPIAYGICQTG----CNTVAVACYAAAG
TRM64791.1           --------------MRVTAILAPVALATAVAAGPIAYGICQTG----CNTLAVACYAAAG
XP_036630286.1       -----------MRFSKLA-AAAVLAALSGVEAGPIAYGLCQTG----CNTVAVACYAAAG
KAF4597889.1         -----------MHFSKLA-PVAVLAALSGVQAGPIAYGLCQTG----CNTVAVACYAGAG
KAF9497993.1         -----------MHFSKLAPVAVVLAALSGVEAGPIAYGLCQTG----CNAVAVACYAAAG
KAG9220390.1         -----------MRFSKLAPVSLVLAALSRVEAGPIAYGLCQTG----CNTVTVACYAAAG
KAF4567255.1         -----------MHFSKLAPAAVVLTALSGVQAGIIAYGICQTG----CNVVAVACYAAAG
KAK0445201.1         ----------MRLSRAFACLATSLVLAPQAYAGPIAYGLCQTG----CNTMAVACYAAAG
SJL12908.1           ----------MRLSRTFVCLATSLVFAPQAYAGPIAYGLCQTG----CNAMAVACYAAAG
PBK94627.1           ----------MRLSRAFACLATSLVLAPQAYAGPIAYGICQTG----CNTMAVACYAAAG
KAK0232361.1         ----------MRLSRAFACLATSLVLAPQAYAGPIAYGICQTG----CNTMAVACYAAAG
KAK0211537.1         ----------MRLSRAFACLATFLVLVPQAYAGPIAYGICQT------------------
KAK0496299.1         ----------MRLSRALACLATSLVLAPQAYAGPIAYGLCQTG----CNTMAVACYAAAG
PBK70903.1           ----------MRLSRAFACLATSLVLAPQAYAGPIAYGLCQTG----CNTMAVACYAAAG
KAK0192253.1         ----------MRLSRAFACLATSLVLAPQAYAGPIAYGICQTG----CNTMAVACYAAAG
KAK0480837.1         ----------MRLSRAFACLATSLVLAPQAYAGPIAYGICQTG----CNTMAVACYAAAG
PBK78949.1           ----------MRLSRAFACLATSLVLAPQAYAGPIAYGICQTG----CNTVAVACYAAAG
KAK0204073.1         ----------MRLSRAFAFLATSLALAPQVHAGPIAYGICQTG----CNTVVVACYAAAG
XP_060325653.1       ----------MRLSRAFAFLATSLALVPQAHAGPIAYGICQTG----CNTVVVACYAAAG
XP_043040323.1       ----------MRLSRVFAFLATSLALAPQAHAGPIAYGICQTG----CNGMAVACYAAAG
KJA16363.1           --------------------AVALASIGSANAGLITYGICQTG----CNTVAVACYAAAG
KJA16366.1           -----------MRLSILAPLAVALASIGSANAGLITYGICQTG----CNTVAVACYAAAG
KZT72154.1           -----------MKTPFALIG-AALAMAAPAFAGPIAYGICQTG----CNTVAVACYAAAG
```

```
KZT72155.1          -----------MKIPFALTG-AALAMAAPVSADPIALVICLIG----CNTVAVACYAAAG
XP_047899200.1      -----------MKIPAALIAATALALATPAFAGPIAYGICQTG----CNTVVVACYAAAG
TFY54996.1          -----------MKTPYALIAALALAVATPASAGPIAYGLCQTG----CNTVVVACYAAAG
KAI0693946.1        -----------MRFAILAALAAIVAVPTAEAG-PLAYAICQTG----CNSLVVACYANAG
RDX45665.1          -----------MRFALVAALVALVAVQTAEAG-PLAYGLCQSG----CNALAVACYGAAG
RPD73715.1          -----------MRFVLATVLVALVAVPTAVDAGPLAYGICQTG----CNAVVVACYGAAG
KAI0744948.1        ---------MNVKLLSIAVVLSTLPALPVYAG-PLAYALCQTG----CNAVVVACYGAAG
KAI0779875.1        ---------MNFKLSALSALAVLYVVPTVEAG-PLAYGLCQTG----CNAVVVACYGAAG
KAI0737528.1        --------MNFARLSLLSAAALYMTVPAVQAG-PIAYGICQTG----CNSVAVACYAAAG
KAI0656637.1        ----------MNFISF-AVLLTLVACAATADAGPIAYGLCQTG----CNAVVVACYAAAG
KAH9894857.1        ----------MNFKSFPALLLTFIACAATVDAGPIAYGLCQTG----CNAVVVACYAAAG
OSC99407.1          ---------MNVKSSALAAVLVIAAAVPATTAGPIAYGICQTG----CNAVAVACYAGAG
KAI9059204.1        ---------MNTKLSALTTVLFVFAAVPVATAGPIAYGICQTG----CNVVAVACYAGAG
KAI8993819.1        ---------MIFKLSAFSVVPLVLAAIQGAEAGPIAYGVCQTG----CNAVAVACYAGAG
XP_007366826.1      -------MNLRLSTLVIVATGL-LAASPIVNAGPVAYGICQTG----CNAVVVACYAGAG
XP_007370845.1      -------MRFHLSTLAIAATSL-LAVFPTVTAGPIAYGLCQTG----CNTVTVACYAGAG
PIL23384.1          -------MHLKLSALTLALVG--LAASPVANAGPIAYGICQTG----CNTVAVACYAGAG
KAI1794409.1        -------MHFKLSALALALVG--LAASPVANAGPIAYGICQTG----CNTVTVACYAAAG
KAI1791328.1        -------MQLKLSALALALAG--LAASPVVNAGPIAYGICQTG----CNAVVVACYAGAG
PIL33906.1          -------MQLKLSFKLSAALAVTLSLPQVANAGPIAYGICQTG----CNVVAVACYAGAG
XP_047873742.1      ----------MHFTPSSLLAAAVLLATGAHAGPVAPYGVCQTG----CVLVAVACYSAAG
XP_047873750.1      ----------MHFKLSSFLAAAALRATCGSQAGPMAYGICQTG----CNKGVVACYAGAG
XP_047873746.1      ----------GELKLSSLLAATALITTGVHAG-PALYGVCQTG----CNAVTVACYAGAG
XP_047873743.1      -----------MQFKLSSLLAAALLATGAQAG-PALYGICQTG----CNTLAFACYAGAG
XP_047873744.1      -----------MQFKLSSLLAAALLATGASAG-PAFYGICQTG----CNTLAVACYAGAG
XP_047873745.1      -----------MQLKPSLLAAAALLATGARAS-SVAYDVCQTAG---CNTVAVACYAGAS
XP_047870886.1      ----------MLFKLSSLVAAAAILATGAHAG-PIAYGICQTG----CNAVVVACYAGAG
XP_047873747.1      ----------MLFKLSSLVAAAAILATGARAG-PIAYGICQTG----CNTVAVACYAGAG
KAF9018149.1        ------------MVRFQRLALLALPFIPAINAGPIAYGICQTG----CNAVAVACYAGAG
KAH8831031.1        -----------MRLSHLFMAFAGMALAPTGAAGPLAYAVCQTG----CNTIAVACYAAAG
KJA27646.1          -----------MRLAVLTTLAVGAATAT---AGPIAYGVCQTE----CNTVAEACYTAAG
KAI0323098.1        -------------MKLSLIVALLAATAPTVFAGPIAYGICQTG----CNVVAVACYAAAG
KAI0323099.1        -------------MKLFSILTLLTLSTPAVIAGPIAYGICQTG----CNVVAVACYAAAG
PFH45591.1          -------MRLTRVFAPLGIIVALSTVPQIVQAGPILYGICQTG----CNSLAVICYAAGG
KIM63914.1          -----------MNFKAIATVVLLATP--VVMAGPIAYGLCQTG----CNILAGACYATAG
EJD44225.1          --------------MTLITSALLLAFAAPASASLILYGICQTG----CNMGAVSCYGVAC
KZV97636.1          --------------MKPSRIVLPLTLVLSANAGLIAYGICQTG----CNMGAVACYAVAG
KAH7096490.1        --------MKLIRPTRLIATTLVLLAPTQVRASLIAYGICQTG----CNIGAVTCYAAAG
KAH7096489.1        --------MKVIRPTRLIASTLVLLAPTQVHAGLIAYGICQTS----CNILAVACYGAAG
KAH7090935.1        --------MKFLHPITRLATALALLAPTQVRAGIIAYGLCRTG----CNVIVMGCYGAAG
KAI6004897.1        -------------MNVKLPLILALGSLPAAMAGPIAYAICQTG----CNVLAGSCYAAAG
KAI6017551.1        -------------MNFRLPLILALSSLPLAMAGPFAYAVCQTG----CNVLAGSCYAAAG
KAI6017553.1        -------------MNFKLPLILALSSLPIAMAGPIAYAVCQTG----CNALVSTCYAAAG
KAI6037129.1        -------------MNFKLPLILALSSLPVAMAGPIAYAVCQSG----CNALVGTCYAAAG
KAI6102071.1        -------------MNFKLLSLLALSSLPVAMAGPIAYGICQTG----CNVVAGACYAAAG
KAI6111997.1        -------------MNFKLLSLLALSSLPVAMAGPIAYGICQTG----CNVLAGACYAAAG
KAI6131194.1        -------------MNPKLLSLLALCSLPVAMAGPIAYGICQTG----CNVVAGACYAAGG
XP_051595080.1      -------------MNLKLLSIITLSSTPVAMAGPLAYAACQTG----CNMIAVGCYSVAG
KAI6155686.1        -------------MNLKLLSIVTLSSLPVAMAGPLAYAACQTG----CNMIAVGCYSVAG
KAI6098867.1        -------------MNLKLPIILALSSLPVATAGPLAYALCQTG----CNMLAVGCYSAAG
KAI6148428.1        ------------------------------MAGPLAYAACQTG----CNMLTVGCYSLAG
KAI6009872.1        -------------MNFKLLSLLALGSLPIAMAGPLAYAVCQTG----CNMVAVTCYSVAG
XP_051595875.1      -------------MNLKLLGLLALSSVPVAMAGPFAYALCQTG----CNMVAVGCYAASG
KAI6147894.1        -------------MNLKLLSLLALSSVPVAMAGPFAYALCQTG----CNTVTVACYAAAG
KAI9568904.1        ---------MNFKSLAALTLTASAAPFAAAGPLA--YAICQTG----CNVLAVSCYGAAG
KAF8132063.1        ---------MNIKSLAALTLAASAAPLVAAGPLA--YALCQTG----CNALVVSCYAAAG
KAF9237865.1        ---------MNFKSLAALTLVASAAPLTMAGPLA--YGLCQTG----CNVLVGSCYAAAG
KIJ66076.1          ---------MNFKALAALTLAASAAPLTMAGPLA--YAACQTG----CNVLAVACYAGAG
KAF8415388.1        ---------MNFKSLAALTLAASAAPLAAAGPLA--YALCQTG----CNTLAVSCYAAAG
KAF8131977.1        ---------MNFKSLAALTLAASAAPLAAAGPLA--YALCQTG----CNTLAVSCYAAAG
KAG6382117.1        ---------MNFKSLAALTLAASAAPLAAAGPLA--YAVCQTG----CNTLAVSCYAAAG
KAF8554845.1        ---------MNLKSLAALTLAASAAPLAAAGPLA--YALCQTG----CNGLAVACYTAAG
EGO00833.1          ---------MNLKSTAALLVVAASAPAALGGPLA--YAMCQTG----CNGLAVACYAAAG
XP_007316625.1      ---------MNLKSTAALLVVAASAPAALGGPLA--YAMCQTD----CNRLAVECYAAAG
KAH7911314.1        ---------MNLKSTAALILVAASAPAVLGGPLA--YAACQTG----CNGLAVACYAAAG
KAH7920598.1        ---------MNLRSTIALFLAAASAPVAIGGPLA--YAACQTG----CNGLAVACYTAAG
XP_007316624.1      ---------MNFKSIAALVLVAATAPTVRGGPLA--YAACQTG----CNVLAVACYGAAG
KAG9312233.1        ---------MNFKSLTAITLAAAAVPLASAGPLA--YAACQTG----CNGLAVACYTGAG
KAH7883341.1        ---------MNLKSLAAIALAAAAAPIAMGGPIA--YGLCQTG----CNVVAVACYAGAG
XP_007768952.1      -------------------VAASTAPAAFGGPLA--YAACQTG----CNTLAVACYAAAG
XP_007769248.1      ---------MNLKLAGALLVAASAAPAAVGGPIA--YGICQTG----CNGLAVACYAGAG
KZP29597.1          ----------MRFTPVALLAIVAAT-PVLGGPIA--YALCQTG----CNGLAVACYAGAG
KZP06955.1          ----------MRFTPVALLAIVAVATPALGGPLA--YAACQTG----CNGLAVACYAGAG
KZP03955.1          ----------MRFTPVALLAIAAAATPVLGGPLA--YAMCQTG----CNGLAVACYSGAG
KZP33060.1          ----------MRITPVTILAVAAVATPALGGPLA--YALCQTG----CNGLAVACYAGAG
KZP05526.1          ----------MRLTHVTLLAIAGAATPAMGGPLA--YAACQTG----CNGLAVACWAAAG
KIM87109.1          ---------MRLTTPLTLLAAAAAATPVAGGPLA--YAACQTG----CNGLAVACYAAAG
KZP33058.1          ----------MRFTLIALLAVIAAATPALGGPLA--ALACQTG----CISLTATCYAAAG
KAF8500117.1        ----------MNLKSHALVLLAALIPAVNGGPVA--YALCQTG----CNAVAVACYGAAG
KAF9222454.1        ---------MNLKSLAALTLAVSATPAVMAGPFAYGLCQTG------CNVLVGACYAGAG
KAF8845957.1        ---------MNLKSLAALTVAASAAPLVMAGPLAYGLCQTG------CNVLVGACYAGAG
KIK73891.1          ---------MNFKALAALTVAASAAPLVTAGPLAYGLCQTG------CNVLVGACYAGAG
```

```
KIJ15625.1          ---------MNLKSLVALTVAASATPLVMAGPIAYGLCQTG------CNSLLGACYAGVG
KIJ15618.1          ---------MNLKSLAVLTAVASAAPLVMAGPLAYGLCQTG------CNALVGVCYAGAG
KIJ15617.1          ---------MNLKSLVVLTVVASAAPLVTA---YYVICQTG------CNVLACACYVGVG
KIJ15626.1          ---------MNLKSLVVLTVVASAAPLVTAGPLAYAICQTG------CNVLACACYGGAG
KIK77344.1          ---------MNLKRLVALTITVSAAPLVMAGPIAYALCQMG------CNVLAYACYAGVG
KAG8220444.1        ---------MNLKSLAALTLAASVP-LASAGPLAYGLCQTASPRRPGCNALVVSCYAGAG
KAF9222481.1        ---------MKLKFVTALAVAASIPPLTIAGPIAYAICQTG------CNTLAVACYAGAG
KAG9311090.1        ---------MNPKSLAALTLAAAAVPLVSAGPIAYAICQTG------CNSLAVVCYSAAG
KIM55694.1          ---------MNFKALAALTALAAAP-VVTAGPLAYALCQTG------CNGLAVACYTAAG
KAG2033209.1        ---------MNFKSLALFLTAAAVPQVVVAGPLA--YGICQTG----CNALVVACYAGAG
XP_041290278.1      ---------MNFKSLALLLTAAAVPQVAVAGPLA--YGICQTG----CNGLAVACYAGAG
XP_041290277.1      ---------MNLKSLALLLTAAAAPQVVVAGPLA--YAICQTG----CNGLAVACYAGAG
XP_041155838.1      ---------MNFKSLALLLTAAAAPQVAVAGPLA--YAICQTG----CNGLVVACYAGAG
KAG2048018.1        ---------MNFKSLALLLTAAAAPQVVVAGPLA--YGVCQTG----CNALVVACYAGAG
KAG1875880.1        ---------MNFKSLALLLTAAAAPQVVVAGPLA--YGICQTG----CNALVVACYAGAG
XP_041155836.1      ---------MNFKSLALLLTAAAVPQVAVAGPLA--YGVCQTG----CNALVVACYAGAG
KAG2033206.1        ---------MNFKFLAVLLTAAAVPQAVVAGPLA--YAICQTAG---CNGLAVACYAGAG
XP_041237613.1      ---------MNFKSLALLLTAAAVPQAVVAGPLA--YAICQTG----CNGLAVACYAGAG
KAG2087743.1        ---------MNFKSLALLLTAAAVPQAVVAGPLA--YGICQTG----CNGLAVACYAAAG
KAG2752971.1        ---------MNFKSLALLLTAAAVPQAVVAGPLA--YAICQTG----CNGLAVACYAGAG
KAG1762396.1        ---------MNFKSLALLLTAAAVPQAVVAGPLA--YAICQTG----CNGLAVACYAGAG
KAG1774592.1        ---------MNFKSLALLLTAAAVPQAVVAGPLA--YAICQTG----CNGLAVACYAGAG
KAG2338305.1        ---------MNFKSLALLLTAAAVPQAVVAGPLA--YAICQTG----CNGLAVACYAGAG
KAG1734527.1        ---------MNFKSIALLLTAAAVPRAVIAGPLA--YAICQTG----CNGLAVACYAGAG
KAG2087744.1        ---------MNFKSLALLLTAAAVPQAVVAGPLA--YGICQTG----CNGLAVACYAGAG
KAG2117666.1        ---------MNFKSLALLLTAAAVPQAVVAGPLA--YGICQTG----CNGLAVACYAGAG
XP_041237615.1      ---------MNFKSLALLLTAAVVPQAVVAGPLA--YGICQTG----CNGLAVACYAGAG
XP_041224654.1      ---------MNFKSLALLLTAAAVPQVVVAGPLA--YGICQTG----CNGLAVACYAGAG
XP_041160366.1      ---------MNFKSLALLLTAAAVPQVVVAGPLA--YGICQTG----CNGLAVACYAGAG
XP_041185215.1      ---------MNFKSLALFLTAAAVPQAVVAGPLA--YGICQTG----CNGLAVACYAGAG
KAG2117668.1        ---------MNFKSLALLLTAAVVPQTVVAGPLA--YAMCQTG----CNGVAVACYSAAG
KAG2087746.1        ---------MNFKSLALLLTAAAVPQTVVAGPLA--YAMCQTG----CNGVAVACYSAAG
XP_041237617.1      ---------MNFKSLALFLTAAAVPQVVVAGPLA--YGICQTG----CNGLAVACYAAAG
KAG1882508.1        ---------MNFKSFALFLTAAAVPQLVVAGPLA--YGICQTG----CNGLAVACYAAAR
XP_041185216.1      ---------MNFKSLALFLTAAAVPQAVVAGPLA--YGICQTG----CNGLAVACYAAAG
XP_041290280.1      ---------MNFKSLALLLTATAIPQVVVAGPLA--YAICQTG----CNGLAVACYAGAG
XP_041160372.1      ---------MNFKSLALLLTATAIPQVVVAGPLA--YAICQTG----CNGLAVACYAGAG
KAG2048022.1        ---------MNFKSLALLLTATAIPQVVVAGPLA--YAICQTG----CNGLAVACYAGAG
XP_041224652.1      ---------MNLRSLALLLTATAIPQVVVAGPLA--YAICQTG----CNGLAVACYAGAG
XP_041312474.1      ---------MNLKSLALLLAATAVPQVVVAGPLG--YAICQTG----CNGLAVACYAGAG
KAG1774591.1        ---------MNLKSIALLLTAAAVPQAVVAGPLA--YGICQTG----CNGLAVACYAGAG
KAG2338307.1        ---------MNLKSVALLLTAAAIPQAVVAGPLA--YGICQTG----CNGLAVACYAGAG
XP_041312481.1      ---------MNFKSLAALLTAAAVPQVVVAGPLS--YAICQTG----CNGVAVACYAGAG
XP_041224653.1      ---------MNLKSLTLLLTAAAVPQVVVAGPCL--YAICQAG----CIGLVVTCYAGAG
KAG2752972.1        ---------MNLKSITVLLTAAAVP--VFAGPLG--YALCQTG----CNGLAVACYAGAG
KIK36068.1          ---------MNLKSITLLLTAAAVP--VFAGPLG--YALCQTG----CNGLAVACYAGAG
XP_041202747.1      ---------MNFKSIAVLLTAAAAP--AYAGPLA--YGICQTG----CNGLAVACYAGAG
XP_041237616.1      ---------------------------------------CSTG----CNGLAVACYAGAG
KAG0702442.1        ---------MNFKSLAVLLTAAAAP--VAAGPLA--YGICQTG----CNGMAVACYAAAG
KAG0702443.1        ---------MNFKSLAVLLTAAAAP--VAAGPLA--YAICQTG----CNGMAVACYAAAG
XP_041171419.1      ---------MNLKSIVVLLTAVAAP--AVAGPLG--YAICQTG----CNSLAVACYGAAG
XP_041169262.1      ---------MNLKSIAVLLTAVAAH--AVTGPIE--YASCQTG----CNGHAVSCYGAAG
XP_041312473.1      ---------MNLKSLALLVTAAVVPHAVVAGPLASDYGICQTG----CNGVAVACYTRAG
KAG0701671.1        ----------MNFRFTTVLLLTAAAAPAVAGPLG--YAICQTG----CNGIAVACYAGAG
XP_041171032.1      ----------MNFKYITVLLAATAAP-AVAGPLG--YAICQTG----CNGIAVACYAGAG
KAG0695044.1        ---------MNSKSTTVTLLCTAAAAPAVAGPLG--YAICQMG----CNGISVACYSAAG
KIK35201.1          --------MKFKSITVAILLLATGP--AVAGPIG--YAICQTAG---CNGIAVACYSAAG
KAG1759280.1        --------MKFKSITVAILLLATGP--AVAGPIG--YAICQTG----CNGIAVACYSAAG
KAG2747675.1        --------MKFKSITVAILLLTTGP--TVAGPIG--YAICQTG----CNGIAVACYSAAG
XP_041291730.1      --------MKFKSISTILLLAATAG-PAVAGPIG--YAICQTG----CNGIAVACYSAAG
XP_041160953.1      --------MKFKSISTILLLTATAV-PAIAGPIG--YAICQTG----CNGIAVACYSAAG
KAG1855037.1        --------MKFKSIGIILVLLAATAGPAVAGPIG--HAICQTAG---CNGIAVACYSAAG
KAG1742009.1        --------MKYKSIAVFLLTATAGP--AVAGPIG--YAICQTG----CNGIAVACYSAAG
KAG1839191.1        --------MKFKSITTILILAATAG-PVVAGPIA--YAICQTG----CNGIAVACYSAAG
KAG1839194.1        --------MKFKSTTTILILAATAG-PVVAGPIA--YAICQTAG---CNGIAVACYSAAS
OJA16009.1          ---------MNIKSAISLITAAAAAPAVVAGPLG--YAICQTG----CNALVVSCYAGAG
OAX33574.1          ---------MNIKSTISLIIAAAAAPAVVAGPIG--YAICQTG----CNALVVSCYAGAG
OJA20466.1          ---------MNIKSTISLIIAAAAAPAVVAGPLG--YAICQTG----CNALVVSCYAGAG
KAJ8594645.1        ---------MNYKHTAILLVAAIASPAVVAGPLG--YAICQTG----CNALAVACYAGAG
KAG2353321.1        ---------MNFKSLALFLTAAAVPQAVVAGPLA--YTICQTG----CNCLAVACYSAAG
KAG2360710.1        ---------MNFKSLALFLTAAAVPQAVVAGPIA--YAICQTG----CNCLAVACYSAAG
KAG2062618.1        ---------MNFKSLALLLTAAAIPQAVVAGPIG--YAICQTG----CNCLAVACYTGAG
KAG2074957.1        ---------MNFRSLALFLAAAAVPQAVVAGPLA--YGICQTG----CNCLAVACYAGAG
KAG2353322.1        ---------MNFKSLALLLTVAAAPQAVMGGPLA--YGICQTG----CNCLAVACYAGAG
KAF8351847.1        ----------MRLSKLLLPIAVALSSTGIVNAGPIAYGLCQTV----CNFGAVACYAAAG
KAF8335236.1        ----------MRPSKLLLPIAVALSTTGIVTAGPIAYGVCQTA----CNAGAVTCYTGVG
KAG6331624.1        -------------MNLRCIAAYSLISLPLAMAGPLAYAACQTG----CNAIVVACYAGAG
KAF8835601.1        ---------MNLKSLATLTLAAP--LVIASPLAYYNYGACQTG----CNTRAGACYTDAG
KIJ13640.1          ---------MNLKSPAALALAAPLVQVIASPLAYYNYGACQTG----CNTGAGACYTAAG
KAH8106389.1        -------MRFSLRVFALAAAAASLPGALGEPFAPA---FCQSS----CYAGIAACYGAAG
KAH8106391.1        -----------MRLSPLALLAAALPGALGGPLAYG---LCQTG----CNAGVVACYGGAG
THH07108.1          ---------MRASLIALAAVSAAFPAVLGGPLAYG---VCQTG----CNVAAGTCYAAAG
```

```
KZT35398.1          ----------MRLPFIALVALVPAMTGSGVAAGPLAYAACQAG----CASLVMACYSAAG
KZS96226.1          ----------MRLPFIALVALVPAMTGSGVAAGPLAYAACQAG----CASLVMACYSAAG
QRV76181.1          --------MKLSVTSLVAFAAVTMNIQQ-VQAGPVAMGLCYSA----CNTGYVTCCTAAG
QRV90993.1          --------MKFSVTSLVALIAVTMNVQQ-AQAGPVAMGLCYSA----CNAGYVTCCTTAG
KAG9082513.1        --------MKFSFTSLVAVVAIALSAER-VQAGPVAMGLCYSA----CNAGYVTCCAAAG
KAG9125829.1        --------MKLSLTSIVAIMTIALSAER-AQAGPVAMGLCYSA----CNAGYVTCCATAG
XP_038910438.1      --------MRFSIASTFAFVAMALNVTH-VQAGPVAMGLCYSA----CNAGYVTCCTAAG
CEL57659.1          --------MKLSITSVVAFVTVALNAGQ-VQAGPIAMGLCYSA----CNAGYVTCCISAG
CAE6407853.1        --------MKFSITSVFAFAALALNAGQ-VQAGPVAMGLCYSA----CNAGYVTCCASAG
EUC53909.1          --------MKFSFTSVVAFAAVALNAGQ-VQAGPVAMGLCYSA----CNAGYVTCCTTAG
KAH7345108.1        --------MKFSFASVVAFAAVALNAGQ-VQAGPVAMGLCYSA----CNAGYVTCCASAG
CUA77418.1          --------MKFSFTSVVAFAAVALNAGQ-VQAGPIAMGLCYSA----CNAGYVTCCASAG
CAE6474563.1        --------MKFSFASLVAFVAVALNAGQ-VQAGPVAMGLCYTA----CNAGYVTCCASAG
CAE6407261.1        --------MKFSLASLAAFAALAFNAGQ-VQAGPIAMGLCYSA----CNAGYVTCCASAG
CAE6512876.1        --------MKFSFASVVAFAAVALNAGQ-VQAGPIAMGMCYSA----CNAGYVTCCTTAG
CAE6512885.1        --------MKFSLASVFAFVAVALNASQ-VQAGPIAMGMCYSA----CNAGYVTCCTAAG
CAE6449415.1        --------MKFSIAPIATLATLALNAGQ-VQAGPIAMGMCYSA----CNARYVTCCASAG
CUA77419.1          --------MKFSIAPMVTLATLALNAGQ-VQAGPIAMGMCYSA----CNAGYVTCCASAG
CAE6474554.1        --------MKFSFAPIVALATLALNAGQ-VQAGPIAMGMCYSA----CNAGYVTCCASAG
CAE7122953.1        --------MKFSVAPIVALATVALNAGQ-VQAGPIAMGLCYSA----CNAGYVTCCTSAG
CAE6506335.1        --------MKFSIAPVVALATLALSAGQ-VQAGPIAMGMCYSA----CNAGYVTCCTAAG
KAH7345109.1        --------MKFSVAPIVALATLALNVGQ-VQAGPIAMGLCYSA----CNAGYVTCCAAAG
EUC53908.1          --------MKFSIAPLVAFATLALNAGQ-VQAGPIAMGMCYSA----CNAGYVTCCVSAG
KAF8707836.1        --------MKFSIASAAALAALSLNIGQ-VEAGPIAMGLCYTA----CNAGYVTCCTVAG
CEL57606.1          --------MKLSVASAVAFAVLALNADQ-VQAGPIAMGLCYTA----CNAGYVTCCVTAG
KAF8604943.1        --------MKFSFTSIVAVVAVALNASQPVQAGPIAMGLCYSA----CNAGYVACCASAG
KAF8604944.1        --------MKLSIRSLVAAVVVSLSAPQPALAGPIAMGLCYSA----CNAGYTTCCTVAG
KAG9075620.1        ---------MKSSLTQLSVIAFALATGRSVQAGPIAMGLCYSA----CNAGYVACCAGAG
CAE6520723.1        --------MKLTITSTLAFVVLTLNARH-VHAGPVAMGACYTA----CNVGYVTCCAGAG
KDN50300.1          --------MKLTITSTLAFVVLTLNARH-VHAGPVAMGACYTA----CNAGYVTCCAGAG
KAH7345111.1        --------MKLTITSAFALVVLALNTRH-VHAGPATMGACYTA----CNVGYVTCCATAG
CAE6449405.1        --------MKLTVSSVFAFVLLALNARH-AHAGPISMGAYYTA----CNVGYATCCATAG
KAJ1311556.1        --------MKLSIASALACVIPTLNARH-VHAGPIFMAACYSA----CNAGYVTCCAAAG
CAE6470859.1        --------MKLTLAPALALVAFTLNARH-VHAGPVAMGACYTA----CNAGYVACCATAG
KAG8697657.1        --------MKLSIAFFSVITAVALHTGN-VRAGPMALALCTAT----CQAGYTTCCAAAG
XP_028477985.1      -------------MKAAIPLALAILASP-VAAGPVAWGLCYTA----CNASYGVCLGALG


KAF8735275.1        -SS-GGTFGTTGAPPAS---PAIAACKTALALCALLFPPGP----------
KAI1785556.1        FSL-GRAQGVSLGPPPS---MLGAICNTAIGSCLAACTAAICQ--------
KAF8661385.1        -----ATFGTVAAPLAP---PAILTCNAALGSCMALCAPLLIAPIP-----
XP_007380194.1      -----FTFGTVAAPLAP---PAILGCNAALGTCSAGCAAFLVAPIP-----
KAH6901377.1        -----TTFATIAALLAP---ATIFVCNSALATCSASCTAAFFYPWSM----
KAH6902106.1        -----FTFGTVAAADAP---PAILACNSALASCSAVCADNHPSGNGKARMG
KAJ3487682.1        -----FTFGTVAAPLAP---PAILACNTALGSCSAACAAAILLPTP-----
KAF8980955.1        -----FTFGTIAAPLAP---VAIIGCNTALGTCSAACAVTALIAPIP----
KAF8977750.1        -----FTFGTIAAPAAP---AAIVGCNTALGSCSAMCAGIALLAPTP----
KAF8980957.1        -----FIFGIYVPPLAP---AAITACNTALATCSAACYMSWFAPTP-----
KLO07918.1          -----ATFGTIAAPAAP---AAVVGCNGALGTCMAACSVAVIAPTP-----
KLO14569.1          -----ATFGTIAAPVAP---AAILGCNSALGTCMAACSVAVFAPTP-----
XP_038910436.1      -----FTFGTVAAAGAP---AVIVACNSALGTCSAGCAALLVTPTP-----
XP_038921664.1      -----FTFGTVAAAGAP---AVIVACNSALGTCSAGCAALLVTPTP-----
XP_038922423.1      -----FTFGTVAAVGAP---SAIVACNSALGTCSAGCAALLIAPAP-----
KAG8993678.1        -----FTLGTVAAPTAP---AAIVACNSALRTCSATCAFLLLAPIP-----
KAG8993679.1        -----FTFGTVAAPVAP---AAIVACNGALGTCSATCASLVLLAPIP----
RXW13811.1          -----FTFGTVAAPLAP---PAIVACNTALGTCSAACATVALLAPIP----
RXW22619.1          -----FTFGTIAAPLAP---PAIVACNAGLGTCSTACATVALLAPTP----
KAF8651157.1        -----FTFGTVAAPLAP---PAIVACNSALGTCSAACAVVALPAPVP----
PPQ74617.1          -----FTFGTIAAPLAP---PAILGCNAALGTCSATCATVALLAPTP----
PPQ74615.1          -----VTFGTVAAALAP---PAIVACNAALGTCSATCATVALLAPTP----
KAF9042500.1        -----FTFGTIAAPLAP---PAIVACNGALGTCSATCAAITLLSPI-----
KAF9042504.1        -----FTFGTVAAPLAP---PAIVACNGALGTCSATCAAITLLSPI-----
KAF9042497.1        -----FTFGAVAAPLAL---PAIVPCNIALGTCSATCAAITFFSPI-----
KAF9042501.1        -----FTFGTVVAPLAP---PAIVACNNALGTCSAACAAITLGSPI-----
KAK0445166.1        -----FTFGTVAAPAAP---AAIIGCNSALGTCSAACASVALLAPTP----
KAK0232397.1        -----FTFGTVAAPAAP---AAIIGCNSALGTCSAACASVALLAPTP----
KAK0192295.1        -----FTFGTVAAPAAP---AAIIGCNSALGTCSAACASVALLAPTP----
KAK0435962.1        -----FTFGTVAAPAAP---AAIIGCNSGLGTCSAACASVALLAPTP----
PBK70936.1          -----FTFGTVAAPAAP---AAIIGCNSGLGTCSAACATVALLAPTP----
SJL10225.1          -----FTFGTVAAPAAP---AAIVGCNSGLGTCSAACATVALLAPTP----
KAK0496260.1        -----FTFGTVAAPAAP---AAIVACNSGLGTCSAACASVALLAPTP----
XP_060325687.1      -----FTFGTVAAPAAP---AAIVACNSGLGTCSAACASVALLAPTP----
KAK0211506.1        -----FTFGTVAAPAAP---AAILGCNSALGTCSAVCASVALLAPTP----
KAK0204044.1        -----FTFGTVAAPAAP---PAIVGCNTALGTCSAACASVVLLAPIP----
KAK0232409.1        -----FTFGTVAAPVAP---VAIIGCNSALGTCSAMCAGVALLAPTP----
KAK0480870.1        -----FTFGTIAAPVAP---VAIIGCNTALGTCSAACATVALFAPTP----
PBK94666.1          -----FTFGTVAAPAAP---AAIIGCNTALGTCSAACASVALLAPTP----
XP_043040322.1      -----FTFGTIAAPAAP---AAIVSCNSALGACSASCASVALFSPTP----
KAK0477555.1        -----FTFGTIAAPAAP---AAIVGCNAALGSCSAICASVALLAPTP----
KAK0477562.1        -----FTFGTVAAPAAP---AAIVGCNAALGSCSTMCATVALLAPTP----
KAH6902114.1        -----FTFGTVAAPAAP---AAIVACNSALGTCSAACAAVALCAPTP----
KAF8219361.1        -----FTFGTVAAPAAP---AAILACNSSLGVCSAACSTVALCAPTP----
```

```
KAH6901375.1        -----FTFGTIAAPVAP---AAIVACNGALGTCSAACATVGLFAPTP----
KAH6901376.1        -----ATFGTIAAPAAP---AAIVACNGALGTCSAACATVGLFAPTP----
KAH6902116.1        -----ATFGTVAAPAAP---AAIVACNTALGTCSAACATVALLAPTP----
KAH6902112.1        -----ATFGTIAAPAAP---AAILGCNSALGSCSAACATVALFAPTP----
TFK18913.1          -----CTFGTVAAPAAP---LAIIGCNSALGTCSTACATVALFAPTP----
KAI0344501.1        -----ATFGTVAAPAAP---AAIVACNGALGTCSSMCATVGLFAPTP----
KAI0344500.1        -----ATFGTIAAPAAP---AAILGCNAALGSCSATCATIGLFAPTP----
KAF9256214.1        -----ATFGTVAAPAAP---AAILGCNSALGSCSAACAATALIAPIP----
XP_043004193.1      -----FTFATVAAPAAP---AALIGCNSALGTCSAACAATALIAPIP----
KAF9014159.1        -----FTFGTVAAAAAP---AALVGCNSALGTCSAACASTTLIAPIP----
TFK72753.1          -----ATFGTIAAPAAP---AAIIACNSALGTCSAACAATALIAPTP----
KAH9948988.1        -----VQFGTIAAPLAP---ATVLGCNTALGTCSAACATVCLLAPTP----
PPR07183.1          -----FTFGTIAAPVAP---AAVVACNAALGTCSAACATVALLAPTP----
KAF8898031.1        -----LTFGTIAAPAAP---AAILGCNAALGTCSATCATVALLAPIP----
KAI0785557.1        ----FTFGTVVASALAP---PAILACNAALGTCSAACASVALLAPTP----
KAF7793897.1        ----FTFGTVVAAALAP---PAILACNAALGTCSAACATVALLAPTP----
KAI0071739.1        ----FTFGTVVAAAAAP---AAVLACNAALGTCSAACATVALFAPTP----
KAF9554637.1        ----FTFG-TVAAADAP---AAVLACNSALGTCSAKCASVTLLAPTS----
KAF4620962.1        ----FTFG-TVAAADAP---AAVLACNSALGTCSEKCASVTLLAPTTS---
KAF9554638.1        ----FTFGTVVAAAAAP---AAVLACNTALGTCSAMCASVALLAPIP----
KAF4621497.1        ----FTFGTVVAAAAAP---AAILACNSALGTCSAACATVALFAPTP----
KJA16367.1          ----ATFGTVVAAAAAP---AAILACNSALGTCSAMCASVALLAPTP----
TDL19155.1          ----FQFGTVVAAAAAP---ATILACNAALGTCSAMCATVALLAPTP----
KAF5322937.1        ----LTFGTIVAAPAAP---AAALACNAGLGTCSAACATVALFAPTP----
KAF5322939.1        ----LTFGTIVAAPAAP---AVALACNAALGTCSAACATVALFAPTP----
KAF5322938.1        ----LTFGTVVAAAAAP---AAALACNAALGTCSTACATVALSAPTP----
KIM37591.1          ----ATFGTVVAAIAAP---PALLACNAALGTCSAACATVALLAPTP----
THG95442.1          ----LTFGTIVAAPLAP---AAALACNVALGTCSAACATVALFAPTP----
KAJ7080818.1        ----LVFGTVVAAPAAP---AAAIACNVALGTCSAACATVALFAPTP----
GAT42696.1          ----LTFGTVVAAAAAP---AAALACNTALGTCSAA---------------
KAF7289020.1        ----LTFGTVVAAAAAP---AAALACNAALGTCSAACATVALLAPTP----
XP_047747560.1      ----LTFGTVVAAPAAP---AAAIAYNAALGTCSAACATVALLAPTP----
KAJ7669782.1        ----LVFGTVVASPAAP---AAALACNVALGQCSAMCATVALLAPTP----
KAJ6517837.1        ----LVFGTVVASPAAP---AAALACNAALGQCSTMCATVALLAPTP----
KAJ7830972.1        ----LVFGTVVASPAAP---AAALVCNAALGKCSAICATVGLFAPTP----
KAJ6579297.1        ----LVFGTVAASPAAP---AAALACNAALGQCSAICATVGLFAPTP----
KAJ7604390.1        ----LVFGTVVAAPAAP---AAALACNAALGTCSATCAAVVLLAPIP----
KAJ6621514.1        ----LVFGTVVAAPAAP---AAALACNVALGTCSATCATVALLAPTP----
KAJ6621484.1        ----LVFGTVVAAPAAP---AAALACNVALGTCSATCATVALFAPTP----
KAJ7036623.1        ----LVFGTVVAAPAAP---AAALACNVALGTCSATCATVALLAPTP----
KAJ7777091.1        ----LVFGTVVAAPAAP---AAALACNIALGTCSATCATVALLAPTP----
KAJ7187815.1        ----LVFGTVIAAPAAP---AAALACNVALGTCSATCATVALFAPTP----
KAJ7147695.1        ----LVFGTVVAAPAAP---AAAVACNVALGTCSATCATVALFAPTP----
KAJ7664006.1        ----LVFGTVVAAPAAP---AAALACNAALGTCSGVCATVALLAPTP----
KAJ6462394.1        ----LVFGTVVAAPAAP---AAALACNAALGTCSGVCATVGLLAPTP----
KAJ6580769.1        ----LVFGTVVAAPAAP---AAALACNAALGTCSATCATVALLAPTP----
KAJ7114020.1        ----LVFGTVVAAPAAP---AAALACNAGLGTCSATCATVALFAPTP----
KAJ7498176.1        ----LVFGTVIAAPAAP---AAALACNAGLGTCSATCATVALLAPTP----
KAJ7712755.1        ----LVFGTVVAAPAAP---AAALACNAGLGTCSATCATVALFAPTP----
KAJ7724508.1        ----LVFGTVIAAPAAP---VAALACNAALGTCSATCATVALFAPTP----
KAJ7330443.1        ----LVFGTVVAAPAAP---AAAIACNVALGTCSATYATVALFAPTP----
KAJ7743748.1        ----LVFGTVVAAPAAP---AAAIACNVALGTCSATCATVALFAPTP----
KAJ7208767.1        ----LIFGTIIAAPLAP---PAAIACNVALGTCSATCATVALFAPTP----
KAJ7483446.1        ----LVFGTVVAAPAAP---AAALACNVALGTCSATCATVALFAPTP----
KAF7345529.1        ----LVFGTVVAAPAAP---AAAITCNVALGTCSATCATVALLAPTP----
KAF8211017.1        ----LVFGTVVAAPAAP---AAALACNAALGTCSATCATVALLAPTP----
KAJ7780066.1        ----LIFGTVVAAPAAP---VAALACNAALGTCSATCATVALFAPTP----
KAJ7691962.1        ----LVFGTVVAAPAAP---AAALACNLALGTCSATCATIGLFAPTP----
KAJ7359841.1        ----LTFGTVVAAPAAP---AAALACNAALGTCCATCATVALFAPTP----
KAJ7772818.1        ----LTFGTVVAAPAAP---AAALACNAALGTCCATCATVALFAPTP----
KAJ7264656.1        ----LTFGTVVAAAAAP---AAALACNAALGTCCATCATVALFAPTP----
KAF7371167.1        ----LTFGTVVAAAAAP---AAAIACNVALGTCSATCATVALLAPTP----
KAJ6471872.1        ----LTFGTVIAAAAAP---AAAIACNIALGTCSATCATVALLAPTP----
KAF7371160.1        ----LTFGTVVAAPAAP---AAALACNAALGTCSATCATVGLFAPTP----
KAJ7330535.1        ----LTFGMVVAVAEAP---AAAIPCNSALGICSANCAAVALVAPTP----
KZV67638.1          ----AVFGTVVAAPAAP---AAILACNAALGTCSAACATVALFAPTP----
VDB91597.1          ----ATFGTVIAAAAAP---AAIIACNSALGTCSAACATVTLLAPTP----
KAI0035134.1        ----LTFGTVVAAPAAP---AAALACNAALGTCSAACATVALFAPTP----
XP_037222833.1      ----FTFGTVVAAAAAP---AAILACNSALGTCSTACATVALLAPTP----
KAF5390836.1        ----FTFGTVVAAAAAP---PMILACNAALGTCSAACATVALFAPTP----
THU93403.1          ----FTFGTVVAAAAAP---PVILACNAALGTCSTACATVALFAPTP----
KAH8107856.1        ----FTFGTVVAAAAAP---AVIIGCNSALGTCSAACATVALFAPTP----
KAI0089603.1        ----FTFGTVVAAPATP---AVIIACNAALGTCSAACATVALFAPTP----
KIY47514.1          ----FTFGTVVASAATP---AVILGCNSALGTCSAACASVALLAPTP----
KXN83432.1          ----FTFG-TIAAPVAP---PAILACNAALGTCSAACAAVVLTPTL-----
KXN83429.1          ----FTFG-TIAAPIAP---PAILACNAALGTCSAACAAVVLTPTL-----
KXN83427.1          ----FTFG-TIVAPVAP---PAILAYNTALGTCSTARSAVASTSTP-----
KXN93170.1          ----FTSG-TIAAPVAV---PAISTRNAAFGICSAACAAVASTPTP-----
KXN81170.1          ----VTFG-TILAVAAP---PAILACNAALGTCSAACAAVALTPTP-----
KXN91089.1          ----FTFG-TVAAAVAP---PAILACNAGLGTCSAACAAVALTPTP-----
KXN83428.1          ----FTFG-TVLAVAAP---PAILACNAALGTCSAACAVVALTPTP-----
KAF5347592.1        ----FTFG-TIAAAAAP---PAIVACNSALGTCSAACATIALTPTP-----
```

```
TFK33514.1          ----FTFGTVVAAAAAP---PAILACNAGLGTCSSACAVVALTPTP-----
KAH9894858.1        ----YTFGTVTAGLGTP---AVIVGCNAALGKCSAACAIVALAPTP-----
KAI0326571.1        ----FTFGTVTAGLGTP---AVILGCNAALGKCSAACAIVALTPTP-----
KAI0656638.1        ----YTFGTVTAGLGTP---AVILGCNAALGKCSAACAIVALTPTP-----
KAJ8495237.1        ----YTFGTVTAGLGTP---AVILGCNAALGKCSAACAIVALTPTP-----
OSC99408.1          ----YTFGTVTAGLGTP---AVVLGCNAALGKCSAACAVVALTPIP-----
KAI9059205.1        ----YTFGTVTAGLGTP---AVILGCNSALGQCSAACAVVALSPIP-----
KAI8993820.1        ----FTFGTVTAGAATP---AVILGCNAALGKCSAACAVVALTPTP-----
CDO70453.1          ----FTFGTVTAGAGVP---AVVLGCNAALGTCMAACAAVALAPIP-----
KAI0373461.1        ----AVFGTVTAGVGTP---AVILGCNAALGKCSAACAIVALTPTP-----
KAI0360878.1        ----ATFGTITAGAGTP---AVILGCNAALGKCSATCAALTLLAPIP----
KAH9851372.1        ----ATFGTVTAGAATP---AIILGCNAALGKCSASCALVTLLPTP-----
KAI0633791.1        ----YTFGTVVAGPAAP---AVIMGCNAALGKCSAACAVTLLAPV------
KAI0737527.1        ----AVFGTVTAGVGTP---AAILACNAALGQCSAACAVVALTPTP-----
KAI0744947.1        ----AVFGTVTAGVGTP---AAILGCNAALGQCSAACAVVALTPTP-----
KAH9851373.1        ----AVFGTVTAGVGTP---AAIVACNVALGQCSAACALIVLAPTP-----
KAI0824053.1        ----AVFGTVTAGVGTP---AAIIACNVALGQCSAACALVALTPTL-----
KAI0373460.1        ----AVFGTITAGVGTP---AAIIACNVALGQCSAACAVVAFTPTP-----
KAI0360877.1        ----AVFGTVTAGVGTP---AAILACNVALGQCSAACALVVLAPTP-----
KAI0768850.1        ----AVFGTVTAGVGTP---AAILGCNVALGQCSAACAVVAFTPTP-----
XP_008034073.1      ----FVFGTVTAGVGTP---AAVLACNVALGQCSAACAVVALTPTP-----
KAI0666621.1        ----AVFGTVTAGVGTP---AAILACNAALGTCATACVAAGFAPTL-----
KAI0644389.1        ----AVFGTVTAGVGTP---VAILACNAALGTCSAACIAAGFAPIP-----
OJT05103.1          ----FTFGTVTAGVGVP---AAIVGCNAGLGVCQAACAAAIFAPTL-----
XP_008033715.1      ----FTFGTVTAGAGVP---AAVVACNAGLGVCMAGCAAAIFAPTP-----
THV03464.1          ----FMFGTMIAVHNVP---PVVLACNTGLGTCSATCVVALLTPD------
XP_036630288.1      ----FTFGTVIATPEAP---AAVLACNAALGACSATCATGLVAPTS-----
KAJ7036624.1        ----LVFGTVIADADAP---VAALACNKALSECSSNCT-------------
KAJ7780067.1        ----LVFGTVVAEPAAP---AAALVCNKALSTCSSVCAKETLSAPTQ----
KAJ7498165.1        ----LVFGTVVATPDAP---PAALACNNNLSTCATNCSTTALLAPTL----
KAJ7832735.1        ----LVFGTVVADAAAL---PVALRCNAALGKCAADCALDTDTTKYV----
KAJ7811222.1        ----LVFGTVVADAAAP---PVALRCNAALGKCAADCALDTDTITSK----
XP_008038531.1      ----RKFGTVTTDENTP---TAILNCNAALGICQQACAKAV----------
XP_008042174.1      ----MVFGAVTAGIATP---VVALACNAVLDRCQAECAIQGADAV------
OCB91128.1          ----LTFGTVIAAPAAP---AAAIACNSVLGVCMAACAASFLAPIP-----
KAF9461010.1        ----FTFGTVIATPAVP---AVILACNASLGTCSAACVAVTLLAPTL----
KAF9255919.1        ----ATFGTVVASPAAP---VAILACNAALGKCSAACATTALIAPTP----
KAI0071738.1        ----AVFG-TVAAPAAP---PAILACNAALGTCSSACAATALIAPIP----
TFL02353.1          ----FTFGTVVAAAATP---AVLVACNTGLGTCSAACAATALIAPIP----
KAF5332937.1        ----ATFGTVVASAATP---AAILACNAALGKCSAACAAVTLVAPTP----
KDR81157.1          ----ATFGTVVAAAAAP---AAILACNSALGTCSAACAAVTLLAPTP----
KDR81156.1          ----ATFGTVVTDADTP---AAILACNAGLGACSAACPAVALPGPTS----
XP_007868697.1      ----FTFGTVVAAPAGP---AAVLACNAALGTCSAACATTALIAPIP----
KZT19035.1          ----FTFGTVVAAAAAP---AAILGCNSALGTCSAACAATALIAPIP----
KDQ57059.1          ----FTFGTVVAAAAAP---AAVLACNAALGTCSAACAATALLAPIP----
KAK1228354.1        ----FTFGTVVAAAAAP---AAILACNSALGTCSAACAATALIAPTP----
KAJ8072958.1        ----FTFGTVIAAPATP---AVILACNAALGTCSAACATVGLFAPTP----
ESK93305.1          ----FTFGTVVAAIAAP---PVILACNAALGTCSAACAATALIAPIP----
KIJ22769.1          ----FVFGTVVAAPLAP---PAIIACNSALGVCSAACAATALIAPIP----
KAF8176973.1        ----FTFGTVVAAPAAP---AAIMACNAALGSCSAMCASVALFAPTP----
KAF8176972.1        ----FTFGTVIAAPAAP---AAIMACNAALGSCSAMCASVALFAPTP----
KAF6743528.1        ----ATFGTVVASAAAP---AVILACNAALGSCSAGCASFALFAPIP----
KAF6748547.1        ----ATFGTVVASAAAP---AAILGCNAALGSCSAGCASFALLAPIP----
XP_040768338.1      ----FQFGTVVASPLVP---ATILACNAALGTCSATCATVVLLAPIP----
XP_040768339.1      ----FQFGTVVATPLAP---ATVLACNAALGTCSATCATVVLFAPIP----
XP_040768337.1      ----FQFGTVVASLLAP---ATILACNTALGTCSATCATVALFAPTP----
KAI0930540.1        ----FQFGTVVAGPLAP---ATILACNAALGTCSAACAGVTLLAPTP----
XP_024343832.1      ----FQFGTVVAAVAAP---ATILACNAALGTCSTMCATVALFAPTP----
EED79690.1          ----FQFGTVVAAAAAP---ATILACNAALGSCSAMCATVALFAPTP----
XP_024343044.1      ----FQFGTVIAAAAAP---ATILACNAALGTCSATCATVALFAPIP----
KIK03653.1          ----FTFGTVIAAAATP---AAILGCNAALGTCSATCATLVLFAPTP----
KIJ99165.1          ----FTFGTVIAAPAAP---AAILGCNAALGTCSATCATLVLLAPTP----
KAF9461003.1        ----FTFGTVIASAATP---AVIVGCNAALGTCSATCASLVLLAPIP----
KAF9528339.1        ----CVFGTVVAAAAAP---AAVLGCNAALGTCSATCATLVLFAPIP----
TFY71517.1          -----FQFGTILAVAAP---ATIIVCNSALGTCSAACAGITLLAPIP----
KAA1466928.1        -----FQFGTVLAAAAP---ASIVACNSALGTCSAACAGITLLAPIP----
KAA1466913.1        -----FQFGTVLAVAAP---ATILACNSALGTCSATCAGITLLAPIP----
KAJ7230373.1        ---VAFEFSVFDAAAAP---DVVLKCNEALGICSKTCAGMELFAPTP----
XP_009549554.1      ----FQFGTVVAAAATP---ATILACNAALGTCSATCATLVLFAPIP----
XP_007306302.1      ----FQFGTVVAAVAAP---ATILACNAALGTCSTACATVALFAPTP----
KDQ08146.1          ----FQFGTVVAAAATP---ATILACNAGLGTCSATCATVALLAPTP----
KAJ7586376.1        ----FTFGTVVAAPAAP---AVILACNAGLGTCSATCATVALFAPTP----
KII83275.1          ----FTFGTVIAAAATP---AALVACNAALGTCSATCATVALFAPTP----
KAF8153443.1        ----FTFGTVVAAPAVP---AVILGCNAGLGTCSAACATVALFAPTP----
RDB15947.1          ----FTFGTVVAAPATP---LVILGCNTGLGTCSAACATVALFAPTP----
KAF8808935.1        ----FTFGTVVASPAAP---AVLLACNAGLGVCSASCATVALFAPTP----
KAF8808937.1        ----YTFGTVVASPAAP---AVVQSCNAGLAACSTACATVALLAPTP----
KAF8869089.1        ----FTFSVIVATPAIP---PALLVCNAGLAACSAVCNTTFFASTP-----
KAF8869829.1        ----FTFGTVVAAPATP---AVLLACNAGLGTCSAACATVALFAPTP----
KAF8872941.1        ----FTFGTVVATVATP---AVIVGCNAGLGTCSAACATVALFAPTP----
KAF8872939.1        ----FTFGTVVAAAATP---AVIVACNAGLGTCSAACATVALFAPTP----
KAF8957624.1        ----FTFGTVIAAPAAP---AAVLACNAALGTCSATCATVALLAPTP----
```

```
KAF9074370.1        ----FTFGTVIAAAAAP---VAVLGCNAALGTCSATCATVALFAPTP----
KAJ3874754.1        ----FTFGTVIAAPATP---AVILGCNAALGSCSAMCATVALLAPTP----
KAJ3868585.1        ----FTFGTVIAAPATP---AVILGCNAALGTCSAMCATVALLAPTP----
KAJ3870732.1        ----FTFGTVVAAPATP---AVILGCNAALGTCSAMCATVALLAPTP----
KAJ4480927.1        ----FTFGTVVAAPATP---AVILGCNAALGTCSAACATVALLAPTP----
KAJ3742886.1        ----FTFGTVIAAPTTP---AVILGCNAALGTCSATCAAVALLAPIP----
KAJ3729090.1        ----FTFGTVTAAPTTP---VVLLGCNAALGTCSATCATVALLAPTP----
KAJ3998398.1        ----FTFGTVVAAPATP---AVILGCNAALGTCSATCAAVALLAPIP----
KAJ3793355.1        ----FTFGTVVAAPATP---AVILGCNAALATCSATCATVALLAPTP----
KAJ3727548.1        ----FTFGTVVAAPATP---AVILGCNAALGTCSATCATVALLAPTP----
KAF9472595.1        ----MTFGTVVAAAAAP---PLILGCNAALGTCSAMCATVALLAPTP----
KAJ7230375.1        ----FTFGTVVAAPAAP---VAVLGCNAALGTCAATCATVALLAPTP----
KIL66627.1          ----FTFGTVVAAPAAP---VAVLACNAALGTCSAACATIGLFAPTP----
THH27703.1          ----FTFGTVIAAPAAP---AAILACNAGLGTCSAACATIGLFAPTP----
KAI0705164.1        ----FVFG--VALPAAP---PAIMACNAALGTCSAACATVALFAPTP----
KAI4520142.1        ----FTMG--VALPAAP---PAILACNAALGTCSAACATIGLFAPTP----
TRM64791.1          ----FTFG--VALPAAP---PVILACNAGLGTCSAACATVALLAPTP----
XP_036630286.1      ----FTFGTVIAAPAAP---AAVLACNAALGACSATCATIGLFAPTP----
KAF4597889.1        ----FTFGTVIAAPAAP---AAVLACNAALGACSATCATIGLFAPTP----
KAF9497993.1        ----FTFGTVIAAPAAP---AAILACNAALGACSATCATIGLFAPTP----
KAG9220390.1        ----FTFGTVIAAPATP---AVLLACNAALGVCSATCATVALFAPTP----
KAF4567255.1        ----FTFG-TVAAPAAP---AAILACNAALGTCSSACATVGLLAPTP----
KAK0445201.1        ----ATFGTVVAAAATP---AVILGCNVALGTCSATCATVGLFAPTP----
SJL12908.1          ----ATFGTVVAAAATP---AVILGCNVALGTCSATCATVGLFAPTP----
PBK94627.1          ----ATFGTVVAAAAAP---AAILACNAALGTCSTACATVGLFAPTP----
KAK0232361.1        ----ATFGTVVAAAAAP---AVILACNASLGTCSTACATVALFAPTP----
KAK0211537.1        ----ATFGTVVAAPAAP---AVILACNAALGTCSATCATVALFAPTP----
KAK0496299.1        ----VTFGTVVAAAATP---AVILTCNASLGVCSATCATVALLAPTP----
PBK70903.1          ----VTFGTVVAAAAAP---AVILGCNAALGTCSATCATVALLAPTP----
KAK0192253.1        ----ATFGTIVAAAAAP---VAILGCNAALGTCSATCATVALLAPTP----
KAK0480837.1        ----ATFGTVIAAAAAP---AAILGCNAALGTCSATCATVGLLAPTP----
PBK78949.1          ----FTFGTVIAAPAVP---AVILTCNAALGTCSAACATVALFAPTP----
KAK0204073.1        ----FTFGTVIAAPAAP---AVVIACNAALGTCSAACATVALFAPTP----
XP_060325653.1      ----FTFGTVIAAPAAP---AAVLACNAALGTCSAACATVALLAPTP----
XP_043040323.1      ----FTFGTVIAAPAAP---AAILACNAALGTCSTACATVALLAPTP----
KJA16363.1          ----FTFGTVIAAPATP---AVILACNAALGTCSTMCATVALLAPTP----
KJA16366.1          ----FTFGTVIAAPATP---AVILACNAALGTCSTMCATVALLAPTP----
KZT72154.1          ----FTFGTVIAAPAAP---AAIIACNAALGTCSAVCASVALFAPTP----
KZT72155.1          ----FTFGTVIAAPATP---VAIIACNAALRTCSAACAGT-----------
XP_047899200.1      ----VTFGTVIAAPATP---AVILGCNAALGTCSAACATIALFAPTP----
TFY54996.1          ----FTFGTVIAAPAVP---AVILGCNAALGTCAATCATVALFAPTP----
KAI0693946.1        ---AVFG-TVTAGVGVA---PAILACNAALGTCFTACATAALCAPTP----
RDX45665.1          ---AVFG-TVTAGVGVA---PAIIGCNAALGTCTAACAATALIAPTP----
RPD73715.1          ---AVFG-TVTAGVGVP---PAILACNAALGICSAACAAAVLAPTP-----
KAI0744948.1        ---AVFG-TVTAGVAVA---PAILACNAALGTCSAACAATALIAPTP----
KAI0779875.1        ---AVFG-TVTAGVAVA---PAIVACNAALGVCSAACAATALIAPTP----
KAI0737528.1        ---VVFG-TVTAGVGVP---PAILACNMALGVCSAACATVGLFAPTP----
KAI0656637.1        ---ATFG-TVTAGVGTP---AAIIACNLALGQCSAACATIALLAPTP----
KAH9894857.1        ---ATFG-TITAGVGTP---AAIIACNVALGQCSAACATIALFAPTP----
OSC99407.1          ---AVMG-TVTAGVGTP---VAVLACNVALGQCSAACATVALFAPTP----
KAI9059204.1        ---AVMG-TVTAGVGTP---VAVLACNVALGQCSAACATVALFAPTP----
KAI8993819.1        ---AVFG-TVTAGIGTP---AAIIACNVALGQCSAACATIALFAPIP----
XP_007366826.1      ---FTFG-TVTAGLGVP---AAIVACNAALGTCSAACATVALFAPTP----
XP_007370845.1      ---FTFG-TVTAGAGVP---AAILACNAALGVCSSTCATVALFAPTP----
PIL23384.1          ---FTFG-TVTAGIGVP---AAILACNAALGTCSSACATVALFAPTP----
KAI1794409.1        ---FTFG-TVTAGVGVP---AVILGCNTALGICSSACATVALFAPTP----
KAI1791328.1        ---FTFG-TVTAGLGVP---AAIIACNAALGTCSAACATVALFAPTP----
PIL33906.1          ---FTFG-TVTAGLGVP---AAVLACNAALGTCSSACATVALFAPTP----
XP_047873742.1      ---FTFGTVKADDPHVP---AAVLNCNAALGTCQAACAKVTLPAATPH---
XP_047873750.1      ---FTFGSISTAGVDVP---AAITTCNATLGVCLAACAAVTFFPST-----
XP_047873746.1      ---FTFGTVVGAG--AP---QAVAACSAAQGKCSSACAAVTLFAPTP----
XP_047873743.1      ---FTFG-TVTAGLGIP---AVIVGCNTALGTCSAACAAVTLLAPTP----
XP_047873744.1      ---FTFG-TVTAGVGIP---AAIAACNSALGTCSAACAAVTLLAPTP----
XP_047873745.1      ---FAFG-TFTAGLGVP---PALVACQTTLEKCSSACASLFPSTP------
XP_047870886.1      ---FTFG-TVTAGLGVP---AAIVACNAALGTCSTACATIGLFAPTP----
XP_047873747.1      ---FTFG-TVTAGLGVP---AAIVACNAALGTCSAACATIGLFAPTP----
KAF9018149.1        ---FTFG-TITAGVGIP---AAIVACNAALGTCSAACASVTLLAPTP----
KAH8831031.1        ---VQFG-TVVAAAGAP---ATVIGCNVALGTCMTACAGTALIAPIP----
KJA27646.1          ---FTFG-TVVAGPETP---AVVLRCNAALGTCAHACATSALRAPTP----
KAI0323098.1        ----ATFGTVVAAAAAP---AAILGCNAALGTCSSMCAGVALLAPTP----
KAI0323099.1        ----ATFG-TVAAPAAP---AAILGCNAALGTCSTMCASVALFAPTP----
PFH45591.1          ----VVFG-TVLAATAP---AAILACNAAQGSCMALCAVTVLPLPTP----
KIM63914.1          ----CVFG-TVAAPTAP---AAILACNSAQGTCMAICAVVALAAPVP----
EJD44225.1          ----ATFGTVVATPLTP---AIILWCNAALGTCMATLCAPLLLIPLP----
KZV97636.1          ----AVFG-TVAAPTAP---AAILACNAAQGMCMATLCAPLLLIPFP----
KAH7096490.1        ----FTFG-TIAAPVAP---LAILGCNAALGTCMAAMCAPLLLIPFF----
KAH7096489.1        ----AVFG-TIAAPVAP---PAILACNAALGTCMGAICAPLLLIPFI----
KAH7090935.1        ----AVFG-TVLAVTAS---PTILACNGAQGLCMTTLFAPLLLTPVP----
KAI6004897.1        ----FTFG-TVAAPAAP---PMIVACNAGLGTCMAACAATALLAPIP----
KAI6017551.1        ----FTFG-TVAAPAAP---PMIVACNAGLGTCMAACAATALLAPIP----
KAI6017553.1        ----FTFG-TVIAGVAP---PMIVACNAGLGTCMAACAATALLAPIP----
KAI6037129.1        ----FTFG-TVIAGAAP---AMIVACNAGLGTCMTGCATTALLAPIP----
```

```
KAI6102071.1        ----VTFG-TVAAPAAP---PLVVACNAALGTCMAACAATALLAPIP----
KAI6111997.1        ----VAFG-TVAAPAAP---PMIVTCNAGLGTCMAACAATALLAPIP----
KAI6131194.1        ----VTFG-TVAAPAAP---LLIASCNAALGTCMAACATTALLAPTP----
XP_051595080.1      ----FTFG-TVAAAVAP---PMILACNAAQGTCMAACAATALWAPIP----
KAI6155686.1        ----FTFG-TVAAAAAP---PMILACNAAQGTCMAACAATALWAPIP----
KAI6098867.1        ----FTFG-TVAAAAAP---PLILACNAAQGTCMAACAATALLAPIP----
KAI6148428.1        ----FTFG-TVAAPAAP---PLILACNAAQGTCMAACAATALWAPIP----
KAI6009872.1        ----FTFG-AVAAPAAP---PLILACNAAQGACMATCAATALLAPIP----
XP_051595875.1      ----FTFG-TVAAPAAP---QMILACNAAQGACMAACAATALLAPIP----
KAI6147894.1        ----FTFG-SVAAAAAP---SLLVGCNTAQGACMAACAATALLAPVP----
KAI9568904.1        -----FTFGVTIV-AAP---PAIMACNAGLGTCMAACATAALFAPTP----
KAF8132063.1        -----CTFGVTIV-AAP---PAIMACNAGLGTCMAACATAALFAPTP----
KAF9237865.1        -----FTFGVTIV-AAP---PAIMACNAGLGTCMATCATVALLAPTP----
KIJ66076.1          -----FTFGVTIV-AVP---PAIMACNAGLGTCMAACATVALFAPTP----
KAF8415388.1        -----FIFGVTII-GVP---PAIMGCNAGLGTCMATCATVALFAPTP----
KAF8131977.1        -----FTFGVTII-GVP---PAIMGCNAGLGTCMAACAVVGLFAPTP----
KAG6382117.1        -----FTFGVTIV-AVP---PAIIGCNVGLGTCMAACAVVGLFAPTP----
KAF8554845.1        -----FTFGVTIV-AAP---PAIIGCNVGLGACMAACAGTALIAPTP----
EGO00833.1          -----FTFGVTIV-AAP---PAIMACNVGLGTCMATCATVGLFAPTP----
XP_007316625.1      -----FTFGVTIV-AAP---PAIMACNVGLGTCMATCATVGLFAPTP----
KAH7911314.1        -----FTFGVTIV-GAP---PAIMACNLGLGTCMATCATIGLFAPTP----
KAH7920598.1        -----FTFGVTIV-AAP---PAIMACNAGLGACMATCATIGLFAPTP----
XP_007316624.1      -----FTFGVTVV-AGP---PALIACNVGLGTCMATCATVALFAPTP----
KAG9312233.1        -----FVFGVTVV-GGP---PAILACNAALGTCMATCATVALFAPTP----
KAH7883341.1        -----FTFGVTIV-GAP---PAIMACNAALGTCMATCATVALFAPTP----
XP_007768952.1      -----FTFGTTIV-AGP---PAIVACNAALGTCMTACATVALFAPTP----
XP_007769248.1      -----FTFGVTII-GGP---PAVIACNVALGTCMAGCATVALFAPTP----
KZP29597.1          -----FTMGVAIV-AAP---PAIMACNAGLGGCMAICATVGLFAPTP----
KZP06955.1          -----FVMGVTIV-GAP---PAVMACNAGLGGCMAICATVGLFAPTP----
KZP03955.1          -----FIMGTTIV-GGP---PAIIACNLGLGTCMATCATVALFAPTP----
KZP33060.1          -----FTMGVAIV-AAP---PAIIACNLALGTCMATCATVALLAPTP----
KZP05526.1          -----FTFGVTIV-LVP---PAILACNVGLGTCMATCATVALFAPTL----
KIM87109.1          -----FTFGVTIV-AVP---PAIMGCNIGLGTCMATCATVGLFAPTP----
KZP33058.1          -----FIFAPTVI-GVP---PAIITCNVALGTCTAG-ATVVLFAPTP----
KAF8500117.1        -----FTFGVTIV-GAP---AAILGCNAALGTCMATCATVALLAPTP----
KAF9222454.1        -----FTFG-VTIVAAP---PAIIACNTGLGACMAACAATALIAPTP----
KAF8845957.1        -----FTFG-VTIVGAP---PAILACNTGLGACMAACAATALIAPTP----
KIK73891.1          -----FTFG-VTIVGAP---PAILACNAGLGTCMAACAATALLAPTP----
KIJ15625.1          -----FTVG-VTIVGAP---PAILACNAGLGTCMAACAVTCLFAPTP----
KIJ15618.1          -----FVFG-ATIVAAP---PAILACNGGLGTCMAACAVTALLAPTP----
KIJ15617.1          -----FTFG-VTIVAAP---PAILACNAGLGTCMAACAATALIGPIP----
KIJ15626.1          -----FVFG-VTIVGVP---PAILACNAGLGTCMAACAATALIAPIP----
KIK77344.1          -----FTFG-MTIIGAP---PTILACNASLGTCMAACAATAPIP-------
KAG8220444.1        -----FTFG-VTVVGAP---AAIIACNAGLGTCMAACAATALIAPIP----
KAF9222481.1        -----FTFG-VTVVGAP---AAIVACNFALGKCMTACALTALPAPIP----
KAG9311090.1        -----FTFGTVAAPAAP---AAILACNAGLGLCMTACAATALIAPIP----
KIM55694.1          -----FTFG-VALPAAP---PMILGCNVALGTCMGACAVTALIAPIP----
KAG2033209.1        -----FTFGVALP-AAP---PVLIACNVGLGTCMAACAAAVFAPTP-----
XP_041290278.1      -----FTFGVALP-LAP---PVLIACNVALGSCMVACATVVFAPTL-----
XP_041290277.1      -----FTFGVAVP-LAP---PALIACNVGLGGCMAACAAVALAPTL-----
XP_041155838.1      -----FTFGVALP-LAP---PALLACNVGLGTCMAACAVVALTPTL-----
KAG2048018.1        -----FTFGVALP-LAP---PAIIACNVGLGTCMAGCAAVALAPTL-----
KAG1875880.1        -----FTFGVALP-LAP---PAIIACNVGLGTCMAACAAVALAPTL-----
XP_041155836.1      -----FTFGVALP-LAP---PALIACNVGLGTCMAACAAVALAPTP-----
KAG2033206.1        -----FTFGVAVP-LAP---PALLACNAGLGGCMAACAVVALTPTP-----
XP_041237613.1      -----FTFGVAVP-LAP---PALLACNAGLGGCMAACAVVALTPTP-----
KAG2087743.1        -----FTFGVAVP-AAP---PVLLACNAGLGGCMAACAAVALTPTP-----
KAG2752971.1        -----FTFGVALP-AAP---PVLIACNAALGGCMAACAVVALTPTL-----
KAG1762396.1        -----FTFGVALP-AAP---PVLIACNVGLGGCMAACAAVALTPTP-----
KAG1774592.1        -----FTFGVALP-AAP---PVLIACNVALGGCMTACAAVALAPTL-----
KAG2338305.1        -----FTFGVALP-AAP---PVLVACNVALGGCMTACAAVALAPTL-----
KAG1734527.1        -----FTFGVALP-AAP---PVLIACNVALGGCMTACAAVALAPTL-----
KAG2087744.1        -----FTFGVALP-AAP---PVLIACNVALGGCMATCAAIALAPTL-----
KAG2117666.1        -----FTFGVALP-AAP---PVLIACNVALGGCMAGCAAVALAPTL-----
XP_041237615.1      -----FTFGVALP-AAP---PVLIACNVGLGGCMAACAAVALAPTL-----
XP_041224654.1      -----FTFGVALP-AVP---PALMACNVGLGGCMAACAAIALAPTLE----
XP_041160366.1      -----FTFGVALP-AVP---PVLMACNVGLGGCMAACAAIALAPTL-----
XP_041185215.1      -----FTFGVALP-LAP---PALIACNVALGGCMATCAVVALAPTL-----
KAG2117668.1        -----FTFGVALP-LAP---PVLIACNVALGGCMATCAAIALAPTL-----
KAG2087746.1        -----FTFGVALP-LAP---PVLIACNVALGGCMATCAAIALAPTL-----
XP_041237617.1      -----FTFGVALP-AAP---PVLIACNVGLGGCMAACAVVALTPTP-----
KAG1882508.1        -----FTFGVALP-AAP---PVLIACNVGLGGCMAACAVVALTPTP-----
XP_041185216.1      -----FTFGVALP-AAP---PALVACNVGLGGCMAACAAVALAPTP-----
XP_041290280.1      -----FTFGVALP-AAP---PVLLACNAALGGCMAACAVVALTPTP-----
XP_041160372.1      -----FTFGVALP-AAP---PVLLACNAALGGCMAACAVVALTPTL-----
KAG2048022.1        -----FTFGVALP-AAP---PALLACNAALGGCMSACAVVALTPTP-----
XP_041224652.1      -----FTFGVALP-AAP---PVLLACNAALGGCMAACAVVALTPTP-----
XP_041312474.1      -----FTFGVALP-AAP---PVLIACNTALGGCMAACAAVALAPTL-----
KAG1774591.1        -----FTFGVALP-AAP---PALIACNAALGGCMAACAVVALAPTL-----
KAG2338307.1        -----FTFGVALP-AAP---PALIACNSALGGCMAACAVVALAPTV-----
XP_041312481.1      -----FVFGVALP-LAP---PAVLACNTALGGCMAACAVVAFMPIP-----
XP_041224653.1      -----FTLVVAPP-LAG---PAVIACNVAFGGCTLACAALMFAPTP-----
```

```
KAG2752972.1          -----FTFGVALP-AAP---PVLLACNSALGGCMAACAVVALTPTL-----
KIK36068.1            -----FTFGVALP-AAP---PVLLACNSALGGCMAACAVVALTPTL-----
XP_041202747.1        -----FTFGVALP-VAP---AAVVGCNVALGGCMAACAVVALTPTL-----
XP_041237616.1        -----FTFGVALP-LAP---PAVIACNVALGGCMAACAVIALAPTL-----
KAG0702442.1          -----CTFGVALP-LAP---AAIVGCNASLGACMVACAAVALAPTP-----
KAG0702443.1          -----FTFGVAMP-LAP---AVIIGCNTSLGTCMVACAAVALAPTL-----
XP_041171419.1        -----FTMGVALP-AVP---AVLVSCNVGLGTCMAACAVVALSPTL-----
XP_041169262.1        -----FTFRVALP-SVP---PVLATCNTGLGTCMAACADLLSERVQL----
XP_041312473.1        -----FTFGVALP-TVP---PALAECNLALGRCMADCAAVTAPIL------
KAG0701671.1          -----FTFGVALP-AAP---PAIIACNAALGACMAACALVALGPTP-----
XP_041171032.1        -----FTFGVALP-AAP---PAIIACNAALGACMAACAVIALGPTP-----
KAG0695044.1          -----FTFGVALP-AAP---PAIVAYNVALGACMAACAAVALGPTP-----
KIK35201.1            -----FTFGVAPP-AAP---PAIIACNAALGACMAACAAVALGPTP-----
KAG1759280.1          -----FTFGVALP-AAP---PEIIACNAALGAYMAACAAVALGPTP-----
KAG2747675.1          -----FTFGVAPP-AAP---PAIIACNAALGACMAACAAVALGPTP-----
XP_041291730.1        -----FTFGVAPP-AAP---PAIIACNTALGACMAACAAVALGPTP-----
XP_041160953.1        -----FTFGVAPP-AAP---PAIIACNTALGACMAACAVVAFGPTP-----
KAG1855037.1          -----FTFGVAPP-AAP---PAIIACNTALGACMAACAAVAFGPTP-----
KAG1742009.1          -----FTFGVAAP-AAP---PAIIACNAALGACMAACALVALGPTP-----
KAG1839191.1          -----FTFGVALP-AAP---PAIMACNAALGACMAACAAIALGPTP-----
KAG1839194.1          -----FTFGVALP-AAP---PAIMACNAALGACVAACAAIALGPTP-----
OJA16009.1            -----FTFGVALP-AAP---PVIMACNAGLGTCMAACAVVALGQPI-----
OAX33574.1            -----FTFGVALP-AAP---PVIMACNAGLGTCMAACAVVALGQPL-----
OJA20466.1            -----FTFGVALP-VAP---AVIVACNAGLGTCMAACAVVAFSPTL-----
KAJ8594645.1          -----FTFGVALP-AAP---PVIIACNAGLGTCMAACAVVALGPTP-----
KAG2353321.1          -----FTFGVLIPGVTP---AAIMACNVGLGTCMAACAAIAFGPTP-----
KAG2360710.1          -----FTFGVPIPGVTP---AAIMACNAGLGTCMAACAAIAFGPTP-----
KAG2062618.1          -----FTFGVPIPGSTP---AAIMACNAGLGTCMAACAA------------
KAG2074957.1          -----FTFGVPIPGSTP---AAIMACNAGLGTCMAACAAIALTPTP-----
KAG2353322.1          -----FTFGVPIPGSTP---AAIMACNAGLGTCMAACAAIALTPTP-----
KAF8351847.1          -----FTFGTVAAPAAP---PIILACNAAQGACMTLCAPLLVAPTF-----
KAF8335236.1          -----FTFG-VTLVAAP---PAILACNSILGACMALCTPFLIAPTP-----
KAG6331624.1          -----FTFG-VALPVAP---PAILACNAALGTCMATCATIALAPTP-----
KAF8835601.1          -----FVFGAVTAAGAP---PAILACNDNLGTCMAACAVTARPVRVR----
KIJ13640.1            -----FVFGNVTTAEAP---PAILTCNASQGTCMAACARASSP--------
KAH8106389.1          -----LTFGTVPVAG-P---PAITVCNALFGTCMAGCEALG----------
KAH8106391.1          -----FTFGTVPVIGAP---AAILGCNALLGTCMAACAGLVLAPTP-----
THH07108.1            -----FTFGTVLVATAP---ASIMACNALLGSCMVACAGMAIAPTL-----
KZT35398.1            -----FVWGATLGVAAP---PTIIACNVGYGTCQAACAGAALVAPTP----
KZS96226.1            -----FVWGATLGVAAP---PAIIACNVGYGTCQAACAGAALVAPTP----
QRV76181.1            ----VTAGTFTLGLGVP---AALLGCSAVQGACMAACTPLLAAPSP-----
QRV90993.1            ----AVAGTFTLGLGVP---AALFACSAVQGACMAACTPLLAAPTP-----
KAG9082513.1          ----AVAGTFTLGLGVP---AALFVCSAVQGTCMAACTPLLAAPTP-----
KAG9125829.1          ----AIAGTFTLGLGVP---AALFTCSVVQGTCMAACTPLLAAPTP-----
XP_038910438.1        ----VTAGTFTLGLGAP---VALIACSLVQGACMSACTPLLAAPTP-----
CEL57659.1            ----AVAGTFTLGLGTP---VALAACSVVQGACMSACTPLLLAPTP-----
CAE6407853.1          ----AVAGTFTLGLGVP---VALAACSVVQGACMSACTPLLLAPTP-----
EUC53909.1            ----AVAGTFTLGLGVP---VALAVCSVVQGTCMAACTPLLAAPTP-----
KAH7345108.1          ----AVAGTFTLGLGIP---AALAVCSVVQGTCMAACTPLLAAPTP-----
CUA77418.1            ----AIAGTFTLGLGVP---AALAVCSVVQGTCMAACTPLLAAPTP-----
CAE6474563.1          ----AVAGTFTLGLGVP---AALAVCSVVQGTCMAACTPLLAAPTP-----
CAE6407261.1          ----ATAGTFTLGLGVP---AALAGCSVIQGACMSACTPLLAAPTP-----
CAE6512876.1          ----ITAGTFTLGLGIP---AAIAACSVVQGTCMAACTPLLVAPTP-----
CAE6512885.1          ----TTAGTFTLGLGVP---AAVAACSVVQGACMAACVPLGVAPIP-----
CAE6449415.1          ----TVAGTFTLGLGVP---AALAACSVVQGTCMVACTPLLAAPSP-----
CUA77419.1            ----TVAGTFTLGLGIP---AALAACSVVQGTCMAACTPLLAAPSP-----
CAE6474554.1          ----AVAGTFTAGLGIP---AALAACSVVQGTCMAACTPLLAAPTP-----
CAE7122953.1          ----AIAGTFTLGLGIP---AALAGCSVIQGTCMAACTPLLAAPSP-----
CAE6506335.1          ----VTAGTFTLGLGIP---AAVAACSVVQGTCMAACTPLLAAPTP-----
KAH7345109.1          ----VTAGTFTLGLGVP---AALAGCSIVQGACMAACTPLLAAPTP-----
EUC53908.1            ----TVAGTFTLGLGVP---AAIAACSAVQGTCMAACTPLLLAPTP-----
KAF8707836.1          ----VTAGTFTLGLGIP---AAVAACSVVQGACMAACTPLLVAPTP-----
CEL57606.1            ----AIAGTFTLGLGIP---AAVAACSVIQGACMAACTPLLVVPTP-----
KAF8604943.1          ----TTAGTFTLGLGVP---AAVAGCSGAQGACMAACTTLIVTPTP-----
KAF8604944.1          ----VTAGTFTFGLGVP---AAIAACSVIQGTCMATCTPLLTAPSP-----
KAG9075620.1          ----ATAGTFTLGLGAP---AALMACSVVQGTCMAACTPFLAAPSP-----
CAE6520723.1          ----ATVGLFTLGLGVP---AALAACSVVQGACMAACVPLGLAPTP-----
KDN50300.1            ----ATVGLFTLGLGVPAALAALAACSVVQGACMAACVPLGLAPTP-----
KAH7345111.1          ----VTAGVFTLGLGVP---AALGACSVVQGACMAACVPLGFAPTP-----
CAE6449405.1          ----TTAGIFTLGLGVP---AALAACSAVQGACMAACVPLGAAPTP-----
KAJ1311556.1          ----ATIGVFTLGLGVP---ATLAACSAVQGTCMAACVPLGVAPTP-----
CAE6470859.1          ----ITAGIFTLGAGVP---AALGACSAVQGVCMAACVPLGLAPTP-----
KAG8697657.1          ----TAIGIFTFGLGTP---VAVAGCSLARGACVAACAPLLAAQGP-----
XP_028477985.1        ----LVAGTFTLGAGTP---VAVVTCSVAQGACMSACSPILMAPTP-----
```

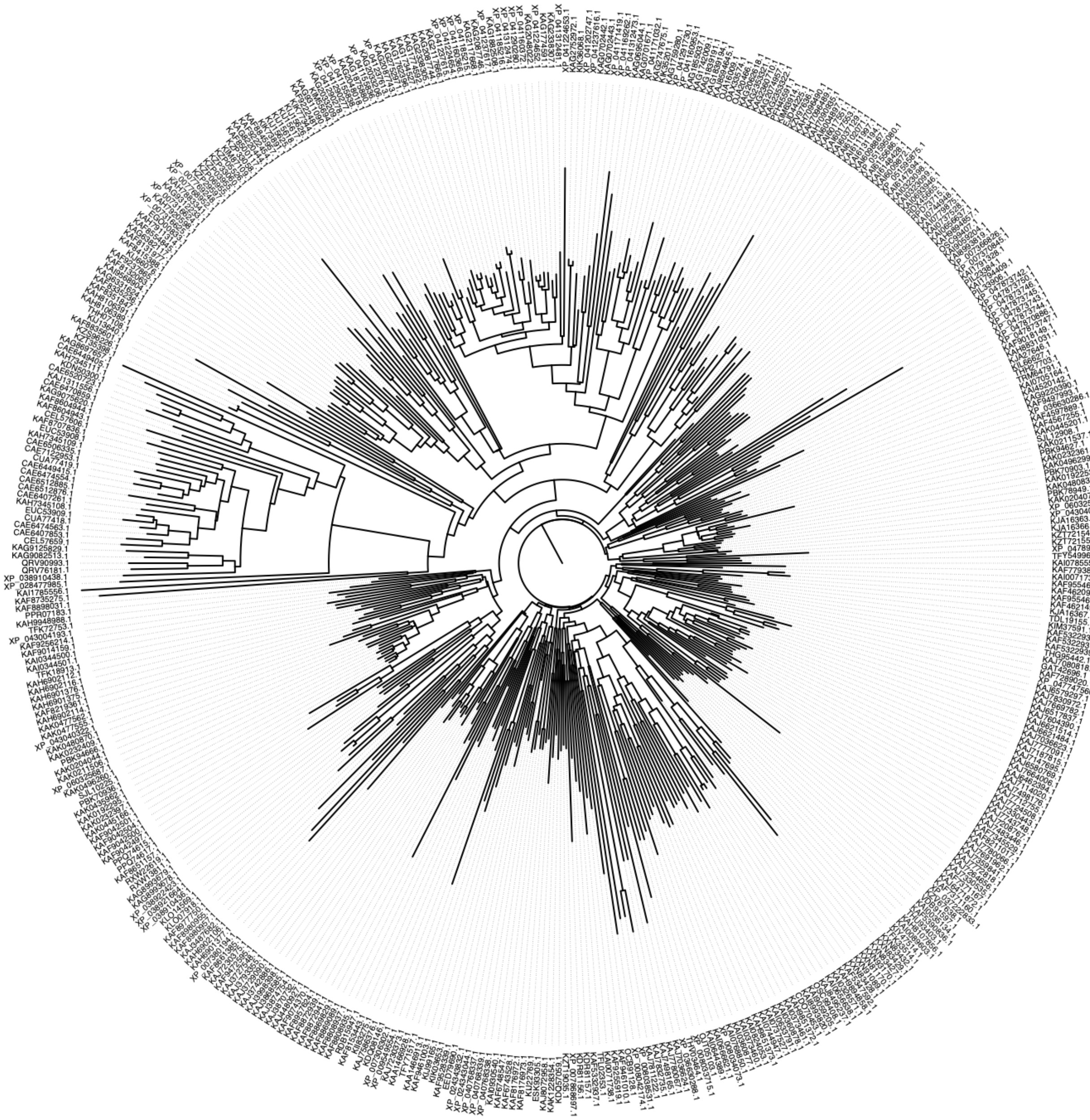

#### Supplementary File S2. Pezizomycotina HLPs: 339 sequences

>KAF8847622.1 hypothetical protein BDZ45DRAFT_315958 [Acephala macrosclerotiorum]
MRLSSILPVIAAVSTPAIAGPAAYGICQAGCASVVMACYGAAGFTWGATLGASAPASIVLCNSAFGTCYAACAAALLLPTP
>KAF2004078.1 hypothetical protein P154DRAFT_427667 [Amniculicola lignicola CBS 123094]
MRVIPLFTLFAEISVVTAGPAAYGICQAGCAAVVTACYAAGGATWGATLGATAGPTIVACNTAFGTCQAACWAALIALTP
>KAF1911237.1 hypothetical protein BDU57DRAFT_462016 [Ampelomyces quisqualis]
MKLTTIVNTVTALLLLPQCASAGPAAYGVCQAGCSAVVMACYGAAGFTWGATLGATAPATILACNAAYGTFQAACAAVLLAPTL
>KAG9233117.1 hypothetical protein BJ875DRAFT_379222 [Amylocarpus encephaloides]
MRPSSLLIPVVTFTTLASAGPAAYGICQAGCAAVVTACYAGAGFTWGATLGASAPATILACNAAFGTCQAACWAALIAPTP
>XP_033393871.1 uncharacterized protein K452DRAFT_235045 [Aplosporella prunicola CBS 121167]
MLRTTILTACVLTLAGTASAGPVGYGICQAGCAGVVMACYTAAGFTWGATLGASAPPTIIACNTAFGSCQAACAAILLAPTP
>RVD82005.1 hypothetical protein DFL_009849 [Arthrobotrys flagrans]
MKPSLIVSACTVISLVSAGPLAYGVCQAGCAAVVTACYGAAGFVWGATLATAAPPAIIACLTMSALGGRRRVR
>RPA84721.1 hypothetical protein BJ508DRAFT_412476 [Ascobolus immersus RN42]
MKLSAIFVFSALAAPALSGPVGYGVCQAGCSAVVMACYSAAGAVWGATAGLGAAPAVLACNAAYGTCQSACAAVLLAPTL
>TGZ81582.1 hypothetical protein EX30DRAFT_340458 [Ascodesmis nigricans]
MTPPYLLFLFTLLLLFTTTVTAGPATVGACQAGCAAVVVACYAAAGFVFGTIMPPSVPPAILACNAAQGTCYAGCYALFFMPTP
>XP_025501802.1 hypothetical protein BO66DRAFT_137353 [Aspergillus aculeatinus CBS 121060]
MKRLLPAICTGLLVCHVQAGPAAYGICQAGCASVVTACYAAAGFTWGATLGATAPASVLACNGAFGICQGACATALLAPTL
>XP_025509214.1 hypothetical protein BO66DRAFT_445109 [Aspergillus aculeatinus CBS 121060]
MMNISYPALGVLALAASCASAGPAACGVCQTGCAAVVMACYSAAGYTWGVALGAGVPATILACNSAFGTCQSACAAVILAPLF
>XP_031903430.1 uncharacterized protein BDW43DRAFT_240834 [Aspergillus alliaceus]
MKILYPAVGIFSTLVNSVYAGPEAYRICQAGYAAVVTACYSAAGFTWGATLTLPEARLA
>KAE8371456.1 hypothetical protein BDV26DRAFT_286779 [Aspergillus bertholletiae]
MRHIPTSIVILLMLAREGFTGPAAYGVCQAGCSAVVMACYSAAGFTWGATLGVSAPAFVIACNTAYGTCQAACAATLLTPTL
>OJJ72525.1 hypothetical protein ASPBRDRAFT_123752 [Aspergillus brasiliensis CBS 101740]
MKNLHLAVYTIPVIASYAYAGPAAYGICQAGCAAVVTACYSAAGYTWGATLGATAPASILACNSAFGTCCAACAATLLTPTL
>XP_025440039.1 hypothetical protein BO95DRAFT_394191 [Aspergillus brunneoviolaceus CBS 621.78]
MKGLLPAICTGLLVCHVQAGPAAYGICQAGCASVVTACYAAAGFTWGATLGATAPASVLACNGAFGICQGACATALLAPTL
>OOF99516.1 hypothetical protein ASPCADRAFT_203296 [Aspergillus carbonarius ITEM 5010]
MRILQAGVSLLIAANGEFRSPAAYGTCQAGCSSVVVSCYAAAGFVFGMVPAVSAPSAIVGCNSAYGNCQAACALVLLAPVL
>XP_025540887.1 hypothetical protein BO79DRAFT_227606 [Aspergillus costaricaensis CBS 115574]
MKNLYLAITTILILADYAYAGPLGYGICQAGCAAVVMACYSAAGYTWGATLGATAPASIVACNTAFGTCCAHCAATLLIPTL
>XP_025383483.1 uncharacterized protein BO83DRAFT_323600 [Aspergillus eucalypticola CBS 122712]
MKNLYLAITTIFIFASYAYAGPLGYGICQAGCAAVVMACYSAAGYTWGATLGATAPASIVACNAAFGTCCAHCAATLLAPTL
>XP_022405400.1 hypothetical protein ASPGLDRAFT_63535 [Aspergillus glaucus CBS 516.65]
MKIQTFFPLVLVLTSPVLAGPAAYGVCQSGCASVVMACYSAAGFTWGATMGASAPASIVACNTAYGTCQAACAATILGPTP
>PYI33230.1 hypothetical protein BP00DRAFT_306968, partial [Aspergillus indologenus CBS 114.80]
VLALASSYASASPAAYGVCQAGCAAVVMACYSGAGYTWGASLGATIPASILACNSAFGTCQSACAAVLLAPFP
>XP_025577596.1 hypothetical protein BO80DRAFT_423204 [Aspergillus ibericus CBS 121593]
LLTLLILTLTLPTPISASPTAYGICQAGCAAVVTACYSAAGFTWGATLGATAPPTILACNSAFGTCQAACAVTLLIPTL
>XP_025569355.1 hypothetical protein BO80DRAFT_369801 [Aspergillus ibericus CBS 121593]
MNLLGAGSGLLVAASNVFAGPSAYAVCQTGCSALVVSCYAAAGFVFGMVPAVSAPPAIIACNSSYGTCQAACASILLVPFP
>XP_025522674.1 zygote-specific protein, partial [Aspergillus japonicus CBS 114.51]
ALAAGYASATPAGYGVCQTGCATVVMACYSAAGFTWGAALGATIPASILACNSAFGTCQSACAAVLLIPFP
>OJZ90505.1 hypothetical protein ASPFODRAFT_40856 [Aspergillus luchuensis CBS 106.47]
MKKLYLLISTILILASYAYAGPMGYGICQAGCAAVVMACYSAAGYTWGATLGATAPASIVACNAAFGTCCAHCAATLLAPTL
>XP_025477608.1 hypothetical protein BO87DRAFT_268285, partial [Aspergillus neoniger CBS 115656]
AGPLGYGICQAGCAAVVMACYSAAGYTWGATLGATAPASIVACNAAFGTCCAHCAATLLMP
**>XP_025460783.1 uncharacterized protein BO96DRAFT_407345 [Aspergillus niger CBS 101883]**
**MKNIYLAVFTILIFVSYASAGPAAYGICQAGCAAVVMACYSAAGYTWGATLGATAPPTIVACNSAFGVCYSSCAATLLAPTL**
>XP_025520823.1 hypothetical protein BO85DRAFT_359315 [Aspergillus piperis CBS 112811]
MKKFYLLISTILILASYAYAGPVGYGICQAGCAAVVMACYSAAGYTWGATLGATAPASIVACNTAFGTCCAHCAATLLIPTL
>RDK39382.1 hypothetical protein M752DRAFT_220502 [Aspergillus phoenicis ATCC 13157]
MKNIYLAAYTIPISAGYASAGPTAYGICQAGCAAVVMACYSAAGYTWGATLGATAPPTIVACNSAFGVCYSSCAATLLAPTL
>XP_040642819.1 uncharacterized protein EURHEDRAFT_470710 [Aspergillus ruber CBS 135680]
MKLSTAILLPLASTAIAGPIGYGICQAGCSGVVMACYSAAGFTWGATLGASAPASIIACNTAYGTCQAACAAVLLAPTP
>XP_025468364.1 hypothetical protein BO94DRAFT_534229 [Aspergillus sclerotioniger CBS 115572]
MNLLRAVINLLIAGSVVVGGPAAYGVCQAGCSSVVVSCYAAAGFVFGMVPAVAAPPAIAGCNSAYGTCQAACATILLAPFP
>XP_024702685.1 zygote-specific protein [Aspergillus steynii IBT 23096]
MSLTNEVSAGPTGYGICQAGCSGVAMACYAAAGCTWGATLGATAPPAIVACNAAFGACQAKCALVLLAPTP
>KAE8155953.1 hypothetical protein BDV40DRAFT_282490 [Aspergillus tamarii]
MKILYPALLILLSSITQVNGGPAAYGICQAGCAAVVTACYTAAGFTWGATLGATAPASIVACNTAFGTCQAACATALLAPTP
>OJI81929.1 hypothetical protein ASPTUDRAFT_45267 [Aspergillus tubingensis CBS 134.48]
MKKLYLLISTILILADYAYAGPLGYGICQAGCAAVVMACYSAAGYTWGATLGATAPASIVACNTAFGTCCAHCAATLLAPTI
>XP_025488249.1 hypothetical protein BO82DRAFT_357645 [Aspergillus uvarum CBS 121591]
MKGLLPVICTSLLACYVQAGPAAYGICQAGCASVVTACYAAAGCTWGATLGATAPASVLACNGAFGICQGACATALLAPTL
>XP_025563775.1 hypothetical protein BO88DRAFT_404318 [Aspergillus vadensis CBS 113365]
MNKLYLAITTILVLASYAHAGPVGYGICQAGCAAVVMACYSAAGYTWGATLGATAPASIVACNTAFGTCCAHCAATLLIPTL
>PYI14498.1 zygote-specific protein, partial [Aspergillus violaceofuscus CBS 115571]
ALAAGYASATPAGYGVCQTGCATVVMACYSAAGFTWGAALGATIPASILACNSAFGACQSACAAVLLIPFP
>XP_040695488.1 uncharacterized protein ASPWEDRAFT_35427 [Aspergillus wentii DTO 134E9]
MKPSLLPLLLVASTASAGPAAYGVCQAGCSAVVMACYSAAGFTWGATAGASAPASILACNSAYGTCQAACAAALLAPTL
>KAF1987320.1 hypothetical protein K402DRAFT_57907 [Aulographum hederae CBS 113979]
MRPRAILIASLLLAAATSVSAGPASYGICQAGCASVVMACYAAAGANWGAVLAFGAPSAVATCNAAFGPCQSACHVVACIPFL
>KAF1972950.1 hypothetical protein BU23DRAFT_554800 [Bimuria novae-zelandiae CBS 107.79]
MRFHTIAATAMVALAGTTAAGPIGYGICQAGCSAVVVACYSAAGFTFGTVAAAAAPAAIAACNTAYGTCQAACAAVLLSPTL
>KAF1972136.1 zygote-specific protein, partial [Bimuria novae-zelandiae CBS 107.79]
VLAGPALAGPATYGVCQAGCPALVMACYSAAGFTWGTTLGASAPATIAACNTAFGTCQAACAAVLLAPMP
>KAF1970296.1 hypothetical protein BU23DRAFT_556961 [Bimuria novae-zelandiae CBS 107.79]

MRLSTLTATALLALTSTVAAGPIGYGVCQAGCSTVVMACYAAAGFTWGATLGATAPASIVACNAAYGTCQAACATVLLGPTP
>KAI1503117.1 hypothetical protein F5X99DRAFT_407449 [Biscogniauxia marginata]
*MKPTTAISVAIASALAPAVSA*GPAAYGVCQAGCSAVVMACYAAGGATWGATLGATAPATIIGCNTAFGSCQAACAAVLIVPTP
>KAI1485417.1 hypothetical protein F5X96DRAFT_674656 [Biscogniauxia mediterranea]
AATVLAALLAPAAVSAGPAAYGICQAGCSAVAMACYAAGGATWGATLGATAPATIVGCNTAFGACQAACWASLIAPTP
>KAI1630725.1 hypothetical protein F4809DRAFT_243437 [Biscogniauxia mediterranea]
MKLSSAIPAAAVLAVLAPAVSAGPAAYGICQAGCSAVAMACYAAGGATWGATLGATAPATIVGCNTAFGVCQAACWAALVAPTP
>KAI0600113.1 hypothetical protein F4775DRAFT_590835 [Biscogniauxia sp. FL1348]
*MKLSTAVPIAIIAALAPAANA*GPAAYGICQAGCSAVVTACYAAGGATWGATLGATAPATIVGCNSAFGACQAACWASLLAPTL
>KAJ5059812.1 hypothetical protein J3E74DRAFT_454481 [Bipolaris maydis]
MAISLLITLITLAFASLASAGPAAYGLCQAGCSAVVMACYSAAGFTWGATLGASAPASIIACNTAFGTCQAACASVLLAPTP
>XP_007694026.1 hypothetical protein COCMIDRAFT_111540 [Bipolaris oryzae ATCC 44560]
MPLLTAIITLAIASPALAGPAAYGVCQAGCSAVVMACYSAAGFTWGATLGASAPASIIACNTAFGTCQAACAAVLLTPML
>XP_007693411.1 hypothetical protein COCMIDRAFT_109719 [Bipolaris oryzae ATCC 44560]
MALSILIATIVFALASPISAGPAGYGLCQAGCSAIVMACYSAAGFTWGATLGASAPASIIACNTAFGTCQAACAAVLLTPTP
>XP_007687291.1 hypothetical protein COCMIDRAFT_36139 [Bipolaris oryzae ATCC 44560]
MKPLSLTKIAIAVLSLTAGQASAGPVGYAVCQAGCAGVVMACYSAAGFTWGATFGATAPASILLCNAAFGKCQAACAFVLLGPTP
**>XP_007693637.1 hypothetical protein COCMIDRAFT_110407 [Bipolaris oryzae ATCC 44560]**
**MALLLLITLITLAFASLASAGPAAYGLCQAGCSAVVMACYSAAGFTWGATLGASAPASIIACNTAFGTCQAACAAVVLTPTP**
>XP_014082052.1 hypothetical protein COCC4DRAFT_129741, partial [Bipolaris maydis ATCC 48331]
AGCAGVVMACYSAAGFTWGATFSASAPASILLCNAAFGKCQAACAVVLLGPTP
>XP_007700491.1 uncharacterized protein COCSADRAFT_42941, partial [Bipolaris sorokiniana ND90Pr]
SLASAGPAAYGLCQAGCSAVVMACYSAAGFTWGATLGASAPASIIACNTAFGTCQAACASVLLAPTP
>XP_014552310.1 hypothetical protein COCVIDRAFT_110563 [Bipolaris victoriae FI3]
MTSSLFIVIISLAFASPALAGPAAYGLCQAGCSAVVMACYSAAGFTWGATLGASAPASIIACNTAFGTCQAACAAVLLSPTP
>XP_014556701.1 hypothetical protein COCVIDRAFT_99125 [Bipolaris victoriae FI3]
*MKPFSLIKATTVVLSLTVRQASA*SRIEYAVCQAGCASLVMACYTAAGFVWGTVGRDTASQAILFCNAAFGKCSAACAEATL
>XP_007712361.1 hypothetical protein COCCADRAFT_96294 [Bipolaris zeicola 26-R-13]
*MKPFSLIKATTVVLSLTVRQASA*GRIEYAVCQAGCASLVMACYTAAGFVWGTVGRDTASQAILFCNAAFGKCSAACAEATL
>XP_007718397.1 hypothetical protein COCCADRAFT_41935 [Bipolaris zeicola 26-R-13]
MELSLFIAIITLALASPASAGPAAYGVCQAGCSAIVMACYSAAGFTWGATLGASAPASIIACNTAFGTCQAACAAVILTPTP
>XP_007718708.1 hypothetical protein COCCADRAFT_112853 [Bipolaris zeicola 26-R-13]
MASSFSIVVIALAFASPALAGPAAYGLCQAGCSAVVMACYSAAGFTWGATLGASAPASIIACNTAFGTCQAACAAVLLSPTP
>KAH8600237.1 hypothetical protein B0O99DRAFT_612058 [Bisporella sp. PMI_857]
MQISSSTITLLMAFTPPIAASLIGYGVCQAGCSTVVMACYGAAGFTWGATLGATAPATIIACNSAFGTCQAACAAVLLVPLP
>TEY32995.1 hypothetical protein BOTCAL_0701g00040 [Botryotinia calthae]
MKPTSILLIACLAGITTAGPVAYGVCQSGCAAVVMACYSAGGATWGATLGATAPATIVACNKAFGTCSATCAGLLVAPIP
>TGO59710.1 hypothetical protein BOTNAR_0157g00030 [Botryotinia narcissicola]
MNPSPSSTLLITVAILAGIATAGPVAYGVCQSGCAAVVMACYGAGGATLEKIYTDGDADVTQL
>KAF7952734.1 hypothetical protein EAE96_005964 [Botrytis aclada]
MKPTSTSILLIAFLAGITTAGPVAYGVCQSGCAAVVMACYGAGGATWGATLGATAPATIVACNTAFGICSAKCAGLLVAPIP
>XP_038729586.1 uncharacterized protein EAE97_009058 [Botrytis byssoidea]
MNPSPSSTFLITVAVLAGIATAGPVAYGVCQSGCAAVVMACYGAGGATWGATLGATAPATIVACNTAFGVCSAKCAGLLVAPIP
**>XP_024549469.1 hypothetical protein BCIN_06g06510 [Botrytis cinerea B05.10]**
**MKPTSPSVLLVAGLAGIATAGPIAYGICQSGCAAVVMACYSAGGATWGATLGATAPATIVACNTAFGTCSATCAGFLVAPIP**
>XP_038808418.1 uncharacterized protein EAE98_007617 [Botrytis deweyae]
MKPSSSSILLIAVFAGITTAGPIAYGVCQSGCAAVVMACYGAGGATWGATAAATAPATIVACNTAFGVCSAKCAVLLAAPIP
>XP_037194411.1 putative zygote-specific protein [Botrytis fragariae]
MKPSSSKSILLIAVLAGIATAGPIAYGVCQSGCAAVVMACYGAGGATWGATAGATAPATIVACNTAFGVCSAKCAGLLVAPIP
>XP_038754834.1 uncharacterized protein EAF02_009333 [Botrytis sinoallii]
MKPSSSSASSILLIAVFAGITTAGPVAYGVCQSGCAAVVMACYGAGGATWGATAAATAPATIVACNTAFGVCSAKCAVLLAAPIP
>KAK0100245.1 hypothetical protein ONS96_007528 [Cadophora gregata f. sp. sojae]
MKMRLSNPSAFLLFMATRPALIHAGPVGYGICKAGCASIVTACYAAAGATWGATLGATAPATVVACNTAFGICQGKCAVVALLPTP
>PVH85672.1 hypothetical protein DL98DRAFT_511255 [Cadophora sp. DSE1049]
MRFPTSAIILLATTPTLIIGGPVAYGICQAGCATVVTACYVAAGATWGATLGATAPATVVGCNTAFGICQAKCAIVALLPTP
>OCK90052.1 hypothetical protein K441DRAFT_666685 [Cenococcum geophilum 1.58]
MHSYALLLTALTLTHFTYAGPAAYGIRQAGCAAVVMACYSAAGFTWGATLGVSAPASILACNAAFGTCQGACAVAFIAPTP
>XP_023457676.1 hypothetical protein CB0940_03666 [Cercospora beticola]
MRISNLKTCLEAATLLITTAHAGPAAYGICQAGCAAVVVACYSAAGAVFGVAAPPAAVPAVLACNSAFGTCQAACWAALMAPTP
>KAF2207598.1 hypothetical protein CERZMDRAFT_102280 [Cercospora zeae-maydis SCOH1-5]
*MRVSNVTFKTLFLAAAYLHTTAHA*GPAAYGICQAGCSAVVAACYAAAGAVFGTVAAPAAPAAILGCNSAYGTCQAACWASLFAPTP
>RPA97680.1 hypothetical protein L873DRAFT_1809479 [Choiromyces venosus 120613-1]
MKLRNILLPASFTTMALAGPISYGICQSGCAGVVVACYSAAGAVFGTVPAAAAAAIPALAACNSAFGTCSHVCATVALLAPIP
>RPA97681.1 hypothetical protein L873DRAFT_1809484 [Choiromyces venosus 120613-1]
MKLQNILLPASFATMALAGPISYGICQGGCAGVVVACYSAAGAVFGTVPAAAAVAIPALAACNSAFGSCSAVCATVALFAPIP
>RPA97678.1 hypothetical protein L873DRAFT_1809475 [Choiromyces venosus 120613-1]
MKLQNLLLPASFATMALAGPISYGICQSGCAGVVVACYSAAGAVFGTVPAAAATAIPALAACNSAFGSCSHFCASVTLLALTP
>KAI9642175.1 hypothetical protein NHQ30_008977 [Ciborinia camelliae]
MKPALLLGAGALTILTSIPTTTAGPVTYGACQSGCAVVVMACYSAAGFTWGATLGATAPATVIACNVAFGKCSATCAGLLLAPTP
>KAF1937296.1 hypothetical protein EJ02DRAFT_458854 [Clathrospora elynae]
MKFLINIITTALAILIFATTASAGPIGYAICQGGCAGVVMACYSAAGFTWGATLGATAPATVLACNAAYGTCQAACAAVLLVPLP
>KAK1955786.1 hypothetical protein LY78DRAFT_542662, partial [Colletotrichum sublineola]
VLGGPGAYGVCQAGCSAVVMACYTAAGATWGATLGLTAAPSVIGCNVAYGTCQAACASVLLAPIP
>KAF0316341.1 hypothetical protein GQ607_016441 [Colletotrichum asianum]
MVAPAVVLAEKCAYGTCQTGCAALVVTCYTVAGGIFGVTSGVAATTAVKECDVAFGKCQASCSRARSAPRC
>KAF0316342.1 hypothetical protein GQ607_016442 [Colletotrichum asianum]
MDLSTILVLYVFLAAPPAVEAGLLGYGICQAGCAGVVTACYAAAGAVWGATAGVGAAPAVIACNVAFGKCQAACAVVALAPTP
>KAK1845575.1 hypothetical protein CCHR01_11807 [Colletotrichum chrysophilum]
MKPLTAILSTCSILAPAVVLAEKCAYGTCQTGCAALVVTCYTVAGGIFGVTSGVAATPAVKECDVAFGKCQASCSQARFAPRC
>KAK1997917.1 hypothetical protein LX36DRAFT_577335 [Colletotrichum falcatum]
MNPATGFLLLLIPALPGVQAGPAAYGICQAGCSAVVMACYAAAGATWGATLGLTAAPSVIACNVAYGTCQAACAAVILAPTP
**>XP_031884111.1 uncharacterized protein CGMCC3_g9227 [Colletotrichum fructicola]**

**MVVIILLLLSIFLAAPPAVEAGPLGYGICQAGCAGVVAACYAAAGAVWGATAGAAAAPAVIVCNLAFGKCQAACAAVALVPTP**
>XP_031884118.1 uncharacterized protein CGMCC3_g9228 [Colletotrichum fructicola]
MKFSTATLPTIYILAPATVLVGPCAYGTCQAGCAILAVACYFGTGATFGVTCGLAATPAVLVCNTAFGKCQASCWLAALAPTC
>KAH9242426.1 hypothetical protein K456DRAFT_1716149 [Colletotrichum gloeosporioides 23]
MKPETLAVALATAKLVMGNMLFGPMIREYAACQMNCATLAYACYIVMEKEWVTKCCIPPKARACYFAFEECQATCAYPIRFESK
>KAH9242424.1 hypothetical protein K456DRAFT_1805227, partial [Colletotrichum gloeosporioides 23]
LSVFLAAPPAVEAGPLGYGICQAGCAGVVTACYAAAGAVWGATAGAAAAPAVITCNVAFGKCQAACAVVALTPIP
>XP_045270162.1 uncharacterized protein GCG54_00010339, partial [Colletotrichum gloeosporioides]
VLLLLSVFLAAPPAVNTGPLSYSICQAGCAGVVTACYAAARAVWGATAGVAAAPAVIACHVAFRKCQAAYAVVALAPIP
>XP_008099621.1 uncharacterized protein GLRG_10745 [Colletotrichum graminicola M1.001]
MTPARSFTLLVIFSLPCVLAGPAAYGVCQRGCSSVVMACYKAAGSTWGATLGLTATPLVIGCNSAYGTCQAACAAVVFTLTP
>KAK2770104.1 hypothetical protein CKAH01_04447 [Colletotrichum kahawae]
MKLSMATLSTLSILAPTTVLAGLCAYGTCQAGCAALAVVCYSGTGATFGVTCGLAATPAVLACNAAFGQCQASCWLAGFAPTC
>KAK2770103.1 hypothetical protein CKAH01_04446 [Colletotrichum kahawae]
MKLNTLTTGLVTVKLAMGNMLFGPMIREYSACQMNCATLAYACYIVVEKEWVTKCCIPPKARACYFAFEECQATCAYPIRFESK
>KAF6819823.1 hypothetical protein CPLU01_12931 [Colletotrichum plurivorum]
MNPTHALATVALATIPPTEAGPIGYGVCQAGCAGVVTACYAAAAAVWSATAGAAAAPAVIACNVAFGKCQAACAAVALLPTP
>TEA19707.1 hypothetical protein C8034_v008990 [Colletotrichum sidae]
MKYTSNLTLLTLALLPAVICGPIEYGICQSGCAAILRACYVPKDARWRQEGGFPAPPSTKKCDDAFWDCQDRCEIAAYAPGL
>TEA19704.1 hypothetical protein C8034_v008993 [Colletotrichum sidae]
MNPSGFITLLTLAALPVAVAGPAAYGVCQAGCASIVVACYAAAGAVFGATAGAAAPPAVVACNIAFGKCQAACAMSAICPIP
>KAF6802731.1 hypothetical protein CSOJ01_11404 [Colletotrichum sojae]
MNPTQALVTIALVTIPTATAGPVGYGICQAGCAAVVVACYAGAGATFGVTAGVATGPAVIACNAAFGQCQAA
>KAK2042028.1 hypothetical protein LZ31DRAFT_471277 [Colletotrichum somersetense]
MKPASNFLFLVTSCLPCALAGPAAYGICQAGCSAVVMACYTAGGATWGATLGAPAGPTIIGCNLAYGTCQAACASVILAPTL
>TDZ35452.1 hypothetical protein C8035_v009313 [Colletotrichum spinosum]
MNPSGFITLFTLAALPVAVAGPAAYGVCQAGCASIVVACYAAAGAVFGATAGAAAPPAVVACNVAFGKCQCACAMSAICPIP
>TDZ41239.1 hypothetical protein CTRI78_v009824 [Colletotrichum trifolii]
MKFTSNLTLLALALLPAVICGPIEYGICQSGCAAILRACYVPKGARWREEGGFPAPPSTKKCDDAFWDCQDRCEIVAYAPGL
>TDZ41237.1 hypothetical protein CTRI78_v009826 [Colletotrichum trifolii]
MNPSGFITLFTLVALPVAVAGPAAYGVCQAGCASIVVACYAAAGAVFGAIAGAAAPPAVVACNVAFGKCQCACAMSAICPIP
>TDZ35453.1 hypothetical protein C8035_v009310 [Colletotrichum spinosum]
MKYTSNLTLLALALLPAVICGPIEYGICQSGCAAILRACYVPKDARWRQEGGFPAPPSTKKCDDAFWDCQDRCEIVAYAPGL
>KAH9242425.1 hypothetical protein K456DRAFT_1820222 [Colletotrichum gloeosporioides 23]
MKFSAAILSTIYILAPATVLAGPCAYGTCQAGCAVLTVACYSGTGATFGVTCGLAATPAVLACNAAFGQCQASCWLAGLAPTC
>XP_060407788.1 uncharacterized protein LY79DRAFT_528443 [Colletotrichum navitas]
MKTARSFTLLVTSFLPCVLAGPAAYGVCQAGCSAVVMACYAAAGATWGATLGLTAAPTVIGCNLAYGTCQAACAAVILAPTP
>KAK2015154.1 hypothetical protein LZ32DRAFT_513037, partial [Colletotrichum eremochloae]
LLLIPCLPGVLGGPGAYGVCQAGCSAVVMACYTAAGATWGAALGLTAAPSVIGCNVAYGTCQAACSSVLLAPIP
>KAK1977458.1 hypothetical protein LZ30DRAFT_664953 [Colletotrichum cereale]
MKWSVRVVTHILWLLSSSLCVLAGPLGYGVCQAGCSAVVMACYTAAGATWGATLGLTASESVISYNLAFGKCQAACAAVPLSPTP
>KAK2031093.1 hypothetical protein LX32DRAFT_637589 [Colletotrichum zoysiae]
MKPASNFLLLVTSCLPFVLAGPAAYGICQAGCSAVVMACYTAGGATWGATLGATAGPTIVGCNLAYGTCQAACAAVILAPTL
>OIW28253.1 hypothetical protein CONLIGDRAFT_577608 [Coniochaeta ligniaria NRRL 30616]
MRASSAIPAISLLFLAQGAVASLVGYGICQAGCAGVVMACYSAAGFTWGATMGVSAPATIIACNSAFGSCQAACATVLLAPIP
>RKU46238.1 hypothetical protein DL546_005261 [Coniochaeta pulveracea]
MRLSRTFTAFTLAPAACAGPLAYGMCQAGCSGIVMACYAAAGFTWGATLGATAPASILLCNSSYGSCQAVCAAIALAPTP
>KAB5545907.1 hypothetical protein GE09DRAFT_222926 [Coniochaeta sp. 2T2.1]
*MRVFSAIPTVSILLLASEVIA*GPVGYGVCQGGCAGVVMACYGAAGFTWGATLGATAPASILACNAAYGSCQAACAAVLLVPLP
>TQW06488.1 hypothetical protein IF2G_05910 [Cordyceps javanica]
*MRDHKIIFPFLVGTAAA*GPLAYGICQAGCAGVVMACYGAAGATWGATAGASAAPTVIACNLAFGKCQAACVVAGCSPTP
>TQV90198.1 hypothetical protein IF1G_11149 [Cordyceps javanica]
MRDQKIILPLLVGLATAGPLAYGVCQAGCSAVVMACYAAAGATWGATAGATAAPAVVACNLAFGKCQAACSVAGFCPSV
>PSN58987.1 hypothetical protein BS50DRAFT_580308 [Corynespora cassiicola Philippines]
MHSSFPATIIAYLALASAVSAGPAAYGVCQAGCATVVMACYSAAGFTWGATLGASAPPTIIACNAAFGTCYSACAATLLAPTL
>PSN59268.1 hypothetical protein BS50DRAFT_580102 [Corynespora cassiicola Philippines]
MHSSFPATIIVYLALASAVSAGPAAYGICQAGCATVVMACYSAAGFTWGATLGASAPPTIIACNAALGTCYSACAATLLAPTL
>USP72936.1 hypothetical protein yc1106_00210 [Curvularia clavata]
MKSTTITKASVALLTFTTGRVSAGHIGYGICQAGCASVVMACYAAAGYTWGATMGATAPASIIACNSAFGTCQAACAAVLLGPTL
>ORY12622.1 hypothetical protein BCR34DRAFT_289991 [Clohesyomyces aquaticus]
MRLSNLSIAAAALLMLPDFTSAGPAAYGVCQAGCAAVVVACYGAAGFTWGATLGASAPATIIACNTTFGTCQAACWAALFTPTP
>XP_047795458.1 uncharacterized protein F4812DRAFT_411739 [Daldinia caldariorum]
MHFGKVSAVVMAVATCATAGPLAYAACQSACAGATLWVPFFGIPAYAGCQSSCAHLLLAPTP
>XP_049156355.1 uncharacterized protein F4817DRAFT_346277 [Daldinia loculata]
MHQAKVSMVVMAIATCATAGPWTYAGCQSACAAGTLWIPAFSTAAYASCQSYCAYFLLAPTP
>KAI2778314.1 hypothetical protein F4815DRAFT_477950 [Daldinia loculata]
MHQAKVSMVVMAIATCATAGPWTYAGCQSACAAGTLWIPVFSTPAYASCQSYCAYYLLAPTP
>KAF1839331.1 hypothetical protein BDW02DRAFT_563871 [Decorospora gaudefroyi]
MKLPFFTTMAAAFLILACLASAGPVAYGICQGGCSAVVMACYSAAGFTWGATLGASAPATIVACNAAFGTCQAACAAILLAPTF
>KAF1356704.1 hypothetical protein BDV97DRAFT_13626 [Delphinella strobiligena]
MRLLALKPFWLTAMFITSCSAGPAAYGICQAGCCGVVTACYSAAGFTWGATLGATAPASILACNSAFGACQAACAVVALAPTL

>KAH7119993.1 hypothetical protein B0J11DRAFT_440189 [Dendryphion nanum]
MRLYNLAKYLPAVFFFNSTVYAGPAAYGICQAGCAGIVAACYGAAGCTWGATLGATAPATIIACNLAFGKCQAACAALLLAPTL

>KAH8743362.1 hypothetical protein F5883DRAFT_441037 [Diaporthaceae sp. PMI_573]
MQLLKSLVATLFLASSVAAGPIAYAICQAGCSSLAVACYSAAGFTFGTVAAAAAPAALLACNSAYGTCQPACVVAALVPTLGKRS
>KAH8759482.1 hypothetical protein F5883DRAFT_564952 [Diaporthaceae sp. PMI_573]
MRFTTLALVLAGITPAIGGPAVSVACHVACATALTACIASGFYFPPMIAQCQAAYGVCQLACTSSAFLPTP
>KAH8767254.1 hypothetical protein F5883DRAFT_419094 [Diaporthaceae sp. PMI_573]
MQPTKTIIAALSLATALTAGPVAYGICQAGCAAVVTACYAGAGFTFGTVLAEAAPPAIIACNSAFGTCQASCAIVALFPCP
>POS77299.1 hypothetical protein DHEL01_v204301 [Diaporthe helianthi]

MQPTKILITAMSLGTALAAEVAYGVCQAICTRQALTCYDKAGVTFGSKHQLGTEFNITTCNVALDTCVRICIFVANGA
>XP_044637434.1 uncharacterized protein KVR01_013659 [Diaporthe batatas]
MQPTGILITAMTVATALAAELTYGVCQAICTRQALTCYDDAGVTVGSKNAYGSAANVTACNVTLDTCGLVCAMVASGGCDSSECC
**>KAI7781638.1 hypothetical protein LA080_014536 [Diaporthe eres]**
**MQLTKIIIPALSLVTAVAAGPIAYGTCQAGCAAVVVACYAGAGFTFGTVLGPAAPASIAACNTAFGFCQAKCAVVALLPTP**
>KAI3393011.1 hypothetical protein diail_4890, partial [Diaporthe ilicicola]
LHESLVATLFLASSVAAGPIAYAVCQAGCSSLAVACYSAAGFTFGTIAAAAAPAALIACNSAYGTCQAACAAAALAPTL
>KAF2262031.1 hypothetical protein CC78DRAFT_535094 [Didymosphaeria enalia]
MRLSSLSVELLLLAGTASAGPAAYGVCQAGCAGVVMACYSAAGFTWGATMGASAPATIVACNAAFGSCQAACAAALIAPTL
>XP_052987644.1 uncharacterized protein J7T55_007670 [Diaporthe amygdali]
MVIITMRFTRTIIAALPLAIAIIAGPNAYSICQAGCSAVLAACYSAAGARVGTVYGSAVPVPIADCNAAFMACNAMCAVITLSQTA
>KAH8747636.1 hypothetical protein F5883DRAFT_437093 [Diaporthaceae sp. PMI_573]
MRLQKSLVANLFLASSVAAGPVVYAVCQAGCSSLAVACYSAAGFTLGIIATTAAPAAVIACKTAYGTCQAACAAAALAATP
>KAI6712870.1 hypothetical protein JHW43_004599 [Diplocarpon mali]
MRLPIISVLFAAPAVVFSGPLAYGVCQGGCASLVMACYTAAGATWGATAGILAPATVVACNTAFGYCQAKCAVVAWLPTL
>KAK2628309.1 hypothetical protein QTJ16_002955 [Diplocarpon rosae]
MLLPAILVLFAAPAVVFSGPIAYGTCQGGCAAVVMACYTAAGATWGATLGLNAPATVLGCNSAFGYCQAKCAVVALLPTF
>PBP25780.1 hypothetical protein BUE80_DR003222 [Diplocarpon rosae]
MRLLVILVLFVAPAVVFSGPVAYGTCQSGCAAVVMACYTAAGATWGTTLGLTAPTTVLGCNTAFGYCQAKCAVVALLPTL
>XP_033524879.1 uncharacterized protein P153DRAFT_366077 [Dothidotthia symphoricarpi CBS 119687]
MRFPTTLTTLTAMLALIPCATAGPIGYAICQAGCSSVVMACYAAAGFTWGATLGATAPASIVACNTAFGTCQAACATVLLGPLP
>KAF2218680.1 hypothetical protein BDZ85DRAFT_180405, partial [Elsinoe ampelina]
VLPGLAQAGPAAYGVCQAGCAGIVMACYAAAGFTWGATLGATAPASIIACNTTFGACQATCAALLLAPTP
>KAF7513366.1 hypothetical protein GJ744_009787 [Endocarpon pusillum]
MKATAFVPLLSLTLTPLVFAGPLSYGICQAGCASIVMACYAAAGATWGATLGATAPASVLACNAAYGSCQAACIAAGCIPIP
>XP_033535919.1 uncharacterized protein P152DRAFT_392925, partial [Eremomyces bilateralis CBS 781.70]
SNLLIIILPFTVPIITVSAGPVLYAGCQVGCAGAVMACYSAAGFTWGATLGITAPATIIACNMAFGSCQAGCAAALLILTL
>KAI1248405.1 hypothetical protein MGN70_009603 [Eutypa lata]
MKVTTPAAALVAILTAAPGAIAGPAAYGICQAGCADVVTACYAAAGFTWGATLGASAPATIVACNTAFGSCQSACWAALVAPTP
>XP_008028533.1 uncharacterized protein SETTUDRAFT_93357 [Exserohilum turcica Et28A]
MNLPSLNNTAVAILILSSQTSAGPLGYAVCQAGCASVVMACYSAAGYVWGNTLSADAPDAVIGCNLAFGKCQAACAEVTLGTAM
>XP_047761909.1 uncharacterized protein CLAFUR5_05392 [Fulvia fulva]
MPSRTFTLLATLTTLVPLANAGPAAYGICQTGCAGVVMACYGAAGFTWGATLGATAPASVLACNSAYGACQAACASTLLVPFF
>XP_047765020.1 uncharacterized protein CLAFUR5_10858 [Fulvia fulva]
MFLPILPILIALVALLPTAIGGPAAYGICQAGCASVVVACYAAAGFVFGVALPAAPPAILACNSAFGTCQAACAVVVFAPTV
>KAF5254884.1 hypothetical protein FANTH_380 [Fusarium anthophilum]
MKSLHLLVAVGSFSTAAVAGPIAYGVCQAGCASVAMACYAAGGATWGATAGATAPATIAACNSAFGVCSAACAQTALIAPTP
>KAF5230862.1 hypothetical protein FAUST_9596 [Fusarium austroamericanum]
MKSSSTIVITTAFLFTVSAGPLAYGACQAGCAGIVMACYSAAGYIWGATARATAPASIIDCNAAFGKCSANETFSETLPCHQE
>KAH6963189.1 hypothetical protein DER45DRAFT_480060 [Fusarium avenaceum]
MKPAAILTTLTCLYTTAAAGPIAYGVCQAGCSAVVMACYAAAGYTWGATLGASAPASIVACNGAFGTCSAACAKIGLFAPTP
>KAF4339535.1 hypothetical protein FBEOM_6561 [Fusarium beomiforme]
MKPLHLFTTITCLSATAHAGPVAYGVCQAGCASVAMACYAAGGATWGATAGATAPATIVACNSAFGACSAACAQTALIAPIP
>KAF5989468.1 hypothetical protein FBULB1_896 [Fusarium bulbicola]
MKPLHLLVAITSLSTTAAAGPIAYGVCQAGCASVAMACYAAGGATWGATAGATAPATIAACNSAFGVCSAACAQTALIAPTP
>QPC57932.1 hypothetical protein HYE67_000163 [Fusarium culmorum]
MKSSSTIAITIALLTTVSAGPLAYGACQAGCASIVMACYSAAGFTWGATAGATAPASIIACNAAFGKCSAVCAAIALGAPTP
>CAG7562008.1 unnamed protein product [Fusarium equiseti]
MKPTTPLITIACVASPVSAGPLAYAACQSGCAGVVMACYSAAGFTWGATFGASAPASILLCNSAFGTCSATCAQVALFAPTP
>XP_045982074.1 uncharacterized protein B0J16DRAFT_165472 [Fusarium flagelliforme]
MKPTTPLIAIACLASTVSAGPLAYAACQAGCATIVMACYSAAGYTWGATLGASAPATILACNAAFGQCSAVCAQVALVAPTP
**>VTO82721.1 unnamed protein product [Fusarium graminearum]**
**MKSSSTIVITTAFLFTVSAGPLAYGACQAGCAGIVMACYSAAGYIWEATAGVTAPASIIDCNAAFGKCSAVCAAIALGTPTS**
>KAJ4012322.1 hypothetical protein NW752_007996 [Fusarium irregulare]
MKPTTPLIAIAYAASTVSAGPLAYAACQSGCAGVVMACYSAAGFTWGATFGATAPASILLCNAAFGTCSATCAQVALLAPTP
>KAF5552569.1 hypothetical protein FMEXI_2720 [Fusarium mexicanum]
MKPMHLLVAITSLSTTAVAGPIAYGVCQAGCASVAMACYAAGGATWGATAGATAPATIAACNSAFGVCSAACAQTALIAPTP
>PNP60693.1 hypothetical protein FNYG_14578 [Fusarium nygamai]
MTSRMMLMSPVATLAILSTTVSAGPVAYGVCQAGYAGVIMACYGAAGYTWGATVGATAPATIIACNSAFRTCSAACAKVALLAPTV
>PNP73544.1 hypothetical protein FNYG_13138 [Fusarium nygamai]
MKPLPLLVAIASFSTTTVAGPIAYGVCQAGCASVAMACYAAGGATWGATAGATAPATIAACNSAFGVCSAACAQTALIAPTP
>EMT71922.1 hypothetical protein FOC4_g10003765 [Fusarium odoratissimum]
MKAISFLATIAGFSTTARAGPIAYGVCQAGCASVVMACYAAGGATWGATAGATAPATIAACNSAFGVCSAACAQTALVAPTP
>EXK76156.1 hypothetical protein FOQG_19086 [Fusarium oxysporum f. sp. raphani 54005]
MKLLSVVITLAILSTTVSTGPVAYGVCQAGCAGVVMACYGAAGYTWGATVGATAPATIVACNSAFGTCSAACAKVALLAPTP
>ENH74830.1 hypothetical protein FOC1_g10003190 [Fusarium oxysporum f. sp. cubense race 1]
MKAISFLTIIAGFSTTARAGPIAYGVCQAGCASVVMACYAAGGATWGATAGATAPVTIAACNSAFGVCSAACAQTALIAPTA
>KAG7410298.1 hypothetical protein Forpi1262_v017615 [Fusarium oxysporum f. sp. raphani]
MKLLSVVTTLAILSTTVSTGPVAYGVCQAGCAGVVMACYGAAGYTWGATVGATAPATIIACNSAFGTCSAACAKVALLAPTP
>RKK72021.1 hypothetical protein BFJ69_g10421 [Fusarium oxysporum]
MKAISFLATIAGFSTTARAGPIAYGVCQAGCASVVMACYAAGGATWGATAGATAPATIAACNSAFGVCSAACAQTALIAPTP
>KAI8397321.1 hypothetical protein FOFC_20593 [Fusarium oxysporum]
MKLLSVVTTLAILSTTVSTGPVAYGVCQAGCAGVVMACYGAAGLQQRVWDMLRGLCQGSLAGPYTMK
>KAH7466833.1 hypothetical protein FOMA001_g16573 [Fusarium oxysporum f. sp. matthiolae]
MKAISFLKIIAGFSTTARAGPIAYGVCQAGCASVVMACYAAGGATWGATAGATAPATIAACNSAFGVCSAACAQTALIAPTP
>XP_059468083.1 uncharacterized protein FOBCDRAFT_233752 [Fusarium oxysporum Fo47]
MKAIGFFTIIAGFSTTARAGPIAYGVCQAGCASVVMACYAAGGATWGATAGATAPATIAACNSAFGVCSAACAQTALIAPTP
>EXK76060.1 hypothetical protein FOQG_19180 [Fusarium oxysporum f. sp. raphani 54005]
MKLLSVVTTLAILSTTVSTGPVAYGICQAGCAGVVMACYGAAGYTWGATVGATAPATIIACNSAFGTCSAACAKVALLAPTP
>QPC69322.1 hypothetical protein HYE68_000074 [Fusarium pseudograminearum]
MKFSSTIAITIALFDTVSAGPLAYGACQAGCASIVMACYSAAGFTWGATAGATAPASIIACNAAFGKCSAVCAAIALGAPTP
>KAF5592368.1 hypothetical protein FPCIR_5689 [Fusarium pseudocircinatum]

MKPLHLLVTIASFSTTAVAGPIAYGVCQAGCASVAMACYAAGGATWGATAGATAPATIAACNSAFGVCSAACAQTALIAPTP
>KAF5574014.1 hypothetical protein FPANT_12003 [Fusarium pseudoanthophilum]
MKPLHLLVATVFFSTTVVAGPIAYGVYQAGYASVAMACYAAGGATWGATAGATAPATIAAYNSAFGVCSAACAQTALIAPTP
>XP_046041803.1 uncharacterized protein BKA55DRAFT_599821 [Fusarium redolens]
MKAISFLAIIAGFSATASAGPIAYGVCQAGCASVAMACYAAGGATWGATAGATAPATIAACNSAFGVCSAACAQTALIAPTP
>XP_036534571.1 uncharacterized protein FSUBG_9910 [Fusarium subglutinans]
MKPLHLLVAITSLSTTAVAGPIAYGVCQAGCASVAMACYAAGGATWGATAGATAPATIAACNSAFGVCSAACAQTALIAPTP
>SPJ71122.1 uncharacterized protein FTOL_00850 [Fusarium torulosum]
MKFSVVLATLACLYTTTTAGPIGYGICQSGCSAVVMACYAAAGYTWGATLGASAPASIIACNSAFGTCSAVCAKVALLAPTP
>KAH7251348.1 hypothetical protein BKA59DRAFT_416074, partial [Fusarium tricinctum]
MKPTAILATLACFYTPTSAGPIAYGVCQAGCSAVVMACYTAAGYTWGATLGASAPASIIACNGAFGTCSAACAKIGLFAPTP
>XP_018761531.1 hypothetical protein FVEG_13353 [Fusarium verticillioides 7600]
MKPLHLLVTITSFSTTVVAGPIAYGVCQAGCASVAMACYAAGGATWGATAGATAPATIAACNSAFGVCSAACAQTALIAPTP
>RBQ81898.1 hypothetical protein FVER14953_21526 [Fusarium verticillioides]
MKPRVTVAVAALFFTGASAGPQAYAACRAGCSAIVMACYGAAGFVWGTRLSATAPPTIIASNNAFGTCSAVCAKVALFAPTP
>KAH7003006.1 hypothetical protein EDB82DRAFT_482357 [Fusarium venenatum]
MKISTITATTALLATAFAGPLAYGTCQAGCASVVVACYSAAGFTWGATAGATAPATIIACNAAFGKCSAVCAAIALAPTP
>KAG5753170.1 hypothetical protein H9Q70_004225 [Fusarium xylarioides]
MKPLRFLVAIASFSTTTVAGPIAYGVCQAGCASVAMACYAAGGATWGATAGATAPATIAACNSAFGVCSAACAQTALIAPTP
>KAH7151049.1 hypothetical protein DER46DRAFT_614204 [Fusarium sp. MPI-SDFR-AT-0072]
MKAISFLATIAGFSATACAGPIAYGVCQAGCASVAMACYAAGGATWGATAGATAPATIAACNSAFGVCSAACAQTALIAPTP
>KAI1061423.1 hypothetical protein LB507_011234 [Fusarium sp. FIESC RH6]
MKPSTPLSAIACAASTVSAGPLAYAACQSGCAGVVMACYSAAGFTWGATFGATAPASILLCNAAFGTCSATCAQVALLAPTP
>CAJ0554546.1 Ff.00g130590.m01.CDS01 [Fusarium sp. VM40]
MKPAAILATLACFYPATSAGPIAYGICQAGCSGVVMACYAAAGYTWGATLGASAPASIVACNGAFGTCSAACAKIGLFAPTP
>XP_009230175.1 hypothetical protein GGTG_13986 [Gaeumannomyces tritici R3-111a-1]
MRPSTSVSALLAAMAFAGLALAGPIAYGICQAGCAALVMACYAAAGATWGATLGATAPATVIACNSAFGTCQAGCATALLLPIP
>XP_009229506.1 hypothetical protein GGTG_13336 [Gaeumannomyces tritici R3-111a-1]
MRRFTSASIMLVTMAFTSPAFAGPAAYGVCQAGCAAVVMACYSAAGFTWGATLGVSAPPTIIACNTSFGTCQAACAAVLLSPTP
>OCL09912.1 hypothetical protein AOQ84DRAFT_290324 [Glonium stellatum]
MHPRRVLLATLTLTHFAYAGPAAYGICQAGCAAVVMACYSAAGFTWGATLGIAAPATILACNSAFGTCQGACAVAFIAPTP
>KAH6664969.1 hypothetical protein B0J14DRAFT_661079 [Halenospora varia]
MRPTILLTPLLLTPALAGPALYGICQAGCASVVMACYGAAGFTWGATLGASAPATVLACNAAFGTCSAACAAAVLAPTL
>KAH8653200.1 zygote-specific protein, partial [Hymenoscyphus varicosporioides]
LASPALAGPALYGICQAGCATVVMACYTAAGFTWGATLGASAPPTILACNAAFGTCSAACAAVILAPTP
>XP_049117280.1 uncharacterized protein GGS25DRAFT_493230 [Hypoxylon fragiforme]
MKSTKFLASTLFAATTIAGPIAYGVCQAGCSAVVMACYSAAGFTWGATLGASAPASIILCNTSYGSCQAACWAALAMPAP
>KAF8463719.1 cysteine-rich protein [Kalaharituber pfeilii]
MRFHRSLSTLLPALLLVTGTEAGPIGYALCQGGCATVVMACYAAAGAVWGTNPWLAAGVPVPAVIVGCNSAYAGCQATCAAVAFSPTP
>KAF8463720.1 hypothetical protein BDZ91DRAFT_627842, partial [Kalaharituber pfeilii]
SIFTLFPALLLVTGTEAGPVGYALCQGGCASVVMACYSAAGAVWGTNPWLAAGVPVPAAIVGCNSAYAGCQAKCAVVALSPTP
>KAF8463718.1 cysteine-rich protein [Kalaharituber pfeilii]
MRFHRSLSTLLPALLLVTGTEAGPIGYALCQGGCATVVMACYTAAGAVWGTNPWLAAGVPVPAVIVGCNSAYAGCQATCAAVALSPTP
>KAF8453073.1 hypothetical protein BDZ91DRAFT_749804, partial [Kalaharituber pfeilii]
*MRFHRSLFTLLPALLLVTGTEA*GPIAYAICQGGCATVVVACYAAAGAVWGTNPWLAAGVSVPAAIVGCNSAYAGCQATCAAVVLLP
>KAF2447884.1 hypothetical protein P171DRAFT_354022 [Karstenula rhodostoma CBS 690.94]
MRFHTIAATAVVGLLVGTAAAGPIGYGICQTGCAAIVVACYSAAGFTFGTVLAAAAPPAILACNSAYGTCQAACAVVLLGPTP
>KAH6977657.1 hypothetical protein EDB80DRAFT_657849 [Ilyonectria destructans]
MKLLSPVPTLAILSTAASAGPVAYGVCQAGCAGVVMACYGAAGYTWGATIGATAPATIVACNSAFGTCSAACAKVALLAPIP
>TVY53193.1 hypothetical protein LCER1_G004518 [Lachnellula cervina]
MKISRIIAALATTTTLVTAGPAAYGLCQTGCAGVVVACYSAAGFTFGTVAAAAAPAAILACDAAYGSCCAACYWALFLPTP
>TVY92812.1 hypothetical protein LAWI1_G003505 [Lachnellula willkommii]
MKISRIIAALATTTTLVAAGPAAYGLCQTGCVGVAVACYSAAGFTFGTVAAATAPAAIVACNSAYGSCCATCHALLFAPTP
>KAF2687625.1 hypothetical protein K458DRAFT_248623, partial [Lentithecium fluviatile CBS 122367]
SISLLVLAGSVSAGPAAYGVCQAGCAAVVTACYSAAGFTWGATLGASAPATIVACNAAFGTCQAACAATLLAPT
>KAF9638679.1 hypothetical protein BFW01_g9576 [Lasiodiplodia theobromae]
MRFQTIVFAGSLIFAGTATAGPIAYGICQAGCASVVMACYAAAGFTWGATLGATAPASIIACNAAFGTCQAACAVVALAPTP
>KAH6709034.1 hypothetical protein BKA61DRAFT_738127 [Leptodontidium sp. MPI-SDFR-AT-0119]
KHFKICLPNPFLILLATSPTLILGGPLAYGICQAGCAAVMTACYAAGGATWGATLGSNAPATIVACNSAFGTCQAACAVVALMHTL
>OCK77742.1 hypothetical protein K432DRAFT_436347 [Lepidopterella palustris CBS 459.81]
MKQCQNLLIIVRFSVPLAIILAGAASTGPIGYGICQGGCATIVMACYGAAGATWGATLGATAPATVIACNSAYGACQAACASVLLLPTP
>XP_037161484.1 uncharacterized protein HO173_009890 [Letharia columbiana]
MRLTKLIPMTTIITSALAGPAAYGICQAGCAGVAVACYAAAGAIMGVTAGAAAPPAVLACNAAFGSCQAACAAALVMPTP
>KAF2648513.1 hypothetical protein K491DRAFT_576600, partial [Lophiostoma macrostomum CBS 122681]
LATVITLSTLAGEVAAGPAAYGVCQAGCSAIVMACYSAAGFTWGATLGASAPATVLGCNTAFGACQAACAATLLAPT
>KAF2490063.1 hypothetical protein BU16DRAFT_518757 [Lophium mytilinum]
MKLRLSILALLTGAAVPISAGPIGYGICQAGCAGVVMACYSAGGATWGATLGATAPATIIGCNTAFGSCQAACAAVLLAPTP
>KAH7025312.1 hypothetical protein B0J12DRAFT_714445 [Macrophomina phaseolina]
MLSSRSLVIGFILATQSLVSAGPGAYGVCQAGCSGVAMACYAAAGFTWGATLGATAPASIVACNSAFGACQAACAAVLLAPTL
>KAH7014050.1 hypothetical protein B0J12DRAFT_586659 [Macrophomina phaseolina]
MPASRSLLVGFILATQSLVSAGPTAYGVCQAECSGVVMACHAAAGFTRGATLGATAPASIVACNSAFGACQAACAVVLLAPTL
>XP_033556128.1 uncharacterized protein BU25DRAFT_353360 [Macroventuria anomochaeta]
MAAAFLILACLASAGPVAYGICQGGCSAVVMACYSAAGFTWGATLGATAPATVVACNAAFGTCQAACAAVLLAPTL
>XP_033563937.1 uncharacterized protein BU25DRAFT_337128 [Macroventuria anomochaeta]
MLLLTKIVLAGPIGYGICQTSCSAFVMACYSATGFTWGATLAATAPASILVCNAAYGTCQTACATVLLGPTL
>KAF2867966.1 hypothetical protein BDV95DRAFT_581004 [Massariosphaeria phaeospora]
MRLSSVLILLAGSVSAGPAAYGACQGGCAKIVMACYAAAGCTWGATMGASAPPTIIACNTAFGSCQAVCAALLLAPTL
>KAA8570577.1 hypothetical protein EYC84_002838 [Monilinia fructicola]
MKLQSASRFSIPLATTLLTLTLFTIIPPATSGPIAYASCQAGCAAVVTACYSAAGFTWGATLGATAPASIVACNAAFGTCYAACAGFLVAPIP
>KAB8299301.1 hypothetical protein EYC80_001377 [Monilinia laxa]
MNLQSVSLFSFLLPLLALFLLNIIPPVTAGPIAYASCQAGCAGIVMACYSAAGFTWGATLGVTAPASIVACNAAFGTCYAACAGFLVAPIP
>RYO93046.1 hypothetical protein DL762_001309 [Monosporascus cannonballus]

MKLTTSILLAIVAATPTIHAGPAAYGVCQAGCSAVVQTCYAAAGFTWGATLGATAPASIVACNSAYGACQAACWTALFSLTP
**>RYP08623.1 hypothetical protein DL764_001751 [Monosporascus ibericus]**
**MQPTIPILLALIATTPTILAGPAAYGVCQAGCSGVVMACYAAAGFTWGATLGATAPASIVACNTAYGACQAACWAALAAPTP**
>RYP51836.1 hypothetical protein DL768_002912 [Monosporascus sp. mg162]
MKPTTPIRLALIATTPTVFAGPAAYGVCQAGCSGVVMACYAAAGFTWGATLGATAPASIIACNSAYGACQAACWAALAAPTP
>RYP76942.1 hypothetical protein DL771_001495 [Monosporascus sp. 5C6A]
MKPTSPILLALIAATPTILAGPAAYGVCQAGCSGVVMACYAAAGFTWGATLGATAPASIVACNTAFGACQAACWAALAAPTP
>RYP75671.1 hypothetical protein DL770_007356 [Monosporascus sp. CRB-9-2]
MKRTSPILLTLIAATPTILAGPAAYGVCQTGCAGVVMACYAAAGFTWGATLGATAPASIVACNSAYGACQAACWAALAAPTP
>RYP24059.1 hypothetical protein DL765_000787 [Monosporascus sp. GIB2]
MKLTTSILLAIVAAVPTIHAGPAAYGACQAGCSAVVQACYAAAGFTWGATLGATAPASIVACNNAYGACQAACWAALFSPTP
>RYP30960.1 hypothetical protein DL767_006002 [Monosporascus sp. MG133]
MKPTTPILLAIVAAMPTILAGPAAYGICQAGCSAVVKACYAAAGFTWGATLGATAPATIVACNGAYGACQASCWAALFAPTP
>KAF2663912.1 hypothetical protein BT63DRAFT_461030 [Microthyrium microscopicum]
*MRLSIPLFAAALLAPSAFA*GPAAYGICQAGCAAVVAACYAAAGATFGVAPPVAIPALAACNTAFGTCQAACWAALLLPTP
>XP_033573400.1 uncharacterized protein BDZ99DRAFT_466009 [Mytilinidion resinicola]
MKLRASILALLTTFTIPIAAGPIGYGVCQAGCAGLVMACYSAAGFTWGATLGATAPATIVACNTAFGSCQAACAAVLLTPTP
>KAI1156788.1 hypothetical protein F4825DRAFT_402532 [Nemania diffusa]
MKPITPFITAILAFVPVGSAGPAAYSLCQGGCSAVVMACYEAAGFTWGATMGASAPATIVACNTAFGTCQASCWAAAIAPTP
>KAI1118733.1 hypothetical protein F5Y14DRAFT_178302 [Nemania sp. NC0429]
*MQPIRAFAATILVFAPIACA*GPAAYGVCQAGCAAVVMACYGAAGCTWGATLGASAPATIIACNAAFGVCQAACWVAAISPGP
>KAI1132034.1 hypothetical protein F5Y10DRAFT_231933 [Nemania abortiva]
*MKLTTTFLAALAVLAPLGNA*GPAAYGLCQAGCASVVTACYAAAGFTWGATLGASAPASIVACNAAFGTCQAACAAIALAPTP
>KAI1176761.1 hypothetical protein F4777DRAFT_545056 [Nemania sp. FL0916]
MKLNTHLVSALAFTSAASAGPIAYGLCQAGCAAVVTACYGAAGFTWGATMGASAPASIVACNTAFGACQAGCWAAAIAPTP
>KAI1187125.1 hypothetical protein F5B17DRAFT_400959 [Nemania serpens]
*MKPTKIFTAAILAFAPIVSA*GPAAYGVCQAGCAAVVTACYAAAGFTWGATLGASAPATIIACNTAFGTCQAACWLAAIAPTP
>KAF2825151.1 zygote-specific protein, partial [Ophiobolus disseminans]
LLTNTAVAGPIGYGICQAGCSGVVMACYSAAGFTWGATLGATAPASILACNSAYGACQAACAAVLLGPWP
>XP_056783524.1 uncharacterized protein N7478_001698 [Penicillium angulare]
MKLSILAAALPLMAANVLAGPIGYAICQAGCASVVMACYAAGGATWGATLGASAPPTIVACNTAYGTCQAACAAVLLGPTP
>KAJ5116294.1 hypothetical protein N7456_000642 [Penicillium angulare]
MKICFLAILAALLLAGDAIAEPAARGKCHSACEKAVKKCYKKAGHKWGAPLANPPAPIIVCNKAFGACQSTCPRS
>XP_056783523.1 uncharacterized protein N7478_001697 [Penicillium angulare]
*MKLYLLAIFAAFLLAGEALA*APSIGSKCHSSCEKAVKKCYKKAGHKWGAPLANPPAAVVTCNKAFGACKSTCPK
>KAJ5116293.1 hypothetical protein N7456_000641 [Penicillium angulare]
*MKLSIIAAAFLLMAVNVLA*GPIGYGICQAGCASLVMACYAAGGATWGATLGATAPPTIVGCNTAFGTCQVACAAVLLVPFP
>XP_056817142.1 uncharacterized protein N7506_001931 [Penicillium brevicompactum]
*MVSSMAMLHLVLTLAPYVPGW*IEYGKCQAGCATLVMACYAGEGAAWGDTLGKTAPPPVKTCNSNFGDCQAECAQRWLWPTS
>XP_056813643.1 uncharacterized protein N7506_005460 [Penicillium brevicompactum]
MRAPWVLINNLLLFIPPVSAGPAAYGICQAGCSAVVMACYSAAGFTWGATMGASAPASVIVCNSAFGTCQAACAAALLAPTL
>KAJ6163906.1 hypothetical protein N7497_003885 [Penicillium chrysogenum]
MRVPWVLINNLLLFIPPVSAGPAAYGVCQAGCAAVVMACYSAAGFTWGATMGARAPASVMAFNLAFGKCQAACAAVLLAPTL
>XP_056549082.1 uncharacterized protein N7496_012271 [Penicillium cataractarum]
MRLSLPIHVLFLCFSTAAVAGPVGYGVCQAGCAGVTMACYSAAGFTWGATAGATAPATIVGCNLAFGKCQAACAAVLLMPTP
>XP_058332260.1 uncharacterized protein N7468_003960 [Penicillium chermesinum]
MKASSILSFLPIFAPAVFAGPAGYGVCQAGCATVVMACYSAAGFTWGATAGLSAPASIIACNTAFGTCQSACAAVLLAPTP
>XP_056573620.1 uncharacterized protein N7489_003563 [Penicillium chrysogenum]
MRVPWVLINNLLLFIPPVSAGPAAYGVCQAGCAAVVMACYSAAGFTWGATMGASAPASVMACNLAFGKCQAACAAVLLAPTL
**>KAJ5277483.1 hypothetical protein N7524_003636 [Penicillium chrysogenum]**
**MRVPWVLINNLLLFIPPVSAGPAAYGVCQAGCAAVVMTCYSAAGFTWGATMGASAPASVMACNLAFGKCQAACAAVLLAPTL**
>XP_014530676.1 hypothetical protein PDIP_86270 [Penicillium digitatum Pd1]
MKSLWAILYACLFFSSDVSAGPAAYGVCQAGCATVVMAFYSAAGFTWGATMGATVPASILACNPAFGKCQAACASVLLAPTF
>KAJ6031359.1 hypothetical protein N7540_002091 [Penicillium herquei]
MKTSTILSLGSLSMVPTVLAGPIGYGICQAGCSSVVMACYTAGGATWGATLGATAPATIVGCNAAFGTCQAACAALLLTPIP
>KAJ5613654.1 hypothetical protein N7528_007308 [Penicillium herquei]
MKTSTLLSLGGLSLAPTVLAGPIGYGICQAGCSSVVMACYVAGGATWGATLGATAPPTIVGCNVAFGTCQAACAAVLLTPTP
>XP_056945834.1 uncharacterized protein N7483_007656 [Penicillium malachiteum]
MKIAKIIPLTLPIVPAANAGQAGYGICQAGCSAVVRACYAVAGVQWGACCGAQASPAVIACNAAHGTCQAACASVLL
>KAJ5738444.1 hypothetical protein N7493_001599 [Penicillium malachiteum]
MKTSTLLCLGTIFLTPTVLAGPIGYGICQAGCSSVVMACYVAGGATWGAVLGATAPATIVGCNAAFGTCQAACAAVLLKPAP
>XP_056945832.1 uncharacterized protein N7483_007654 [Penicillium malachiteum]
MKASTFLSLGTLSLVPTVLAGPIGYAICQGGCSSVVMACYAAGGATLGATLGATAPATIVGCNVAFGTCQAACAAVLLTPIP
>KAJ5710925.1 hypothetical protein N7488_005081 [Penicillium malachiteum]
MKTSTLLSIVGLSLKPTVLAGPIGYGICQAGCSSVVMACYAAGGATWGATLGATAPPTIIACNVAFGTCQAACAAVLLTPTP
>XP_056989915.1 uncharacterized protein N7511_001757 [Penicillium nucicola]
MRAPWVLIYGLLLFIPFVFAGPAAYGICQAGCAAVVMACYSAAGFTWGATMGASAPASVLVCNSAFGTCQAACAAALLAPTL
>XP_057023110.1 uncharacterized protein N7466_005032 [Penicillium verhagenii]
MRVLPLLIPIYILLSFAAAGPISYGICQAGCAAVVMACYSAAGFTWGATMGATAPASILLCNTAFGKCQAAAAAVLLSPTP
>CDM32893.1 unnamed protein product [Penicillium roqueforti FM164]
MRVSRGKFLNLLLFSPTVLAGPAAYGVCQAGCAAVVMACYSAAGFTWGATMGISAPASIVACNSAFGTCQGACASVLLAPTP
>CDM32632.1 unnamed protein product [Penicillium roqueforti FM164]
MRVPWVLINNLLLFIPPVSAGPAAYGVCQVGCAAVIMACYSAVGFTWGATMGASAPASVMACNLAFGKCQAACAAVLLAPIL
>XP_057021283.1 uncharacterized protein N7466_006287 [Penicillium verhagenii]
MRSHWAQFWLPLLLATNVSAGPAAYGVCQAGCAALVMACYSAAGFTWGVAMGATIPASIVTCNSAFGTCQAACASVLLAPTL
>KAJ5955379.1 hypothetical protein N7501_009658 [Penicillium viridicatum]
MRAPWVLINNLLLFIPPVSAGPAAYGICQAGCSAVVMACYSAAGFTWGATMGVSAPASVIVCNSAFGTCQAACAAALLAPTL
>XP_022582555.1 hypothetical protein ASPZODRAFT_158867 [Penicilliopsis zonata CBS 506.65]
MKILEIGAVFSAVSLVMAGPAAYGICQAGCAAVVTACYSAAGFTWGATLGASAPASILLCNSAFGTCQAACAATLLAPTL
>KAH7066527.1 hypothetical protein FB567DRAFT_458571 [Paraphoma chrysanthemicola]
MACRAFLLVTTTLAFAGPVHAGPIGYGICQAACSAVVMACYSAAGFTWGATLGATAPSSILACNPAFGTCQAACAAVLLAPTP
>KAF9740932.1 hypothetical protein PMIN01_00471 [Paraphaeosphaeria minitans]

*MRLPTMTATAMVALAGTAAAGF*IGYRICQAGCSDVVVACYSGAGFTFGAVAAAAAPPALIACNTAYGRCQAACATALLVPAP
>KAH5618967.1 hypothetical protein HBI23_246000 [Parastagonospora nodorum]
LLETIATTATAILLLTQSALAGPAAYGICQAGCSAVVMACYSAAGFTWGATLGATAPASILVCNAAFGTCQAACAAVLLAPTL
>XP_001803436.1 hypothetical protein SNOG_13225 [Parastagonospora nodorum SN15]
MKLATTTVTTAATAALLVSMVSAGPIGYGICQAGCSAVVMACYSAAGFTWGAVLGATAPATIVVCNTASGTCQAACAAVLLGPTP
>PVH94442.1 hypothetical protein DM02DRAFT_618639 [Periconia macrospinosa]
MHFPSISTVSAGFLALAGTVSAGPAAYGICQAGCAGIVMACYSAAGFTWGATAGAMVPATNIACNASFGKSQGACVLVLL
>CAI6339829.1 unnamed protein product [Periconia digitata]
MRFHYLTPLALLGLAGTTQAGPAAYGICQAGCADVVMACYSGAGFTWGATPGATAPATIIACNAAFGTCQAACAAVIFASTP
>KAI5806420.1 hypothetical protein DFH27DRAFT_651256 [Peziza echinospora]
LPTTTTTALVLLTLTLPTAHAGPLAYGICQAGCAAVVMACYTAAGAVWGVAPPLALAAPLACDSAFGVCQAACWAAVIAPTP
>KAH8691071.1 hypothetical protein GQ44DRAFT_720093 [Phaeosphaeriaceae sp. PMI808]
MKFPSFSIIIIDCLVLARSASAGPAAYGICQAGCSAVVMACYTAAGFTWGATLGASAPATIIGCNAAFGTCQAACAATLLAPTP
>KAK2073227.1 hypothetical protein P8C59_007522 [Phyllachora maydis]
*MQPIKILFPAMLIFGHAAHA*GPVAYGLCQAGCAAVVTACYAAGGATWGVTGGASVPPTIVACDSAFGACQAACWAAVIAPTP
>KAG7005615.1 hypothetical protein G7Y79_00018g044220 [Physcia stellaris]
MHFIKFSTFVAISSLMTTAVAGPVAYGICQAGCATVATACYAAAGATFGTITAGAGTPAVILGCNSAFGVCSAKCAAIALLPTP
>KAF2705155.1 hypothetical protein K504DRAFT_460423 [Pleomassaria siparia CBS 279.74]
MHLYSFVTSTTLLVLASSVSAGPAAYGICQAGCAGVVMACYAAGGSTWGATLGASAPPTIVACNTAYGTCQAACAAIALAPTP
>KAI1006201.1 hypothetical protein K3495_g2025 [Podosphaera aphanis]
MRISLPLLVAISTAAPVIAGPAAYGICQAGCAGVVMACYTAAGFTWGATLGATAPATIIACNAAYGTCQAACAVAIFAPTP
>KAI1811651.1 hypothetical protein GGS20DRAFT_91490 [Poronia punctata]
MKLIATILLANALAVEAGPVGYGICQAGCSGVVMACYGAAGFTWGATLGASAPASIIACNTAFGTCQAACAAVLLTPTP
>TLD26839.1 hypothetical protein PspLS_05170 [Pyricularia sp. CBS 133598]
*MQYSTITKAIVASAILNGGVLA*GPAAYGVCQSGCSAVVMACYAAAGFTWGATLGASAPASIIACNTAFGFCQAGCHAALVMPTP
>CCX30515.1 Similar to predicted protein [Postia placenta Mad-698-R]; acc. no. XP_002475098 [Pyronema omphalodes CBS 100304]
MKVSALTLTVAAMFASSVTAGPLAYAACQAGCATVVMACYTAGGATWGATLGATAPATIIGCNSAYASCQAVCASVALFAPTP
>CAE7186653.1 hypothetical protein PTTW11_07002 [Pyrenophora teres f. teres]
MKLPTLATVATASLIFAHSASAGPIGYGICQAGCSAVVMACYSAAGFTWGATLGATAPASILACNAAYGTCQAACAAVLLGPTP
>KAA8618168.1 hypothetical protein PtrV1_09675 [Pyrenophora tritici-repentis]
MKLPALTAVTTASLFFAHSASAGPIGYGICQAGCSAVVMACYSAAGFTWGATLGATAPASILACNAAYGTCQAACAAVLLSPVP
>KAI7911282.1 zygote-specific protein [Pyricularia oryzae]
*MQHSTIAHAIMASAVLSGVALA*GPAAYGICQAGCSGVVMACYGAAGFTWGATLGASAPASILACNTAFGACQASCHAVLFIPGP
>XP_030984416.1 uncharacterized protein PgNI_03574 [Pyricularia grisea]
*MHHLTIAKAITATAILSGPALA*GPGAYGVCQAGCCAVVMACYAAAGATWGATAGATAPATVVACNTAFGACQASCHAALIFPIP
>XP_029751233.1 hypothetical protein PpBr36_01438 [Pyricularia pennisetigena]
MHYSAIAKVIMATAVLNGLALAGPAAYGVCQAGCCAVVMACYTAAGATWGATAGATAPATVVACNTAFGSCQAACHAALLMPVP
>KAI5802037.1 hypothetical protein FPQ18DRAFT_251069 [Pyronema domesticum]
MKVSALTLTVAAMFASSVTAGPLAYAACQAGCATVVMACYTAGGATWGATLGATAPATIIGCNSAYASCQAVCASVALFALTP
>KAI5789794.1 hypothetical protein FPQ18DRAFT_36823 [Pyronema domesticum]
MKVCTPLTVAAMFASSVTAGPLAYAACQAGCTTVVVACYAAGGATWGATVGATAPATIIACNSAYASCQAVCATVGLFTPTP
>KAI5816642.1 hypothetical protein BZA77DRAFT_293242 [Pyronema omphalodes]
ILFGSNVTAGPLAYAACQAGCATVVMACYSAAGFTWGATLAATAPATVIACNSAYGSCQTVCATIGLLAPTP
>KAI5818345.1 hypothetical protein BZA77DRAFT_306718 [Pyronema omphalodes]
MKVRTSVTVAALLFGSSVTAGPLAYAACQAGCATVVMACYSAAGFTWGATLGATAPATVIACNSAYASCQTVCATVGLFAPTP
>KAH8711905.1 hypothetical protein GQ44DRAFT_689847 [Phaeosphaeriaceae sp. PMI808]
MKLLHSFSLVTGIFLLGQSVSAGPAAYGVCQAGCSAIVVACYSAAGFTWGATLAATAPASILACNSAFGTCQAACAAVLLAATP
>CCX16445.1 Similar to Proteophosphoglycan 5 [Rhodotorula glutinis ATCC 204091]; acc. no. EGU11658 [Pyronema omphalodes CBS 100304]
MKVCTPLTVAAMFASSVTAGPLAYAACQAGCTTVVVACYAAGGATWGATVGATAPATIIACNSAYASCQAVCATKCFEGNVLDLAT
**>CZS94272.1 uncharacterized protein RAG0_04318 [Rhynchosporium agropyri]**
**MNLLIVILVLLAASLSLVVGGPVSYGVCQGGCAAVVMACYAAGGATWGATLGLTATPTIVGCNTAFGVCQASCWVAVLNPLF**
>CZS94274.1 uncharacterized protein RAG0_04319 [Rhynchosporium agropyri]
MRLPSPPLFLLAISPTLALGGPAAYGLCQAGCAAVVTACYAAGGATWGATLGASAPATIIGCNSAFGTCQAACAVVALLPTI
>CZT42857.1 uncharacterized protein RSE6_02809 [Rhynchosporium secalis]
MRLPSPPLFLLAISPTLALGGPAVYGLCQAGCAAVVTACYAAGGATWGATLGASAPATIIGCNSAFGTCQAACAVVALLPTI
>KAH7363888.1 hypothetical protein BKA65DRAFT_489870 [Rhexocercosporidium sp. MPI-PUGE-AT-0058]
MRICIALSLLAAASPALASHLEYSMCQAGCMPATCACYAAPAAVFGTVMTGFASTAVLACNKAQGWCYSNCAAKHLP
>PQE30004.1 hypothetical protein CJF32_00000670 [Rutstroemia sp. NJR-2017a WRK4]
MKLSFTATLLTLAPLLPFTTAGPAAYGVCQSGCAAVVMACYTAAGFTWGATLGATAPATILACNAAFGTCSATCAALLLAPTP
>ESZ95438.1 hypothetical protein SBOR_4179 [Sclerotinia borealis F-4128]
MKPKSSLLIFFFFVSTLMGTATAGPIAYGICQSGCAAVVMACYAAGGATWGATLGATAPATIVACNTAFGVCSAKCWVAAALPFF
>XP_001596882.1 predicted protein [Sclerotinia sclerotiorum 1980 UF-70]
MKPTSTSILLLTAITTFMGMATAGPVAYGVCQSGCAGVVMACYSAAGFTWGATLGATAPPTIIACNTAFGLCSAKCAGFLVAPIP
>KAJ8059346.1 hypothetical protein OCU04_012303 [Sclerotinia nivalis]
MKPTSTSILLLTTLTTFMGMATAGPVAYGVCQSCCAIVVMACYSAAGFTWGATLGATAPPTIIACNAAFGLCCAKCAGFLVAPIP
>CAD6445730.1 e41de491-f048-4506-b0d3-36f1b4dfaa10 [Sclerotinia trifoliorum]
MKPTSILLTTLASFMGMASAGPIAYGACQSGCAAVVMACYLAAGFTWGATLGATAPPTIIACNTAFGLCSAKCAGFLVAPIP
>KAF2024935.1 hypothetical protein EK21DRAFT_77712 [Setomelanomma holmii]
MNRFAITVFSVLALADSGTAGPAAYGICQAGCSAIVMACYSAAGFTWGATMGASAPATILACNAAFGTCQAACAAALLAPTL
>KAA8906278.1 hypothetical protein FN846DRAFT_907066 [Sphaerosporella brunnea]
MKPITLLLALPATVTAGPLAYAACQGGCAAVVMACYGAAGYTWGATLGVAAPATVLACNAAYATCQATCATICLFAPTP
>XP_016763988.1 uncharacterized protein SEPMUDRAFT_114931 [Sphaerulina musiva SO2202]
MRFTTLTLATLATLILPSLAGPAAYGICQAGCSAVVMACYAAGGATWGAALGATAPPTIVACNTAYGICYAACHAAIFAPTP
>KEY69198.1 hypothetical protein S7711_01656 [Stachybotrys chartarum IBT 7711]
MKLILGLRGAFFLLFAGSAAAGPVAYGICQAGCASVVMACYTAGGATWGATAGATAPATIVGCNTAFGSCQAACAYIALAPTP
>KAF7871723.1 hypothetical protein EAF04_003830 [Stromatinia cepivora]
MKPTSTSILLLTTTFIGIATAGPVAYGVCQSGCAGVVMACYSAAGFTWGATLGATAPPTIIACNTAFGFCSAKCAGFLVAPIP
>KAI4247966.1 MAG: hypothetical protein LQ352_006032 [Teloschistes flavicans]
*MHPLKSLPLLALIPLVTA*GPAAYGICQAGCSAVVVACYSGAGFTFGTVTAGAGIPAAVVACNSAYGTCQAACAAILLAPTP
>MCJ1226585.1 hypothetical protein [Toensbergia leucococca]

MIPTTLLAILTLPLLLLPTAVTAGPLAYATCQAGCSAVVMACYAAGGATWGATLGATAPATIVGCNSAFGLCQAGCAAALLLPMP
>XP_051335166.1 uncharacterized protein BZA05DRAFT_449074 [Tricharina praecox]
MKLSLVPLLALLATSASAGPAAYGVCQAGCACVVMACYAAGGATWGATLALTAPATIIGCNGAYATCQSACALVVFAPTL
>UKZ60045.1 hypothetical protein TrAtP1_001332 [Trichoderma atroviride]
MKLTNITLAAIVLPGTAMAGPLAYAACQAACSTTLAAGPAGVALYAACQSACAPLLVMPCP
>XP_024774938.1 hypothetical protein M431DRAFT_495476 [Trichoderma harzianum CBS 226.95]
MHIFKVLSISTRVATSNAGPIAYRICQAGCAGVVMACYGAAGATWGATAAASAPATVIACNRTFGVCQAACWITATLPYF
>XP_024774939.1 hypothetical protein M431DRAFT_508663 [Trichoderma harzianum CBS 226.95]
MQIVKVLAILTLAATSNAGPVAYGICQAGCAAVVTACYAAGGATWGATAGATAGPTIIGCNSAFGSCQAACWAAATFPCP
**>KAF3063233.1 hypothetical protein CFAM422_010070 [Trichoderma lentiforme]**
**MRITKVVTISALAATSNAGPVAYGICQAGCATVVTACYAAGGATWGATAGATAGPTILACNSAFGSCQAACWAAAIFPCP**
>PTB76422.1 hypothetical protein M440DRAFT_1401876 [Trichoderma longibrachiatum ATCC 18648]
MKMKFVNIASSVIVLSGIAMAGPVAYATCQAACAVSLAAPGGVAIYAACQSHCAALLVAPCP
>QYS96969.1 hypothetical protein H0G86_004206 [Trichoderma simmonsii]
MQIVEVLAILTLATTSNAGPVAYGICQAGCAAVVTACYAAGGATWGATAGATASPTIIGCNTAFGSCQAACYAAATFPCP
>XP_013959331.1 hypothetical protein TRIVIDRAFT_31923 [Trichoderma virens Gv29-8]
MRILKVLAISTLVTTSAAGPIAYRICQAGCAGVVMACYAAAGATWGATAAASAPVTVLACNSAFGTCQAAYWVAATLPCF
>UKZ75591.1 hypothetical protein TrVFT333_003279 [Trichoderma virens FT-333]
MRILKALAISTLVTTSAAGPIAYRICQAGCAGVVMACYAAAGATWGATAAASAPVTVLACNSAFGTCQAAYWVAATLPCF
>XP_040731585.1 uncharacterized protein BHQ10_003081 [Talaromyces amestolkiae]
MKLLSAALAALLLTTSVTAGPAAYGVCQTGCAAVVMACYSAAGFTWGATLGATAPASIIACNTAYGTCQAACAAVLLTPTL
>KAF8434007.1 cysteine-rich protein [Terfezia claveryi]
MRFSLIIPAVAAIFGTASAGPLAYGICQAGCSALVVSCYTAGGLTFGTITAGAGAPAIALACNASYGVCQAACAAAILAPTP
>KAF8424820.1 hypothetical protein EV426DRAFT_532606 [Tirmania nivea]
MRSSFIFPLVAMLGTASAGPLAYGICQAGCSTLVVSCYAAGGLTFGTVTAGLGAPAVALACNASYGACQAACAAAILAPTP
>KAF2434089.1 hypothetical protein EJ08DRAFT_582745 [Tothia fuscella]
MRVIAILAAAFVGMTAAGPIAYGVCQAGCSAVVMACYTAGGATWGATAGATAPATIIGCNSAFGTCQAACAVAFLLPTL
>PWW74214.1 hypothetical protein C7212DRAFT_210569 [Tuber magnatum]
MKLQNLLLLPTSFAAVALAGPISYGICQGGCAAVAVACYSGAGFIFGTVPATAAAAIPAVLACNSAFGTCSATCAGVTLLAPIP
>KAG0129985.1 hypothetical protein HOY82DRAFT_563331 [Tuber indicum]
MKLQNLLLPMSFATVALAGPISYGICQSGCAAIVCVCYSAAGAVFGTVPAAAAAAMPAIVACNSAFGTCSATCAGVTLLAPIP
>KAG0639743.1 hypothetical protein HOY80DRAFT_961797 [Tuber brumale]
MKLQNLLLPMSFAAVALAGPISYGICQGGCAAVACACYSAAGVVFGTVPGAVAAAMPAIAACNGAFGTCSATCAGVTLLAPIP
>ABO93224.1 hypothetical protein Tbz1 [Tuber borchii]
MKLQTLLLPASFAAVALAGPISYGICQSGCAAVVVACYTGAGAVFGTVPAAAAAAMPAIAACNGAFGTCSATCATITLLAPIP
>XP_002839789.1 uncharacterized protein GSTUM_00007935001 [Tuber melanosporum]
MKLQNLLLPMSFATVALAGPISYGICQSGCAAVVCVCYSAAGAVFGTVPAAAAAAMPALAACNGAFGTCSATCAGVTLLAPIP
>XP_043001095.1 uncharacterized protein UV8b_07663 [Ustilaginoidea virens]
RHRTFATVAVFVPCVVAGPALYGVCQAGCAAVVMACYSAAGATWGATAGITAPASVLACNAAFGKCSLACYIAAGAPTP
>ROV87140.1 hypothetical protein VSDG_09989 [Valsa sordida]
MRPIKILAILAMATGTTAGPIAYGICQAGCAAVVTACYAAAGATFGTVAAPAAPAAIVACNSAFGTCQAACAVVALAPTP
>KUI64379.1 hypothetical protein VM1G_11180 [Valsa mali]
MQPIKMLAVLTMATTATAGPIGYGICQAGCSAVVTACYAAAGVTFGTIAALAAPAAIVGCNTAFGTCQAACAAVLLTPTP
>KUI62490.1 hypothetical protein VP1G_09610 [Valsa mali var. pyri (nom. inval.)]
MQPIKMLAVLTMATTATAGPIAYGICQAGCSAVVTACYAAAGATFGTVAAPAAPAAIVGCNTAFGTCQAACAAVLLTPTP
>QDS77261.1 hypothetical protein FKW77_003660 [Venturia effusa]
MHPSLLLTITALATTTTAGPLGYGICQAGCSSIVMACYAAAGFTWGATLGASAPATIVACNAAYGTCQAACASVLLTPTP
>KAE9964269.1 hypothetical protein EG328_010656 [Venturia inaequalis]
MRTSILITAAALAATISAGPLGYAVCQAGCASVVMACYAAGGATWGATLGATAPATIVACNAAYGTCQAACAAVLLIPFL
>XP_031874101.1 Uncharacterized protein BP5553_01424 [Venustampulla echinocandica]
MRIFTPAFALAVLCLLDSATAGPVGYGVCQAGCAAVVTACYGAAGFTWGATLGATAPASIIACNAAFGTCQAACAAVLLTPTP
>KAF3353751.1 L-rhamnonate dehydratase [Verticillium dahliae VDG1]
MRPTKIVQLTALAAVPASAVGPVAYGICQAGCASVVIACYGAAGFTWGATLGATAPASVLACNAAFGTCCAACAAVPLTPTP
>XP_003008200.1 conserved hypothetical protein [Verticillium alfalfae VaMs.102]
MRPTKIVQLAALAVVPASAGLVAYGICQAGCASVVTACYGAAGFTWGATLGATAPASVLACNAAFGTCCAACAAVALTPTP
>XP_028490889.1 uncharacterized protein D7B24_002946 [Verticillium nonalfalfae]
MHPTKIVQLAALAAVPASAGLVAYGICQAGCASVVTACYGAAGFIWGATLGATAPASVLACNAAFGTCCAACAAVALTPTP
>KAG7104944.1 hypothetical protein HYQ44_016256 [Verticillium longisporum]
MRPTNIFQLAALTAVPASAGLVAYGICQAGCASVVTACYGAAGFTWGATLGATAPASVLACNAAFGTCCAACAAVALTPTP
**>PNH30441.1 hypothetical protein BJF96_g6371 [Verticillium dahliae]**
**MRPTKIVQLTALVVVPASAGPVAYGICQAGCAGVVTACYGAAGFTWGATLGATAPASVLACNAAFGTCCAACAAVALTPTP**
>XP_009651807.1 uncharacterized protein VDAG_02859 [Verticillium dahliae VdLs.17]
MRPTKIVQLTALAVVPASAGPVAYGICQAGCAGVVTACYGAAGFTWGATLGATAPASVLACNAAFGTCCAACAAVALTPTP
>KAG7104378.1 hypothetical protein HYQ44_015690 [Verticillium longisporum]
MRPPKIVQLTALAAIPASAGPVAYGICQAGCASVVTACYGAAGFTWGATLGATAPASVLACNAAFGTCCAACAAVALTPTP
>KAF8244270.1 hypothetical protein K440DRAFT_610160 [Wilcoxina mikolae CBS 423.85]
MKLSLIVTAALLSASGAHAGPLAYAACQAGCAGVVMACYSAAGATWGATLGATAPATVLGCNAAYASCQGVCATVALCAPIP
>KAI1338517.1 hypothetical protein F5Y15DRAFT_386103 [Xylariaceae sp. FL0016]
MKLNLTPLSPVLLALASVTSAGPIGYGICQAGCASVVTACYGAAGATWGATAGATAPATVLACNSAFGTCQAACAAVLLTPTP
>KAI4258584.1 MAG: hypothetical protein L6R42_005003 [Xanthoria sp. 1 TBL-2021]
MKPTILLLPLTTLSALTSASPAAYGICQAGCSALVIACYSGAGFTFGTVTAGAAIPAAIVACNSAFGTCQAACAAVMIAPTP
>KAI0817434.1 hypothetical protein GGR55DRAFT_620392 [Xylaria sp. FL0064]
MRLTTTFAPVLLGFASIASAGPAAYAACQTGCAGLVTACYAAAGFTWGATLGATAPASILACNSAFGTCQAACWVAAIAPTP
>KAI0459375.1 hypothetical protein F5B21DRAFT_499555 [Xylaria acuta]
*MKPTTPFVTAILASTPIVSA*GPAAYGICQTGCAAVVTACYAAAGFTWGATAGISAPATIVACNSAFGTCQAACWVAAIAPTP
>KAI0972575.1 hypothetical protein F4678DRAFT_460143 [Xylaria arbuscula]
*MRPTTSLVATLIGVAPIVSA*GPAAYGICQAGCAAVVTACYAAAGFTWGATMGASAPATVIGCNTAFGACQAACWAAAIAPTP
>KAI4224460.1 MAG: hypothetical protein LQ349_007235 [Xanthoria aureola]
MKLTNLLTTFASLTTLTSAGPAGYGVCQAGCSAVVVACYSGAGFTFGTVTAGAAVPAAIVACNSAYGTCQAACAAVLIAPTP
>KAI1756746.1 hypothetical protein F4782DRAFT_526421 [Xylaria castorea]
MKPTTPFVTAILASTSIVSAGPAAYGICQAGCAAVVTACYAAGGFTWGVTAGLSIPATIIACNTAFGTCQAACWVAAIAPTP
>KAI0859552.1 hypothetical protein F4860DRAFT_249266 [Xylaria cubensis]

*MKPTTSLFAALLASTPIVSA*GPAAYGICQAGCAAVVTACYAAAGFTWGATAGLSIPATIVACNTGFGTCQAACCAMTG
>KAI0555338.1 hypothetical protein F4679DRAFT_578725 [Xylaria curta]
MKTTTALVTALLASTSIVSAGPAAYGICQAGCAAVVTACYAAAGFTWGATVGVSAPATIIACNSAFGTCEAACWAAAIAPTP
>TRX93855.1 hypothetical protein FHL15_005237 [Xylaria flabelliformis]
MKPTTSLFAAILASTPIVSAGPAAYGICQAGCAAVVTACYAAAGFTWGATAGISAPATIVACNTASGTCQAACPFFFSAQEG
>KAI0486197.1 hypothetical protein F4859DRAFT_511198 [Xylaria cf. heliscus]
*MKPTALLATSILALAPLASA*GPAAYGICQAGCAGVVMACYGAAGFTWGATAGLTLPASVIACNTTFGACQAACWAAAIAPTP
>KAI8947707.1 hypothetical protein F4801DRAFT_559745 [Xylaria longipes]
*MKPSTPFVTALLTFTPIVSA*GPASYSICQAGCAAVVKACYAAAGFTWGATLGISAPASIVACNAAFGTCQASCWVAAIVPTP
**>KAI0404163.1 hypothetical protein F4802DRAFT_568168 [Xylaria palmicola]**
**MKVTTPLISAVLAFSPIASAGPLAYGLCQAGCAAVVTACYSAAGFTWGATMGASAPATVVACNAAFGTCQAACWAAATAPTL**
>KAH8159225.1 hypothetical protein CIB48_g9022 [Xylaria polymorpha]
MKLTIPFLTATLAPIVSAGPAAYGICQTGCAAVVVACYAAAGFTWGATAGISAPATILACNSAFGTCQASCWVAAIAPTP
>KAI1736724.1 hypothetical protein F4680DRAFT_246167 [Xylaria scruposa]
MKATTALVTSLLASTPIVSAGPAAYGICQAGCAAVVTACYAAAGFTWGATAGVSAPATIIACNSAFGTCQAACYVAAFAPTP
>KAI4226682.1 MAG: hypothetical protein L6R36_002995 [Xanthoria steineri]
MKLAKLLTTLATLTAVTSAGPAGYGVCQAGCSAVVVACYSGAGFTFGTVTAGAAVPAAIVACNSAFGTCQAACAAVLIAPTP
>KAI0447941.1 hypothetical protein F4803DRAFT_498254 [Xylaria telfairii]
MNLTTSFLTATLALAPIVSAGPAAYGICQAGCAAVVTACYAGAGFTWGATAGISVPATIIACNSAFGTCQASCWVAAIAPTP
>KAI1312583.1 hypothetical protein F5Y03DRAFT_340889 [Xylaria venustula]
MRPATSLIAALIGFAPTVSAGPAAYGLCQTGCAAVVTACYAAAGYTWGATLGASAPATILACNTAFGTCQAACWAAAIAPTP
>KAF2175953.1 hypothetical protein K469DRAFT_608836 [Zopfia rhizophila CBS 207.26]
MQIKDVLAAGLFALPIVVAGPLAYGVCQAGCSTVVVACYAAAGATFGTIAAAAAPPAIIGCNTAYGACQATCASVLLAPTP
>XP_003847476.1 uncharacterized protein MYCGRDRAFT_51607 [Zymoseptoria tritici IPO323]
MSFTFITFSGLVFMAAIAQAGPVGYGLCQAGCSAVVMACYTAGGATWGATAGATAPATIIGCNSAFGTCQAACAAVIFAPTP
>SMQ45019.1 unnamed protein product [Zymoseptoria tritici ST99CH_3D7]
MRLLKMSLATLAVIAPAISAGPAAYGVCQAGCAGVVVACYAAGGFVFGAAAPPVLPPVIVACNTAFGACQAACWAALMMPTP
>SMR41375.1 unnamed protein product [Zymoseptoria tritici ST99CH_1E4]
*MRLLKMSLATLAVIAPAISA*GPAAYGVCQGGCAKVVKACYAAAGFIWGSAFDGPVPATVQACNGAFGICQGACWTALLMPVP

```
RPA97680.1        -------------MKLRNILLPASFTTMALAGPISYG--ICQSGCAGVVVACYSAAGAVFGTV--PAAAAAAIPALAACNSAFGTCSHVCATVALLAPIP----
RPA97681.1        -------------MKLQNILLPASFATMALAGPISYG--ICQGGCAGVVVACYSAAGAVFGTV--PAAAAVAIPALAACNSAFGSCSAVCATVALFAPIP----
RPA97678.1        -------------MKLQNLLLPASFATMALAGPISYG--ICQSGCAGVVVACYSAAGAVFGTV--PAAAATAIPALAACNSAFGSCSHFCASVTLLALTP----
KAG0129985.1      -------------MKLQNLLLPMSFATVALAGPISYG--ICQSGCAAIVCVCYSAAGAVFGTV--PAAAAAAMPAIVACNSAFGTCSATCAGVTLLAPIP----
XP_002839789.1    -------------MKLQNLLLPMSFATVALAGPISYG--ICQSGCAAVVCVCYSAAGAVFGTV--PAAAAAAMPALAACNGAFGTCSATCAGVTLLAPIP----
KAG0639743.1      -------------MKLQNLLLPMSFAAVALAGPISYG--ICQGGCAAVACACYSAAGVVFGTV--PGAVAAAMPAIAACNGAFGTCSATCAGVTLLAPIP----
ABO93224.1        -------------MKLQTLLLPASFAAVALAGPISYG--ICQSGCAAVVVACYTGAGAVFGTV--PAAAAAAMPAIAACNGAFGTCSATCATITLLAPIP----
PWW74214.1        ------------MKLQNLLLLPTSFAAVALAGPISYG--ICQGGCAAVAVACYSGAGFIFGTV--PATAAAAIPAVLACNSAFGTCSATCAGVTLLAPIP----
POS77299.1        -------------MQPTKILITAMSLGTALAAEVAYG--VCQAICTRQALTCYDKAGVTFGSK--HQLGTEFN--ITTCNVALDTCVRICIFVANGA-------
XP_044637434.1    -------------MQPTGILITAMTVATALAAELTYG--VCQAICTRQALTCYDDAGVTVGSK--NAYGSAAN--VTACNVTLDTCGLVCAMVASGGCDSSECC
KAH7363888.1      -------------MRICIALSLLAAASPALASHLEYS--MCQAGCMPATCACYAAPAAVFGTV--MTGFASTA--VLACNKAQGWCYSNCAAKHLP--------
TVY53193.1        ------------MKISRIIAALATTTTLVTAGPAAYG--LCQTGCAGVVVACYSAAGFTFGT----VAAAAAPAAILACDAAYGSCCAACYWALFLPTP-----
TVY92812.1        ------------MKISRIIAALATTTTLVAAGPAAYG--LCQTGCVGVAVACYSAAGFTFGT----VAAATAPAAIVACNSAYGSCCATCHALLFAPTP-----
TGZ81582.1        ---------MTPPYLLFLFTLLLLFTTTVTAGPATVG--ACQAGCAAVVVACYAAAGFVFGT----IMPPSVPPAILACNAAQGTCYAGCYALFFMPTP-----
XP_025569355.1    ------------MNLLGAGSGLLVAASNVFAGPSAYA--VCQTGCSALVVSCYAAAGFVFGM----VPAVSAPPAIIACNSSYGTCQAACASILLVPFP-----
XP_025468364.1    ------------MNLLRAVINLLIAGSVVVGGPAAYG--VCQAGCSSVVVSCYAAAGFVFGM----VPAVAAPPAIAGCNSAYGTCQAACATILLAPFP-----
OOF99516.1        ------------MRILQAGVSLLIAANGEFRSPAAYG--TCQAGCSSVVVSCYAAAGFVFGM----VPAVSAPSAIVGCNSAYGNCQAACALVLLAPVL-----
KAF2175953.1      ------------MQIKDVLAAGLFALPIVVAGPLAYG--VCQAGCSTVVVACYAAAGATFGT----IAAAAAPPAIIGCNTAYGACQATCASVLLAPTP-----
KAH8743362.1      ------------MQLLKSLVATLFLASSVAAGPIAYA--ICQAGCSSLAVACYSAAGFTFGT----VAAAAAPAALLACNSAYGTCQPACVVAALVPTLGKRS-
KAI3393011.1      --------------LHESLVATLFLASSVAAGPIAYA--VCQAGCSSLAVACYSAAGFTFGT----IAAAAAPAALIACNSAYGTCQAACAAAALAPTL-----
KAH8747636.1      ------------MRLQKSLVANLFLASSVAAGPVVYA--VCQAGCSSLAVACYSAAGFTLGI----IATTAAPAAVIACKTAYGTCQAACAAAALAATP-----
KAF1972950.1      ----------MRFHTIAATAMVALAG-TTAAGPIGYG--ICQAGCSAVVVACYSAAGFTFGT----VAAAAAPAAIAACNTAYGTCQAACAAVLLSPTL-----
KAF9740932.1      ----------MRLPTMTATAMVALAG-TAAAGFIGYR--ICQAGCSDVVVACYSGAGFTFGA----VAAAAAPPALIACNTAYGRCQAACATALLVPAP-----
KAF2447884.1      ----------MRFHTIAATAVVGLLVGTAAAGPIGYG--ICQTGCAAIVVACYSAAGFTFGT----VLAAAAPPAILACNSAYGTCQAACAVVLLGPTP-----
RVD82005.1        -------------MKPSLIVSACTVISLVSAGPLAYG--VCQAGCAAVVTACYGAAGFVWGA----TLATAAPPAIIACLTMSALGGRRRVR------------
XP_031903430.1    -----------MKILYPAVGIFSTLVNSVYAGPEAYR--ICQAGYAAVVTACYSAAGFTWGA----TLTLPEARLA----------------------------
XP_014556701.1    --------MKPFSLIKATTVVLSLTVRQASASRIEYA--VCQAGCASLVMACYTAAGFVWGT----VGRDTASQAILFCNAAFGKCSAACAEATL---------
XP_007712361.1    --------MKPFSLIKATTVVLSLTVRQASAGRIEYA--VCQAGCASLVMACYTAAGFVWGT----VGRDTASQAILFCNAAFGKCSAACAEATL---------
XP_008028533.1    ---------MNLPSLNNTAVAILILSSQTSAGPLGYA--VCQAGCASVVMACYSAAGYVWGN----TLSADAPDAVIGCNLAFGKCQAACAEVTLGTAM-----
KAF1987320.1      ----------MRPRAILIASLLLAAATSVSAGPASYG--ICQAGCASVVMACYAAAGANWGA----VLAFGAPSAVATCNAAFGPCQSACHVVACIPFL-----
XP_051335166.1    -------------MKLSLVPLLALLATSASAGPAAYG--VCQAGCACVVMACYAAGGATWGA----TLALTAPATIIGCNGAYATCQSACALVVFAPTL-----
KAK0100245.1      -------MKMRLSNPSAFLLFMATRPALIHAGPVGYG--ICKAGCASIVTACYAAAGATWGA----TLGATAPATVVACNTAFGICQGKCAVVALLPTP-----
PVH85672.1        -----------MRFPTSAIILLATTPTLIIGGPVAYG--ICQAGCATVVTACYVAAGATWGA----TLGATAPATVVGCNTAFGICQAKCAIVALLPTP-----
CZS94274.1        -----------MRLPSPPLFLLAISPTLALGGPAAYG--LCQAGCAAVVTACYAAGGATWGA----TLGASAPATIIGCNSAFGTCQAACAVVALLPTI-----
CZT42857.1        -----------MRLPSPPLFLLAISPTLALGGPAVYG--LCQAGCAAVVTACYAAGGATWGA----TLGASAPATIIGCNSAFGTCQAACAVVALLPTI-----
KAH6709034.1      -------KHFKICLPNPFLILLATSPTLILGGPLAYG--ICQAGCAAVMTACYAAGGATWGA----TLGSNAPATIVACNSAFGTCQAACAVVALMHTL-----
KAF0316341.1      ----------------------MVAPAVVLAEKCAYG--TCQTGCAALVVTCYTVAGGIFGV----TSGVAATTAVKECDVAFGKCQASCSRARSAPRC-----
KAK1845575.1      ----------MKPLTAILSTCSILAPAVVLAEKCAYG--TCQTGCAALVVTCYTVAGGIFGV----TSGVAATPAVKECDVAFGKCQASCSQARFAPRC-----
KAK2770104.1      ----------MKLSMATLSTLSILAPTTVLAGLCAYG--TCQAGCAALAVVCYSGTGATFGV----TCGLAATPAVLACNAAFGQCQASCWLAGFAPTC-----
KAH9242425.1      ----------MKFSAAILSTIYILAPATVLAGPCAYG--TCQAGCAVLTVACYSGTGATFGV----TCGLAATPAVLACNAAFGQCQASCWLAGLAPTC-----
XP_031884118.1    ----------MKFSTATLPTIYILAPATVLVGPCAYG--TCQAGCAILAVACYFGTGATFGV----TCGLAATPAVLVCNTAFGKCQASCWLAALAPTC-----
XP_056817142.1    ------------MVSSMAMLHLVLTLAPYVPGWIEYG--KCQAGCATLVMACYAGEGAAWGD----TLGKTAPPPVKTCNSNFGDCQAECAQRWLWPTS-----
KAK2628309.1      -------------MLLPAILVLFAAPAVVFSGPIAYG--TCQGGCAAVVMACYTAAGATWGA----TLGLNAPATVLGCNSAFGYCQAKCAVVALLPTF-----
PBP25780.1        -------------MRLLVILVLFVAPAVVFSGPVAYG--TCQSGCAAVVMACYTAAGATWGT----TLGLTAPTTVLGCNTAFGYCQAKCAVVALLPTL-----
KAI6712870.1      -------------MRLPIISVLFAAPAVVFSGPLAYG--VCQGGCASLVMACYTAAGATWGA----TAGILAPATVVACNTAFGYCQAKCAVVAWLPTL-----
CZS94272.1        -----------MNLLIVILVLLAASLSLVVGGPVSYG--VCQGGCAAVVMACYAAGGATWGA----TLGLTATPTIVGCNTAFGVCQASCWVAVLNPLF-----
XP_023457676.1    -------MRIS--NLKTCLEAATLLITTAHAGPAAYG--ICQAGCAAVVVACYSAAGAVFGV----AAPPAAVPAVLACNSAFGTCQAACWAALMAPTP-----
KAF2207598.1      -------MRVSNVTFKTLFLAAAYLHTTAHAGPAAYG--ICQAGCSAVVAACYAAAGAVFGT----VAAPAAPAAILGCNSAYGTCQAACWASLFAPTP-----
KAF2663912.1      ------------MRLSIPLFAAALLAPSAFAGPAAYG--ICQAGCAAVVAACYAAAGATFG-----VAPPVAIPALAACNTAFGTCQAACWAALLLPTP-----
KAI5806420.1      ----------LPTTTTTALVLLTLTLPTAHAGPLAYG--ICQAGCAAVVMACYTAAGAVWG-----VAPPLALAAPLACDSAFGVCQAACWAAVIAPTP-----
XP_047765020.1    ----------MFLPILPILIALVALLPTAIGGPAAYG--ICQAGCASVVVACYAAAGFVFG-----VALPAAPPAILACNSAFGTCQAACAVVVFAPTV-----
XP_037161484.1    -------------MRLTKLIPMTTIITSALAGPAAYG--ICQAGCAGVAVACYAAAGAIMGV----TAGAAAPPAVLACNAAFGSCQAACAAALVMPTP-----
KAH9242424.1      ------------------LSVFLAAPPAVEAGPLGYG--ICQAGCAGVVTACYAAAGAVWGA----TAGAAAAPAVITCNVAFGKCQAACAVVALTPIP-----
XP_045270162.1    --------------VLLLLSVFLAAPPAVNTGPLSYS--ICQAGCAGVVTACYAAARAVWGA----TAGVAAAPAVIACHVAFRKCQAAYAVVALAPIP-----
XP_031884111.1    ----------MVVIILLLLSIFLAAPPAVEAGPLGYG--ICQAGCAGVVAACYAAAGAVWGA----TAGAAAAPAVIVCNLAFGKCQAACAAVALVPTP-----
KAF0316342.1      ----------MDLSTILVLYVFLAAPPAVEAGLLGYG--ICQAGCAGVVTACYAAAGAVWGA----TAGVGAAPAVIACNVAFGKCQAACAVVALAPTP-----
KAF6819823.1      -----------MNPTHALATVALATIPPTEAGPIGYG--VCQAGCAGVVTACYAAAAAVWSA----TAGAAAAPAVIACNVAFGKCQAACAAVALLPTP-----
KAF6802731.1      -----------MNPTQALVTIALVTIPTATAGPVGYG--ICQAGCAAVVVACYAGAGATFGV----TAGVATGPAVIACNAAFGQCQAA---------------
TDZ35452.1        -----------MNPSGFITLFTLAALPVAVAGPAAYG--VCQAGCASIVVACYAAAGAVFGA----TAGAAAPPAVVACNVAFGKCQCACAMSAICPIP-----
TDZ41237.1        -----------MNPSGFITLFTLVALPVAVAGPAAYG--VCQAGCASIVVACYAAAGAVFGA----IAGAAAPPAVVACNVAFGKCQCACAMSAICPIP-----
TEA19704.1        -----------MNPSGFITLLTLAALPVAVAGPAAYG--VCQAGCASIVVACYAAAGAVFGA----TAGAAAPPAVVACNIAFGKCQAACAMSAICPIP-----
TQW06488.1        --------------MRDHKIIFPFLVGTAAAGPLAYG--ICQAGCAGVVMACYGAAGATWGA----TAGASAAPTVIACNLAFGKCQAACVVAGCSPTP-----
TQV90198.1        --------------MRDQKIILPLLVGLATAGPLAYG--VCQAGCSAVVMACYAAAGATWGA----TAGATAAPAVVACNLAFGKCQAACSVAGFCPSV-----
XP_013959331.1    -------------MRILKVLAISTLVTTSAAGPIAYR--ICQAGCAGVVMACYAAAGATWGA----TAAASAPVTVLACNSAFGTCQAAYWVAATLPCF-----
UKZ75591.1        -------------MRILKALAISTLVTTSAAGPIAYR--ICQAGCAGVVMACYAAAGATWGA----TAAASAPVTVLACNSAFGTCQAAYWVAATLPCF-----
XP_024774938.1    -------------MHIFKVLSISTRVATSNAGPIAYR--ICQAGCAGVVMACYGAAGATWGA----TAAASAPATVIACNRTFGVCQAACWITATLPYF-----
XP_024774939.1    -------------MQIVKVLAILTLAATSNAGPVAYG--ICQAGCAAVVTACYAAGGATWGA----TAGATAGPTIIGCNSAFGSCQAACWAAATFPCP-----
QYS96969.1        -------------MQIVEVLAILTLATTSNAGPVAYG--ICQAGCAAVVTACYAAGGATWGA----TAGATASPTIIGCNTAFGSCQAACYAAATFPCP-----
KAF3063233.1      -------------MRITKVVTISALAATSNAGPVAYG--ICQAGCATVVTACYAAGGATWGA----TAGATAGPTILACNSAFGSCQAACWAAAIFPCP-----
RPA84721.1        ------------MKLSAIFVFSALAAPALS-GPVGYG--VCQAGCSAVVMACYSAAGAVWGA----TAGLGAAPAVLACNAAYGTCQSACAAVLLAPTL-----
XP_056945834.1    ------------MKIAKIIPLTLPIVPAANAGQAGYG--ICQAGCSAVVRACYAVAGVQWGA----CCGAQASPAVIACNAAHGTCQAACASVLL---------
KAK1955786.1      ----------------------------VLGGPGAYG--VCQAGCSAVVMACYTAAGATWGA----TLGLTAAPSVIGCNVAYGTCQAACASVLLAPIP-----
KAK2015154.1      -------------------LLLIPCLPGVLGGPGAYG--VCQAGCSAVVMACYTAAGATWGA----ALGLTAAPSVIGCNVAYGTCQAACSSVLLAPIP-----
KAK1997917.1      -----------MNPATGFLLLLIPALPGVQAGPAAYG--ICQAGCSAVVMACYAAAGATWGA----TLGLTAAPSVIACNVAYGTCQAACAAVILAPTP-----
XP_008099621.1    -----------MTPARSFTLLVIFSLPCVLAGPAAYG--VCQRGCSSVVMACYKAAGSTWGA----TLGLTATPLVIGCNSAYGTCQAACAAVVFTLTP-----
XP_060407788.1    -----------MKTARSFTLLVTSFLPCVLAGPAAYG--VCQAGCSAVVMACYAAAGATWGA----TLGLTAAPTVIGCNLAYGTCQAACAAVILAPTP-----
KAK2042028.1      -----------MKPASNFLFLVTSCLPCALAGPAAYG--ICQAGCSAVVMACYTAGGATWGA----TLGAPAGPTIIGCNLAYGTCQAACASVILAPTL-----
KAK2031093.1      -----------MKPASNFLLLVTSCLPFVLAGPAAYG--ICQAGCSAVVMACYTAGGATWGA----TLGATAGPTIVGCNLAYGTCQAACAAVILAPTL-----
KAK1977458.1      --------MKWSVRVVTHILWLLSSSLCVLAGPLGYG--VCQAGCSAVVMACYTAAGATWGA----TLGLTASESVISYNLAFGKCQAACAAVPLSPTP-----
KAF2004078.1      -------------MRVIPLFTLFAEISVVTAGPAAYG--ICQAGCAAVVTACYAAGGATWGA----TLGATAGPTIVACNTAFGTCQAACWAALIALTP-----
XP_016763988.1    -----------MRFTTLTLATLATLILPSLAGPAAYG--ICQAGCSAVVMACYAAGGATWGA----ALGATAPPTIVACNTAYGICYAACHAAIFAPTP-----
KAF2705155.1      ----------MHLYSFVTSTTLLVLASSVSAGPAAYG--ICQAGCAGVVMACYAAGGSTWGA----TLGASAPPTIVACNTAYGTCQAACAAIALAPTP-----
KAI1485417.1      ---------------AATVLAALLAPAAVSAGPAAYG--ICQAGCSAVAMACYAAGGATWGA----TLGATAPATIVGCNTAFGACQAACWASLIAPTP-----
KAI1630725.1      ---------MKLSSAIPAAAVLAVLAPAVSAGPAAYG--ICQAGCSAVAMACYAAGGATWGA----TLGATAPATIVGCNTAFGVCQAACWAALVAPTP-----
KAI0600113.1      ----------MKLSTAVPIAIIAALAPAANAGPAAYG--ICQAGCSAVVTACYAAGGATWGA----TLGATAPATIVGCNSAFGACQAACWASLLAPTL-----
KAI1503117.1      ----------MKPTTAISVAIASALAPAVSAGPAAYG--VCQAGCSAVVMACYAAGGATWGA----TLGATAPATIIGCNTAFGSCQAACAAVLIVPTP-----
MCJ1226585.1      --------MIPTTLLAILTLPLLLLPTAVTAGPLAYA--TCQAGCSAVVMACYAAGGATWGA----TLGATAPATIVGCNSAFGLCQAGCAAALLLPMP-----
KAF2434089.1      --------------MRVIAILAAAFVGMTAAGPIAYG--VCQAGCSAVVMACYTAGGATWGA----TAGATAPATIIGCNSAFGTCQAACAVAFLLPTL-----
XP_003847476.1    -----------MSFTFITFSGLVFMAAIAQAGPVGYG--LCQAGCSAVVMACYTAGGATWGA----TAGATAPATIIGCNSAFGTCQAACAAVIFAPTP-----
KEY69198.1        ----------MKLILGLRGAFFLLFAGSAAAGPVAYG--ICQAGCASVVMACYTAGGATWGA----TAGATAPATIVGCNTAFGSCQAACAYIALAPTP-----
KAH6664969.1      -------------MRPTILLTPLLLTPALA-GPALYG--ICQAGCASVVMACYGAAGFTWGA----TLGASAPATVLACNAAFGTCSAACAAAVLAPTL-----
KAH8653200.1      -----------------------LASPALA-GPALYG--ICQAGCATVVMACYTAAGFTWGA----TLGASAPPTILACNAAFGTCSAACAAVILAPTP-----
XP_003008200.1    -----------MRPTKIVQLAALAVVPASA-GLVAYG--ICQAGCASVVTACYGAAGFTWGA----TLGATAPASVLACNAAFGTCCAACAAVALTPTP-----
XP_028490889.1    -----------MHPTKIVQLAALAAVPASA-GLVAYG--ICQAGCASVVTACYGAAGFIWGA----TLGATAPASVLACNAAFGTCCAACAAVALTPTP-----
KAG7104944.1      -----------MRPTNIFQLAALTAVPASA-GLVAYG--ICQAGCASVVTACYGAAGFTWGA----TLGATAPASVLACNAAFGTCCAACAAVALTPTP-----
PNH30441.1        -----------MRPTKIVQLTALVVVPASA-GPVAYG--ICQAGCAGVVTACYGAAGFTWGA----TLGATAPASVLACNAAFGTCCAACAAVALTPTP-----
XP_009651807.1    -----------MRPTKIVQLTALAVVPASA-GPVAYG--ICQAGCAGVVTACYGAAGFTWGA----TLGATAPASVLACNAAFGTCCAACAAVALTPTP-----
KAG7104378.1      -----------MRPPKIVQLTALAAIPASA-GPVAYG--ICQAGCASVVTACYGAAGFTWGA----TLGATAPASVLACNAAFGTCCAACAAVALTPTP-----
KAF3353751.1      -----------MRPTKIVQLTALAAVPASAVGPVAYG--ICQAGCASVVIACYGAAGFTWGA----TLGATAPASVLACNAAFGTCCAACAAVPLTPTP-----
KAF8847622.1      -----------MRLSSILPVIAAVSTPAIA-GPAAYG--ICQAGCASVVMACYGAAGFTWGA----TLGASAPASIVLCNSAFGTCYAACAAALLLPTP-----
XP_025501802.1    -----------MKRLLPAICTGLL-VCHVQAGPAAYG--ICQAGCASVVTACYAAAGFTWGA----TLGATAPASVLACNGAFGICQGACATALLAPTL-----
XP_025440039.1    -----------MKGLLPAICTGLL-VCHVQAGPAAYG--ICQAGCASVVTACYAAAGFTWGA----TLGATAPASVLACNGAFGICQGACATALLAPTL-----
XP_025488249.1    -----------MKGLLPVICTSLL-ACYVQAGPAAYG--ICQAGCASVVTACYAAAGCTWGA----TLGATAPASVLACNGAFGICQGACATALLAPTL-----
KAE8155953.1      -----------MKILYPALLILLSSITQVNGGPAAYG--ICQAGCAAVVTACYTAAGFTWGA----TLGATAPASIVACNTAFGTCQAACATALLAPTP-----
XP_025577596.1    --------------LLTLLILTLTLPTPISASPTAYG--ICQAGCAAVVTACYSAAGFTWGA----TLGATAPPTILACNSAFGTCQAACAVTLLIPTL-----
KAF1356704.1      ----------MRLLALKPFWLTAMFITSCSAGPAAYG--ICQAGCCGVVTACYSAAGFTWGA----TLGATAPASILACNSAFGACQAACAVVALAPTL-----
XP_025460783.1    -----------MKNIYLAVFTILIFVSYASAGPAAYG--ICQAGCAAVVMACYSAAGYTWGA----TLGATAPPTIVACNSAFGVCYSSCAATLLAPTL-----
RDK39382.1        -----------MKNIYLAAYTIPISAGYASAGPTAYG--ICQAGCAAVVMACYSAAGYTWGA----TLGATAPPTIVACNSAFGVCYSSCAATLLAPTL-----
OJJ72525.1        -----------MKNLHLAVYTIPVIASYAYAGPAAYG--ICQAGCAAVVTACYSAAGYTWGA----TLGATAPASILACNSAFGTCCAACAATLLTPTL-----
XP_025540887.1    -----------MKNLYLAITTILILADYAYAGPLGYG--ICQAGCAAVVMACYSAAGYTWGA----TLGATAPASIVACNTAFGTCCAHCAATLLIPTL-----
XP_025383483.1    -----------MKNLYLAITTIFIFASYAYAGPLGYG--ICQAGCAAVVMACYSAAGYTWGA----TLGATAPASIVACNAAFGTCCAHCAATLLAPTL-----
OJZ90505.1        -----------MKKLYLLISTILILASYAYAGPMGYG--ICQAGCAAVVMACYSAAGYTWGA----TLGATAPASIVACNAAFGTCCAHCAATLLAPTL-----
XP_025520823.1    -----------MKKFYLLISTILILASYAYAGPVGYG--ICQAGCAAVVMACYSAAGYTWGA----TLGATAPASIVACNTAFGTCCAHCAATLLIPTL-----
OJI81929.1        -----------MKKLYLLISTILILADYAYAGPLGYG--ICQAGCAAVVMACYSAAGYTWGA----TLGATAPASIVACNTAFGTCCAHCAATLLAPTI-----
XP_025563775.1    -----------MNKLYLAITTILVLASYAHAGPVGYG--ICQAGCAAVVMACYSAAGYTWGA----TLGATAPASIVACNTAFGTCCAHCAATLLIPTL-----
XP_025477608.1    ------------------------------AGPLGYG--ICQAGCAAVVMACYSAAGYTWGA----TLGATAPASIVACNAAFGTCCAHCAATLLMP-------
KAH7119993.1      ---------MRLYNLAKYLPAVFFFNSTVYAGPAAYG--ICQAGCAGIVAACYGAAGCTWGA----TLGATAPATIIACNLAFGKCQAACAALLLAPTL-----
XP_022582555.1    -------------MKILEIGAVFSAVSLVMAGPAAYG--ICQAGCAAVVTACYSAAGFTWGA----TLGASAPASILLCNSAFGTCQAACAATLLAPTL-----
XP_033563937.1    ---------------------MLLLTKIVLAGPIGYG--ICQTSCSAFVMACYSATGFTWGA----TLAATAPASILVCNAAYGTCQTACATVLLGPTL-----
KAH5618967.1      ----------LLETIATTATAILLLTQSALAGPAAYG--ICQAGCSAVVMACYSAAGFTWGA----TLGATAPASILVCNAAFGTCQAACAAVLLAPTL-----
KAF1911237.1      ---------MKLTTIVNTVTALLLLPQCASAGPAAYG--VCQAGCSAVVMACYGAAGFTWGA----TLGATAPATILACNAAYGTFQAACAAVLLAPTL-----
KAH8711905.1      ---------MKLLHSFSLVTGIFLLGQSVSAGPAAYG--VCQAGCSAIVVACYSAAGFTWGA----TLAATAPASILACNSAFGTCQAACAAVLLAATP-----
XP_025509214.1    ----------MMNISYPALGVLALAASCASAGPAACG--VCQTGCAAVVMACYSAAGYTWGV----ALGAGVPATILACNSAFGTCQSACAAVILAPLF-----
```

```
PYI33230.1          --------------------VLALASSYASASPAAYG--VCQAGCAAVVMACYSGAGYTWGA----SLGATIPASILACNSAFGTCQSACAAVLLAPFP-----
XP_025522674.1      ----------------------ALAAGYASATPAGYG--VCQTGCATVVMACYSAAGFTWGA----ALGATIPASILACNSAFGTCQSACAAVLLIPFP-----
PYI14498.1          ----------------------ALAAGYASATPAGYG--VCQTGCATVVMACYSAAGFTWGA----ALGATIPASILACNSAFGACQSACAAVLLIPFP-----
XP_056813643.1      -----------MRAPWVLINNLLLFIPPVSAGPAAYG--ICQAGCSAVVMACYSAAGFTWGA----TMGASAPASVIVCNSAFGTCQAACAAALLAPTL-----
KAJ5955379.1        -----------MRAPWVLINNLLLFIPPVSAGPAAYG--ICQAGCSAVVMACYSAAGFTWGA----TMGVSAPASVIVCNSAFGTCQAACAAALLAPTL-----
XP_056989915.1      -----------MRAPWVLIYGLLLFIPFVFAGPAAYG--ICQAGCAAVVMACYSAAGFTWGA----TMGASAPASVLVCNSAFGTCQAACAAALLAPTL-----
XP_056573620.1      -----------MRVPWVLINNLLLFIPPVSAGPAAYG--VCQAGCAAVVMACYSAAGFTWGA----TMGASAPASVMACNLAFGKCQAACAAVLLAPTL-----
KAJ5277483.1        -----------MRVPWVLINNLLLFIPPVSAGPAAYG--VCQAGCAAVVMTCYSAAGFTWGA----TMGASAPASVMACNLAFGKCQAACAAVLLAPTL-----
KAJ6163906.1        -----------MRVPWVLINNLLLFIPPVSAGPAAYG--VCQAGCAAVVMACYSAAGFTWGA----TMGARAPASVMAFNLAFGKCQAACAAVLLAPTL-----
CDM32632.1          -----------MRVPWVLINNLLLFIPPVSAGPAAYG--VCQVGCAAVIMACYSAVGFTWGA----TMGASAPASVMACNLAFGKCQAACAAVLLAPIL-----
CDM32893.1          -----------MRVSRGKFLNLLLFSPTVLAGPAAYG--VCQAGCAAVVMACYSAAGFTWGA----TMGISAPASIVACNSAFGTCQGACASVLLAPTP-----
XP_014530676.1      -----------MKSLWAILYACLFFSSDVSAGPAAYG--VCQAGCATVVMAFYSAAGFTWGA----TMGATVPASILACNPAFGKCQAACASVLLAPTF-----
XP_057021283.1      -----------MRSHWAQFWLPLLLATNVSAGPAAYG--VCQAGCAALVMACYSAAGFTWGV----AMGATIPASIVTCNSAFGTCQAACASVLLAPTL-----
XP_040695488.1      --------------MKPSLLPLLLVASTASAGPAAYG--VCQAGCSAVVMACYSAAGFTWGA----TAGASAPASILACNSAYGTCQAACAAALLAPTL-----
KAE8371456.1        -----------MRHIPTSIVILLMLAREGFTGPAAYG--VCQAGCSAVVMACYSAAGFTWGA----TLGVSAPAFVIACNTAYGTCQAACAATLLTPTL-----
KAF2648513.1        ---------------LATVITLSTLAGEVAAGPAAYG--VCQAGCSAIVMACYSAAGFTWGA----TLGASAPATVLGCNTAFGACQAACAATLLAPT------
PSN58987.1          ----------MHSSFPATIIAYLALASAVSAGPAAYG--VCQAGCATVVMACYSAAGFTWGA----TLGASAPPTIIACNAAFGTCYSACAATLLAPTL-----
PSN59268.1          ----------MHSSFPATIIVYLALASAVSAGPAAYG--ICQAGCATVVMACYSAAGFTWGA----TLGASAPPTIIACNAALGTCYSACAATLLAPTL-----
KAF2687625.1        ------------------SISLLVLAGSVSAGPAAYG--VCQAGCAAVVTACYSAAGFTWGA----TLGASAPATIVACNAAFGTCQAACAATLLAPT------
KAH8691071.1        ---------MKFPSFSIIIIDCLVLARSASAGPAAYG--ICQAGCSAVVMACYTAAGFTWGA----TLGASAPATIIGCNAAFGTCQAACAATLLAPTP-----
KAF2024935.1        -----------MNRFAITVFSVLALADSGTAGPAAYG--ICQAGCSAIVMACYSAAGFTWGA----TMGASAPATILACNAAFGTCQAACAAALLAPTL-----
KAJ5059812.1        -----------MAISLLITLITLAFASLASAGPAAYG--LCQAGCSAVVMACYSAAGFTWGA----TLGASAPASIIACNTAFGTCQAACASVLLAPTP-----
XP_007693637.1      -----------MALLLLITLITLAFASLASAGPAAYG--LCQAGCSAVVMACYSAAGFTWGA----TLGASAPASIIACNTAFGTCQAACAAVVLTPTP-----
XP_007700491.1      --------------------------SLASAGPAAYG--LCQAGCSAVVMACYSAAGFTWGA----TLGASAPASIIACNTAFGTCQAACASVLLAPTP-----
XP_014552310.1      -----------MTSSLFIVIISLAFASPALAGPAAYG--LCQAGCSAVVMACYSAAGFTWGA----TLGASAPASIIACNTAFGTCQAACAAVLLSPTP-----
XP_007718708.1      -----------MASSFSIVVIALAFASPALAGPAAYG--LCQAGCSAVVMACYSAAGFTWGA----TLGASAPASIIACNTAFGTCQAACAAVLLSPTP-----
XP_007693411.1      -----------MALSILIATIVFALASPISAGPAGYG--LCQAGCSAIVMACYSAAGFTWGA----TLGASAPASIIACNTAFGTCQAACAAVLLTPTP-----
XP_007718397.1      -----------MELSLFIAIITLALASPASAGPAAYG--VCQAGCSAIVMACYSAAGFTWGA----TLGASAPASIIACNTAFGTCQAACAAVILTPTP-----
XP_007694026.1      -------------MPLLTAIITLAIASPALAGPAAYG--VCQAGCSAVVMACYSAAGFTWGA----TLGASAPASIIACNTAFGTCQAACAAVLLTPML-----
XP_009229506.1      ---------MRRFTSASIMLVTMAFTSPAFAGPAAYG--VCQAGCAAVVMACYSAAGFTWGA----TLGVSAPPTIIACNTSFGTCQAACAAVLLSPTP-----
KAF1972136.1        -----------------------VLAGPALAGPATYG--VCQAGCPALVMACYSAAGFTWGT----TLGASAPATIAACNTAFGTCQAACAAVLLAPMP-----
XP_058332260.1      ------------MKASSILSFLPIFAPAVFAGPAGYG--VCQAGCATVVMACYSAAGFTWGA----TAGLSAPASIIACNTAFGTCQSACAAVLLAPTP-----
XP_022405400.1      ------------MKIQTFFPLVLVLTSPVLAGPAAYG--VCQSGCASVVMACYSAAGFTWGA----TMGASAPASIVACNTAYGTCQAACAATILGPTP-----
XP_040731585.1      ------------MKLLSAALAALLLTTSVTAGPAAYG--VCQTGCAAVVMACYSAAGFTWGA----TLGATAPASIIACNTAYGTCQAACAAVLLTPTL-----
XP_031874101.1      ----------MRIFTPAFALAVLCLLDSATAGPVGYG--VCQAGCAAVVTACYGAAGFTWGA----TLGATAPASIIACNAAFGTCQAACAAVLLTPTP-----
KAF2262031.1        ------------MRLSSLSVELLLLAGTASAGPAAYG--VCQAGCAGVVMACYSAAGFTWGA----TMGASAPATIVACNAAFGSCQAACAAALIAPTL-----
KAF2867966.1        ---------------MRLSSVLILLAGSVSAGPAAYG--ACQGGCAKIVMACYAAAGCTWGA----TMGASAPPTIIACNTAFGSCQAVCAALLLAPTL-----
XP_033393871.1      -----------MLRTTILTACVLTLAGTASAGPVGYG--ICQAGCAGVVMACYTAAGFTWGA----TLGASAPPTIIACNTAFGSCQAACAAILLAPTP-----
XP_007687291.1      --------MKPLSLTKIAIAVLSLTAGQASAGPVGYA--VCQAGCAGVVMACYSAAGFTWGA----TFGATAPASILLCNAAFGKCQAACAFVLLGPTP-----
XP_014082052.1      ------------------------------------------AGCAGVVMACYSAAGFTWGA----TFSASAPASILLCNAAFGKCQAACAVVLLGPTP-----
XP_057023110.1      ------------MRVLPLLIPIYILLSFAAAGPISYG--ICQAGCAAVVMACYSAAGFTWGA----TMGATAPASILLCNTAFGKCQAAAAVLLSPTP-----
KAF2490063.1        -----------MKLRLSILALLTGAAVPISAGPIGYG--ICQAGCAGVVMACYSAGGATWGA----TLGATAPATIIGCNTAFGSCQAACAAVLLAPTP-----
XP_033573400.1      -----------MKLRASILALLTTFTIPIAAGPIGYG--VCQAGCAGLVMACYSAAGFTWGA----TLGATAPATIVACNTAFGSCQAACAAVLLTPTP-----
XP_056549082.1      -----------MRLSLPIHVLFLCFSTAAVAGPVGYG--VCQAGCAGVTMACYSAAGFTWGA----TAGATAPATIVGCNLAFGKCQAACAAVLLMPTP-----
OIW28253.1          ----------MRASSAIPAISLLFLAQGAVASLVGYG--ICQAGCAGVVMACYSAAGFTWGA----TMGVSAPATIIACNSAFGSCQAACATVLLAPIP-----
KAB5545907.1        ----------MRVFSAIPTVSILLLASEVIAGPVGYG--VCQGGCAGVVMACYGAAGFTWGA----TLGATAPASILACNAAYGSCQAACAAVLLVPLP-----
XP_040642819.1      --------------MKLSTAILLPLASTAIAGPIGYG--ICQAGCSGVVMACYSAAGFTWGA----TLGASAPASIIACNTAYGTCQAACAAVLLAPTP-----
KAI1811651.1        --------------MKLIATILLANALAVEAGPVGYG--ICQAGCSGVVMACYGAAGFTWGA----TLGASAPASIIACNTAFGTCQAACAAVLLTPTP-----
KAH7066527.1        ----------MACRAFLLVTTTLAFAGPVHAGPIGYG--ICQAACSAVVMACYSAAGFTWGA----TLGATAPSSILACNPAFGTCQAACAAVLLAPTP-----
KAF1970296.1        -----------MRLSTLTATALLALTSTVAAGPIGYG--VCQAGCSTVVMACYAAAGFTWGA----TLGATAPASIVACNAAYGTCQAACATVLLGPTP-----
XP_033524879.1      ---------MRFPTTLTTLTAMLALIPCATAGPIGYA--ICQAGCSSVVMACYAAAGFTWGA----TLGATAPASIVACNTAFGTCQAACATVLLGPLP-----
KAH8600237.1        -----------MQISSSTITLLMAFTPPIAASLIGYG--VCQAGCSTVVMACYGAAGFTWGA----TLGATAPATIIACNSAFGTCQAACAAVLLVPLP-----
USP72936.1          --------MKSTTITKASVALLTFTTGRVSAGHIGYG--ICQAGCASVVMACYAAAGYTWGA----TMGATAPASIIACNSAFGTCQAACAAVLLGPTL-----
KAH7025312.1        ----------MLSSRSLVIGFILATQSLVSAGPGAYG--VCQAGCSGVAMACYAAAGFTWGA----TLGATAPASIVACNSAFGACQAACAAVLLAPTL-----
KAH7014050.1        ----------MPASRSLLVGFILATQSLVSAGPTAYG--VCQAECSGVVMACHAAAGFTRGA----TLGATAPASIVACNSAFGACQAACAVVLLAPTL-----
XP_024702685.1      ----------------------MSLTNEVSAGPTGYG--ICQAGCSGVAMACYAAAGCTWGA----TLGATAPPAIVACNAAFGACQAKCALVLLAPTP-----
CAE7186653.1        ---------MKLPTLATVATASLIFAHSASAGPIGYG--ICQAGCSAVVMACYSAAGFTWGA----TLGATAPASILACNAAYGTCQAACAAVLLGPTP-----
KAA8618168.1        ---------MKLPALTAVTTASLFFAHSASAGPIGYG--ICQAGCSAVVMACYSAAGFTWGA----TLGATAPASILACNAAYGTCQAACAAVLLSPVP-----
KAF2825151.1        -----------------------LLTNTAVAGPIGYG--ICQAGCSGVVMACYSAAGFTWGA----TLGATAPASILACNSAYGACQAACAAVLLGPWP-----
KAF1937296.1        --------MKFLINIITTALAILIFATTASAGPIGYA--ICQGGCAGVVMACYSAAGFTWGA----TLGATAPATVLACNAAYGTCQAACAAVLLVPLP-----
XP_001803436.1      --------MKLATTTVTTAATAALLVSMVSAGPIGYG--ICQAGCSAVVMACYSAAGFTWGA----VLGATAPATIVVCNTASGTCQAACAAVLLGPTP-----
KAF1839331.1        ---------MKLPFFTTMAAAFLILACLASAGPVAYG--ICQGGCSAVVMACYSAAGFTWGA----TLGASAPATIVACNAAFGTCQAACAAILLAPTF-----
XP_033556128.1      -----------------MAAAFLILACLASAGPVAYG--ICQGGCSAVVMACYSAAGFTWGA----TLGATAPATVVACNAAFGTCQAACAAVLLAPTL-----
OCK77742.1          ----MKQCQNLLIIVRFSVPLAIILAGAASTGPIGYG--ICQGGCATIVMACYGAAGATWGA----TLGATAPATVIACNSAYGACQAACASVLLLPTP-----
KAI1338517.1        ----------MKLNLTPLSPVLLALASVTSAGPIGYG--ICQAGCASVVTACYGAAGATWGA----TAGATAPATVLACNSAFGTCQAACAAVLLTPTP-----
XP_056783524.1      ------------MKLSILAAALPLMAANVLAGPIGYA--ICQAGCASVVMACYAAGGATWGA----TLGASAPPTIVACNTAYGTCQAACAAVLLGPTP-----
KAJ5116293.1        ------------MKLSIIAAAFLLMAVNVLAGPIGYG--ICQAGCASLVMACYAAGGATWGA----TLGATAPPTIVGCNTAFGTCQVACAAVLLVPFP-----
KAJ6031359.1        -----------MKTSTILSLGSLSMVPTVLAGPIGYG--ICQAGCSSVVMACYTAGGATWGA----TLGATAPATIVGCNAAFGTCQAACAALLLTPIP-----
XP_056945832.1      -----------MKASTFLSLGTLSLVPTVLAGPIGYA--ICQGGCSSVVMACYAAGGATLGA----TLGATAPATIVGCNVAFGTCQAACAAVLLTPIP-----
KAJ5613654.1        -----------MKTSTLLSLGGLSLAPTVLAGPIGYG--ICQAGCSSVVMACYVAGGATWGA----TLGATAPPTIVGCNVAFGTCQAACAAVLLTPTP-----
KAJ5710925.1        -----------MKTSTLLSIVGLSLKPTVLAGPIGYG--ICQAGCSSVVMACYAAGGATWGA----TLGATAPPTIIACNVAFGTCQAACAAVLLTPTP-----
KAJ5738444.1        -----------MKTSTLLCLGTIFLTPTVLAGPIGYG--ICQAGCSSVVMACYVAGGATWGA----VLGATAPATIVGCNAAFGTCQAACAAVLLKPAP-----
QDS77261.1          -------------MHPSLLLTITALATTTTAGPLGYG--ICQAGCSSIVMACYAAAGFTWGA----TLGASAPATIVACNAAYGTCQAACASVLLTPTP-----
KAE9964269.1        -------------MRTSILITAAALAATISAGPLGYA--VCQAGCASVVMACYAAGGATWGA----TLGATAPATIVACNAAYGTCQAACAAVLLIPFL-----
KAF9638679.1        -----------MRFQTIVFAGSLIFAGTATAGPIAYG--ICQAGCASVVMACYAAAGFTWGA----TLGATAPASIIACNAAFGTCQAACAVVALAPTP-----
CAI6339829.1        -----------MRFHYLTPLALLGLAGTTQAGPAAYG--ICQAGCADVVMACYSGAGFTWGA----TPGATAPATIIACNAAFGTCQAACAAVIFASTP-----
TEY32995.1          -------------MKPTSILLIACLAGITTAGPVAYG--VCQSGCAAVVMACYSAGGATWGA----TLGATAPATIVACNKAFGTCSATCAGLLVAPIP-----
KAF7952734.1        -----------MKPTSTSILLIAFLAGITTAGPVAYG--VCQSGCAAVVMACYGAGGATWGA----TLGATAPATIVACNTAFGICSAKCAGLLVAPIP-----
XP_038808418.1      -----------MKPSSSSILLIAVFAGITTAGPIAYG--VCQSGCAAVVMACYGAGGATWGA----TAAATAPATIVACNTAFGVCSAKCAVLLAAPIP-----
XP_038754834.1      --------MKPSSSSASSILLIAVFAGITTAGPVAYG--VCQSGCAAVVMACYGAGGATWGA----TAAATAPATIVACNTAFGVCSAKCAVLLAAPIP-----
XP_037194411.1      ----------MKPSSSKSILLIAVLAGIATAGPIAYG--VCQSGCAAVVMACYGAGGATWGA----TAGATAPATIVACNTAFGVCSAKCAGLLVAPIP-----
XP_024549469.1      -----------MKPTSPSVLLVAGLAGIATAGPIAYG--ICQSGCAAVVMACYSAGGATWGA----TLGATAPATIVACNTAFGTCSATCAGFLVAPIP-----
TGO59710.1          ---------MNPSPSSTLLITVAILAGIATAGPVAYG--VCQSGCAAVVMACYGAGGATLEK----IYTDGDADVTQL--------------------------
XP_038729586.1      ---------MNPSPSSTFLITVAVLAGIATAGPVAYG--VCQSGCAAVVMACYGAGGATWGA----TLGATAPATIVACNTAFGVCSAKCAGLLVAPIP-----
ESZ95438.1          --------MKPKSSLLIFFFFVSTLMGTATAGPIAYG--ICQSGCAAVVMACYAAGGATWGA----TLGATAPATIVACNTAFGVCSAKCWVAAALPFF-----
XP_001596882.1      --------MKPTSTSILLLTAITTFMGMATAGPVAYG--VCQSGCAGVVMACYSAAGFTWGA----TLGATAPPTIIACNTAFGLCSAKCAGFLVAPIP-----
KAJ8059346.1        --------MKPTSTSILLLTTLTTFMGMATAGPVAYG--VCQSCCAIVVMACYSAAGFTWGA----TLGATAPPTIIACNAAFGLCCAKCAGFLVAPIP-----
KAF7871723.1        --------MKPTSTSILLLT--TTFIGIATAGPVAYG--VCQSGCAGVVMACYSAAGFTWGA----TLGATAPPTIIACNTAFGFCSAKCAGFLVAPIP-----
CAD6445730.1        -----------MKPTSILLTTLASFMGMASAGPIAYG--ACQSGCAAVVMACYLAAGFTWGA----TLGATAPPTIIACNTAFGLCSAKCAGFLVAPIP-----
KAI9642175.1        --------MKPALLLGAGALTILTSIPTTTAGPVTYG--ACQSGCAVVVMACYSAAGFTWGA----TLGATAPATVIACNVAFGKCSATCAGLLLAPTP-----
PQE30004.1          ----------MKLSFTATLLTLAPLLPFTTAGPAAYG--VCQSGCAAVVMACYTAAGFTWGA----TLGATAPATILACNAAFGTCSATCAALLLAPTP-----
KAA8570577.1        MKLQSASRFSIPLATTLLTLTLFTIIPPATSGPIAYA--SCQAGCAAVVTACYSAAGFTWGA----TLGATAPASIVACNAAFGTCYAACAGFLVAPIP-----
KAB8299301.1        MNLQSVSLFSFLLP--LLALFLLNIIPPVTAGPIAYA--SCQAGCAGIVMACYSAAGFTWGA----TLGVTAPASIVACNAAFGTCYAACAGFLVAPIP-----
XP_033535919.1      -------------SNLLIILPFTVPIITVSAGPVLYA--GCQVGCAGAVMACYSAAGFTWGA----TLGITAPATIIACNMAFGSCQAGCAAALLILTL-----
PVH94442.1          ---------MHFPSISTVSAGFLALAGTVSAGPAAYG--ICQAGCAGIVMACYSAAGFTWGA----TAGAMVPATNIACNASFGKSQGACVLVLL---------
KAF5552569.1        ------------MKPMHLLVAITSLSTTAVAGPIAYG--VCQAGCASVAMACYAAGGATWGA----TAGATAPATIAACNSAFGVCSAACAQTALIAPTP----
XP_036534571.1      ------------MKPLHLLVAITSLSTTAVAGPIAYG--VCQAGCASVAMACYAAGGATWGA----TAGATAPATIAACNSAFGVCSAACAQTALIAPTP----
KAF5989468.1        ------------MKPLHLLVAITSLSTTAAAGPIAYG--VCQAGCASVAMACYAAGGATWGA----TAGATAPATIAACNSAFGVCSAACAQTALIAPTP----
PNP73544.1          ------------MKPLPLLVAIASFSTTTVAGPIAYG--VCQAGCASVAMACYAAGGATWGA----TAGATAPATIAACNSAFGVCSAACAQTALIAPTP----
KAG5753170.1        ------------MKPLRFLVAIASFSTTTVAGPIAYG--VCQAGCASVAMACYAAGGATWGA----TAGATAPATIAACNSAFGVCSAACAQTALIAPTP----
KAF5592368.1        ------------MKPLHLLVTIASFSTTAVAGPIAYG--VCQAGCASVAMACYAAGGATWGA----TAGATAPATIAACNSAFGVCSAACAQTALIAPTP----
XP_018761531.1      ------------MKPLHLLVTITSFSTTVVAGPIAYG--VCQAGCASVAMACYAAGGATWGA----TAGATAPATIAACNSAFGVCSAACAQTALIAPTP----
KAF5574014.1        ------------MKPLHLLVATVFFSTTVVAGPIAYG--VYQAGYASVAMACYAAGGATWGA----TAGATAPATIAAYNSAFGVCSAACAQTALIAPTP----
KAF5254884.1        ------------MKSLHLLVAVGSFSTAAVAGPIAYG--VCQAGCASVAMACYAAGGATWGA----TAGATAPATIAACNSAFGVCSAACAQTALIAPTP----
EMT71922.1          ------------MKAISFLATIAGFSTTARAGPIAYG--VCQAGCASVVMACYAAGGATWGA----TAGATAPATIAACNSAFGVCSAACAQTALVAPTP----
RKK72021.1          ------------MKAISFLATIAGFSTTARAGPIAYG--VCQAGCASVVMACYAAGGATWGA----TAGATAPATIAACNSAFGVCSAACAQTALIAPTP----
XP_046041803.1      ------------MKAISFLAIIAGFSATASAGPIAYG--VCQAGCASVAMACYAAGGATWGA----TAGATAPATIAACNSAFGVCSAACAQTALIAPTP----
KAH7151049.1        ------------MKAISFLATIAGFSATACAGPIAYG--VCQAGCASVAMACYAAGGATWGA----TAGATAPATIAACNSAFGVCSAACAQTALIAPTP----
ENH74830.1          ------------MKAISFLTIIAGFSTTARAGPIAYG--VCQAGCASVVMACYAAGGATWGA----TAGATAPVTIAACNSAFGVCSAACAQTALIAPTA----
KAH7466833.1        ------------MKAISFLKIIAGFSTTARAGPIAYG--VCQAGCASVVMACYAAGGATWGA----TAGATAPATIAACNSAFGVCSAACAQTALIAPTP----
XP_059468083.1      ------------MKAIGFFTIIAGFSTTARAGPIAYG--VCQAGCASVVMACYAAGGATWGA----TAGATAPATIAACNSAFGVCSAACAQTALIAPTP----
KAF4339535.1        ------------MKPLHLFTTITCLSATAHAGPVAYG--VCQAGCASVAMACYAAGGATWGA----TAGATAPATIVACNSAFGACSAACAQTALIAPIP----
KAG7410298.1        ------------MKLLSVVTTLAILSTTVSTGPVAYG--VCQAGCAGVVMACYGAAGYTWGA----TVGATAPATIIACNSAFGTCSAACAKVALLAPTP----
EXK76060.1          ------------MKLLSVVTTLAILSTTVSTGPVAYG--ICQAGCAGVVMACYGAAGYTWGA----TVGATAPATIIACNSAFGTCSAACAKVALLAPTP----
EXK76156.1          ------------MKLLSVVITLAILSTTVSTGPVAYG--VCQAGCAGVVMACYGAAGYTWGA----TVGATAPATIVACNSAFGTCSAACAKVALLAPTP----
PNP60693.1          --------MTSRMMLMSPVATLAILSTTVSAGPVAYG--VCQAGYAGVIMACYGAAGYTWGA----TVGATAPATIIACNSAFRTCSAACAKVALLAPTV----
KAH6977657.1        ------------MKLLSPVPTLAILSTAASAGPVAYG--VCQAGCAGVVMACYGAAGYTWGA----TIGATAPATIVACNSAFGTCSAACAKVALLAPIP----
KAI8397321.1        ------------MKLLSVVTTLAILSTTVSTGPVAYG--VCQAGCAGVVMACYGAAGLQQRV----WDMLRG------------LCQGSLAGPYTMK-------
KAH7251348.1        ------------MKPTAILATLACFYTPTSAGPIAYG--VCQAGCSAVVMACYTAAGYTWGA----TLGASAPASIIACNGAFGTCSAACAKIGLFAPTP----
CAJ0554546.1        ------------MKPAAILATLACFYPATSAGPIAYG--ICQAGCSGVVMACYAAAGYTWGA----TLGASAPASIVACNGAFGTCSAACAKIGLFAPTP----
KAH6963189.1        ------------MKPAAILTTLTCLYTTAAAGPIAYG--VCQAGCSAVVMACYAAAGYTWGA----TLGASAPASIVACNGAFGTCSAACAKIGLFAPTP----
SPJ71122.1          ------------MKFSVVLATLACLYTTTTAGPIGYG--ICQSGCSAVVMACYAAAGYTWGA----TLGASAPASIIACNSAFGTCSAVCAKVALLAPTP----
RBQ81898.1          ------------MKPRVTVAVAALFFTGASAGPQAYA--ACRAGCSAIVMACYGAAGFVWGT----RLSATAPPTIIASNNAFGTCSAVCAKVALFAPTP----
```

```
KAJ4012322.1        ------------MKPTTPLIAIAYAASTVSAGPLAYA--ACQSGCAGVVMACYSAAGFTWGA----TFGATAPASILLCNAAFGTCSATCAQVALLAPTP----
KAI1061423.1        ------------MKPSTPLSAIACAASTVSAGPLAYA--ACQSGCAGVVMACYSAAGFTWGA----TFGATAPASILLCNAAFGTCSATCAQVALLAPTP----
CAG7562008.1        ------------MKPTTPLITIACVASPVSAGPLAYA--ACQSGCAGVVMACYSAAGFTWGA----TFGASAPASILLCNSAFGTCSATCAQVALFAPTP----
XP_045982074.1      ------------MKPTTPLIAIACLASTVSAGPLAYA--ACQAGCATIVMACYSAAGYTWGA----TLGASAPATILACNAAFGQCSAVCAQVALVAPTP----
CCX30515.1          -----------MKVSALTLTVAAMFASSVTAGPLAYA--ACQAGCATVVMACYTAGGATWGA----TLGATAPATIIGCNSAYASCQAVCASVALFAPTP----
KAI5802037.1        -----------MKVSALTLTVAAMFASSVTAGPLAYA--ACQAGCATVVMACYTAGGATWGA----TLGATAPATIIGCNSAYASCQAVCASVALFALTP----
KAI5789794.1        ------------MKVCTPLTVAAMFASSVTAGPLAYA--ACQAGCTTVVVACYAAGGATWGA----TVGATAPATIIACNSAYASCQAVCATVGLFTPTP----
CCX16445.1          ------------MKVCTPLTVAAMFASSVTAGPLAYA--ACQAGCTTVVVACYAAGGATWGA----TVGATAPATIIACNSAYASCQAVCATKCFEGNVLDLAT
KAI5816642.1        ----------------------ILFGSNVTAGPLAYA--ACQAGCATVVMACYSAAGFTWGA----TLAATAPATVIACNSAYGSCQTVCATIGLLAPTP----
KAI5818345.1        -----------MKVRTSVTVAALLFGSSVTAGPLAYA--ACQAGCATVVMACYSAAGFTWGA----TLGATAPATVIACNSAYASCQTVCATVGLFAPTP----
KAF8244270.1        ------------MKLSLIVTAALLSASGAHAGPLAYA--ACQAGCAGVVMACYSAAGATWGA----TLGATAPATVLGCNAAYASCQGVCATVALCAPIP----
KAA8906278.1        ---------------MKPITLLLALPATVTAGPLAYA--ACQGGCAAVVMACYGAAGYTWGA----TLGVAAPATVLACNAAYATCQATCATICLFAPTP----
KAF5230862.1        ------------MKSSSTIVITTAFLFTVSAGPLAYG--ACQAGCAGIVMACYSAAGYIWGA----TARATAPASIIDCNAAFGKCSANETFSETLPCHQE---
VTO82721.1          ------------MKSSSTIVITTAFLFTVSAGPLAYG--ACQAGCAGIVMACYSAAGYIWEA----TAGVTAPASIIDCNAAFGKCSAVCAAIALGTPTS----
QPC57932.1          ------------MKSSSTIAITIALLTTVSAGPLAYG--ACQAGCASIVMACYSAAGFTWGA----TAGATAPASIIACNAAFGKCSAVCAAIALGAPTP----
QPC69322.1          ------------MKFSSTIAITIALFDTVSAGPLAYG--ACQAGCASIVMACYSAAGFTWGA----TAGATAPASIIACNAAFGKCSAVCAAIALGAPTP----
KAH7003006.1        -------------MKISTITATTALLATAFAGPLAYG--TCQAGCASVVVACYSAAGFTWGA----TAGATAPATIIACNAAFGKCSAVCAAIALAPTP-----
KAG9233117.1        ------------MRPSSLLIPVVTFTTLASAGPAAYG--ICQAGCAAVVTACYAGAGFTWGA----TLGASAPATILACNAAFGTCQAACWAALIAPTP-----
KAI1248405.1        ---------MKVTTPAAALVAILTAAPGAIAGPAAYG--ICQAGCADVVTACYAAAGFTWGA----TLGASAPATIVACNTAFGSCQSACWAALVAPTP-----
OCK90052.1          ------------MHSYALLLTALTLTHFTYAGPAAYG--IRQAGCAAVVMACYSAAGFTWGA----TLGVSAPASILACNAAFGTCQGACAVAFIAPTP-----
OCL09912.1          ------------MHPRRVLLATLTLTHFAYAGPAAYG--ICQAGCAAVVMACYSAAGFTWGA----TLGIAAPATILACNSAFGTCQGACAVAFIAPTP-----
RYO93046.1          -----------MKLTTSILLAIVAATPTIHAGPAAYG--VCQAGCSAVVQTCYAAAGFTWGA----TLGATAPASIVACNSAYGACQAACWTALFSLTP-----
RYP24059.1          -----------MKLTTSILLAIVAAVPTIHAGPAAYG--ACQAGCSAVVQACYAAAGFTWGA----TLGATAPASIVACNNAYGACQAACWAALFSPTP-----
RYP30960.1          -----------MKPTTPILLAIVAAMPTILAGPAAYG--ICQAGCSAVVKACYAAAGFTWGA----TLGATAPATIVACNGAYGACQASCWAALFAPTP-----
RYP08623.1          -----------MQPTIPILLALIATTPTILAGPAAYG--VCQAGCSGVVMACYAAAGFTWGA----TLGATAPASIVACNTAYGACQAACWAALAAPTP-----
RYP76942.1          -----------MKPTSPILLALIAATPTILAGPAAYG--VCQAGCSGVVMACYAAAGFTWGA----TLGATAPASIVACNTAFGACQAACWAALAAPTP-----
RYP75671.1          -----------MKRTSPILLTLIAATPTILAGPAAYG--VCQTGCAGVVMACYAAAGFTWGA----TLGATAPASIVACNSAYGACQAACWAALAAPTP-----
RYP51836.1          -----------MKPTTPIRLALIATTPTVFAGPAAYG--VCQAGCSGVVMACYAAAGFTWGA----TLGATAPASIIACNSAYGACQAACWAALAAPTP-----
KAI1006201.1        -----------MRISLPLLVAISTAAP-VIAGPAAYG--ICQAGCAGVVMACYTAAGFTWGA----TLGATAPATIIACNAAYGTCQAACAVAIFAPTP-----
KAI1156788.1        -----------MKPITPFITAILAFVPVGSAGPAAYS--LCQGGCSAVVMACYEAAGFTWGA----TMGASAPATIVACNTAFGTCQASCWAAAIAPTP-----
KAI8947707.1        -----------MKPSTPFVTALLTFTPIVSAGPASYS--ICQAGCAAVVKACYAAAGFTWGA----TLGISAPASIVACNAAFGTCQASCWVAAIVPTP-----
KAI0972575.1        -----------MRPTTSLVATLIGVAPIVSAGPAAYG--ICQAGCAAVVTACYAAAGFTWGA----TMGASAPATVIGCNTAFGACQAACWAAAIAPTP-----
KAI1312583.1        -----------MRPATSLIAALIGFAPTVSAGPAAYG--LCQTGCAAVVTACYAAAGYTWGA----TLGASAPATILACNTAFGTCQAACWAAAIAPTP-----
KAI1187125.1        -----------MKPTKIFTAAILAFAPIVSAGPAAYG--VCQAGCAAVVTACYAAAGFTWGA----TLGASAPATIIACNTAFGTCQAACWLAAIAPTP-----
KAI0459375.1        -----------MKPTTPFVTAILASTPIVSAGPAAYG--ICQTGCAAVVTACYAAAGFTWGA----TAGISAPATIVACNSAFGTCQAACWVAAIAPTP-----
KAI1756746.1        -----------MKPTTPFVTAILASTSIVSAGPAAYG--ICQAGCAAVVTACYAAGGFTWGV----TAGLSIPATIIACNTAFGTCQAACWVAAIAPTP-----
KAI0555338.1        -----------MKTTTALVTALLASTSIVSAGPAAYG--ICQAGCAAVVTACYAAAGFTWGA----TVGVSAPATIIACNSAFGTCEAACWAAAIAPTP-----
KAI1736724.1        -----------MKATTALVTSLLASTPIVSAGPAAYG--ICQAGCAAVVTACYAAAGFTWGA----TAGVSAPATIIACNSAFGTCQAACYVAAFAPTP-----
KAH8159225.1        -------------MKLTIPFLTATLAPIVSAGPAAYG--ICQTGCAAVVVACYAAAGFTWGA----TAGISAPATILACNSAFGTCQASCWVAAIAPTP-----
KAI0447941.1        -----------MNLTTSFLTATLALAPIVSAGPAAYG--ICQAGCAAVVTACYAGAGFTWGA----TAGISVPATIIACNSAFGTCQASCWVAAIAPTP-----
KAI0859552.1        -----------MKPTTSLFAALLASTPIVSAGPAAYG--ICQAGCAAVVTACYAAAGFTWGA----TAGLSIPATIVACNTGFGTCQAACCAMTG----------
TRX93855.1          -----------MKPTTSLFAAILASTPIVSAGPAAYG--ICQAGCAAVVTACYAAAGFTWGA----TAGISAPATIVACNTASGTCQAACPFFFSAQEG-----
KAI1176761.1        -----------MKLNTHLVS-ALAFTSAASAGPIAYG--LCQAGCAAVVTACYGAAGFTWGA----TMGASAPASIVACNTAFGACQAGCWAAAIAPTP-----
KAI0404163.1        -----------MKVTTPLISAVLAFSPIASAGPLAYG--LCQAGCAAVVTACYSAAGFTWGA----TMGASAPATVVACNAAFGTCQAACWAAATAPTL-----
KAK2073227.1        -----------MQPIKILFPAMLIFGHAAHAGPVAYG--LCQAGCAAVVTACYAAGGATWGV----TGGASVPPTIVACDSAFGACQAACWAAVIAPTP-----
KAI1118733.1        -----------MQPIRAFAATILVFAPIACAGPAAYG--VCQAGCAAVVMACYGAAGCTWGA----TLGASAPATIIACNAAFGVCQAACWVAAISPGP-----
KAI1132034.1        -----------MKLTTTFLAALAVLAPLGNAGPAAYG--LCQAGCASVVTACYAAAGFTWGA----TLGASAPASIVACNAAFGTCQAACAAIALAPTP-----
KAI0817434.1        -----------MRLTTTFAPVLLGFASIASAGPAAYA--ACQTGCAGLVTACYAAAGFTWGA----TLGATAPASILACNSAFGTCQAACWVAAIAPTP-----
SMQ45019.1          -----------MRLLKMSLATLAVIAPAISAGPAAYG--VCQAGCAGVVVACYAAGGFVFGA----AAPPVLPPVIVACNTAFGACQAACWAALMMPTP-----
SMR41375.1          -----------MRLLKMSLATLAVIAPAISAGPAAYG--VCQGGCAKVVKACYAAAGFIWGS----AFDGPVPATVQACNGAFGICQGACWTALLMPVP-----
XP_043001095.1      --------------RHRTFATVAVFVPCVVAGPALYG--VCQAGCAAVVMACYSAAGATWGA----TAGITAPASVLACNAAFGKCSLACYIAAGAPTP-----
RKU46238.1          -------------MRLSRTFTAFTLAPAACAGPLAYG--MCQAGCSGIVMACYAAAGFTWGA----TLGATAPASILLCNSSYGSCQAVCAAIALAPTP-----
KAF7513366.1        -----------MKATAFVPLLSLTLTPLVFAGPLSYG--ICQAGCASIVMACYAAAGATWGA----TLGATAPASVLACNAAYGSCQAACIAAGCIPIP-----
XP_047761909.1      ----------MPSRTFTLLATLTTLVPLANAGPAAYG--ICQTGCAGVVMACYGAAGFTWGA----TLGATAPASVLACNSAYGACQAACASTLLVPFF-----
KAF2218680.1        -----------------------VLPGLAQAGPAAYG--VCQAGCAGIVMACYAAAGFTWGA----TLGATAPASIIACNTTFGACQATCAALLLAPTP-----
KAI0486197.1        -----------MKPTALLATSILALAPLASAGPAAYG--ICQAGCAGVVMACYGAAGFTWGA----TAGLTLPASVIACNTTFGACQAACWAAAIAPTP-----
ORY12622.1          ---------MRLSNLSIAAAALLMLPDFTSAGPAAYG--VCQAGCAAVVVACYGAAGFTWGA----TLGASAPATIIACNTTFGTCQAACWAALFTPTP-----
XP_049117280.1      -------------MKSTKFLASTLFAATTIAGPIAYG--VCQAGCSAVVMACYSAAGFTWGA----TLGASAPASIILCNTSYGSCQAACWAALAMPAP-----
TLD26839.1          ---------MQYSTITKAIVASAILNGGVLAGPAAYG--VCQSGCSAVVMACYAAAGFTWGA----TLGASAPASIIACNTAFGFCQAGCHAALVMPTP-----
KAI7911282.1        ---------MQHSTIAHAIMASAVLSGVALAGPAAYG--ICQAGCSGVVMACYGAAGFTWGA----TLGASAPASILACNTAFGACQASCHAVLFIPGP-----
XP_030984416.1      ---------MHHLTIAKAITATAILSGPALAGPGAYG--VCQAGCCAVVMACYAAAGATWGA----TAGATAPATVVACNTAFGACQASCHAALIFPIP-----
XP_029751233.1      ---------MHYSAIAKVIMATAVLNGLALAGPAAYG--VCQAGCCAVVMACYTAAGATWGA----TAGATAPATVVACNTAFGSCQAACHAALLMPVP-----
XP_009230175.1      ---------MRPSTSVSALLAAMAFAGLALAGPIAYG--ICQAGCAALVMACYAAAGATWGA----TLGATAPATVIACNSAFGTCQAGCATALLLPIP-----
KAF8434007.1        ------------MRFSLIIPAVAAIFGTASAGPLAYG--ICQAGCSALVVSCYTAGGLTFGTI---TAGAGAPAIALACNASYGVCQAACAAAILAPTP-----
KAF8424820.1        -------------MRSSFIFPLVAMLGTASAGPLAYG--ICQAGCSTLVVSCYAAGGLTFGTV---TAGLGAPAVALACNASYGACQAACAAAILAPTP-----
KAG7005615.1        ----------MHFIKFSTFVAISSLMTTAVAGPVAYG--ICQAGCATVATACYAAAGATFGTI---TAGAGTPAVILGCNSAFGVCSAKCAAIALLPTP-----
KAI4224460.1        ------------MKLTNLLTTFASLTTLTSAGPAGYG--VCQAGCSAVVVACYSGAGFTFGTV---TAGAAVPAAIVACNSAYGTCQAACAAVLIAPTP-----
KAI4226682.1        ------------MKLAKLLTTLATLTAVTSAGPAGYG--VCQAGCSAVVVACYSGAGFTFGTV---TAGAAVPAAIVACNSAFGTCQAACAAVLIAPTP-----
KAI4258584.1        ------------MKPTILLLPLTTLSALTSASPAAYG--ICQAGCSALVIACYSGAGFTFGTV---TAGAAIPAAIVACNSAFGTCQAACAAVMIAPTP-----
KAI4247966.1        -------------MHPLKSLPLLALIPLVTAGPAAYG--ICQAGCSAVVVACYSGAGFTFGTV---TAGAGIPAAVVACNSAYGTCQAACAAILLAPTP-----
KAH8767254.1        ------------MQPTKTIIAALSLATALTAGPVAYG--ICQAGCAAVVTACYAGAGFTFGT----VLAEAAPPAIIACNSAFGTCQASCAIVALFPCP-----
KAI7781638.1        ------------MQLTKIIIPALSLVTAVAAGPIAYG--TCQAGCAAVVVACYAGAGFTFGT----VLGPAAPASIAACNTAFGFCQAKCAVVALLPTP-----
XP_052987644.1      -------MVIITMRFTRTIIAALPLAIAIIAGPNAYS--ICQAGCSAVLAACYSAAGARVGT----VYGSAVPVPIADCNAAFMACNAMCAVITLSQTA-----
KUI64379.1          -------------MQPIKMLAVLTMATTATAGPIGYG--ICQAGCSAVVTACYAAAGVTFGT----IAALAAPAAIVGCNTAFGTCQAACAAVLLTPTP-----
KUI62490.1          -------------MQPIKMLAVLTMATTATAGPIAYG--ICQAGCSAVVTACYAAAGATFGT----VAAPAAPAAIVGCNTAFGTCQAACAAVLLTPTP-----
ROV87140.1          -------------MRPIKILAILAMATGTTAGPIAYG--ICQAGCAAVVTACYAAAGATFGT----VAAPAAPAAIVACNSAFGTCQAACAVVALAPTP-----
KAF8463719.1        ---------MRFHRSLSTLLPALLLVTGTEAGPIGYA--LCQGGCATVVMACYAAAGAVWGTNPWLAAGVPVPAVIVGCNSAYAGCQATCAAVAFSPTP-----
KAF8463718.1        ---------MRFHRSLSTLLPALLLVTGTEAGPIGYA--LCQGGCATVVMACYTAAGAVWGTNPWLAAGVPVPAVIVGCNSAYAGCQATCAAVALSPTP-----
KAF8453073.1        ---------MRFHRSLFTLLPALLLVTGTEAGPIAYA--ICQGGCATVVVACYAAAGAVWGTNPWLAAGVSVPAAIVGCNSAYAGCQATCAAVVLLP-------
KAF8463720.1        --------------SIFTLFPALLLVTGTEAGPVGYA--LCQGGCASVVMACYSAAGAVWGTNPWLAAGVPVPAAIVGCNSAYAGCQAKCAVVALSPTP-----
KAJ5116294.1        -----------MKICFLAILAALLLAGDAIAEPAARG--KCHSACEKAVKKCYKKAGHKWGAP-----LANPPAPIIVCNKAFGACQSTCPRS-----------
XP_056783523.1      -----------MKLYLLAIFAAFLLAGEALAAPSIGS--KCHSSCEKAVKKCYKKAGHKWGAP-----LANPPAAVVTCNKAFGACKSTCPK------------
TEA19707.1          -----------MKYTSNLTLLTLALLPAVICGPIEYG--ICQSGCAAILRACYVPKDARWRQ----EGGFPAPPSTKKCDDAFWDCQDRCEIAAYAPGL-----
TDZ35453.1          -----------MKYTSNLTLLALALLPAVICGPIEYG--ICQSGCAAILRACYVPKDARWRQ----EGGFPAPPSTKKCDDAFWDCQDRCEIVAYAPGL-----
TDZ41239.1          -----------MKFTSNLTLLALALLPAVICGPIEYG--ICQSGCAAILRACYVPKGARWRE----EGGFPAPPSTKKCDDAFWDCQDRCEIVAYAPGL-----
XP_049156355.1      ------------MHQAKVSMVVMAIATCATAGPWTYA--GCQSACAAGTLWIPAFSTAA-----------------------YASCQSYCAYFLLAPTP-----
KAI2778314.1        ------------MHQAKVSMVVMAIATCATAGPWTYA--GCQSACAAGTLWIPVFSTPA-----------------------YASCQSYCAYYLLAPTP-----
XP_047795458.1      ------------MHFGKVSAVVMAVATCATAGPLAYA--ACQSACAGATLWVPFFGIPA-----------------------YAGCQSSCAHLLLAPTP-----
UKZ60045.1          ------------MKLTNITLAAIVLPGTAMAGPLAYA--ACQAACST-TLAAGPAGVAL-----------------------YAACQSACAPLLVMPCP-----
PTB76422.1          ----------MKMKFVNIASSVIVLSGIAMAGPVAYA--TCQAACAV-SLAA-PGGVAI-----------------------YAACQSHCAALLVAPCP-----
KAH8759482.1        ------------MRFTTLALVLAGIT-PAIGGPAVSV--ACHVACATALTACIASGFYFPPM-------------IAQCQAAYGVCQLACTSSAFLPTP-----
KAH9242426.1        ---------MKPETLAVALATAKLVMGNMLFGPMIREYAACQMNCATLAYACYIVMEKEWVT------KCCIPPKARACYFAFEECQATCAYPIRFESK-----
KAK2770103.1        ---------MKLNTLTTGLVTVKLAMGNMLFGPMIREYSACQMNCATLAYACYIVVEKEWVT------KCCIPPKARACYFAFEECQATCAYPIRFESK-----
```

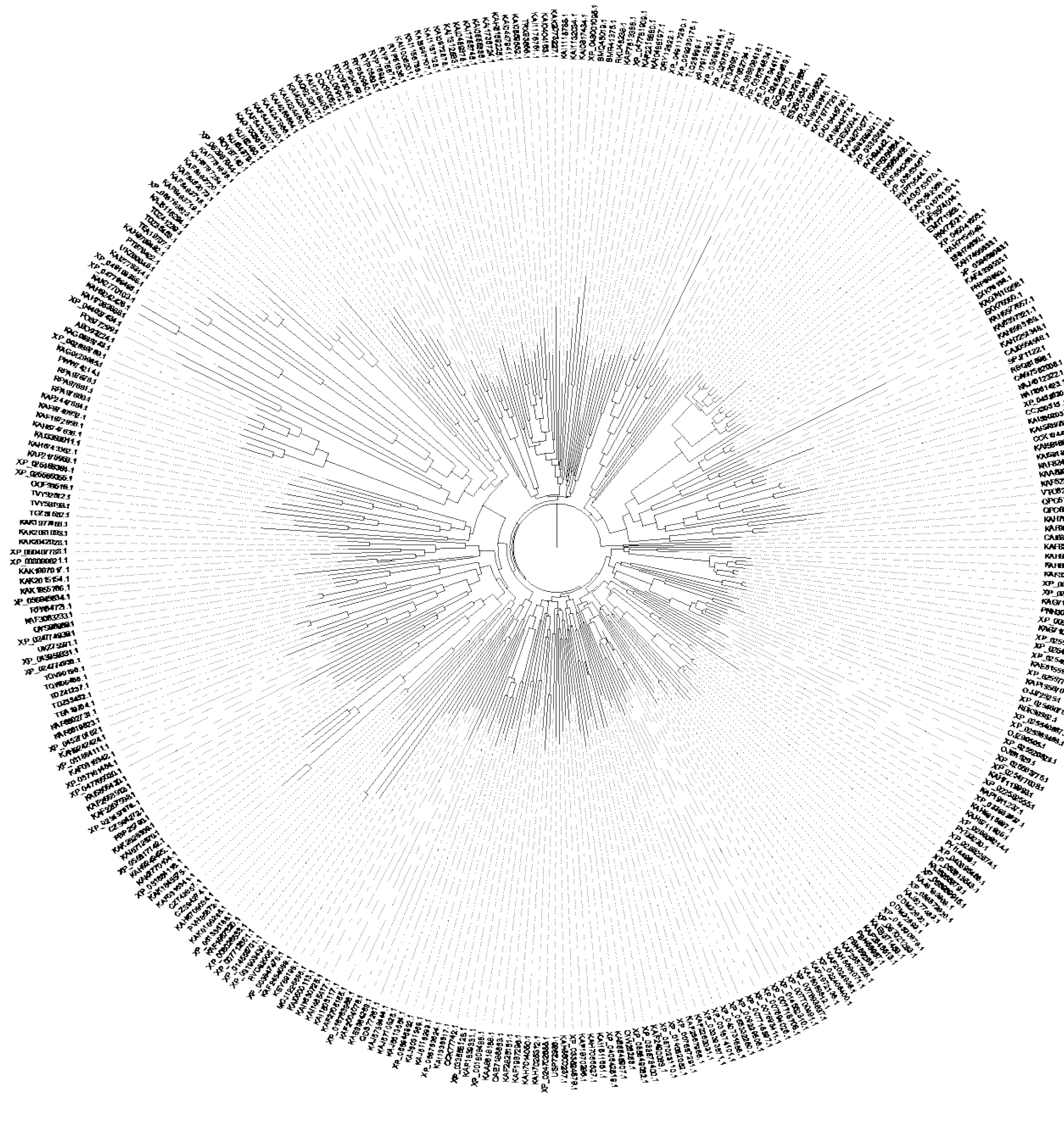

#### Supplementary File S3. Glomeromycota HLPs: 47 sequences

**>CAG8591141.1 2777_t:CDS:2 [Ambispora gerdemannii]**

**MNAKQQLFIAFVVLLCALHGTHAGPIAYAVCQTACNLGWVSCYASAGLVAGTVTGGLGAPLAAIACNVAQGVCMAACVGLIAAPTP**

>CAG8529223.1 4579_t:CDS:2 [Ambispora gerdemannii]

MIPKYFFIVFLVFFCVFQQTVAGPIAYAVCQTACNLKWASCYLSAGLVAGTVTGGLCAPFAALACNAAQGVCMAACAGLLVAPTP

>CAG8495269.1 6133_t:CDS:2 [Ambispora leptoticha]

MNSIFFALLVILCAIHSTYAGPLAYAACQTACNRGWVSCYSSADLVAGATGDLAALPAAVACNVAQGVCMASCVALLTAPSP

>CAG8626826.1 14487_t:CDS:2 [Ambispora leptoticha]

MNSKQQKFFTLLVLLCAIHSTYAGPLTYAACQTACNLGRDSCYALAGFVSGTVFVGLAPPAILACNAVQGFCIASCAGLLSAPIP

>CAG8476239.1 2835_t:CDS:2 [Ambispora leptoticha]

MNSKQQIFFAFLVLLCVIHGTYAGPVAYAVCQTACNLGWVSCYASAGLVAGTVTGGLGAPLAAIACNVAQGVCMAACVGLLTAPTP

>CAG8545532.1 13786_t:CDS:2 [Cetraspora pellucida]

MTRQFIFIILSVLLLSSIITEAGPVAYMSCQSACNAGWVKCYAVMGLVAGTITGGIGAPAGAITCNVAQAACMASCAVLLLAPTL

>CAG8656190.1 3098_t:CDS:2 [Cetraspora pellucida]

MKKLVLAILLVLLFSSITEAGPLAYAICQTACNAGWVACYAAGGLVAGTVTGGVGAPVAAILCNVAQGACMAACAAVILAPTP

>CAG8523360.1 9071_t:CDS:2 [Dentiscutata heterogama]

*MNKKIFAIAIIFISFSLIVNA*GPLAYGACQTLCNIGWVSCYAAFGYVAGTVTAGAGTPLVILGCNAAQGACMVLCAPLLVAPTP

>CAG8607399.1 6810_t:CDS:2 [Dentiscutata heterogama]

*MSKKLIAIIFVLLVLTSSVTVA*GPLCQTACNIGWVKCYAALGFIAGTVTGGTGAPLVVHACNIAQGAFVDYSTTELDT

>CAG8783504.1 11490_t:CDS:2 [Dentiscutata erythropus]

IILGLLDFLRCRSVLHIFNYLSLELLNDQNAYGACQTACNIGWVSCYATFGYVAGTVTAGAGTPLVILGCNAAQGACTVLCAPLLVPTP

>CAG8544857.1 1021_t:CDS:2 [Diversispora eburnea]

MSDGPRAVTGGALPAGAITCNRVQGVCATAGATLPAGAVACNVVQGVCMASCAASFLCPIP

>RHZ44576.1 hypothetical protein Glove_718g57 [Diversispora epigaea]

MAKLSTFVFVLTLLLISFNNVSAGPFAYALCQTACNMGWCSCYAAIGLTAGAATGGVALPAGAVACNVVQGVCMASCAASFLCPIP

>RHZ70068.1 hypothetical protein Glove_275g57 [Diversispora epigaea]

MAKLSIFVFVTILFLISLNNVSAGPITYALCQTACNVGWCTCYGTLGLTAGAATGGAALPAAAIACNVAQGVCMASCAASFFCPVP

>CAJ0758742.1 4610_t:CDS:2 [Entrophospora sp. SA101]

MIQKTSTGQIILVFIVLFFVIATTEAGPLEYGCCQTACNLAWVSCYAASGLTAGTLTGGASLPAAALACNVCQGVCMTVCAGCFLAPTP

>CAG8641126.1 5766_t:CDS:2 [Entrophospora candida]

MDLLFGRKKAPAEIPHEHQRALQRARREMGQAGPLAYGCCQAACNLAWVSCYAASGLTAGAALPTAALACNVCQGVCMTAYAGCFLAPTP

>CAG8540520.1 8579_t:CDS:2 [Entrophospora candida]

MIVTKVNAGLLAYGCCQTACNTAWVSCYAAAGFTAGATAGVALPASVVACNIAQGGCMAVCAACFLAPTP

**>CAG8456294.1 2747_t:CDS:2 [Entrophospora candida]**

**MIQKTSTGQIILVFIVLFFVIATTEAGPLAYGCCQTACNLAWVSCYAASGLTAGKLTGGAALPEAALACNVCQGVCMTVCASCFLAPTP**

>CAG8625918.1 7260_t:CDS:2 [Entrophospora candida]

MIQKTSTGQIILVFIVLFFVIATTEAGPLAYGCCQAACNLAWVSCYAASGLTAGTLTGGAALPTAALACNVCQGVCMTAYAGCFLAPTP

>CAG8439459.1 13992_t:CDS:2 [Funneliformis caledonium]

MAKFSCIFILISVIIFALQPANVSAGPISYAICQSACNVGWVSCYASAGLVAGTVTGGLGAPLAAIACNVAQGVCMAACVGLLTAPTP

**>CAI2165577.1 14144_t:CDS:2 [Funneliformis geosporum]**

**MAKFSHISVLIAMIVFALQPANVNAGPIAYAVCQTACNVGWVSCYASAGLVAGTVTGGLGAPFAAIACNVAQGVCMAACVGLLTAPTP**

>CAI2195945.1 19742_t:CDS:2 [Funneliformis geosporum]

MAKFIYFFVLISLIIFSLQPSNVNAGPIAYASCQTACNLGWGTCYAAAGLIAGTVTGGLATPLAAITCNLAQGACMTGCVVLLAAPTP

>CAI2195120.1 3870_t:CDS:2 [Funneliformis geosporum]

MAKSTSTYCFVLIFLIIFSLQPSSVNAGPLAYAACQTACNLGWVSCYGTAGLAAGTVTGGLATPLAAITCNLAQGACMTGCVVLLAAPTP

>KAF0496523.1 cysteine-rich protein [Gigaspora margarita]

MNQKAFAIVIIFLSFSLIVNAGPIAYGICQTGCNVIWVSCYAAAGFVAGTVTAGAGTPLVIIGCNAAQGVCMAGCVGLLFAPTP

>KAF0517298.1 hypothetical protein F8M41_016942 [Gigaspora margarita]

MKKFITIFLLVLVILPFANAGPVTYTLCQSTCNSGWVSCYAAAGIIGAAIPALGAPSVVALCYIAHGTCMASCAASTLVPTP

>KAF0501944.1 hypothetical protein F8M41_019871 [Gigaspora margarita]

MSKKLIATIFVLLVLISSITEAGPLCQAACCAVLTQCNIAAVSVVGIVTAGIGAPLAVLACNAAYGACIAVCAASPL

**>KAF0496522.1 cysteine-rich protein [Gigaspora margarita]**

**MAQRIFAVFIIFVSLSLLVNAGPLAYGACQTVCNIGWVSCYAAFGYVAGTVTVGVGTPLVILGCNAAQGACMGLCAPLLFLPTP**

>CAG8763021.1 245_t:CDS:2, partial [Gigaspora margarita]

MNRKLIVAILLVLLISSMADAGPLTYILCQSACNAGWVSCYAAAGLVAGTVTGGVGAPAAAILCNVGQGACMAACAASFMTPTP

>KAF0517297.1 hypothetical protein F8M41_016941 [Gigaspora margarita]

MNRKLIVAILLVLLISSMADAGPLTYILCQSACNAGWVSCYAAAGLVAGTVTGGIGAPAAAILCNVGQGACMAACAASFMTPTP

>KAF0507781.1 hypothetical protein F8M41_018921 [Gigaspora margarita]

MNSNLMTSIILILLISSTANAGPIVYIVCQSSCNAGWVACYAAGGLVAGTVTGGLGAPAAAILCNIAQGACMAACAASLLAPTP

>CAG8577285.1 3171_t:CDS:2 [Gigaspora rosea]

MNQKTFAIAIIFLSFSLIVNAGPIAYGLCQTGCNVIWVSCYAAAGFVAGTVTAGAGTPLVILGCNAAQGVCMAGCVALLVAPTP

>RIB10631.1 hypothetical protein C2G38_2205609 [Gigaspora rosea]

MNQRIFAVIIIFINLSLLVNAGPLAYGACQTVCNIGWVSCYAAFGYVAGTVTAGAGTPLVILGCNAAQGACMALCAPLLFAPTP

>RIB08867.1 hypothetical protein C2G38_315484 [Gigaspora rosea]

MKKFITIFLFILVILPFSNAGPVTYTLCQSTCNSGWVSCYAAAGVIGAAIPALGAPSVVALCYIAHGTCMASCAASTLMPTP

>RIB08799.1 hypothetical protein C2G38_1982314 [Gigaspora rosea]

MDSKLVASVILILLLSSTANAGPILYAVCQTACNAGWVACYAAGGLVAGTVTGGLGAPAAAILCNIAQGGCMAACAASLFAPTP
>RIB08800.1 hypothetical protein C2G38_319280 [Gigaspora rosea]
MNSKLITSIILILLLSSIANAGPIVYIACQSSCNAGWVACYAAAGLIAGTVTGGLGAPATAILCNIAQGTCMAACAASLLAPTP
>RIB08866.1 hypothetical protein C2G38_1982239 [Gigaspora rosea]
MDRKLIVAIFLVLLLSSMADAGPLTYILCQSACNAGWVSCYAAAGLVAGTVTGGVGAPAAAILCNVGQGACMAACAASFLTPTP
>RIA99691.1 hypothetical protein C1645_684595 [Glomus cerebriforme]
MILFVFQPADVSAGPIAYAICQTACNLGWVSCYASAGLVAGTITGGLGAPIAAIACNVAQGVCMGACAGLLVAPTP
>GET04047.1 cysteine-rich protein [Rhizophagus clarus]
*MAKISFVILLIMIIFSFQPANVNA*GPIAYAVCQSACNLGWVSCYASAGLVAGTVTGGLGAPLAAIACNVAQAACMAGCVALLTAPSP
>GET03365.1 cysteine-rich protein [Rhizophagus clarus]
MAKISFFVLFMIIIIAFQTINVSAGPIAFAVCQTACNLGWVSCYASAGLVAGTVTAGIGAPFAAIACNVAQGICMGACGSLLVAPTP
**>RGB31413.1 hypothetical protein C1646_764100 [Rhizophagus diaphanus] [Rhizophagus sp. MUCL 43196]**
**MSFIILFIMVIFAFQPANVNAGPIAYAVCQSACNLGWVSCYASAGLAAGTVTAGFGAPLAAIACNVAQGACMAGCVGLLTAPTP**
>RGB42500.1 hypothetical protein C1646_617691 [Rhizophagus diaphanus] [Rhizophagus sp.MUCL 43196]
MAKLSFIALFIIVIFAFQTTNVSAGPIAYAVCQTACNLGWVSCYASAGLVAGTVTGGLGAPFAAIACNVAQGVCMGACAGLLVAPTP
>XP_025164911.1 hypothetical protein GLOIN_2v1734183 [Rhizophagus irregularis DAOM 181602=DAOM 197198]
MSKLSFIALFIIVIFAFQTTNVSAGPIAYAVCQTACNLGWVSCYASAGIVAGTVTGGLGAPFAAIACNVAQGVCMGACAGLLVAPTP
>EXX60164.1 hypothetical protein RirG_182470 [Rhizophagus irregularis DAOM 197198w]
*MAKMSFIILFIMVIFAFQPANVNA*GPIAYAVCQSACNIGWVSCYASAGLVAGTVTGGLGTPFAAIACNVAQGACMAGCIGLLTAPTP
>CAB4412621.1 unnamed protein product [Rhizophagus irregularis]
MAKMSFIILFIMVIFAFQPANVNAGPIAYAVCQSACNIGWVSCYASAGLVAGTITGGLGAPFAAIACNVAQGACMAGCIGLLAAPTP
>CAG8595072.1 2363_t:CDS:2 [Scutellospora calospora]
MNKTTAIFFFVLFLSIANIADAGPITYILCQTACNAGWVSCYAAAGLVAGTVTGGLGAPAAAILCNVAQGACMAACAASFLTPTP
>CAG8771382.1 36661_t:CDS:2, partial [Racocetra persica]
LLLLSTTEAGPLTFVLCQSACNAGWVSCYAGAGLVAGTVTGGLGAPAAALLCNVAQGTCIAACAASFLTPTP
**>CAG8544979.1 30828_t:CDS:2 [Racocetra persica]**
**MTKQLFFAILLSLLLLSTTEAGPITFILCQSACNAGWVSCYAAAGLVAGTVTGGLGAPAAALLCNVAQGACMAACAASFLTPIP**
>CAG8736504.1 7442_t:CDS:2, partial [Racocetra persica]
MPKFALIFLLVLSILSLANAGPVTYTLCQSTCNSGWVACYAASGIIGTICIPGLGSPSVVALCYIAHSACMATCAASALTPAP

```
CAG8591141.1      ---------MNAKQQLFIAFVVLLCALHGTHAGPIAYAVCQTACNLGWVSCYASAGLVAGTVTGGLGAPLAAIACNVAQGVCMAACVGLIAAPTP
CAG8476239.1      ---------MNSKQQIFFAFLVLLCVIHGTYAGPVAYAVCQTACNLGWVSCYASAGLVAGTVTGGLGAPLAAIACNVAQGVCMAACVGLLTAPTP
CAG8495269.1      ------------MNSIFFALLVILCAIHSTYAGPLAYAACQTACNRGWVSCYSSADLVAG-ATGDLAALPAAVACNVAQGVCMASCVALLTAPSP
CAG8626826.1      ---------MNSKQQKFFTLLVLLCAIHSTYAGPLTYAACQTACNLGRDSCYALAGFVSG-TVFVGLAPPAILACNAVQGFCIASCAGLLSAPIP
CAG8529223.1      ----------MIPKYFFIVFLVFFCVFQQTVAGPIAYAVCQTACNLKWASCYLSAGLVAGTVTGGLCAPFAALACNAAQGVCMAACAGLLVAPTP
CAG8439459.1      -------MAKFSCIFILISVIIFALQPANVSAGPISYAICQSACNVGWVSCYASAGLVAGTVTGGLGAPLAAIACNVAQGVCMAACVGLLTAPTP
CAI2165577.1      -------MAKFSHISVLIAMIVFALQPANVNAGPIAYAVCQTACNVGWVSCYASAGLVAGTVTGGLGAPFAAIACNVAQGVCMAACVGLLTAPTP
RGB42500.1        --------MAKLSFIALFIIVIFAFQTTNVSAGPIAYAVCQTACNLGWVSCYASAGLVAGTVTGGLGAPFAAIACNVAQGVCMGACAGLLVAPTP
XP_025164911.1    --------MSKLSFIALFIIVIFAFQTTNVSAGPIAYAVCQTACNLGWVSCYASAGIVAGTVTGGLGAPFAAIACNVAQGVCMGACAGLLVAPTP
RIA99691.1        -------------------MILFVFQPADVSAGPIAYAICQTACNLGWVSCYASAGLVAGTITGGLGAPIAAIACNVAQGVCMGACAGLLVAPTP
EXX60164.1        --------MAKMSFIILFIMVIFAFQPANVNAGPIAYAVCQSACNIGWVSCYASAGLVAGTVTGGLGTPFAAIACNVAQGACMAGCIGLLTAPTP
CAB4412621.1      --------MAKMSFIILFIMVIFAFQPANVNAGPIAYAVCQSACNIGWVSCYASAGLVAGTITGGLGAPFAAIACNVAQGACMAGCIGLLAAPTP
RGB31413.1        -----------MSFIILFIMVIFAFQPANVNAGPIAYAVCQSACNLGWVSCYASAGLAAGTVTAGFGAPLAAIACNVAQGACMAGCVGLLTAPTP
CAI2195945.1      -------MAKFIYFFVLISLIIFSLQPSNVNAGPIAYASCQTACNLGWGTCYAAAGLIAGTVTGGLATPLAAITCNLAQGACMTGCVVLLAAPTP
CAI2195120.1      -----MAKSTSTYCFVLIFLIIFSLQPSSVNAGPLAYAACQTACNLGWVSCYGTAGLAAGTVTGGLATPLAAITCNLAQGACMTGCVVLLAAPTP
CAG8545532.1      ----------MTRQFIFIILSVLLLSSIITEAGPVAYMSCQSACNAGWVKCYAVMGLVAGTITGGIGAPAGAITCNVAQAACMASCAVLLLAPTL
KAF0507781.1      -----------MNSNLMTSIILILLISSTANAGPIVYIVCQSSCNAGWVACYAAGGLVAGTVTGGLGAPAAAILCNIAQGACMAACAASLLAPTP
RIB08800.1        -----------MNSKLITSIILILLLSSIANAGPIVYIACQSSCNAGWVACYAAAGLIAGTVTGGLGAPATAILCNIAQGTCMAACAASLLAPTP
RIB08799.1        -----------MDSKLVASVILILLLSSTANAGPILYAVCQTACNAGWVACYAAGGLVAGTVTGGLGAPAAAILCNIAQGGCMAACAASLFAPTP
CAG8656190.1      ------------MKKLVLAILLVLLFSSITEAGPLAYAICQTACNAGWVACYAAGGLVAGTVTGGVGAPVAAILCNVAQGACMAACAAVILAPTP
CAG8763021.1      -----------MNRKLIVAILLVLLISSMADAGPLTYILCQSACNAGWVSCYAAAGLVAGTVTGGVGAPAAAILCNVGQGACMAACAASFMTPTP
KAF0517297.1      -----------MNRKLIVAILLVLLISSMADAGPLTYILCQSACNAGWVSCYAAAGLVAGTVTGGIGAPAAAILCNVGQGACMAACAASFMTPTP
RIB08866.1        -----------MDRKLIVAIFLVLLLSSMADAGPLTYILCQSACNAGWVSCYAAAGLVAGTVTGGVGAPAAAILCNVGQGACMAACAASFLTPTP
CAG8771382.1      -----------------------LLLLSTTEAGPLTFVLCQSACNAGWVSCYAGAGLVAGTVTGGLGAPAAALLCNVAQGTCIAACAASFLTPTP
CAG8544979.1      -----------MTKQLFFAILLSLLLLSTTEAGPITFILCQSACNAGWVSCYAAAGLVAGTVTGGLGAPAAALLCNVAQGACMAACAASFLTPIP
CAG8595072.1      ----------MNKTTAIFFFVLFLSIANIADAGPITYILCQTACNAGWVSCYAAAGLVAGTVTGGLGAPAAAILCNVAQGACMAACAASFLTPTP
CAG8607399.1      ----------MSKKLIAIIFVLLVLTSSVTVAGP----LCQTACNIGWVKCYAALGFIAGTVTGGTGAPLVVHACNIAQGAFVDYSTTELDT---
KAF0501944.1      ----------MSKKLIATIFVLLVLISSITEAGP----LCQAACCAVLTQCNIAAVSVVGIVTAGIGAPLAVLACNAAYGACIAVCAASPL----
KAF0517298.1      ------------MKKFITIFLLVLVILPFANAGPVTYTLCQSTCNSGWVSCYAAAGIIG-AAIPALGAPSVVALCYIAHGTCMASCAASTLVPTP
RIB08867.1        ------------MKKFITIFLFILVILPFSNAGPVTYTLCQSTCNSGWVSCYAAAGVIG-AAIPALGAPSVVALCYIAHGTCMASCAASTLMPTP
CAG8736504.1      ------------MPKFALIFLLVLSILSLANAGPVTYTLCQSTCNSGWVACYAASGIIGTICIPGLGSPSVVALCYIAHSACMATCAASALTPAP
KAF0496522.1      -----------MAQRIFAVFIIFVSLSLLVNAGPLAYGACQTVCNIGWVSCYAAFGYVAGTVTVGVGTPLVILGCNAAQGACMGLCAPLLFLPTP
RIB10631.1        -----------MNQRIFAVIIIFINLSLLVNAGPLAYGACQTVCNIGWVSCYAAFGYVAGTVTAGAGTPLVILGCNAAQGACMALCAPLLFAPTP
CAG8523360.1      -----------MNKKIFAIAIIFISFSLIVNAGPLAYGACQTLCNIGWVSCYAAFGYVAGTVTAGAGTPLVILGCNAAQGACMVLCAPLLVAPTP
CAG8783504.1      -----IILGLLDFLRCRSVLHIFNYLSLELLNDQNAYGACQTACNIGWVSCYATFGYVAGTVTAGAGTPLVILGCNAAQGACTVLCAPLLVPTP-
KAF0496523.1      -----------MNQKAFAIVIIFLSFSLIVNAGPIAYGICQTGCNVIWVSCYAAAGFVAGTVTAGAGTPLVIIGCNAAQGVCMAGCVGLLFAPTP
CAG8577285.1      -----------MNQKTFAIAIIFLSFSLIVNAGPIAYGLCQTGCNVIWVSCYAAAGFVAGTVTAGAGTPLVILGCNAAQGVCMAGCVALLVAPTP
KAG0733020.1      ------------MKKLIVIYLITALLVSTSYAGLLAYGICQTGCNALAVACYGAAGFTFGTVTGGAAVPPVIAGCNVALGACMAGCVAAGAAPVP
RHZ44576.1        ---------MAKLSTFVFVLTLLLISFNNVSAGPFAYALCQTACNMGWCSCYAAIGLTAGAATGGVALPAGAVACNVVQGVCMASCAASFLCPIP
RHZ70068.1        ---------MAKLSIFVFVTILFLISLNNVSAGPITYALCQTACNVGWCTCYGTLGLTAGAATGGAALPAAAIACNVAQGVCMASCAASFFCPVP
CAG8544857.1      -----------------MSDGPRAVTGGALPAGAITCNRVQGVC------------------ATAGATLPAGAVACNVVQGVCMASCAASFLCPIP
CAJ0758742.1      ------MIQKTSTGQIILVFIVLFFVIATTEAGPLEYGCCQTACNLAWVSCYAASGLTAGTLTGGASLPAAALACNVCQGVCMTVCAGCFLAPTP
CAG8456294.1      ------MIQKTSTGQIILVFIVLFFVIATTEAGPLAYGCCQTACNLAWVSCYAASGLTAGKLTGGAALPEAALACNVCQGVCMTVCASCFLAPTP
CAG8625918.1      ------MIQKTSTGQIILVFIVLFFVIATTEAGPLAYGCCQAACNLAWVSCYAASGLTAGTLTGGAALPTAALACNVCQGVCMTAYAGCFLAPTP
CAG8641126.1      MDLLFGRKKAPAEIPHEHQRALQRARREMGQAGPLAYGCCQAACNLAWVSCYAASGLTAG-----AALPTAALACNVCQGVCMTAYAGCFLAPTP
CAG8540520.1      ------------------------MIVTKVNAGLLAYGCCQTACNTAWVSCYAAAGFTAG-ATAGVALPASVVACNIAQGGCMAVCAACFLAPTP
```

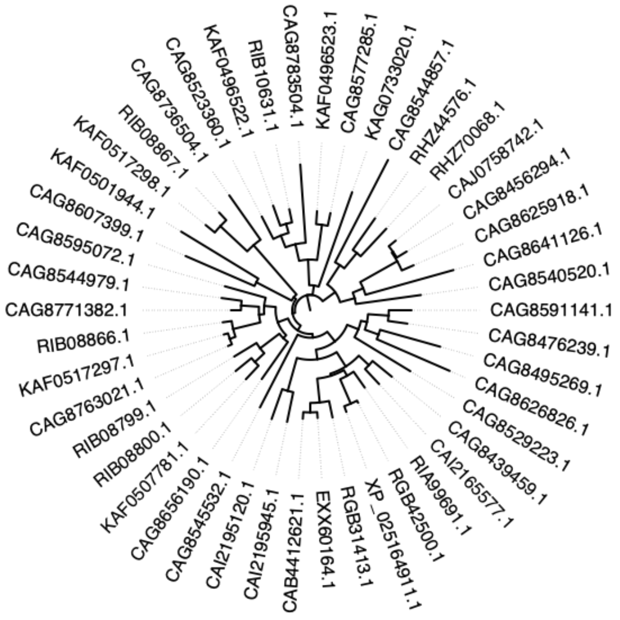
CAG8783504.1
KAF0496523.1
CAG8577285.1
KAG0733020.1
CAG8544857.1
RHZ44576.1
RHZ70068.1
CAJ0758742.1
CAG8456294.1
CAG8625918.1
CAG8641126.1
CAG8540520.1
CAG8591141.1
CAG8476239.1
CAG8495269.1
CAG8626826.1
CAG8529223.1
CAG8439459.1
CAI2165577.1
RIA99691.1
RGB42500.1
XP_025164911.1
RGB31413.1
EXX60164.1
CAB4412621.1
CAI2195945.1
CAI2195120.1
CAG8545532.1
CAG8656190.1
KAF0507781.1
RIB08800.1
RIB08799.1
CAG8763021.1
KAF0517297.1
RIB08866.1
CAG8771382.1
CAG8544979.1
CAG8595072.1
CAG8607399.1
KAF0501944.1
KAF0517298.1
RIB08867.1
CAG8736504.1
CAG8523360.1
KAF0496522.1
RIB10631.1

### Supplementary File S4. Mucoromycota HLPs: 36 sequences

>KAI9487037.1 MAG: hypothetical protein EXX96DRAFT_517501 [Benjaminiella poitrasii]  
MRLKNNSILILLATLCLFFHVSKAGPIAYGVCQTGCNGLAVACYGAAGYTFGTVTAGVGIPVAIVQCNAALGACMGACVAAGLIPLF  
>KAI8349664.1 hypothetical protein BD560DRAFT_410117 [Blakeslea trispora]  
MMKLFLFFAIACLFISQSEAGPISYAICQTGCNALAVACYAAAGAVFGTVTAGAGTPAVILGCNAAQGVCMSACAVVAVVPIP  
**>KAI8355641.1 hypothetical protein EDC96DRAFT_484348 [Choanephora cucurbitarum]**  
**MLKLTFVLLIVCLLMGLSEAGPLAYGICQTGCNALAVACYAGAGFTFGTITAGAGVPAVILGCNAALGTCMAACVAAGLAPIP**  
>OBZ83011.1 hypothetical protein A0J61_08936 [Choanephora cucurbitarum]  
*MFKTTILIVLAASLTLIQA*GPASYGICQAGCNAVAVACYAGAGFTFGTVTAGAGIPASIAACNTALGTCMAHCVMAGICPIP  
>OBZ83014.1 hypothetical protein A0J61_08935 [Choanephora cucurbitarum]  
MGLSEAGPASYGICQAGCNAVAVACYAGAGFTFGTVTAGAGIPAVILGCNAALGTCMTACVAAGLAPIP  
>KAI8355638.1 hypothetical protein EDC96DRAFT_576932 [Choanephora cucurbitarum]  
MKIADAMAGPASYGVCQAGCNAVAVACYAAAGFTFGTVTAGAGIPASIAACNTALGTCMAKCVLAGLCPIP  
>XP_051437264.1 uncharacterized protein B0P05DRAFT_531801 [Gilbertella persicaria]  
MVKFNSEIVVVFGFIGFVSAYGICQAGCIGAFIACEGTVGSLSVWISGLRQLYSTSICARAHRACQEGCVSLGLLPIPY  
**>XP_051439056.1 uncharacterized protein B0P05DRAFT_524115 [Gilbertella persicaria]**  
**MFKAVLILSIILCMSNLVHAGPALYGLCQTGCNTLVVACYAAAGFVFGTITAGAGTPLVILGCNSAQGVCMAACAAALLAPTP**  
>KAI9270929.1 hypothetical protein EDC94DRAFT_596572 [Helicostylum pulchrum]  
MKVVLLSAIFLSLIGVSYCGLLTYGLCQSGCNALAVACYTAGGATFGTVTAGLGIPSVIAGCNSALGACMLGCIAAGCAPTP  
>RUP47770.1 hypothetical protein BC936DRAFT_145348 [Jimgerdemannia flammicorona]  
MLAYSFIICSIMTLLFGTTEAGPLAYGLCQTACNAAWVSCYASLGLVAGWLHEICAGVASALGCNFIQGVCMTSCAASFLLPIP  
>RUP43601.1 cysteine-rich protein [Jimgerdemannia flammicorona]  
MKTPIFATLMTLVIMASLLNGADGSILLYGLCQTACNAAWVSCYAAAGLVAGAATGGLGIPAAAIACNIGQGVCMASCAASFLVPFP  
>KAF9925183.1 hypothetical protein FBU30_004990 [Linnemannia zychae]  
*MQVKLPLILSIVGFANA*GPGLYGICQTGCNALVVACYSAAGATFGTVTAGVGVAPAIIACNAALGTCMAGCVAAGFSPTP  
**>GAN10857.1 zygote-specific protein [Mucor ambiguus]**  
**MNKFYICCLLVLGLISSSYAGPLAYGICQTGCNALVVSCYAAAGFTFGTITAGAGLPAVLVSCNAGLGVCMAGCVAAGLSPTL**  
>EPB81143.1 hypothetical protein HMPREF1544_12152 [Mucor circinelloides 1006PhL]  
MRKFYIACITFLGLISSNYAGLLAYGICQTGCNAVAVACYSAAGFTFGTVTAGAGVPAVILGCNTALGVCMAGCVAAGCAPIP  
>KAF1797016.1 hypothetical protein FB192DRAFT_1401759 [Mucor lusitanicus]  
MNKLFIYCVLVLGLISSACAGPLAYGVCQTGCNALVVSCYAAAGFTFGTVTAGAGIPAALVSCNAALGVCMAGCVAAGFAPTL  
>KAF1797015.1 hypothetical protein FB192DRAFT_1291223 [Mucor lusitanicus]  
*MLKTVVYCLVLLGFLSYVSA*GPLAYGICQTGCNALAVACYSAAGFTFGTVTAGAGVPAVILACNAAQGFCMAGCVAAGCAPIP  
>XP_051460734.1 uncharacterized protein EV154DRAFT_459735 [Mucor mucedo]  
MVNAFLKLCLFVVVVSCLVGQSYAGPLAYGICQTGCNALVVTCYTAAGAVFGTVTAGAGVPAAILGCNAGLGLCMAGCIAAGFAPTP  
>KAG2208688.1 hypothetical protein INT47_007787 [Mucor saturninus]  
MVNAFLKLCLFVVVVSCLVGQSYAGPLAYGICQTGCNALVVTCYTAAGAVFGTVTAGAGVPAAILGCNAGLGLCMAGCVAAGFSPTP  
>KAG2208689.1 hypothetical protein INT47_007788 [Mucor saturninus]  
MKIIYVTQAVPLMLALVQVSQAGPLVYAICQSGCNSLAVACYASAGSVFGTVTAGLGTPAAILGCNTALGTCMAGCIACGCAPTL  
>XP_052938655.1 uncharacterized protein BDF20DRAFT_814749 [Mycotypha africana]  
MNKLFVFFFTIFCLIFNVYAGRLAYGICQTGCNAVATACYAAAGSTFGTVTAGAGIPAAIMACNAALGACMAACIAAGFAPTP  
**>CEP19814.1 hypothetical protein [Parasitella parasitica]**  
**MNKLYFSIFVLLTFFISNSNAGPLAYGICQTGCNALAVSCYAAAGFTFGTITAGAAIPAAIVSCNAALGVCMAGCIAAGFAPTL**  
>KAI8636407.1 hypothetical protein BD408DRAFT_426185 [Parasitella parasitica]  
MNKSFFLISILLLCFTSITHAGPLAYGICQTGCNALAVSCYAAAGFTFGTVTAGTAIPAVIAGCNTSLGLCMAACIAAGFAPTL  
>KAI9337204.1 hypothetical protein BD770DRAFT_331857 [Pilaira anomala]  
MKGSYGGPAAYGVCQSGCNAVAVACYAAAGSTFGTVTLGLGIPFAIMGCNAALGTCMAACVAAGCAPTP  
>KAI9363540.1 cysteine-rich protein [Pilaira anomala]  
*MFLKSALYLTLFSCFLTFTYS*DPLSYGICQTGCNSVAAACYAAAGYTFGTVTAGLCLSPALVACNSALGACMAACVAAGCSPVP  
**>KAI9354162.1 hypothetical protein BD770DRAFT_392522 [Pilaira anomala]**  
***MVFSGNSFKVVLFLAVVLCLVNSSDA*GPISYGICQTGCNALAVSCYAAAGAVFGTVTAGVGVPAAIIGCNVGLGLCMSGCVAAGLSPIP**  
>KAI8976490.1 hypothetical protein BDB01DRAFT_693912, partial [Pilobolus umbonatus]  
YAGPLAYGICQTGCNALVVSCYAAAGFTFGTITAGVGTPAVVVGCNAALGVCMSGCVAAGLAPT  
>KAG0733020.1 hypothetical protein G6F23_013748 [Rhizopus arrhizus]  
MKKLIVIYLITALLVSTSYAGLLAYGICQTGCNALAVACYGAAGFTFGTVTGGAAVPPVIAGCNVALGACMAGCVAAGAAPVP  
>EIE89593.1 hypothetical protein RO3G_14304 [Rhizopus delemar RA 99-880]  
MNKIYLLLLIVAALLGFSAEAGLLSYVICQTGCNTLDATCYAASGLTFGTVTTGAGAPAVALAYNAALGLCMSACVAAGCVPIP  
>EIE76361.1 hypothetical protein RO3G_01065 [Rhizopus delemar RA 99-880]  
MKKLILIYLITALLISTSYAGLLAYGICQTGCNALAVACYGAAGFTFGTVTGGAAVPPVIAACNLALGTCMAGCVAAGAAPVP  
>KAG1436808.1 hypothetical protein G6F56_013401 [Rhizopus delemar]  
MKCKLVFFLVVFTLIQSCVAGPLSYAICQTGCNAVVVACYAGAGFVFGTITGGLGAPPAILACNAGLGVCMSGCVAAGFAPIP  
**>KAG1470267.1 hypothetical protein G6F56_002783 [Rhizopus delemar]**  
**MKIQVFVLLILSFLVCICQAGPISYAICQTGCNAVGVACYSAAGFVFGTITGGLGAPPAVIACNAGLGVCMAACVAAGCTPTP**  
>KAG1142548.1 hypothetical protein G6F38_007664 [Rhizopus arrhizus]  
MKKLIVIYLITALLVSTSYAGLLAYGICQTGCNALAVACYSAAGFTFGTVTGGLAVPPAIACCNVALGTCMAGCVAAGAAPIP  
>RCI02276.1 hypothetical protein CU098_008242 [Rhizopus stolonifer]  
MKYQVVALLIIVFLACSCYAGPLSYGLCQSGCNVVAVACYSAAGFTFGTVSAGLLAPPAIIGCNVALGTCMTACVAAGCAPVP  
>KAI9281927.1 zygote-specific protein-like protein [Sporodiniella umbellata]  
MKAYFFLFILCFISGAYCGPLAYAVCQSGCNALVVSCYAAAGAVFGTVTGGSGVPAAITSCNTGLGICMSACIAAGCAPTL  
>KAG2229776.1 hypothetical protein INT48_006256 [Thamnidium elegans]  
MKLLLPFVLFATLIETSYCGSVVFGIYQQSCISIAAKCYDAAGVSTPLAVIGCNSVLVATSCNSALGVCMAGCVAAGFAPTL  
>KAI8064103.1 hypothetical protein BDF21DRAFT_428225 [Thamnidium elegans]  
MKLLLFLVVFVSLLGLSYCGPLAYGICQTGCNAVAVACYAAAGVTFGTVSAGLCTPLAVIGCNSALGVCMAGCVAAGFAPTL

```
KAI8349664.1     ------MMKLFLFFAIACLFISQSEAGPISYAICQTGCNALAVACYAAAGAVFGTVTAGAGTPAVILGCNAAQGVCMSACAVVAVVPIP-
XP_051439056.1   ------MFKAVLILSIILCMSNLVHAGPALYGLCQTGCNTLVVACYAAAGFVFGTITAGAGTPLVILGCNSAQGVCMAACAAALLAPTP-
KAI8355641.1     ------MLKLTFVLLIVCLLMGLSEAGPLAYGICQTGCNALAVACYAGAGFTFGTITAGAGVPAVILGCNAALGTCMAACVAAGLAPIP-
OBZ83014.1       --------------------MGLSEAGPASYGICQAGCNAVAVACYAGAGFTFGTVTAGAGIPAVILGCNAALGTCMTACVAAGLAPIP-
OBZ83011.1       -------MFKTTILIVLAASLTLIQAGPASYGICQAGCNAVAVACYAGAGFTFGTVTAGAGIPASIAACNTALGTCMAHCVMAGICPIP-
KAI8355638.1     ------------------MKIADAMAGPASYGVCQAGCNAVAVACYAAAGFTFGTVTAGAGIPASIAACNTALGTCMAKCVLAGLCPIP-
EIE89593.1       -----MNKIYLLLLIVAALLGFSAEAGLLSYVICQTGCNTLDATCYAASGLTFGTVTTGAGAPAVALAYNAALGLCMSACVAAGCVPIP-
XP_052938655.1   ------MNKLFVFFFTIFCLIFNVYAGRLAYGICQTGCNAVATACYAAAGSTFGTVTAGAGIPAAIMACNAALGACMAACIAAGFAPTP-
KAI9337204.1     --------------------MKGSYGGPAAYGVCQSGCNAVAVACYAAAGSTFGTVTLGLGIPFAIMGCNAALGTCMAACVAAGCAPTP-
KAI9487037.1     --MRLKNNSILILLATLCLFFHVSKAGPIAYGVCQTGCNGLAVACYGAAGYTFGTVTAGVGIPVAIVQCNAALGACMGACVAAGLIPLF-
KAG2208689.1     ----MKIIYVTQAVPLMLALVQVSQAGPLVYAICQSGCNSLAVACYASAGSVFGTVTAGLGTPAAILGCNTALGTCMAGCIACGCAPTL-
KAI9281927.1     --------MKAYFFLFILCFISGAYCGPLAYAVCQSGCNALVVSCYAAAGAVFGTVTGGSGVPAAITSCNTGLGICMSACIAAGCAPTL-
KAI9270929.1     -------MKVVLLSAIFLSLIGVSYCGLLTYGLCQSGCNALAVACYTAGGATFGTVTAGLGIPSVIAGCNSALGACMLGCIAAGCAPTP-
GAN10857.1       ------MNKFYICCLLVLGLISSSYAGPLAYGICQTGCNALVVSCYAAAGFTFGTITAGAGLPAVLVSCNAGLGVCMAGCVAAGLSPTL-
KAF1797016.1     ------MNKLFIYCVLVLGLISSACAGPLAYGVCQTGCNALVVSCYAAAGFTFGTVTAGAGIPAALVSCNAALGVCMAGCVAAGFAPTL-
KAI8976490.1     ------------------------YAGPLAYGICQTGCNALVVSCYAAAGFTFGTITAGVGTPAVVVGCNAALGVCMSGCVAAGLAPT--
CEP19814.1       -----MNKLYFSIFVLLTFFISNSNAGPLAYGICQTGCNALAVSCYAAAGFTFGTITAGAAIPAAIVSCNAALGVCMAGCIAAGFAPTL-
KAI8636407.1     -----MNKSFFLISILLLCFTSITHAGPLAYGICQTGCNALAVSCYAAAGFTFGTVTAGTAIPAVIAGCNTSLGLCMAACIAAGFAPTL-
EPB81143.1       ------MRKFYIACITFLGLISSNYAGLLAYGICQTGCNAVAVACYSAAGFTFGTVTAGAGVPAVILGCNTALGVCMAGCVAAGCAPIP-
KAF1797015.1     ------MLKTVVYCLVLLGFLSYVSAGPLAYGICQTGCNALAVACYSAAGFTFGTVTAGAGVPAVILACNAAQGFCMAGCVAAGCAPIP-
EIE76361.1       ------MKKLILIYLITALLISTSYAGLLAYGICQTGCNALAVACYGAAGFTFGTVTGGAAVPPVIAACNLALGTCMAGCVAAGAAPVP-
KAG1142548.1     ------MKKLIVIYLITALLVSTSYAGLLAYGICQTGCNALAVACYSAAGFTFGTVTGGLAVPPAIACCNVALGTCMAGCVAAGAAPIP-
XP_051460734.1   --MVNAFLKLCLFVVVVSCLVGQSYAGPLAYGICQTGCNALVVTCYTAAGAVFGTVTAGAGVPAAILGCNAGLGLCMAGCIAAGFAPTP-
KAG2208688.1     --MVNAFLKLCLFVVVVSCLVGQSYAGPLAYGICQTGCNALVVTCYTAAGAVFGTVTAGAGVPAAILGCNAGLGLCMAGCVAAGFSPTP-
KAI9354162.1     MVFSGNSFKVVLFLAVVLCLVNSSDAGPISYGICQTGCNALAVSCYAAAGAVFGTVTAGVGVPAAIIGCNVGLGLCMSGCVAAGLSPIP-
KAF9925183.1     ---------MQVKLPLILSIVGFANAGPGLYGICQTGCNALVVACYSAAGATFGTVTAGVGVAPAIIACNAALGTCMAGCVAAGFSPTP-
KAG1436808.1     ------MKCKLVFFLVVFTLIQSCVAGPLSYAICQTGCNAVVVACYAGAGFVFGTITGGLGAPPAILACNAGLGVCMSGCVAAGFAPIP-
KAG1470267.1     ------MKIQVFVLLILSFLVCICQAGPISYAICQTGCNAVGVACYSAAGFVFGTITGGLGAPPAVIACNAGLGVCMAACVAAGCTPTP-
RCI02276.1       ------MKYQVVALLIIVFLACSCYAGPLSYGLCQSGCNVVAVACYSAAGFTFGTVSAGLLAPPAIIGCNVALGTCMTACVAAGCAPVP-
KAG2229776.1     -------MKLLLPFVLFATLIETSYCGSVVFGIYQQSCISIAAKCYDAAGVSTPLAVIGCNSVLVATSCNSALGVCMAGCVAAGFAPTL-
KAI8064103.1     -------MKLLLFLVVFVSLLGLSYCGPLAYGICQTGCNAVAVACYAAAGVTFGTVSAGLCTPLAVIGCNSALGVCMAGCVAAGFAPTL-
KAI9363540.1     -----MFLKSALYLTLFSCFLTFTYSDPLSYGICQTGCNSVAAACYAAAGYTFGTVTAGLCLSPALVACNSALGACMAACVAAGCSPVP-
RUP47770.1       -----MLAYSFIICSIMTLLFGTTEAGPLAYGLCQTACNAAWVSCYASLGLVAGWLHEICAGVASALGCNFIQGVCMTSCAASFLLPIP-
RUP43601.1       --MKTPIFATLMTLVIMASLLNGADGSILLYGLCQTACNAAWVSCYAAAGLVAGAATGGLGIPAAAIACNIGQGVCMASCAASFLVPFP-
GET04047.1       --MAKISFVILLIMIIFSFQPANVNAGPIAYAVCQSACNLGWVSCYASAGLVAGTVTGGLGAPLAAIACNVAQAACMAGCVALLTAPSP-
GET03365.1       --MAKISFFVLFMIIIIAFQTINVSAGPIAFAVCQTACNLGWVSCYASAGLVAGTVTAGIGAPFAAIACNVAQGICMGACGSLLVAPTP-
XP_051437264.1   ----------MVKFNSEIVVVFGFIGFVSAYGICQAGCIGAFIACEGTVGSLSVWIS-GLRQLYSTSICARAHRACQEGCVSLGLLPIPY
```

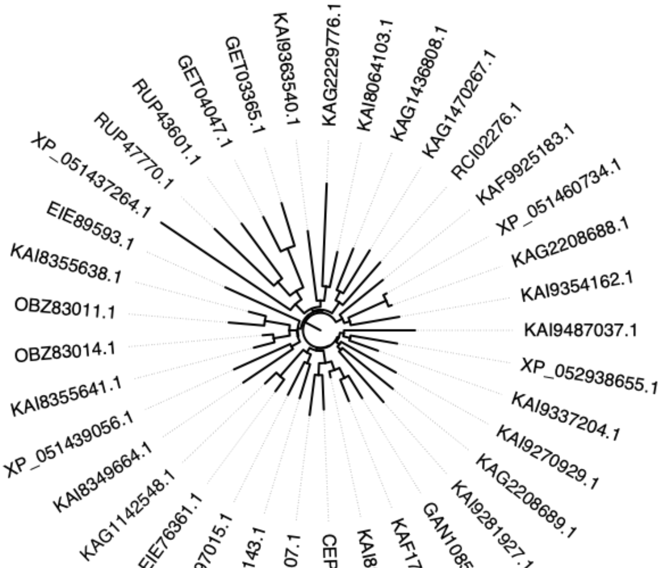

#### Supplementary File S5. Mortierellomycota HLPs: 22 sequences

**>KAG0297619.1 hypothetical protein BGZ98_000539 [Dissophora globulifera]**
**MNCQILFYVVTLLGIVGQTNAGPIAYAVCQSGCNALAVACYSAAGFTFGTVTGGTGTPLVIVGCNTALGTCMAACIAAGFAPTP**
>KAI8605814.1 hypothetical protein EDD21DRAFT_362605 [Dissophora ornata]
MLYYLVVLLAILGVSNAGPLAYGICQTGCNSLVVACYAAAGYTFGTVTGGAAIPAVIANCNIGLGVCMTACVAAGCAPTL
>KAI8601253.1 hypothetical protein EDD21DRAFT_112568 [Dissophora ornata]
MFLRQKLAAALLLLAIAPSPAFGGLITYAVCQSYCNVLAVGCYTFFGFTFGTITVGLGTPLVILGCNAMLGTCMSTCILLGFTPTP
>KAF9998262.1 hypothetical protein BGZ79_008065 [Entomortierella chlamydospora]
MSHTSYTKSDAGPLVYGICQSGCNAIVVACYAAAGFTFGTITAGLGTPAAIVGCNAGLGTCMVACVAAGFAPTP
>KAF9998261.1 hypothetical protein BGZ79_008064 [Entomortierella chlamydospora]
MYKSIKAFLFQLFNLLAILGLSSAGPIAYGICQTGCNALAVACYAAAGFTFGTVTAGAGIPAAIIGCNAALGTCMAAYAVALLASTP
>XP_051411982.1 uncharacterized protein BC939DRAFT_397347 [Gamsiella multidivaricata]
MKYTMLRHLAVVCTILGVSNAGPLAYGICQSGCNALVVACYAAAGFTFGTITAGAAVPAVIAGCNAALGTYMVGCVAAGCAPTP
>KAG0044043.1 hypothetical protein BGZ83_010726 [Gryganskiella cystojenkinii]
MSSRAIFYFTVIFCMILGLANAGPLAYGICQSGCNALVVACYSGAGFTFGTVTGGAGIPAVIVACNAALGTCMVSCVAAGFAPTV
>KAH7042872.1 hypothetical protein BKA57DRAFT_483025 [Linnemannia elongata]
MSSCAFSLCASCALFLLSLPRPLAYGICQTGCNSLAVACYGAAGFTFGTLTAGAGIPAVIVACNAGLGTCMVGCIAAGFAPTL
>KAH7054856.1 hypothetical protein BKA57DRAFT_389634, partial [Linnemannia elongata]
LVFLSILGLTTAGPLAYGICQTGCNAVAVACYAAAGFTFGTVTAGAGIPAVIIGCNAALGTCMVACVAAGFAPTP
**>OAQ22948.1 cysteine-rich protein [Linnemannia elongata AG-77]**
***MIFRSILTYVVILFMIFGLTNA*GPLAYGICQTGCNSLAVACYGAAGFTFGTLTAGAGIPAVIVACNAGLGTCMVGCIAAGFAPTL**
>KAF9276334.1 hypothetical protein BGZ88_001800 [Linnemannia elongata]
MTSRSIFSYVMVLFAFLGLANAGPLAYGICQTGCNALAVACYSAAGVTFGTVTAGAGIPAAVVACNTALGTCMVACVAAGCAPTP
>KAH7054791.1 hypothetical protein BKA57DRAFT_405270 [Linnemannia elongata]
MTSRYNLFYVVLFFMILGLANAGPLAYGICQSGCNSLVVACYAAAGVTFGTITAGAGIPAAVVACNTALGTCMVACVAAGCAPTL
**>XP_021879085.1 hypothetical protein BCR41DRAFT_358057 [Lobosporangium transversale]**
**MLSRKSFVVLLAMVGLSNAGPLAYGICQSGCNALVVACYAGAGFTFGTVTAGAGVPAAIVACNAGLGTCMVGCIAAGFTPTP**
>XP_021879084.1 cysteine-rich protein [Lobosporangium transversale]
MLSRKSFVVLLAMLGFSNAGPLAYGICQTGCNALVVACYAGAGFTFGTVTAGAGLPVAIAACNAGLGTCMVGCVAAGLTPIP
>KAF9352520.1 hypothetical protein BGX26_009688, partial [Mortierella sp. AD094]
MSQYLIRAFIVQLFVLLSILGLSSAGPLAYGLCQTGCNALAVACYAAAGFTFGTVTAGAGIPAVIAGCNAALGTCM
**>KAF9114376.1 hypothetical protein BGX27_011009 [Mortierella sp. AM989]**
**MSYLFTYLIVLLTILGLSSAGPLAYGICQTGCNGLAVACYTAAGFTFGTLTAGLGIPAVIVGCNTGLGTCMVACVVAGFAPTP**
>KAG0209727.1 hypothetical protein BGX28_010021 [Mortierella sp. GBA30]
MLIGLSNAGPLAYGICQTGCNALAVACYAAAGFTFGTVTAGAGIPAVIVGCNAGLGTCMVACVAAILAPTP
**>KAF9951023.1 hypothetical protein BGZ72_007357 [Mortierella alpina]**
**MKISYPYSLIALLAILTKSEAGPAAYGICQTGCNGLAVACYGAGGFVFGTVTAGAGIPAAVVACNVGLGACMAGCVLAGLAPTP**
>KAG0076862.1 hypothetical protein BGZ92_002214 [Podila epicladia]
MSYILATAGPLAYGICQTGCNAVVVACYSAAGFTFGTVTAGAGIPAVIVGCNTGLGVCMAGCIAAGFAPTL
>KAG0088707.1 hypothetical protein BGZ92_005820 [Podila epicladia]
MFLYCLLILLTILCQANAGPLAYGICQTGCNAIAVACYAAAGFTFGTVTAGAGIPAVIAGCNTALGVCMAACVAAGLAPTP
>KAG0018422.1 hypothetical protein BGZ81_010247 [Podila clonocystis]
VFFYYLCIFLTIIGLATAGPLAYGICQTGCNAVAVACYAAAGFTFGTVTGGAGIPAVIAGCNTALGVCMAACIAAGCAPTP
**>KAI9238970.1 MAG: hypothetical protein BYD32DRAFT_248401 [Podila humilis]**
**MKFQAILYYVVLFLMIIGLTNAGPLAYGICQTGCNAVVVACYTGAGATFGTVTAGAGIPAAIIGCNSALGVCMAACVAAGFAPTL**

```
XP_021879085.1 ------MLSRKSFVVLLAMVGLSNAGPLAYGICQSGCNALVVACYAGAGFTFGTVTAGAGVPAAIVACNAGLGTCMVGCIAAGFTPTP
XP_021879084.1 ------MLSRKSFVVLLAMLGFSNAGPLAYGICQTGCNALVVACYAGAGFTFGTVTAGAGLPVAIAACNAGLGTCMVGCVAAGLTPIP
KAF9951023.1   ----MKISYPYSLIALLAILTKSEAGPAAYGICQTGCNGLAVACYGAGGFVFGTVTAGAGIPAAVVACNVGLGACMAGCVLAGLAPTP
KAI8605814.1   --------MLYYLVVLLAILGVSNAGPLAYGICQTGCNSLVVACYAAAGYTFGTVTGGAAIPAVIANCNIGLGVCMTACVAAGCAPTL
XP_051411982.1 ----MKYTMLRHLAVVCTILGVSNAGPLAYGICQSGCNALVVACYAAAGFTFGTITAGAAVPAVIAGCNAALGTYMVGCVAAGCAPTP
KAF9276334.1   ---MTSRSIFSYVMVLFAFLGLANAGPLAYGICQTGCNALAVACYSAAGVTFGTVTAGAGIPAAVVACNTALGTCMVACVAAGCAPTP
KAH7054791.1   ---MTSRYNLFYVVLFFMILGLANAGPLAYGICQSGCNSLVVACYAAAGVTFGTITAGAGIPAAVVACNTALGTCMVACVAAGCAPTL
KAG0044043.1   ---MSSRAIFYFTVIFCMILGLANAGPLAYGICQSGCNALVVACYSGAGFTFGTVTGGAGIPAVIVACNAALGTCMVSCVAAGFAPTV
KAH7042872.1   -----MSSCAFSLCASCALFLLSLPRPLAYGICQTGCNSLAVACYGAAGFTFGTLTAGAGIPAVIVACNAGLGTCMVGCIAAGFAPTL
OAQ22948.1     ---MIFRSILTYVVILFMIFGLTNAGPLAYGICQTGCNSLAVACYGAAGFTFGTLTAGAGIPAVIVACNAGLGTCMVGCIAAGFAPTL
KAG0209727.1   -----------------MLIGLSNAGPLAYGICQTGCNALAVACYAAAGFTFGTVTAGAGIPAVIVGCNAGLGTCMVACVAAILAPTP
KAF9998261.1   -MYKSIKAFLFQLFNLLAILGLSSAGPIAYGICQTGCNALAVACYAAAGFTFGTVTAGAGIPAAIIGCNAALGTCMAAYAVALLASTP
KAF9352520.1   MSQYLIRAFIVQLFVLLSILGLSSAGPLAYGLCQTGCNALAVACYAAAGFTFGTVTAGAGIPAVIAGCNAALGTCM------------
KAG0076862.1   -----------------MSYILATAGPLAYGICQTGCNAVVVACYSAAGFTFGTVTAGAGIPAVIVGCNTGLGVCMAGCIAAGFAPTL
KAI9238970.1   ---MKFQAILYYVVLFLMIIGLTNAGPLAYGICQTGCNAVVVACYTGAGATFGTVTAGAGIPAAIIGCNSALGVCMAACVAAGFAPTL
KAG0088707.1   -------MFLYCLLILLTILCQANAGPLAYGICQTGCNAIAVACYAAAGFTFGTVTAGAGIPAVIAGCNTALGVCMAACVAAGLAPTP
KAG0018422.1   -------VFFYYLCIFLTIIGLATAGPLAYGICQTGCNAVAVACYAAAGFTFGTVTGGAGIPAVIAGCNTALGVCMAACIAAGCAPTP
KAH7054856.1   -------------LVFLSILGLTTAGPLAYGICQTGCNAVAVACYAAAGFTFGTVTAGAGIPAVIIGCNAALGTCMVACVAAGFAPTP
KAF9114376.1   -----MSYLFTYLIVLLTILGLSSAGPLAYGICQTGCNGLAVACYTAAGFTFGTLTAGLGIPAVIVGCNTGLGTCMVACVVAGFAPTP
KAF9998262.1   --------------MSHTSYTKSDAGPLVYGICQSGCNAIVVACYAAAGFTFGTITAGLGTPAAIVGCNAGLGTCMVACVAAGFAPTP
KAG0297619.1   ----MNCQILFYVVTLLGIVGQTNAGPIAYAVCQSGCNALAVACYSAAGFTFGTVTGGTGTPLVIVGCNTALGTCMAACIAAGFAPTP
KAI8601253.1   --MFLRQKLAAALLLLAIAPSPAFGGLITYAVCQSYCNVLAVGCYTFFGFTFGTITVGLGTPLVILGCNAMLGTCMSTCILLGFTPTP
```

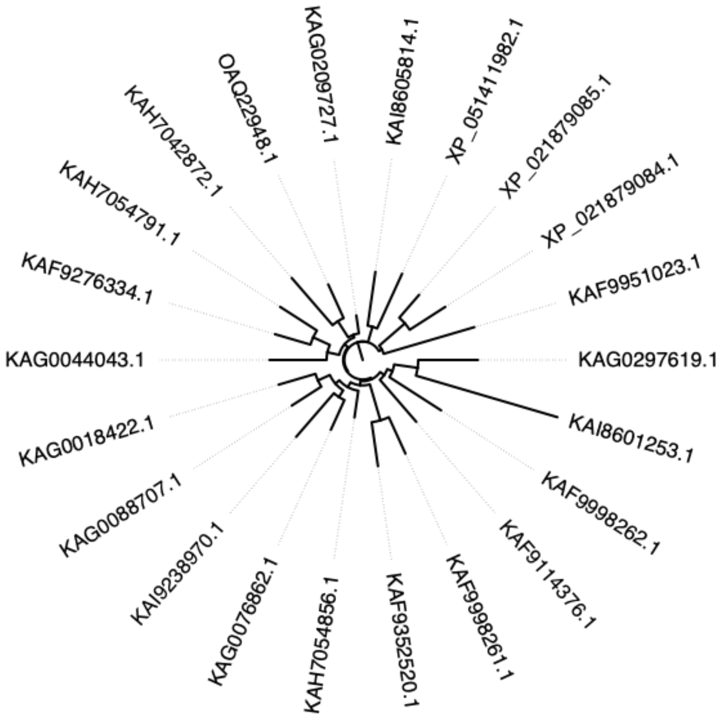
KAG0209727.1
KAI8605814.1
XP_051411982.1
OAQ22948.1
XP_021879085.1
KAH7042872.1
XP_021879084.1
KAH7054791.1
KAF9951023.1
KAF9276334.1
KAG0044043.1
KAG0297619.1
KAG0018422.1
KAI8601253.1
KAG0088707.1
KAF9998262.1
KAI9238970.1
KAF9114376.1
KAG0076862.1
KAF9998261.1
KAH7054856.1
KAF9352520.1

#### Supplementary File S6. Other phyla and sub-phyla HLPs: 46 sequences

##### Pucciniomycotina

>XP_066824081.1 uncharacterized protein P389DRAFT_65519 [Cystobasidium minutum MCA 4210]
MKFSILTTFFTLAALSTQAVAGPALYGICQSGCASVVCACYSAAGFTFGTVVAGPATPAVILACNSAFGACSAKCALVTMAAPTLVTMAAPTTFR
>ORY89663.1 hypothetical protein BCR35DRAFT_261989, partial [Leucosporidium creatinivorum]
QRMKLLTLVAALLAASPLAVQGGPLAYAGCQAGCAGLVVACYSAAGMVFGTVVASAAAPPAILACNAAFGSCQAACAVALLAPTP
>GAA5916100.1 hypothetical protein JCM6882_003936 [Rhodosporidiobolus microsporus]
MKTSFLLVAALALFVNSANGGPVAYGLCQAGCSAPTVACYSAAGAVFGTVTVGVGTPAAILACNGAFGTCCATCASVALFAPTP
>BGP16525.1 hypothetical protein JCM10213v2_004527 [Rhodosporidiobolus nylandii]
MKTSTVAGILLASATTVHGGPLAYAACQAGCAGLQVACYAAAGFTFGTVTAGAGTPAAILACNSAFGACYAACVPALVAPTP
>GAA5985469.1 hypothetical protein JCM11641_007078 [Rhodosporidiobolus odoratus]
MKISTTTLAAVAGLIISAPSAAAGPLSYGICQAGCAAVVVACYSAAGAVFGTVIAGVGTPVAILACNSAFGTCSSACVAAGCLPIP
>GAA5985472.1 hypothetical protein JCM11641_007079 [Rhodosporidiobolus odoratus]
MKPSCILPAAFALVAFASSVEAGPIAYGICQAGCAGIVVACYAAAGFTFGTVTAGAGTPAAIIACNGAFGSCQAACAAIALAPTP
>GAA6001600.1 hypothetical protein JCM10207_006745 [Rhodosporidiobolus poonsookiae]
MKPNSLFLAVFLASFAHLAYGGPLAYAICQAGCAAVVVACYSAAGFTFGTVVAGPATPAVLLACNAAQGTCYAACAVSALAPVP
>GAA6028274.1 hypothetical protein JCM8097_006951 [Rhodosporidiobolus ruineniae]
MKLSFAVLVSALAFASSAQGLLLGYGICQAGCASLVVACYSAAGAVFGCVAAVAAPPAILACNSAFGSCQAACVVAGLAPIP
>GAA6052904.1 hypothetical protein JCM3770_004401 [Rhodotorula araucariae]
MHKLVALVALVALLSLTKSASAGPLAYGTCQAGCACLVVACYAAAGAVFGTITAGAGTAPAILKCNSAYGVCQAACAAAAAAPAP
>GAA5833889.1 hypothetical protein JCM9279_001651 [Rhodotorula babjevae]
MLHLVLLFLLALLGAQVAQAGPLAYAACQACCSAGVVTCYGGAGFVFGTVTAGASTPAVILGCNSAFGACSSACAWMLLAPTP
>GAA5833885.1 hypothetical protein JCM9279_001650 [Rhodotorula babjevae]
MLRHGLALALFLLVLAQVAQAGPAAYGACQSGCSALAVGCYAAAGFAYGTVRKGVGAPAAIMACNRALSSCSASYAPLLHAPSA
>TNY24758.1 cysteine-rich protein [Rhodotorula diobovata]
MLAFLPTVVLSLAVAQTGFAGPIAYATCQAGCSTAAVSCYAAAGFVYGTITAGIGTPAAILSCNAVLGTCSAACAVLSLAPTP
>GAA5943749.1 hypothetical protein JCM3775_000941 [Rhodotorula graminis]
MLHLLVLLVVALLGAQVAKAGPLAYAACQACCSAGVVTCYGAAGFVFGTITAGASTPAVILGCNAAFGSCSSACAWMLLAPTP
>GAA5943751.1 hypothetical protein JCM3775_000942 [Rhodotorula graminis]
MLHLVLAVALSLLVLAQVVQAGPAAYGACQSTCSAIAAACYAAAGFTYGTVRKCVGAPAAIIACNRALSSCSSTCAPLLHSPTA
>BGP40412.1 hypothetical protein JCM10449v2_004374 [Rhodotorula kratochvilovae]
MVHFLKLVTALLLAQTASAGPIAYAVCQAGCAGLVVACYTAAGALRAVTADTGTPAAILGCNAAFGSCQAACAAVALAPTP
>XP_016275080.1 uncharacterized protein RHTO_07020 [Rhodotorula toruloides NP11]
MRFSLLNSALLLLAVTTQSVQAGPLAYAVCQAGYSAVVVACYSAAGFTFGTVTAGAAIPIALVKCNAAYGACQAACATAALFAPTP
>GAA5832040.1 hypothetical protein JCM5353_000730 [Sporobolomyces roseus]
MKFSFPRVTLIAAVFANTVQGMAPYGICQSGCAAVVVPCYSAAGFVFGTVAAPAAPATILACNSGFGACSAKCAGLFLAAPA
>GAA5970736.1 hypothetical protein JCM3765_004696 [Sporobolomyces pararoseus]
MKLSLPFAALATLAFAQSVNGGPIAYGICQSGCAAVVVACYSAAGAVFGTVPAAAIAAGSALAGCNAAFGTCSATCASVALLAPIP
>GAA5873123.1 hypothetical protein JCM16303_006949 [Sporobolomyces ruberrimus]
MKYTFLSPLFVTLALSATTVNGGPIVHGLCQAGFSAVVVACYSAAGAVFGCVPAAGLAAGSALLACNSAFGTCSATCATVALLAPTP

##### Entomophthoromycotina

>KAJ9069409.1 hypothetical protein DSO57_1018795 [Entomophthora muscae]
MKTKILILSLSVVFAGPLAYGICQTGCNAIVVACYAAAGTTFGTVTAGNGAPAAVVSCDAALGTCMAACVAAGFAPTP
>KAJ9064622.1 hypothetical protein DSO57_1028539 [Entomophthora muscae]
MKIKILILSLSVVFAGPLAYGICQTGCNAIVVACYAAAGATFGTVTAGIGAPAAVVSCDAALGTCMAACVAAGFAPTP
>KAJ9069407.1 hypothetical protein DSO57_1018793 [Entomophthora muscae]
MKITFHIASAVLGGPLAYGICQTGCNAVVVACYAAGGATFGTVTAGAGVPAVILGCNVALGTCMAGCVAAGLAPTP
>KAJ9069408.1 hypothetical protein DSO57_1018794 [Entomophthora muscae]
MKFIAFVISTVLAGPLAYGICQTGCNAVAVACYTAGGATFGTVTAGAGVPAVILGCNVGLGACMAGCVAAGFAPTP

##### Blastocladiomycota

>KAI9168152.1 hypothetical protein H9P43_007523 [Blastocladiella emersonii ATCC 22665]
MKLSPLATVLFALLVLALLAAGPAHAGPLGMLAYGLCQTGCNTAAVACYSAAGLTFGVSVGTLGPVAMAAAAGCSVAQGSCMAACAAMAIAPTP

##### Chytridiomycota

>KAJ3320696.1 hypothetical protein HDV06_005104 [Boothiomyces sp. JEL0866]
MQIAYVSLISLASAGLLSYGICQTGCNSVVVACYAGAGLTFGTVTAGAGMPAAAIACNAALGVCMTACVAAGCAPVP
>KAJ3313366.1 hypothetical protein HDV04_002172 [Boothiomyces sp. JEL0838]
MKVSTPILLSFVAAGPLSYGICQTGCNTVVVACYAGAGLTFGTVTAGAGMPAAAIACNSALGVCMVACVAAGCAPTP
>KAJ3250057.1 hypothetical protein HK103_004094, partial [Boothiomyces macroporosus]
MTPLTLGGPLAYGICQTGCNAVVVSCYTAAGATFGTVTAGAGVPAIILGCNAGLGVCMAACVAAG
>ORZ40009.1 hypothetical protein BCR44DRAFT_35654 [Catenaria anguillulae PL171]
MRSATIIFTILVTILLCGIQAVNAGLVTYAACQSGCNIAVVACYAAAGVVFGVGSVPICNVQQGLCMATCAAMALTPTP
>KAI8837667.1 hypothetical protein BJ741DRAFT_602937, partial [Chytriomyces cf. hyalinus JEL632]
MKSVALFAVTSLSMAIGAAAGPLPYVICISACNAGWVACYAGAGLVAGTVTAGIGAPAAAIMCNAAQGACMTACGAALLAPDPSWACAL
>KAI8837668.1 hypothetical protein BJ741DRAFT_602939, partial [Chytriomyces cf. hyalinus JEL632]
MKSAALFVATCLTLAVTTVAGPLPYVLCVSACNAGWVSCYAGAGLVAGTVTAGLGAPAAAMMCNAAQAACMTACGGTLLAPDPSWACTVM

```
>KAJ3137369.1 hypothetical protein HDU90_002156 [Geranomyces variabilis]
MIVKFSILAILFTLFTNALAGPAAVVACITACNAGVVICYSGLGFVFGTVTFGIAAPAAAVTCSAAQGACMAACAPLVIAPTP
>KAJ2986305.1 hypothetical protein HDV02_006766, partial [Globomyces sp. JEL0801]
MKLCLTTILASFCIAGPLSYGICQTGCNAVVVACYAGAGLTFGTVTGGAGVPAAALACNAGLGVCMAACVAAG
>KAI8891779.1 hypothetical protein BC833DRAFT_533663 [Globomyces pollinis-pini]
MKLRFTALLASFCYAGPLSYRICQTGCNTVVVACYAGAELTLGSVTAGAGTKLVASKA
>KAI8902834.1 hypothetical protein BC833DRAFT_613799 [Globomyces pollinis-pini]
MKLCLTTILAPFCIAGPLSYGICQTGCNAVVVACYAGAGLTFGTVTGGAGVPAAALACNAGLGVCMAACVAAGCSPTP
>KAI8893339.1 hypothetical protein BC833DRAFT_531893 [Globomyces pollinis-pini]
MGLCFAGPVSYGICQTGCNALAVACYAGAGLTFGTVTGGVGIPAAAAACNSALGFCMASCVVAGCIPSL
>KAI8892977.1 cysteine-rich protein [Globomyces pollinis-pini]
MSLNIIPIMMSLCFSGPISFGICQNGCNALAISCYATVGLTFGMGDVGVPAVAACNSALGFCMNRCVDAGFIPYV
>KAI9324622.1 hypothetical protein BDR26DRAFT_255637 [Obelidium mucronatum]
MRVSFVPAIFLVHQALAGPVAWAACQTACNAGAVYCYATAGLVFGTVTLGAGAAGPVGWWAWFFGGGAAAATAAATACSAAQGVCMAACTPLLIAPTP
>KAJ3092448.1 hypothetical protein HDU96_002710 [Phlyctochytrium bullatum]
MKFPAATVLAGALLAVTLYAHEAHAGPIAMGSCYTACNAGYVTCCISAGVVAGTFTLGLGAPAALVTCSAIQGACMAACT
PLLVAPTL
>KAJ3103909.1 hypothetical protein HK100_004133 [Physocladia obscura]
MDAYKRNLQQKEKKKMSPPLALACVGACNAGWVACYAAGGLIAGTVSGGIGVPALALACNTAQGACMAACPALIAVPDPGWACTIM
>KAJ3085415.1 hypothetical protein HK102_014195 [Quaeritorhiza haematococci]
MASAKNISFFFFLATLLFALSAPQAKAGVVGIVTYSLCQSGCNTAYVACVGAAGFTAGTFTLGIGAPAALIACSCAQGACMAACAAMALAPTP
>KAJ3068446.1 hypothetical protein HDU99_003210 [Rhizoclosmatium hyalinum]
MHVLKRLTIIVVVGTAADVYAGPLAYATCQTACNLGACACYAAAGLTFGTVTLGAVAGGPITWWAWFFGGAAATGTAAATACSAAQGICMAACTPLLVAPT
P
>KAJ3011798.1 hypothetical protein HDU68_001522 [Siphonaria sp. JEL0065]
MRFTSLPLLSLFVLAASTQVSAGPIAWGVCQTACNAGVVVCYAAAGLTFGTVTIVAGPVSWWAWLFGGGATAATGAAAACSAVQGACMSACTPLLIAPTP
>KAI9359667.1 hypothetical protein DFJ73DRAFT_793749 [Zopfochytrium polystomum]
MDGHQLFLLILAIFLTALAPPTTAGPLAYGVCQTGCNALVASCYMAAGAVFGTVTAGLGTAPAILSCNAAHGTCMAACAAAVLLPA
```

#### Rozellomycota

```
>RKP17103.1 cysteine-rich protein, partial [Rozella allomycis CSF55]
LLGIIGLAFGGPISYGTCQAGCASVVVACYAAAGAVFGTVTLGAGAPPALIACSAAYAKCQAVCAGLLLFPTP
>EPZ35264.1 hypothetical protein O9G_000660 [Rozella allomycis CSF55]
MNIRPSLILLLCIIGLALGGPISYGTCQAGCAAVVVACYAAAGAVFGTVTLGVGAPPALVACSAAFGKCQAICAGLLILPTP
```

#### Ustilagomycotina

```
>XP_025355203.1 uncharacterized protein FA14DRAFT_188945, partial [Meira miltonrushii]
MNVKLLVSLVFATTLAQLAIAGPTAYGICQAGCASPSVACYTAAGATFGTVAAAAAPAAILGCNSAF
```

```
XP_025355203.1      ---------MNVKLLVSLVFATTLAQLAIAGPTAYGICQAGCASPSVACYTAAGATFGTV
GAA6028274.1        ----------MKLSFAVLVSALAFASSAQGLLLGYGICQAGCASLVVACYSAAGAVFGCV
BGP40412.1          -----------MVHFLKLVTALLLAQTASAGPIAYAVCQAGCAGLVVACYTAAGALRAVT
XP_066824081.1      ---------MKFSILTTFFTLAALSTQAVAGPALYGICQSGCASVVCACYSAAGFTFGTV
GAA5832040.1        ---------MKFSFPR-VTLIAAVFANTVQGMAPYGICQSGCAAVVVPCYSAAGFVFGTV
GAA5970736.1        --------MKLSLPFAALATLAFAQS-VNGGPIAYGICQSGCAAVVVACYSAAGAVFGTV
GAA5873123.1        --------MKYTFLSPLFVTLALSATTVNGGPIVHGLCQAGFSAVVVACYSAAGAVFGCV
GAA5916100.1        ----------MKTSFLLVAALALFVNSANGGPVAYGLCQAGCSAPTVACYSAAGAVFGTV
XP_016275080.1      --------MRFSLLNSALLLLAVTTQSVQAGPLAYAVCQAGYSAVVVACYSAAGFTFGTV
GAA6001600.1        ---------MKPNSLFLAVFLASFAHLAYGGPLAYAICQAGCAAVVVACYSAAGFTFGTV
ORY89663.1          --------QRMKLLTLVAALLAASPLAVQGGPLAYAGCQAGCAGLVVACYSAAGMVFGTV
BGP16525.1          -----------MKTSTVAGILLASATTVHGGPLAYAACQAGCAGLQVACYAAAGFTFGTV
GAA5985472.1        --------MKPSCILPAAFALVAFASSVEAGPIAYGICQAGCAGIVVACYAAAGFTFGTV
KAJ3320696.1        ----------------MQIAYVSLISLASAGLLSYGICQTGCNSVVVACYAGAGLTFGTV
KAJ3313366.1        ----------------MKVSTPILLSFVAAGPLSYGICQTGCNTVVVACYAGAGLTFGTV
KAJ2986305.1        ---------------MKLCLTTILASFCIAGPLSYGICQTGCNAVVVACYAGAGLTFGTV
KAI8902834.1        ---------------MKLCLTTILAPFCIAGPLSYGICQTGCNAVVVACYAGAGLTFGTV
KAI8891779.1        ---------------MKLRFTALLASFCYAGPLSYRICQTGCNTVVVACYAGAELTLGSV
KAI8893339.1        ------------------------MGLCFAGPVSYGICQTGCNALAVACYAGAGLTFGTV
KAI8892977.1        ---------------MSLNIIPIMMSLCFSGPISFGICQNGCNALAISCYATVGLTFG--
KAJ9069407.1        -----------------MKITFHIASAVLGGPLAYGICQTGCNAVVVACYAAGGATFGTV
KAJ9069408.1        -----------------MKFIAFVISTVLAGPLAYGICQTGCNAVAVACYTAGGATFGTV
KAJ3250057.1        -----------------------MTPLTLGGPLAYGICQTGCNAVVVSCYTAAGATFGTV
KAJ9069409.1        ---------------MKTKILILSLSVVFAGPLAYGICQTGCNAIVVACYAAAGTTFGTV
KAJ9064622.1        ---------------MKIKILILSLSVVFAGPLAYGICQTGCNAIVVACYAAAGATFGTV
GAA5985469.1        -------MKISTTTLAAVAGLIISAPSAAAGPLSYGICQAGCAAVVVACYSAAGAVFGTV
GAA6052904.1        --------MHKLVALVALVALLSLTKSASAGPLAYGTCQAGCACLVVACYAAAGAVFGTI
KAI9359667.1        ------MDGHQLFLLILAIFLTALAPPTTAGPLAYGVCQTGCNALVASCYMAAGAVFGTV
RKP17103.1          --------------------LLGIIGLAFGGPISYGTCQAGCASVVVACYAAAGAVFGTV
EPZ35264.1          -----------MNIRPSLILLLCIIGLALGGPISYGTCQAGCAAVVVACYAAAGAVFGTV
GAA5833889.1        ----------MLHLVLLFLLALLGAQVAQAGPLAYAACQACCSAGVVTCYGGAGFVFGTV
GAA5943749.1        ----------MLHLLVLLVVALLGAQVAKAGPLAYAACQACCSAGVVTCYGAAGFVFGTI
```

```
TNY24758.1          ----------MLAFLPTVVLSLAVAQTGFAGPIAYATCQAGCSTAAVSCYAAAGFVYGTI
GAA5833885.1        ---------MLRHGLALALFLLVLAQVAQAGPAAYGACQSGCSALAVGCYAAAGFAYGTV
GAA5943751.1        ---------MLHLVLAVALSLLVLAQVVQAGPAAYGACQSTCSAIAAACYAAAGFTYGTV
KAI8837667.1        ----------MKSVALFAVTSLSMAIGAAAGPLPYVICISACNAGWVACYAGAGLVAGTV
KAI8837668.1        ----------MKSAALFVATCLTLAVTTVAGPLPYVLCVSACNAGWVSCYAGAGLVAGTV
KAJ3103909.1        --------------MDAYKRNLQQKEKKKMSPPLALACVGACNAGWVACYAAGGLIAGTV
KAJ3137369.1        ----------MIVKFSILAILFTLFTNALAGPAAVVACITACNAGVVICYSGLGFVFGTV
KAJ3092448.1        -----MKFPAATVLAGALLAVTLYAHEAHAGPIAMGSCYTACNAGYVTCCISAGVVAGTF
KAI9324622.1        -------------MRVSFVPAIFLVHQALAGPVAWAACQTACNAGAVYCYATAGLVFGTV
KAJ3068446.1        ---------MHVLKRLTIIVVVGTAADVYAGPLAYATCQTACNLGACACYAAAGLTFGTV
KAJ3011798.1        --------MRFTSLPLLSLFVLAASTQVSAGPIAWGVCQTACNAGVVVCYAAAGLTFGTV
KAI9168152.1        -MKLSPLATVLFALLVLALLAAGPAHAGPLGMLAYGLCQTGCNTAAVACYSAAGLTFGVS
ORZ40009.1          ----MRSATIIFTILVTILLCG--IQAVNAGLVTYAACQSGCNIAVVACYAAAGVVFGVG
KAJ3085415.1        MASAKNISFFFFLATLLFALSAPQAKAGVVGIVTYSLCQSGCNTAYVACVGAAGFTAGTF
                                                         *          *        .


XP_025355203.1      AAA-------------------AAPAAILGCNSAF-------------------------
GAA6028274.1        AAV-------------------AAPPAILACNSAFGSCQAACVVAGLAPIP---------
BGP40412.1          ADT-------------------GTPAAILGCNAAFGSCQAACAAVALAPTP---------
XP_066824081.1      VAG------------------PATPAVILACNSAFGACSAKCALVTMAAPTLVTMAAPTT
GAA5832040.1        AA-------------------PAAPATILACNSGFGACSAKCAGLFLAAPA---------
GAA5970736.1        PAAA-----------------IAAGSALAGCNAAFGTCSATCASVALLAPIP--------
GAA5873123.1        PAAG-----------------LAAGSALLACNSAFGTCSATCATVALLAPTP--------
GAA5916100.1        TVG------------------VGTPAAILACNGAFGTCCATCASVALFAPTP--------
XP_016275080.1      TAG------------------AAIPIALVKCNAAYGACQAACATAALFAPTP--------
GAA6001600.1        VAG------------------PATPAVLLACNAAQGTCYAACAVSALAPVP---------
ORY89663.1          VAS------------------AAAPPAILACNAAFGSCQAACAVALLAPTP---------
BGP16525.1          TAG------------------AGTPAAILACNSAFGACYAACVPALVAPTP---------
GAA5985472.1        TAG------------------AGTPAAIIACNGAFGSCQAACAAIALAPTP---------
KAJ3320696.1        TAG------------------AGMPAAAIACNAALGVCMTACVAAGCAPVP---------
KAJ3313366.1        TAG------------------AGMPAAAIACNSALGVCMVACVAAGCAPTP---------
KAJ2986305.1        TGG------------------AGVPAAALACNAGLGVCMAACVAAG--------------
KAI8902834.1        TGG------------------AGVPAAALACNAGLGVCMAACVAAGCSPTP---------
KAI8891779.1        TAG------------------AGTKLVASKA-----------------------------
KAI8893339.1        TGG------------------VGIPAAAAACNSALGFCMASCVVAGCIPSL---------
KAI8892977.1        MGD------------------VGVP-AVAACNSALGFCMNRCVDAGFIPYV---------
KAJ9069407.1        TAG------------------AGVPAVILGCNVALGTCMAGCVAAGLAPTP---------
KAJ9069408.1        TAG------------------AGVPAVILGCNVGLGACMAGCVAAGFAPTP---------
KAJ3250057.1        TAG------------------AGVPAIILGCNAGLGVCMAACVAAG--------------
KAJ9069409.1        TAG------------------NGAPAAVVSCDAALGTCMAACVAAGFAPTP---------
KAJ9064622.1        TAG------------------IGAPAAVVSCDAALGTCMAACVAAGFAPTP---------
GAA5985469.1        IAG------------------VGTPVAILACNSAFGTCSSACVAAGCLPIP---------
GAA6052904.1        TAG------------------AGTAPAILKCNSAYGVCQAACAAAAAAPAP---------
KAI9359667.1        TAG------------------LGTAPAILSCNAAHGTCMAACAAAVLLPA----------
RKP17103.1          TLG------------------AGAPPALIACSAAYAKCQAVCAGLLLFPTP---------
EPZ35264.1          TLG------------------VGAPPALVACSAAFGKCQAICAGLLILPTP---------
GAA5833889.1        TAG------------------ASTPAVILGCNSAFGACSSACAWMLLAPTP---------
GAA5943749.1        TAG------------------ASTPAVILGCNAAFGSCSSACAWMLLAPTP---------
TNY24758.1          TAG------------------IGTPAAILSCNAVLGTCSAACAVLSLAPTP---------
GAA5833885.1        RKG------------------VGAPAAIMACNRALSSCSASYAPLLHAPSA---------
GAA5943751.1        RKC------------------VGAPAAIIACNRALSSCSSTCAPLLHSPTA---------
KAI8837667.1        TAG------------------IGAPAAAIMCNAAQGACMTACGAALLAPDPSWACAL---
KAI8837668.1        TAG------------------LGAPAAAMMCNAAQAACMTACGGTLLAPDPSWACTVM--
KAJ3103909.1        SGG------------------IGVPALALACNTAQGACMAACPALIAVPDPGWACTIM--
KAJ3137369.1        TFG------------------IAAPAAAVTCSAAQGACMAACAPLVIAPTP---------
KAJ3092448.1        TLG------------------LGAPAALVTCSAIQGACMAACTPLLVAPTL---------
KAI9324622.1        TLGAGAAGPVGWWAWFFGGGAAAATAAATACSAAQGVCMAACTPLLIAPTP---------
KAJ3068446.1        TLGAVAGGPITWWAWFFGGAAATGTAAATACSAAQGICMAACTPLLVAPTP---------
KAJ3011798.1        TI---VAGPVSWWAWLFGGGATAATGAAAACSAVQGACMSACTPLLIAPTP---------
KAI9168152.1        VGTLGP----------------VAMAAAAGCSVAQGSCMAACAAMAIAPTP---------
ORZ40009.1          S--------------------------VPICNVQQGLCMATCAAMALTPTP---------
KAJ3085415.1        TLGIG------------------APAALIACSCAQGACMAACAAMALAPTP---------
                                                  .
```

**Supplementary File S7.**

**Sequences of *Phytophthora* AMPs present in the NCBI protein database.**
**Total: 22 sequences. The sequence shown in bold font was used for BLAST search.**

>KAG6959506.1 hypothetical protein JG688_00010035 [Phytophthora aleatoria]
MNFKTCLAVALVAVVATVATAEDPLYCQAIGCPTLYSEANLAVSKECRDQGKLGDDFHRCCEEQCGSTTPAPA
**>ALC04447.1 phytotoxic protein PcF precursor [Phytophthora cactorum]**
**MNFKTCLAVALVAVVATVATAEDPLYCQAIGCPTLYSEANLAVSKECRDQGKLGDDFHRCCEEQCGSTTPASA**
>ALC04451.1 small cysteine-rich secretory protein SCR82 [Phytophthora capsici]
MNFKTCFAVVLAAVVATVATAEDPLYCQATGCPTLYSESNLAVSRDCRDDFVPNLDESPEDSLARQVKEFHDCCGVKCAKPL
>XP_067783902.1 small cysteine-rich protein SCR91 [Phytophthora cinnamomi]
MNFKTCFALILAAVVAVSVTAEEDQAPKLLYCQATGCPPLYSKANLDVSQKCRNEGHTGDDFHDCCETKCGATPQPGQ
>KAK1945614.1 hypothetical protein P3T76_002662 [Phytophthora citrophthora]
MNFKTCFALVLATVVATVATAEDPLYCQAVGCPTLYSESNLAVSRDCRDDPVPDLANDGGEEDKAKVIKAFHACCETKCAKPL
>XP_002907995.1 PcF and SCR74-like cys-rich secreted peptide, [Phytophthora infestans T30-4]
MNFKTCFAVLLAAVIATVATAEDPLYCQATGCPSLYPEANLALSRECRNQGKVGDDFHTCCTDKCGEKSS
>KAG3118405.1 hypothetical protein PI125_g2929 [Phytophthora idaei]
MNFKTCLAVALVAVVATVATAEDPLYCQAIGCPTLYSEANLAVSKECRDQGKLGDDFHRCCEGQCGSTTPASA
>AAU21448.1 phytotoxin-like SCR74 [Phytophthora infestans]
MNLKIYAIVALTAVLATPITAQQQQLCKAVGCAYEYSHANDVVSKCCQAINPDPVAFHDCCGKSCNTGIPCKSV
>AAU21460.1 phytotoxin-like SCR74 [Phytophthora infestans]
MNFKIYAIVALTAVLATPITAQQQQLCRADGCAYEHSHANKVISKCCQAINPDPVAFYDCCGKSCNTGIPCKSV
>AAU21452.1 phytotoxin-like SCR74 [Phytophthora infestans]
MNLKIYAIVALTAVLATPITAQQQQLCKAVGCAYEYSHANDVVSKCCQAINADPIAFHDCCSKSCNTGSPCKSV
>AAU21440.1 phytotoxin-like SCR74 [Phytophthora infestans]
MNFKIYAIVALTAVLATPITAQQQQLCKADGCAYEYSLANDVVSKCCKAINADPIAFHDCCSKSCNTGSPCKSV
>AAU21455.1 phytotoxin-like SCR74 [Phytophthora infestans]
MNFKIYAIVALTAVLATPITAQQQQLCRADGCAYEYSLANKVISKCCQAINPDPVAFYDCCRISCNMGSPCKAV
>XP_002899161.1 PcF and SCR74-like cys-rich secreted peptide, [Phytophthora infestans T30-4]
MNLGIYAVAALSAMIATTTNAQQLCSDSGCAYVYSESNLKTSECCRKQSMSFNECCRVSCNFVSPC
>AAU21461.1 phytotoxin-like SCR74 [Phytophthora infestans]
MNLKIYAIVALTAVLATPITAQQQQLCKAVGCAYEYSHANDVVSQCCKAINAEPVAFNDCCSKSCNTGSPCKSV
>AAU21462.1 phytotoxin-like SCR74 [Phytophthora infestans]
MNFKIYAIVALTAVLATPITAQQQQLCKAAGCAYEHSSANGVVSQCCKAINAEPVAFNDCCSKSCNTGSPCRSV
>OWZ16433.1 PcF and SCR74-like cys-rich secreted peptide [Phytophthora megakarya]
MNYFAVFLVAVVATTSSAQQQLCSAPGCASRFSDSNVRTSECCKKRPGNFDDCCRMSCNSGSPC
>OWY95656.1 PcF and SCR74-like cys-rich secreted peptide [Phytophthora megakarya]
MNFKTCFAVLLAAAIATSVPANAQQVCRVQACGSPHSDSTVRISDRCKGRSGDFDECCSTSCRFGNPC
>OWY91458.1 PcF and SCR74-like cys-rich secreted peptide [Phytophthora megakarya]
MIATAATAQQYCGARGCAVLYSEDNLRVSKCCASQPRGDFNECCRLSCNIGTPCR
>XP_008893788.1 hypothetical protein PPTG_21087 [Phytophthora nicotianae INRA-310]
MNFKTWFAFVFAAVVATVATAEESSLSGDPQYCQGDGCPPLYCEANIQISQACRNDIAVKGGTFEACCKTKCGASA
>XP_008897797.1 hypothetical protein PPTG_05820 [Phytophthora nicotianae INRA-310]
MDLKTCLHVVLFAVVATAAAAEDPLYKPPPYCDLSTGCQPLYSEANLAVSKACRDEGNTGNDFHICCVEKCGVTFAPNSVTK
>KAL3668925.1 hypothetical protein V7S43_006213 [Phytophthora oleae]
MNFKSCFAIVLAAVVATVATAEDPLYCLATGCPTLYSEANLAVSKDCRDEGNTGKAFHDCCETKCGKPL
>KAE8987386.1 hypothetical protein PR002_g22066 [Phytophthora rubi]
MNFKTCFALVLAAVVATAVSADDSPLYCQAVGCPTLYSEVNLAVSQQCRNEGNTGEAFHSCCVTKCGAPK

```
AAU21461.1     MNLKIYAIVALTAVLATPITA          QQQQLCKAVG-CAYEYSHANDVVSQCCKAINAE--------------PVAFNDCCSKSCNTGSPCKSV
AAU21462.1     MNFKIYAIVALTAVLATPITA          QQQQLCKAAG-CAYEHSSANGVVSQCCKAINAE--------------PVAFNDCCSKSCNTGSPCRSV
AAU21448.1     MNLKIYAIVALTAVLATPITA          QQQQLCKAVG-CAYEYSHANDVVSKCCQAINPD--------------PVAFHDCCGKSCNTGIPCKSV
AAU21452.1     MNLKIYAIVALTAVLATPITA          QQQQLCKAVG-CAYEYSHANDVVSKCCQAINAD--------------PIAFHDCCSKSCNTGSPCKSV
AAU21440.1     MNFKIYAIVALTAVLATPITA          QQQQLCKADG-CAYEYSLANDVVSKCCKAINAD--------------PIAFHDCCSKSCNTGSPCKSV
AAU21460.1     MNFKIYAIVALTAVLATPITA          QQQQLCRADG-CAYEHSHANKVISKCCQAINPD--------------PVAFYDCCGKSCNTGIPCKSV
AAU21455.1     MNFKIYAIVALTAVLATPITA          QQQQLCRADG-CAYEYSLANKVISKCCQAINPD--------------PVAFYDCCRISCNMGSPCKAV
XP_002899161.1 MNLGIYAVAALSAMIATTTNA            QQLCSDSG-CAYVYSESNLKTSECCRKQS-----------------MSFNECCRVSCNFVSPC
OWY91458.1     MIATAATA                         QQYCGARG-CAVLYSEDNLRVSKCCASQPR----------------GDFNECCRLSCNIGTPCR
OWZ16433.1     MNYFAVFLVAVVATTSSA              QQQLCSAPG-CASRFSDSNVRTSECCKKRP-----------------GNFDDCCRMSCNSGSPC
OWY95656.1     MNFKTCFAVLLAAAIATSVPANA          QQVCRVQA-CGSPHSDSTVRISDRCKGRS-----------------GDFDECCSTSCRFGNPC
ALC04447.1     MNFKTCLAVALVAVVATVATA          EDPLYCQAIG-CPTLYSEANLAVSKECRDQ--------------GKLGDDFHRCCEEQCGSTTPASA
KAG3118405.1   MNFKTCLAVALVAVVATVATA          EDPLYCQAIG-CPTLYSEANLAVSKECRDQ--------------GKLGDDFHRCCEGQCGSTTPASA
KAG6959506.1   MNFKTCLAVALVAVVATVATA          EDPLYCQAIG-CPTLYSEANLAVSKECRDQ--------------GKLGDDFHRCCEEQCGSTTPAPA
XP_002907995.1 MNFKTCFAVLLAAVIATVATA          EDPLYCQATG-CPSLYPEANLALSRECRNQ--------------GKVGDDFHTCCTDKCGEKSS
ALC04451.1     MNFKTCFAVVLAAVVATVATA          EDPLYCQATG-CPTLYSESNLAVSRDCRDDFVPNLD-ESPEDSLARQVKEFHDCCGVKCAKPL
KAK1945614.1   MNFKTCFALVLATVVATVATA          EDPLYCQAVG-CPTLYSESNLAVSRDCRDDPVPDLANDGGEEDKAKVIKAFHACCETKCAKPL
KAL3668925.1   MNFKSCFAIVLAAVVATVATA          EDPLYCLATG-CPTLYSEANLAVSKDCRDE--------------GNTGKAFHDCCETKCGKPL
XP_067783902.1 MNFKTCFALILAAVVAVSVTA     EEDQAPKLLYCQATG-CPPLYSKANLDVSQKCRNE--------------GHTGDDFHDCCETKCGATPQPGQ
KAE8987386.1   MNFKTCFALVLAAVVATAVSA         DDSPLYCQAVG-CPTLYSEVNLAVSQQCRNE--------------GNTGEAFHSCCVTKCGAPK
XP_008897797.1 MDLKTCLHVVLFAVVATAAAA     EDPLYKPPPYCDLSTGCQPLYSEANLAVSKACRDE--------------GNTGNDFHICCVEKCGVTFAPNSVTK
XP_008893788.1 MNFKTWFAFVFAAVVATVATA    EESSLSGDPQYCQGDG-CPPLYCEANIQISQACRND-------------IAVKGGTFEACCKTKCGASA
```

**Sequences of fungal Anti-Fungal Proteins present in the NCBI protein database. Total: 99 Sequences. The sequence shown in bold font was used for BLAST search.**

>KAK7983606.1 hypothetical protein PG989_011008 [Apiospora arundinis]
MQFSTVALFLFAVVGAVANPVEGSADGIDAREVQITYDGTCSRSKNECKYKGQNGRPTIVKCPSFANKKVFIQTP
>XP_066727522.1 antifungal protein [Apiospora kogelbergensis]
MQFATAALFLFAAMGAIANPVEGNSEGIDVRATETIFHGTCSKVKDECNYKGEHGKLHHVKCPPKDKCTKKGAKCTFNSKDKKVICH
>XP_066727524.1 uncharacterized protein PG998_011654 [Apiospora kogelbergensis]
MQFSTVTLFLVAAMGAIATPVNSADGVEARAEQGTLLEYTGTCTKAKNECKFKGQTGATTFVKCPTKPNNHRCFRDGNKCTFDSYSRKVVCT
>KAK7994358.1 hypothetical protein PG991_015946 [Apiospora marii]
MQFTTAALSLLAAMGAIAIPLNTTMTLETQGEMGMFITYPGKCTLKTQTCRYKGQNGQTTLAKCPKSPANKRCFGDGHSCSFDSVTRKVSCS
>Aall_KAE8393168.1 antifungal protein precursor [Aspergillus alliaceus]
MKASSFVSLIFIFATALGVAASPTNTNSVPSNDLAMEDEAEIKIKYYGKCFIDEKMKMTKCKYDGPNGRTSFCRCHFKRCTKNGAKCHFDSVNKDCYCS
>Abra_GKZ20515.1 hypothetical protein AbraCBS73388_006092 [Aspergillus brasiliensis]
MHITSIAIVLFAAMGAIANPIAAEADDLLAREAELSKYGGECSLEHNTCTYRKDGKNHVVACPTAANLRCKTDRHHCEYDDHHKTVDCQTPV
>Abra_XP_067479519.1 uncharacterized protein ASPBRDRAFT_41941 [Aspergillus brasiliensis CBS 101740]
MGAIANPIAAEADDLLAREAQLSKYGGECSLEHNTCTYRKDGKNHVVACPTAANLRCKTDRHHCEYDDHHKTVDCQTPV
>Acae_XP_031928216.1 antifungal protein precursor [Aspergillus caelatus]
MHLTTVVLFLFAAMGAVATPIESEAFGLDARAEASTLIKYPGQCTKAKNECKYKSQNKKNTFVKCPSFANKRCTKDGNWCQFDSYSRNVECK
>Acam_XP_024690694.1 uncharacterized protein P168DRAFT_329071 [Aspergillus campestris IBT 28561]
MQLISLASMGLVLFAAVGAVASPVDNNALDVNDNLEVHDEAATLITYNGSCSKKNNSCKYKGQKGKTSFCHCKFKKCGKDGNKCHFDSYSRDCKCI
>Aeuc_XP_025382252.1 uncharacterized protein BO83DRAFT_383499 [Aspergillus eucalypticola CBS 122712]
MKLTSIAIILFAAMGAIANPIAAESDDLLARDVQLSKYGGECSLEHNTCTYRKDGKNHVVSCPSATNLRCKTDRHHCEYDDHHKTVDCQTPV
>Afla_KOC12588.1 hypothetical protein AFLA70_818g000280 [Aspergillus flavus AF70]
MQITTVVLFLFAAMGAVATPIESESFGLDTRAEASTLIKYPGKCSKAKNECKFKGQTKKDTFVKCPSFANKRCTKDGNPCHFDSYSRTVDCK
>Agig_CAA37523.1 antifungal protein, partial [Aspergillus giganteus]
VATPVEADSLTAGGLDARDESAVLATYNGKCYKKDNICKYKAQSGKTAICKCYVKKCPRDGAKCEFDSYKGKCYC
>Ahan_KAF7592939.1 hypothetical protein BBP40_012264 [Aspergillus hancockii]
MKFTSLSLGLVLFSALGAVASPVDAASANRVEVREEAGDAGILIKYDGTCSKKNNECKYKAQNGKTAFCKCQVKKCGSDGGKCFYDSANRQCTCY
>Ajap_XP_025526306.1 antimicrobial peptide [Aspergillus japonicus CBS 114.51]
MKISPVSIGFILLAAMGVAATPLNHAESVGVRSENNVQVKYDGQCRKSENQCRYTAQSGRTAICKCQFRKCSKDGAKCNFDSYNRDCNCY
>Amel_KAK1138769.1 hypothetical protein N8T08_002000 [Aspergillus melleus]
MKLTSLASLGIVLFTAMGTLGSPVDSGSLASNDLDARDEAGIMIRYDGKCSKKDNSCRYKSQNGRTAFCKCQFKRCAKDGNKCHYESYNGNCQCI
>Apho_RDK38155.1 hypothetical protein M752DRAFT_339122 [Aspergillus phoenicis ATCC 13157]
MQLTSIAIILFAAMGAIANPITAEANNLVAREEELSKYGGECSVEHNTCTYLKGGKDHIVSCPSAANLRCKTERHHCEYDEHHKTVDCQTPV
>Aser_KAE8333019.1 antifungal protein [Aspergillus sergii]
MQITSVAIVLFAAMGAVANPIATQSDDLSARDIQLSKFGGECSLKHNTCTYLKGGKNHVVNCGSADNQKCKEERHHCEYDDHHKTVNCQTPV
>Aspa_SMQ11440.1 antifungal protein [Aspergillus spathulatus]
MQITKISLFLFVGVGVVASPIHAESDGLNARAVNAADLEYKGECFTKDNTCKYKIDGKTYLAKCPSAANTKCEKDGNKCTYDSYNRKVKCDFRH
>Aste_XP_024699894.1 antifungal protein precursor [Aspergillus steynii IBT 23096]
MKFLSIASLSLILFTAMGVLGSPIESEALASNDLDARDEAGILIKYPGTCSKKNNNCRYKSQNGRTAFCKCKFKKCAKDGNKCHFDSYNQDCQCI
>Atai_PLN82996.1 hypothetical protein BDW42DRAFT_192584 [Aspergillus taichungensis]
MQLISLASMGLVLFAAVGAVASPVDNNALDIDNNLEVRDEAASLIKYHGVCSKKNNSCKFKGQNGKTSFCHCKFKKCGKENNKCHFDSYNRDCKCI
>Atam_KAE8159784.1 antifungal protein [Aspergillus tamarii]
MQLTSIAIFLFAAMGAVANPIATESDDLDTRDIELSKFGGECSVEHNTCTYLKGGKNHVVKCGSAANKKCKSDRHHCEYDEHHKTVDCQTPV
>Awen_KAI9925351.1 hypothetical protein MW887_006279 [Aspergillus wentii]
MKIISLASFGIALFTALAAATPVEPNTVEARAEQGVLIKYDGKCSKKDNSCKFKGQGGKTTFCHCKFKKVFLPPSDIVDL
>Avir_XP_043122868.1 uncharacterized protein Aspvir_003682 [Aspergillus viridinutans]
MHTSIAIVLIAAMGAIANPIVTESDDLDARDVQLSKFGGECSLKYNTCTYLKGGKNQVVNCGSAANKRCKTDRRHCEYDEYHKTVDFQTPV
>KAI0202405.1 antifungal protein [Astrocystis sublimbata]
MQIINAALFLFAAMGAVATPLEAGSDGLDARGADAVLITYKGKCTKSSNTCKYTGQNGKTTIVSCPTARNLVCTNDNKECTFDSVDKKVTCS
**>PMB64038.1 Cicadin [Beauveria bassiana]**
**MQIISIALSLLAATGAIAAATPEQFEARDGAGAMIKYHGVSKKPPIHVHYGEKLTIDSTSKICTKAKNECKFKGQNGRDTFVKCPSFANKRCTKDYNECSYDSVSRAV**
**VCH**
>KAF0329504.1 antifungal protein [Colletotrichum asianum]
MQIAKIALFLFAAIGVAANPVDIDDASGIDAGVSGGEDMNTFLTYTGKCTRGRNYKEDTCKFKGQKGKTTIVRCPRFANQRVSFDSLSSTQCPRNGSKCTWDSYKRTT
KCNYK
>XP_053031537.1 uncharacterized protein COL26b_011876 [Colletotrichum chrysophilum]
MQIAKVAFFLFAAIGVAANPVDVDGSGIDAGVSGGEDMNTLITYTGKCTRGRNYKEDTCKFKGQKGKTTIVRCPRFANQRVSFDTQYTGAINGPKKNIGQSKNFNATT
A
>KAJ0271918.1 hypothetical protein CBS470a_012941, partial [Colletotrichum nupharicola]
MQITKVALFLFAAIGVAANPVNVDGSGIDVGVSGGEDMNTFITYTGKCSRGRNYKEDTCKFKGQKGKTTIVRCPRFANQR
>TEA21567.1 Antifungal protein [Colletotrichum sidae]
MQFANNFVFLLAAMGAIASPVQPNSNDVGPADLEGDGITYTGKCTRKTNECNFTASRKKKSVKCPSLPNKTCTKDGAKCTYDILSQEIVCY
>KAF4823470.1 Antifungal protein [Colletotrichum tropicale]
MQIAKVAFFLFAAIGVAANPVDVDGSGIDAGVSGGGRNYKEDTCKFKGQKGKTTIVRCPRFANQRLQADYQVQL
>KAF4922418.1 hypothetical protein CGCVW01_v005291 [Colletotrichum viniferum]
MQIVKVALFLFAAIGVAANPVDVDGSGMDVGVSGGEDMNTFITYTGKCTRGRNYKEDTCKFKGQKGKTTIVRCPRFANQRVSFDMLSSIQIHKLNRSDIMVFSSSALG
MAVSAHGTVTSGLPSATTNSFSSLEAAATTTRSHSEA
>KAF4920109.1 Antifungal protein [Colletotrichum viniferum]
MQIVKVALFLFAAIGVAANPVDVDGSGMDVGVSGGEDMNTFITYTGKCTRGRNYKEDTCKFKGQKGKTTIVRCPRFANQRCPRNGSKCTWDSYKRTTKCNYK
>XP_018702882.1 Antifungal protein [Cordyceps fumosorosea ARSEF 2679]
MQITSIALFLLTATGAVAAATPEQFDARDGLSAQIKYHGICTKAKNECKFKGQNGRDTFVKCPNFANKRCTKDHNECSYDSVSRAVVCH
>XP_006667310.1 uncharacterized protein CCM_02093 [Cordyceps militaris CM01]
MQIPTFALLLLTAAAAAMPLSDPLDARDAASAQITYYGICTKANNECKYKNQNGKDTFVKCPRFANKKRQQSRAVPLDSVNVAEPFEHFPRLFVPIDI

>USP79915.1 antifungal protein precursor [Curvularia clavata]
MQLASVFLLSFAALGAVANPINSQGDSIDVRGENNVFITYTGTCTKANNQCRYKAQNGKTAFAKCPKFANKKVHLKHTYVTKLD
>XP_033521083.1 uncharacterized protein P153DRAFT_368761 [Dothidotthia symphoricarpi CBS 119687]
MSAVATPIDSKLNDIDTRGVFITYTGKCDASTQQCRYNGQNGVVTIAKCGIAANKKCTTLPTTGGTCEYDSASKVLTCH
>AWO72254.1 antifungal protein [Epichloe festucae]
MQITVVAVFLLSAMGGVATPINSRINPVDARAETGILITYEGTCSRAKNECKYKNQNNKDTFVKCPSFANKKCTKDNAKCSFDSYSRAVTCH
>Faga_KAF4500017.1 antifungal [Fusarium agapanthi]
MGVVATPIDSPAMALEVRGNLEKRLDYKGTCTRSSNTCRYKGPNGRIAFKKCGTFANQKCTKDGAPCVWQSDKGAGDNADGICLDIGLS
>Faus_KAF5228166.1 hypothetical protein FAUST_11285 [Fusarium austroamericanum]
MKFSSVTLLLFVATGAVATPVDSLPNQLDARDGLFPRRTFPGKCTRSDNRCKYENSNGKTVTISCGTAANKKCTKDNADCVYDDANRSVKCD
>Fave_KAH6970200.1 antifungal protein [Fusarium avenaceum]
MQFSTITLFLAATMGVAASPVDTPAQELDARGNLFPRLDYHGTCTKSTNRCRYINDKKRTVIISCPKFANKKCTKDGNKCTYDSAARSVICR
>Fbul_KAF5965014.1 antifungal protein [Fusarium bulbicola]
MQFSAITIFLVAAMGVAATPIDSPAIALDARGSLEKRLDYKGTCTRSSNTCRYKGPNGRTAFKKCGTFANQKCTKDGAPCVWQSDKGVGGKITCK
>Fchl_KAL4723034.1 hypothetical protein ACLX1H_010275 [Fusarium chlamydosporum]
MQFSSITLLLAAAIGAVATPVDSPPSQLDTRSRLFPRRPYEGTCTRADNRCKYDNSNGKTVTISCGTAANRKCTKDGAKCVYDDADRSVKCD
>Fgra_KAF4995278.1 hypothetical protein FGRMN_5243 [Fusarium graminum]
MQFSTIALVFVAAMGAVATPVDPPAQDLDARGDLYPRLDYWGTCTKANNRCRYKNDKGRTVLQNCPKFTNKKCTKDGNRCKWDSAAKDLICY
>Fgre_CAF3538390.1 unnamed protein product [Fusarium graminearum]
MQFSTIVPLFVAAMGVVATPVNSPAQELDARGNLFPRLEYWGKCTKAENRCKYKNDKGKDVLQNCPKFDNKKCTKDGNSCKWDSASKALTCY
>Flan_GKU13684.1 unnamed protein product, partial [Fusarium langsethiae]
MQISTILPLFVAAMGVVATPIDSPAQELDARGNLFPRLEYWGSCTKAENRCKYKNDKGKEVLQNCPKFDNKKVQSHKVPNSFHTN
>Flon_RGP60307.1 antifungal [Fusarium longipes]
MHLSTITLFFVAAMSAVATPVDSSAQNLDARANLGRRRDFPGTCTKSDNRCKYKNSNDKYVTIACPKFDNKKCTKDGNSCTYDDADRSVKCD
>Fmex_KAF5551368.1 hypothetical protein FMEXI_3486 [Fusarium mexicanum]
MQFSAITLFLVAAMGVAATPIDPPAIALDARGSLEKRLDYKGTCTRSSNTCRYKGPNGRTAFKKCGTFANQKCTKDGAPCVWQSDKGVGGKITCK
>Fnap_KAF5554466.1 hypothetical protein FNAPI_6417 [Fusarium napiforme]
MGVAATPIDSPAMALDARGDLEKRLDYKGTCTKSSNTCRYKGPNGRTTFKKCGTFANQKVQSRGFSAMLN
>Fpro_XP_031085614.1 uncharacterized protein FPRO_15745 [Fusarium proliferatum ET1]
MQLSAITLFLVAAMGVAATPIDSPVMALDARGNLEKRLDYKGTCTKSSNTCRYKGPNGRTAFKKCGTFANQKDGAPCVWQSDKGVGGKIICK
>Fpse_XP_009263607.1 hypothetical protein FPSE_12215 [Fusarium pseudograminearum CS3096]
MQFSTILPLFVAAMGIVATPVNSPAQELDARGNLLPRLEYWGKCTKAENRCKYKNDKGRDVLQNCPKFDNKKCTKDGNSCKWDSASKALTCY
>Fsam_KAL6912397.1 hypothetical protein FSST1_010157 [Fusarium sambucinum]
MQFSTILPLFIAAMGVVATPIDAPAQELDARGNLFPRLEYWGSCTKSDNRCKYKNDKGNSVLQNCPSFDNKKCTKDGNSCKWDSAKKELTCY
>Ftri_KAH7263759.1 hypothetical protein BKA59DRAFT_506774 [Fusarium tricinctum]
MQFSTITLFLAATMGVAATPVDSPAQELDARGNLFPRLDYHGTCTKSTNRCRYINDKKRTVIISCPKFANKKVQSSKMLTDNFY
>Fven_XP_025591605.1 antifungal protein [Fusarium venenatum]
MQFSTIFSLFIAAMGIVATPIDSPTQELDARGNLFPRLEYWGSCTKSNNRCKYKNDDDNSVLQNCLSFNNKKCTKDGNNCKWDSAKKELNYY
>Fver_XP_018757790.1 hypothetical protein FVEG_16817 [Fusarium verticillioides 7600]
MQFSTITLLLLASTGVAATPIYSPAMPLDARRNLEKRLEYKGTCTKSSNTCRYKGPNGRTTFKKCGTFANQKCTKDGAPCVWESEKGVGGKVTCK
>Fxyl_KAG5755042.1 hypothetical protein H9Q70_002367 [Fusarium xylarioides]
MQFSTITLLLVAAMGVAATPIDSPAMALDARGNLEKRLDYKGTCTKSSNTCRYKGPNGRTAFKKCGTFANQKCTKDGASGFENLAGFYMSSNMTVTGICEIVIVGTGP
GTNIGI
>KAH7010366.1 antifungal protein, partial [Ilyonectria destructans]
MQIATATLFLIAAMGAVASPVEPNANGMDARDGIYARITYSGTCTRSDNTCKYKNQNNGNTFIKCPTAANKKCTNDGKAC
SYDSVSKAVTC
>AHA86567.1 MAFP1 [Monascus pilosus]
MQFTKIAIFLFAAMGAVANPIAAESGDLDVRDVQLSKYGGECSLQHNTCTYLKGGKNQVVHCGSAANQKCKSDRHHCEYDEHHKTVNCQTPV
>PFH58491.1 hypothetical protein XA68_13599 [Ophiocordyceps unilateralis]
MVSLYSIAFALLAVSGAVASPAGDSPSLEQRDAAITYNGKCSAKANTCRYVGQSGRPSICKCYVKKCSGDGKACHYDSYK
NQCLCV
>KAJ9304236.1 hypothetical protein DTO217A2_6319 [Paecilomyces variotii]
MQITKISLFLFAAIAAVANPIDAESDGVVERDVDAADITYTGECFRKNNECRYVANGKTHYVKCPSKFANKRCQMDKHKC
TFDSYSRVVNCNA
>KAH8696995.1 antifungal protein [Phaeosphaeriaceae sp. PMI808]
MQITTTVLFLSAAIGVIATPIQPERTSVDVRDNASSRIEYTGKCTRSTNICRYEGQNNKIILISCPSAANLRCTNDGRAC
SYESSTKKVTCG
>Psp_KAJ5413027.1 hypothetical protein N7465_005332 [Penicillium sp. CMV-2018d]
MQITSIAIVLFAAMGAVANPIPTESDDLVARDVQLSKFGGECSLKHNTCSYRKGGKTRIVNCGSAANKKCKSDRHHCEYDEHHKRVDCQTPV
>Psp_KAJ5455444.1 antifungal protein-domain-containing protein [Penicillium sp. IBT 31633x]
MQITTIALFFFAAMGAVANPIASEASIASEANELDARAEAGTLISYSGKCYKSKNECKFKGQNGKTTFVKCPKFANKKCTKDGASCKYDSYDGKVTCN
>Psp_KAJ6113509.1 hypothetical protein N7523_006826 [Penicillium sp. IBT 18751x]
MQITSIAIVLFAAMGAIANPIGTESDDLDARDVQLSKYGGECSLTHNTCTYLKGGKHIAVNCGSAANKRCKTDRHHCEYDEHHKTVDCQTPV
>Psp_KAJ5466821.1 Antifungal protein [Penicillium sp. IBT 31633x]
MQITRIAIVLFAAMGAVANPIAAESNSLDTQAFGSKYGGECSKEHNTCTYRKDGKDHKVKCPSADNLKCKTDRHHCEYDEHHKKVDCQTPV
>Patr_XP_056739537.1 Peptidyl-prolyl cis-trans isomerase D [Penicillium atrosanguineum]
MQITKVSLFLFAAMGAIASPIDAESDGLNARVENAANIEYTGKCVAKDNNCRYGIGGKTHLVKCPSAANTKCEKDGNKCTYDSYNGKVKCDFRH
>Pbra_CEJ62478.1 Putative Antifungal protein [Penicillium brasilianum]
MQVAKISLFLFAAMGTVASPIDAESEGLSVRGVNAADIQYTGKCYTNGNNCKYDFDGKTHFVKCPSAANTKCEKNGNKCTYDSYNGKVKCDFRH
>Pcon_XP_056577899.1 uncharacterized protein N7517_003919 [Penicillium concentricum]
MQLTTVALFLFAAMGAVASPIESVENGLDARAEAAAQAKYTGTCTRSKNECRYKNDRGKTTFIKCPTKIANKRCTRDGAKCTVDTYNNSVDCD
>Pchr_2NB0_A Chain A, Antifungal protein [Penicillium chrysogenum]
AKYTGKCTKSKNECKYKNSAGKDTFIKCPKFDNKKCTKDNNKCTVDTYNNAVDCD
>Pcin_XP_058307805.1 uncharacterized protein N7498_006552 [Penicillium cinerascens]
MQITKISLFLFAAMGAVASPVDAESGLNARAENAADIKYTGKCYTKDNECKYEADGKTHLVKCPSAANTKCEKNGNKCTYDSYDRKVKCDFRH
>Pcop_XP_056527855.1 uncharacterized protein N7500_009332 [Penicillium coprophilum]
MQITKVALFFFAAMGAVATPIEPVENGLDARAEAGVLVKYTGCPSKIANKRCTKDGAKCTVDTYNNSVDCD
>Pcop_XP_056527837.1 Antifungal protein [Penicillium coprophilum]
MQITRIAFFLFAAMGVVASPIEANSNGLDAQALSKYGGECSKEHNTCTYRKDGKEHKVKCPSADNLKCKTDRHHCEYDDHHKKVDCQTPV
>Pdig_XP_014531969.1 Antifungal protein Afp [Penicillium digitatum]
MQITSIAIILFTAMGAVANPIATASDDLDARDVQLSKYGGQCSLKHNTCTYLKGGRNVIVNCGSAANKRCKSDRHHCEYDEHHRRVDCQTPV
>Pexp_XP_016603682.1 Antifungal protein [Penicillium expansum]

```
MQITKIALFLFAAMGAVASPIEAEAESGINARAENGANVLYTGQCFKKDNICKYKVNGKQNIAKCPSAANKRCEKDKNKCTFDSYDRKVTCDFRK
>Pexp_9FQG_A Chain A, PeAfpB chimeric [Penicillium expansum]
LSKYGGECSKKDNTCTYRKDGKDHIVKCPSADNKKCEKDKNKCEYDDHHKTVDCQTPV
>Pexp_XP_016599536.1 Antifungal protein [Penicillium expansum]
MQITRIAIFLFAAMGAVASPIVAESRDVDAQALSKYGGECSKEHNTCTYRKDGKDHIVKCPSADNKKCKTDRHHCEYDDHHKTVDCQTPV
>Pfim_KAJ5520788.1 hypothetical protein N7463_001241 [Penicillium fimorum]
MQITTVALFLFAAIGVVATPIESVANGLDARAEAGVLAKYTGTCTRSKNECRYKNDRGKTTFIKCPSKIANKRCTKDGAKCTVDTYNNSVDCD
>Pgla_CAI7625829.1 unnamed protein product [Penicillium glandicola]
MQIISIAIALFVAMGAVANPIATESNGLDAREAQLSKYGGECSLKNNTCTYKKDGKDHIVNCPTSTNKKCKTDRHHCEYDDHHKTVDCQTPV
>Pgri_KAJ5189542.1 hypothetical protein N7472_008556 [Penicillium cf. griseofulvum]
MKITSIAIVLFAAMGAVANLIATESDDLVARDVQLDIFGGECSLKHNTCTYKKGGKDQVVKCGSAANTRCKADRNRCQWDDHHKRVECQPPYP
>Pita_KGO72077.1 Antifungal protein [Penicillium italicum]
MQITRIAIFFFAAMGAVANPITNDLNAQALSKYGGECSKEHNTCTYRKDGKDHKVKCPSADNLKCKTDRHHCEYDDHHKKVDCQTPV
>Plon_KAJ5671293.1 hypothetical protein N7507_000420 [Penicillium longicatenatum]
MQITSITIVLFAAIGAVANPIATESDDLNARDLQLSKYGGECSLKHNTCTYRKGGKDHVVNCGSATNRKCKTDRHHCEYDEHHNTVDCQTPV
>Pnuc_XP_056982840.1 uncharacterized protein N7511_007395 [Penicillium nucicola]
MQITNIAIVLFAAMGAVATPIAAESDGLDARDTQLSKYGGECSLQHNTCTYKKGGKDHVVNCPTATNKKCKTDRHHCEYDDHHKTVDCQTPV
>Poxa_EPS29334.1 hypothetical protein PDE_04283 [Penicillium oxalicum 114-2]
MQITKISLFLFAAMGAVASPIDAESDGLNARAVNAANIQYTEKCYTKDNNCKYENDGKTHFVKCPSAANTKCEKDGNRCTHESYNGNVKCDFRH
>Poxa_S8AKE6.1 RecName: Full=Antifungal protein opdH; AltName: Full=Oxopyrrolidines biosynthesis cluster
protein H; Flags: Precursor [Penicillium oxalicum 114-2]
MQFSSLSLVFLAVIGAIANPIAVDSELENRDVQLSKYGGECNLKTNACRYTKGGKSVFVPCGTAANKRCKSDRHHCEYDEHHKRVDCQTPV
>Ppsy_XP_057045999.1 uncharacterized protein N7518_003622 [Penicillium psychrosexuale]
MQIIKVTLFLCAAMGTVATPIDSVSDGLDARAESSALRDYNGVCFRAKNECRYKNDREKTSYVKCSSTIANNRPGNP
>Prub_XP_002557660.1 uncharacterized protein N7525_001733 [Penicillium rubens]
MHITSIAIVFFAAMGAVASPIATESDDLDARDVQLSKFGGECSLKHNTCTYLKGGKNHVVNCGSAANKKCKSDRHHCEYDEHHKRVDCQTPV
>Psal_CAG8925930.1 unnamed protein product [Penicillium salamii]
MQITKVAIFLFAAVGVVANPIATESNDIDARESTKQHGQCDTKNNVCTFDLKGKTNKIKCGSAANKKCRKDRDTCIYDTHHKTVECQI
>Psal_CAG7953647.1 unnamed protein product [Penicillium salamii]
MQITKVAIFLFAAVGVVANPIAIESNDIDARASTKQQGQCDKKNNVCTYVVKGKTNKVKCGSATNKKCRNDRDACTYDTHHKKVDCQL
>Psam_XP_057132942.1 uncharacterized protein N7471_009340 [Penicillium samsonianum]
MQITTVALFLFAAMGAVATPIESVSNGLDARAEAGILAKYTGKCTKSKNECKYKNDAGKDTFIKCPKFDNKKCTKDGNKCTVDTYNNAVDCD
>Psop_XP_057103581.1 antifungal protein precursor [Penicillium soppii]
MQITNLALCFFVAMGAVASPIDTASGGLEARDEAAALRDFPGKCVRSNNTCKYKDDRNKTVIRKCNTKFANQRCTKDGNPCTVDTYGGVVKCS
>Psop_XP_057098855.1 uncharacterized protein N7529_004033 [Penicillium soppii]
MQITKIALFLFTAMRAVASPIDAEPDSLGVRAEDSPSIEYTGKCYKEDNNCKYQADRKTHFVKCPSAANTRCEQDGNKCTYDSYNRKVKCDFRHL
>Psub_XP_057012187.1 uncharacterized protein N7473_002593 [Penicillium subrubescens]
MQITSIAIVLFAAMGAVANPTATESDGLDARDVELSKYGGECSLAHNTCTYLKGGKNQVVACGTAANKRCKTDRHHCEYDEYHKMVDCQTPV
>Pvul_XP_057107628.1 Antifungal protein [Penicillium vulpinum]
MQITRIAIVLFAAMGAVANPVATESNDLDAEAFGSKYGGECSKQHNTCKYRKNGKTHIIKCPSANNLKCKTDRHHCEYDEHHKKVDCQTPV
>EFQ93821.1 hypothetical protein PTT_08687 [Pyrenophora teres f. teres 0-1]
MQFTTATLIFLTALNAVATPIDSASVPTEVRLKFTGTCTKSTDQCSFTRNGKTSISKCSTATAVNYRCTKDKNSCTYDDVDGKTRCT
>XP_001934325.1 antifungal protein [Pyrenophora tritici-repentis]
MQFTTATLLFLTAITVVASPVESVPGDIRIKFDGKCTKSTDQCSFTRNGKTSISKCSTATAVNYRCTKDKNPCTYDDVDGKTRCT
>KAH7355634.1 antifungal protein [Pyrenochaeta sp. MPI-SDFR-AT-0127]
MQITTAALMLFAALGVVATPIDSEPNSIDMRDEVDILIKYSGTCTRDKNECKYKNQENKDTFVKCPTLANKKCTNDGKTCTWDSVSKVVTCD
>KAH7321797.1 antifungal protein [Rhexocercosporidium sp. MPI-PUGE-AT-0058]
MQISKVTLFFIAAMGAVASPIEPSSDGIDARAELITYTGTCTRSDNTCKYKGQSGANTFIKCPTAANKKCTNDGRACSYDSASKVVTCD
>KAL6806198.1 antifungal protein [Trichoderma sp. SZMC 28012]
MKLITLSSIGFALFMAMGAVAVPTNPGSHVVDTHAEGVDAGIHITYYGTCTQKTGQCKYKGQTGMTTICKCPDGCSKDGQSCRFDSVTKLCFCF
>KAI0904534.1 antifungal protein [Ustulina deusta]
MQIATAALFLFAAVGAVATPVESNPNSVDVRDGAGILITYTGTCTKVKNECTYKGENGKDTFVKCPSKKKCTKDGAKCAYDSVTKDVMCE
>KAJ2993564.1 hypothetical protein NUW58_g1805 [Xylaria curta]
MQFATATLFLFAAMGALAIPAESNPNGVDVRDDANILIKYDGTCDKEKNECKYKSQGGGTAFVKCPTFANKRCTKDGNKCTYDSVDKSVTCD
```

```
KAK7983606.1              ---MQFSTVALFLFAVVGAVANPVEGSAD---------GIDAREVQITYDG----------------------T
XP_006667310.1            ---MQIPTFALLLLTAA-AAAMPLSDPLD---------ARDAASAQITYYG----------------------I
PMB64038.1                ---MQIISIALSLLAATGAIAAATPEQFE---------ARDGAGAMIKYHGVSKKPPIHVHYGEKLTIDSTSKI
XP_018702882.1            ---MQITSIALFLLTATGAVAAATPEQFD---------ARDGLSAQIKYHG----------------------I
XP_066727522.1            ---MQFATAALFLFAAMGAIANPVEGNSE------GIDVR---ATETIFHGT----------------------
KAI0904534.1              ---MQIATAALFLFAAVGAVATPVESNPN------SVDVRDGAGILITYTGT----------------------
KAJ2993564.1              ---MQFATATLFLFAAMGALAIPAESNPN------GVDVRDDANILIKYDGT----------------------
Acae_XP_031928216.1       ---MHLTTVVLFLFAAMGAVATPIESEA------FGLDARAEASTLIKYPGQ----------------------
Afla_KOC12588.1           ---MQITTVVLFLFAAMGAVATPIESES------FGLDTRAEASTLIKYPGK----------------------
Psp_KAJ5455444.1          ---MQITTIALFFFAAMGAVANPIASEASIASEANELDARAEAGTLISYSGK----------------------
AWO72254.1                ---MQITVVAVFLLSAMGGVATPINSRI------NPVDARAETGILITYEGT----------------------
KAH7355634.1              ---MQITTAALMLFAALGVVATPIDSEP------NSIDMRDEVDILIKYSGT----------------------
Pcon_XP_056577899.1       ---MQLTTVALFLFAAMGAVASPIESVEN------GLDARAEAAAQAKYTGT----------------------
Pfim_KAJ5520788.1         ---MQITTVALFLFAAIGVVATPIESVAN------GLDARAEAGVLAKYTGT----------------------
Pcop_XP_056527855.1       ---MQITKVALFFFAAMGAVATPIEPVEN------GLDARAEAGVLVKYTG-----------------------
Pchr_2NB0_A               ----------------------------------------------AKYTGK----------------------
Psam_XP_057132942.1       ---MQITTVALFLFAAMGAVATPIESVSN------GLDARAEAGILAKYTGK----------------------
Ppsy_XP_057045999.1       ---MQIIKVTLFLCAAMGTVATPIDSVSD------GLDARAESSALRDYNGV----------------------
Psop_XP_057103581.1       ---MQITNLALCFFVAMGAVASPIDTASG------GLEARDEAAALRDFPGK----------------------
KAH7010366.1              ---MQIATATLFLIAAMGAVASPVEPNAN------GMDARDGIYARITYSGT----------------------
KAH7321797.1              ---MQISKVTLFFIAAMGAVASPIEPSSD------GIDAR---AELITYTGT----------------------
KAH8696995.1              ---MQITTTVLFLSAAIGVIATPIQPERT------SVDVRDNASSRIEYTGK----------------------
KAI0202405.1              ---MQIINAALFLFAAMGAVATPLEAGSD------GLDARGADAVLITYKGK----------------------
USP79915.1                ---MQLASVFLLSFAALGAVANPINSQGD------SIDVRGENNVFITYTGT----------------------
XP_033521083.1            ----------------MSAVATPIDSKLN------DIDTRG---VFITYTGK----------------------
Fnap_KAF5554466.1         ----------------MGVAATPIDSPAM------ALDARGDLEKRLDYKGT----------------------
Fxyl_KAG5755042.1         ---MQFSTITLLLVAAMGVAATPIDSPAM------ALDARGNLEKRLDYKGT----------------------
Faga_KAF4500017.1         ----------------MGVVATPIDSPAM------ALEVRGNLEKRLDYKGT----------------------
Fbul_KAF5965014.1         ---MQFSAITIFLVAAMGVAATPIDSPAI------ALDARGSLEKRLDYKGT----------------------
Fmex_KAF5551368.1         ---MQFSAITLFLVAAMGVAATPIDPPAI------ALDARGSLEKRLDYKGT----------------------
Fpro_XP_031085614.1       ---MQLSAITLFLVAAMGVAATPIDSPVM------ALDARGNLEKRLDYKGT----------------------
Fver_XP_018757790.1       ---MQFSTITLLLLASTGVAATPIYSPAM------PLDARRNLEKRLEYKGT----------------------
Fave_KAH6970200.1         ---MQFSTITLFLAATMGVAASPVDTPAQ------ELDARGNLFPRLDYHGT----------------------
Ftri_KAH7263759.1         ---MQFSTITLFLAATMGVAATPVDSPAQ------ELDARGNLFPRLDYHGT----------------------
Fgre_CAF3538390.1         ---MQFSTIVPLFVAAMGVVATPVNSPAQ------ELDARGNLFPRLEYWGK----------------------
Fpse_XP_009263607.1       ---MQFSTILPLFVAAMGIVATPVNSPAQ------ELDARGNLLPRLEYWGK----------------------
Fsam_KAL6912397.1         ---MQFSTILPLFIAAMGVVATPIDAPAQ------ELDARGNLFPRLEYWGS----------------------
Fven_XP_025591605.1       ---MQFSTIFSLFIAAMGIVATPIDSPTQ------ELDARGNLFPRLEYWGS----------------------
Flan_GKU13684.1           ---MQISTILPLFVAAMGVVATPIDSPAQ------ELDARGNLFPRLEYWGS----------------------
Fgra_KAF4995278.1         ---MQFSTIALVFVAAMGAVATPVDPPAQ------DLDARGDLYPRLDYWGT----------------------
Faus_KAF5228166.1         ---MKFSSVTLLLFVATGAVATPVDSLPN------QLDARDGLFPRRTFPGK----------------------
Fchl_KAL4723034.1         ---MQFSSITLLLAAAIGAVATPVDSPPS------QLDTRSRLFPRRPYEGT----------------------
Flon_RGP60307.1           ---MHLSTITLFFVAAMSAVATPVDSSAQ------NLDARANLGRRRDFPGT----------------------
Abra_GKZ20515.1           ---MHITSIAIVLFAAMGAIANPIAAEAD--------DLLAREAELSKYGGE----------------------
Abra_XP_067479519.1       ----------------MGAIANPIAAEAD--------DLLAREAQLSKYGGE----------------------
Aeuc_XP_025382252.1       ---MKLTSIAIIILFAAMGAIANPIAAESD--------DLLARDVQLSKYGGE----------------------
Apho_RDK38155.1           ---MQLTSIAIIILFAAMGAIANPITAEAN--------NLVAREEELSKYGGE----------------------
Aser_KAE8333019.1         ---MQITSVAIVLFAAMGAVANPIATQSD--------DLSARDIQLSKFGGE----------------------
Plon_KAJ5671293.1         ---MQITSITIVLFAAIGAVANPIATESD--------DLNARDLQLSKYGGE----------------------
Avir_XP_043122868.1       ----MHTSIAIVLIAAMGAIANPIVTESD--------DLDARDVQLSKFGGE----------------------
Psp_KAJ6113509.1          ---MQITSIAIVLFAAMGAIANPIGTESD--------DLDARDVQLSKYGGE----------------------
Psub_XP_057012187.1       ---MQITSIAIVLFAAMGAVANPTATESD--------GLDARDVELSKYGGE----------------------
Psp_KAJ5413027.1          ---MQITSIAIVLFAAMGAVANPIPTESD--------DLVARDVQLSKFGGE----------------------
Prub_XP_002557660.1       ---MHITSIAIVFFAAMGAVASPIATESD--------DLDARDVQLSKFGGE----------------------
Pdig_XP_014531969.1       ---MQITSIAIILFTAMGAVANPIATASD--------DLDARDVQLSKYGGQ----------------------
Atam_KAE8159784.1         ---MQLTSIAIFLFAAMGAVANPIATESD--------DLDTRDIELSKFGGE----------------------
AHA86567.1                ---MQFTKIAIFLFAAMGAVANPIAAESG--------DLDVRDVQLSKYGGE----------------------
Pgla_CAI7625829.1         ---MQIISIAIALFVAMGAVANPIATESN--------GLDAREAQLSKYGGE----------------------
Pnuc_XP_056982840.1       ---MQITNIAIVLFAAMGAVATPIAAESD--------GLDARDTQLSKYGGE----------------------
Psp_KAJ5466821.1          ---MQITRIAIVLFAAMGAVANPIAAESN--------SLDTQAFGS-KYGGE----------------------
Pvul_XP_057107628.1       ---MQITRIAIVLFAAMGAVANPVATESN--------DLDAEAFGS-KYGGE----------------------
Pcop_XP_056527837.1       ---MQITRIAFFLFAAMGVVASPIEANSN--------GLDAQALS--KYGGE----------------------
Pita_KGO72077.1           ---MQITRIAIFFFAAMGAVANPITNDLN-----------AQALS--KYGGE----------------------
Pexp_XP_016599536.1       ---MQITRIAIFLFAAMGAVASPIVAESR--------DVDAQALS--KYGGE----------------------
Pexp_9FQG_A               -------------------------------------------LS--KYGGE----------------------
Pgri_KAJ5189542.1         ---MKITSIAIVLFAAMGAVANLIATESD--------DLVARDVQLDIFGGE----------------------
Poxa_S8AKE6.1             ---MQFSSLSLVFLAVIGAIANPIAVDSE--------LE-NRDVQLSKYGGE----------------------
Psal_CAG8925930.1         ---MQITKVAIFLFAAVGVVANPIATESN--------DIDAREST--KQHGQ----------------------
Psal_CAG7953647.1         ---MQITKVAIFLFAAVGVVANPIAIESN--------DIDARAST--KQQGQ----------------------
Pbra_CEJ62478.1           ---MQVAKISLFLFAAMGTVASPIDAESE------GLSVRGVNAADIQYTGK----------------------
Poxa_EPS29334.1           ---MQITKISLFLFAAMGAVASPIDAESD------GLNARAVNAANIQYTEK----------------------
Pcin_XP_058307805.1       ---MQITKISLFLFAAMGAVASPVDAES-------GLNARAENAADIKYTGK----------------------
Patr_XP_056739537.1       ---MQITKVSLFLFAAMGAIASPIDAESD------GLNARVENAANIEYTGK----------------------
Aspa_SMQ11440.1           ---MQITKISLFLFVGVGVVASPIHAESD------GLNARAVNAADLEYKGE----------------------
Psop_XP_057098855.1       ---MQITKIALFLFTAMRAVASPIDAEPD------SLGVRAEDSPSIEYTGK----------------------
Pexp_XP_016603682.1       ---MQITKIALFLFAAMGAVASPIEAEAES-----GINARAENGANVLYTGQ----------------------
KAJ9304236.1              ---MQITKISLFLFAAIAAVANPIDAESD------GVVERDVDAADITYTGE----------------------
EFQ93821.1                ---MQFTTATLIFLTALNAVATPIDSAS---------VPTEVR---LKFTGT----------------------
XP_001934325.1            ---MQFTTATLLFLTAITVVASPVES-----------VPGDIR---IKFDGK----------------------
TEA21567.1                ---MQFANNFVFLLAAMGAIASPVQPNSN------DVGPADLEGDGITYTGK----------------------
XP_066727524.1            ---MQFSTVTLFLVAAMGAIATPVNSADG-------VEARAEQGTLLEYTGT----------------------
KAK7994358.1              ---MQFTTAALSLLAAMGAIAIPLNTTMT-------LETQGEMGMFITYPGK----------------------
```

```
KAJ0271918.1            ---MQITKVALFLFAAIGVAANPVNVDGS--GIDVGVSGGEDMNTFITYTGKCSR------------------G
KAF4922418.1            ---MQIVKVALFLFAAIGVAANPVDVDGS--GMDVGVSGGEDMNTFITYTGKCTR------------------G
KAF4920109.1            ---MQIVKVALFLFAAIGVAANPVDVDGS--GMDVGVSGGEDMNTFITYTGKCTR------------------G
KAF0329504.1            ---MQIAKIALFLFAAIGVAANPVDIDDAS-GIDAGVSGGEDMNTFLTYTGKCTR------------------G
XP_053031537.1          ---MQIAKVAFFLFAAIGVAANPVDVDGS--GIDAGVSGGEDMNTLITYTGKCTR------------------G
KAF4823470.1            ---MQIAKVAFFLFAAIGVAANPVDVDGS--GIDAGVSGG---------------------------------G
Amel_KAK1138769.1       MKLTSLASLGIVLFTAMGTLGSPVDSGSLASND---LDARDEAGIMIRYDGKCSK-------------------
Aste_XP_024699894.1     MKFLSIASLSLILFTAMGVLGSPIESEALASND---LDARDEAGILIKYPGTCSK-------------------
Aall_KAE8393168.1       MKASSFVSLIFIFATALGVAASPTNTNSVPSND---LAMEDEAEIKIKYYGKCFID------------------
Acam_XP_024690694.1     MQLISLASMGLVLFAAVGAVASPVDNNALDVNDN--LEVHDEAATLITYNGSCSK-------------------
Atai_PLN82996.1         MQLISLASMGLVLFAAVGAVASPVDNNALDIDNN--LEVRDEAASLIKYHGVCSK-------------------
Ahan_KAF7592939.1       -MKFTSLSLGLVLFSALGAVASPVDAASANRVEV--REEAGDAGILIKYDGTCSK-------------------
Ajap_XP_025526306.1     -MKISPVSIGFILLAAMGVAATPLNHAES-------VGVRSENNVQVKYDGQCRK-------------------
PFH58491.1              --MVSLYSIAFALLAVSGAVASPAGDSPS-------LEQR---DAAITYNGKCSA-------------------
Agig_CAA37523.1         ---------------VATPVEADSLTAGG-------LDARDESAVLATYNGKCYK-------------------
KAL6806198.1            MKLITLSSIGFALFMAMGAVAVPTNPGSHVVDT---HAEGVDAGIHITYYGTCTQ-------------------
Awen_KAI9925351.1       MKIISLASFGIALFTAL-AAATPVEPNTV--------EARAEQGVLIKYDGKCSK-------------------


KAK7983606.1            CSRSKNECKYKGQNGRPTIVKCPS---FANKKVFIQTP--------
XP_006667310.1          CTKANNECKYKNQNGKDTFVKCPR---FANKKRQQSRAVPLDSVNVAEPFEHFPRLFVPIDI------------
PMB64038.1              CTKAKNECKFKGQNGRDTFVKCPS---FANKRCTKDYNECSYDSVSRAVVCH----------------------
XP_018702882.1          CTKAKNECKFKGQNGRDTFVKCPN---FANKRCTKDHNECSYDSVSRAVVCH----------------------
XP_066727522.1          CSKVKDECNYKGEHGKLHHVKCP----PK-DKCTKKGAKCTFNSKDKKVICH----------------------
KAI0904534.1            CTKVKNECTYKGENGKDTFVKCP----SK-KKCTKDGAKCAYDSVTKDVMCE----------------------
KAJ2993564.1            CDKEKNECKYKSQGGGTAFVKCPT--FAN-KRCTKDGNKCTYDSVDKSVTCD----------------------
Acae_XP_031928216.1     CTKAKNECKYKSQNKKNTFVKCPS--FAN-KRCTKDGNWCQFDSYSRNVECK----------------------
Afla_KOC12588.1         CSKAKNECKFKGQTKKDTFVKCPS--FAN-KRCTKDGNPCHFDSYSRTVDCK----------------------
Psp_KAJ5455444.1        CYKSKNECKFKGQNGKTTFVKCPK--FAN-KKCTKDGASCKYDSYDGKVTCN----------------------
AWO72254.1              CSRAKNECKYKNQNNKDTFVKCPS--FAN-KKCTKDNAKCSFDSYSRAVTCH----------------------
KAH7355634.1            CTRDKNECKYKNQENKDTFVKCPT--LAN-KKCTNDGKTCTWDSVSKVVTCD----------------------
Pcon_XP_056577899.1     CTRSKNECRYKNDRGKTTFIKCPT--KIANKRCTRDGAKCTVDTYNNSVDCD----------------------
Pfim_KAJ5520788.1       CTRSKNECRYKNDRGKTTFIKCPS--KIANKRCTKDGAKCTVDTYNNSVDCD----------------------
Pcop_XP_056527855.1     ---------------------CPS--KIANKRCTKDGAKCTVDTYNNSVDCD----------------------
Pchr_2NB0_A             CTKSKNECKYKNSAGKDTFIKCPK--FDN-KKCTKDNNKCTVDTYNNAVDCD----------------------
Psam_XP_057132942.1     CTKSKNECKYKNDAGKDTFIKCPK--FDN-KKCTKDGNKCTVDTYNNAVDCD----------------------
Ppsy_XP_057045999.1     CFRAKNECRYKNDREKTSYVKCSS--TIANN---RPGNP-----------------------------------
Psop_XP_057103581.1     CVRSNNTCKYKDDRNKTVIRKCNT--KFANQRCTKDGNPCTVDTYGGVVKCS----------------------
KAH7010366.1            CTRSDNTCKYKNQNNGNTFIKCP---TAANKKCTNDGKACSYDSVSKAVTC-----------------------
KAH7321797.1            CTRSDNTCKYKGQSGANTFIKCP---TAANKKCTNDGRACSYDSASKVVTCD----------------------
KAH8696995.1            CTRSTNICRYEGQNNKIILISCP---SAANLRCTNDGRACSYESSTKKVTCG----------------------
KAI0202405.1            CTKSSNTCKYTGQNGKTTIVSCP---TARNLVCTNDNKECTFDSVDKKVTCS----------------------
USP79915.1              CTKANNQCRYKAQNGKTAFAKCP---KFANKKVHLKHTYVTKLD------------------------------
XP_033521083.1          CDASTQQCRYNGQNGVVTIAKCG---IAANKKCTTLPTTGGTCEYDSASKVLTCH-------------------
Fnap_KAF5554466.1       CTKSSNTCRYKGPNGRTTFKKCG---TFANQKVQSRGFSAMLN-------------------------------
Fxyl_KAG5755042.1       CTKSSNTCRYKGPNGRTAFKKCG---TFANQKCTKDGASGFENLAGFYMSSNMTVTGICEIVIVGTGPGTNIGI
Faga_KAF4500017.1       CTRSSNTCRYKGPNGRIAFKKCG---TFANQKCTKDGAPCVWQSDKGAGDNADGICLDIGLS------------
Fbul_KAF5965014.1       CTRSSNTCRYKGPNGRTAFKKCG---TFANQKCTKDGAPCVWQSDKGVGGKITCK-------------------
Fmex_KAF5551368.1       CTRSSNTCRYKGPNGRTAFKKCG---TFANQKCTKDGAPCVWQSDKGVGGKITCK-------------------
Fpro_XP_031085614.1     CTKSSNTCRYKGPNGRTAFKKCG---TFANQ---KDGAPCVWQSDKGVGGKIICK-------------------
Fver_XP_018757790.1     CTKSSNTCRYKGPNGRTTFKKCG---TFANQKCTKDGAPCVWESEKGVGGKVTCK-------------------
Fave_KAH6970200.1       CTKSTNRCRYINDKKRTVIISCP---KFANKKCTKDGNKCTYDSAARSVICR----------------------
Ftri_KAH7263759.1       CTKSTNRCRYINDKKRTVIISCP---KFANKKVQSSKMLTDNFY------------------------------
Fgre_CAF3538390.1       CTKAENRCKYKNDKGKDVLQNCP---KFDNKKCTKDGNSCKWDSASKALTCY----------------------
Fpse_XP_009263607.1     CTKAENRCKYKNDKGRDVLQNCP---KFDNKKCTKDGNSCKWDSASKALTCY----------------------
Fsam_KAL6912397.1       CTKSDNRCKYKNDKGNSVLQNCP---SFDNKKCTKDGNSCKWDSAKKELTCY----------------------
Fven_XP_025591605.1     CTKSNNRCKYKNDDDNSVLQNCL---SFNNKKCTKDGNNCKWDSAKKELNYY----------------------
Flan_GKU13684.1         CTKAENRCKYKNDKGKEVLQNCP---KFDNKKVQSHKVPNSFHTN-----------------------------
Fgra_KAF4995278.1       CTKANNRCRYKNDKGRTVLQNCP---KFTNKKCTKDGNRCKWDSAAKDLICY----------------------
Faus_KAF5228166.1       CTRSDNRCKYENSNGKTVTISCG---TAANKKCTKDNADCVYDDANRSVKCD----------------------
Fchl_KAL4723034.1       CTRADNRCKYDNSNGKTVTISCG---TAANRKCTKDGAKCVYDDADRSVKCD----------------------
Flon_RGP60307.1         CTKSDNRCKYKNSNDKYVTIACP---KFDNKKCTKDGNSCTYDDADRSVKCD----------------------
Abra_GKZ20515.1         CSLEHNTCTYR-KDGKNHVVACPT---AANLRCKTDRHHCEYDDHHKTVDCQTPV-------------------
Abra_XP_067479519.1     CSLEHNTCTYR-KDGKNHVVACPT---AANLRCKTDRHHCEYDDHHKTVDCQTPV-------------------
Aeuc_XP_025382252.1     CSLEHNTCTYR-KDGKNHVVSCPS---ATNLRCKTDRHHCEYDDHHKTVDCQTPV-------------------
Apho_RDK38155.1         CSVEHNTCTYL-KGGKDHIVSCPS---AANLRCKTERHHCEYDEHHKTVDCQTPV-------------------
Aser_KAE8333019.1       CSLKHNTCTYL-KGGKNHVVNCGS---ADNQKCKEERHHCEYDDHHKTVNCQTPV-------------------
Plon_KAJ5671293.1       CSLKHNTCTYR-KGGKDHVVNCGS---ATNRKCKTDRHHCEYDEHHNTVDCQTPV-------------------
Avir_XP_043122868.1     CSLKYNTCTYL-KGGKNQVVNCGS---AANKRCKTDRRHCEYDEYHKTVDFQTPV-------------------
Psp_KAJ6113509.1        CSLTHNTCTYL-KGGKHIAVNCGS---AANKRCKTDRHHCEYDEHHKTVDCQTPV-------------------
Psub_XP_057012187.1     CSLAHNTCTYL-KGGKNQVVACGT---AANKRCKTDRHHCEYDEYHKMVDCQTPV-------------------
Psp_KAJ5413027.1        CSLKHNTCSYR-KGGKTRIVNCGS---AANKKCKSDRHHCEYDEHHKRVDCQTPV-------------------
Prub_XP_002557660.1     CSLKHNTCTYL-KGGKNHVVNCGS---AANKKCKSDRHHCEYDEHHKRVDCQTPV-------------------
Pdig_XP_014531969.1     CSLKHNTCTYL-KGGRNVIVNCGS---AANKRCKSDRHHCEYDEHHRRVDCQTPV-------------------
Atam_KAE8159784.1       CSVEHNTCTYL-KGGKNHVVKCGS---AANKKCKSDRHHCEYDEHHKTVDCQTPV-------------------
AHA86567.1              CSLQHNTCTYL-KGGKNQVVHCGS---AANQKCKSDRHHCEYDEHHKTVNCQTPV-------------------
Pgla_CAI7625829.1       CSLKNNTCTYK-KDGKDHIVNCPT---STNKKCKTDRHHCEYDDHHKTVDCQTPV-------------------
Pnuc_XP_056982840.1     CSLQHNTCTYK-KGGKDHVVNCPT---ATNKKCKTDRHHCEYDDHHKTVDCQTPV-------------------
Psp_KAJ5466821.1        CSKEHNTCTYR-KDGKDHKVKCPS---ADNLKCKTDRHHCEYDEHHKKVDCQTPV-------------------
Pvul_XP_057107628.1     CSKQHNTCKYR-KNGKTHIIKCPS---ANNLKCKTDRHHCEYDEHHKKVDCQTPV-------------------
Pcop_XP_056527837.1     CSKEHNTCTYR-KDGKEHKVKCPS---ADNLKCKTDRHHCEYDDHHKKVDCQTPV-------------------
```

```
Pita_KGO72077.1          CSKEHNTCTYR-KDGKDHKVKCPS---ADNLKCKTDRHHCEYDDHHKKVDCQTPV-------------------
Pexp_XP_016599536.1      CSKEHNTCTYR-KDGKDHIVKCPS---ADNKKCKTDRHHCEYDDHHKTVDCQTPV-------------------
Pexp_9FQG_A              CSKKDNTCTYR-KDGKDHIVKCPS---ADNKKCEKDKNKCEYDDHHKTVDCQTPV-------------------
Pgri_KAJ5189542.1        CSLKHNTCTYK-KGGKDQVVKCGS---AANTRCKADRNRCQWDDHHKRVECQPPYP------------------
Poxa_S8AKE6.1            CNLKTNACRYT-KGGKSVFVPCGT---AANKRCKSDRHHCEYDEHHKRVDCQTPV-------------------
Psal_CAG8925930.1        CDTKNNVCTFD-LKGKTNKIKCGS---AANKKCRKDRDTCIYDTHHKTVECQI---------------------
Psal_CAG7953647.1        CDKKNNVCTYV-VKGKTNKVKCGS---ATNKKCRNDRDACTYDTHHKKVDCQL---------------------
Pbra_CEJ62478.1          CYTNGNNCKYD-FDGKTHFVKCPS---AANTKCEKNGNKCTYDSYNGKVKCDFRH-------------------
Poxa_EPS29334.1          CYTKDNNCKYE-NDGKTHFVKCPS---AANTKCEKDGNRCTHESYNGNVKCDFRH-------------------
Pcin_XP_058307805.1      CYTKDNECKYE-ADGKTHLVKCPS---AANTKCEKNGNKCTYDSYDRKVKCDFRH-------------------
Patr_XP_056739537.1      CVAKDNNCRYG-IGGKTHLVKCPS---AANTKCEKDGNKCTYDSYNGKVKCDFRH-------------------
Aspa_SMQ11440.1          CFTKDNTCKYK-IDGKTYLAKCPS---AANTKCEKDGNKCTYDSYNRKVKCDFRH-------------------
Psop_XP_057098855.1      CYKEDNNCKYQ-ADRKTHFVKCPS---AANTRCEQDGNKCTYDSYNRKVKCDFRHL------------------
Pexp_XP_016603682.1      CFKKDNICKYK-VNGKQNIAKCPS---AANKRCEKDKNKCTFDSYDRKVTCDFRK-------------------
KAJ9304236.1             CFRKNNECRYV-ANGKTHYVKCPS--KFANKRCQMDKHKCTFDSYSRVVNCNA---------------------
EFQ93821.1               CTKSTDQCSFT-RNGKTSISKCSTAT-AVNYRCTKDKNSCTYDDVDGKTRCT----------------------
XP_001934325.1           CTKSTDQCSFT-RNGKTSISKCSTAT-AVNYRCTKDKNPCTYDDVDGKTRCT----------------------
TEA21567.1               CTRKTNECNFT-ASRKKKSVKCP--S-LPNKTCTKDGAKCTYDILSQEIVCY----------------------
XP_066727524.1           CTKAKNECKFKGQTGATTFVKCPT--KPNNHRCFRDGNKCTFDSYSRKVVCT----------------------
KAK7994358.1             CTLKTQTCRYKGQNGQTTLAKCPK--SPANKRCFGDGHSCSFDSVTRKVSCS----------------------
KAJ0271918.1             RNYKEDTCKFKGQKGKTTIVRCPR---FANQR------------------------------------------
KAF4922418.1             RNYKEDTCKFKGQKGKTTIVRCPR---FANQRVSFDMLSSIQIHKLNRSDIMVFSSSALGMAVSAHGTVTSGLP
KAF4920109.1             RNYKEDTCKFKGQKGKTTIVRCPR---FANQRCPRNGSKCTWDSYKRTTKCNYK--------------------
KAF0329504.1             RNYKEDTCKFKGQKGKTTIVRCPR---FANQRVSFDSLSSTQCPRNGSKCTWDSYKRTTKCNYK----------
XP_053031537.1           RNYKEDTCKFKGQKGKTTIVRCPR---FANQRVSFDTQYTGAINGPKKNIGQSKNFNATTA-------------
KAF4823470.1             RNYKEDTCKFKGQKGKTTIVRCPR---FANQRLQADYQVQL---------------------------------
Amel_KAK1138769.1        ---KDNSCRYKSQNGRTAFCKCQFKR------CAKDGNKCHYESYNGNCQCI----------------------
Aste_XP_024699894.1      ---KNNNCRYKSQNGRTAFCKCKFKK------CAKDGNKCHFDSYNQDCQCI----------------------
Aall_KAE8393168.1        EKMKMTKCKYDGPNGRTSFCRCHFKR------CTKNGAKCHFDSVNKDCYCS----------------------
Acam_XP_024690694.1      ---KNNSCKYKGQKGKTSFCHCKFKK------CGKDGNKCHFDSYSRDCKCI----------------------
Atai_PLN82996.1          ---KNNSCKFKGQNGKTSFCHCKFKK------CGKENNKCHFDSYNRDCKCI----------------------
Ahan_KAF7592939.1        ---KNNECKYKAQNGKTAFCKCQVKK------CGSDGGKCFYDSANRQCTCY----------------------
Ajap_XP_025526306.1      ---SENQCRYTAQSGRTAICKCQFRK------CSKDGAKCNFDSYNRDCNCY----------------------
PFH58491.1               ---KANTCRYVGQSGRPSICKCYVKK------CSGDGKACHYDSYKNQCLCV----------------------
Agig_CAA37523.1          ---KDNICKYKAQSGKTAICKCYVKK------CPRDGAKCEFDSYKGKCYC-----------------------
KAL6806198.1             ---KTGQCKYKGQTGMTTICKCPDG-------CSKDGQSCRFDSVTKLCFCF----------------------
Awen_KAI9925351.1        ---KDNSCKFKGQGGKTTFCHCKFKK------VFLPPS----DIVDL---------------------------
                                                            *
```

**Supplementary File S8.**
**Sequences of *A. pisum* HLPs and their most similar orthologues in Fungi, used for the alignment of Figure 5.**

```
>Apis_XP_003248175.1 uncharacterized protein LOC100573341 [Acyrthosiphon pisum]
MVAQKILSLMLVGLLIASSANAGPIAAGICYAGCAGVTVACFTAAGFTFGTVPGAVIAATPALAACNAAFGICEASCVAALLLPTP
>Apis_XP_029343601.1 uncharacterized protein LOC100163777 [Acyrthosiphon pisum]
MVGQKMWSLMLVGLLIASSANAGPIAAGICYAGCAGVTVACFSAAGFTFGTVPGALIAATPALAACNAAFGVCEASCMAALFVPVP
>Apis_XP_016657431.1 uncharacterized protein LOC100164849 [Acyrthosiphon pisum]
MVAQKFLSLMLAGLLIASSANAGPIAAGICYAGCAGVTVACFAAAGFTFGTVPGAVIAATPALAACNAAFGICEASCVAALVVPVP
>Apis_XP_016657429.1 uncharacterized protein LOC107882876 [Acyrthosiphon pisum]
MVAQKIWSLMLVGLLIASSANAGPIAAGICYAGCAGVTVACFAAAGFTFGTVPGAVIAATPALAACNAAFGICEASCIAALVVPVP
>Apis_XP_003247138.1 uncharacterized protein LOC100165615 [Acyrthosiphon pisum]
MIAQKLWSLIFVGLLISSSANAGPIAAGICYAGCAAVTVACFSAAGFTFGTVPGAVIAATPMLAACNAAFGICEASCVAALIVPVP

>Pfij_XP_007922500.1 uncharacterized protein MYCFIDRAFT_43367 [Pseudocercospora fijiensis
CIRAD86]
MQFQKIFTLSMLATYVTAGPAAYGICQAGCAGVTVACYSAAGFVFGVALPAAPPAILACNAAFGSCQAACWAALIAPTP
>Pcit_KAK7545839.1 hypothetical protein IWX46DRAFT_92887 [Phyllosticta citricarpa]
MRLTNLMTSLAVVTSATAGPLGYGICQAGCSGVVVACYAAAGFTFGTVLAVAAPPAILVCNSAYGTCQAACAAVLLAPTL
>Apru_XP_033393871.1 uncharacterized protein K452DRAFT_235045 [Aplosporella prunicola CBS
121167]
MLRTTILTACVLTLAGTASAGPVGYGICQAGCAGVVMACYTAAGFTWGATLGASAPPTIIACNTAFGSCQAACAAILLAPTP
>Peum_KXT04810.1 hypothetical protein AC578_9754 [Pseudocercospora eumusae]
MKFTKIALPILASFVPLVKAGPAAYGVCQAGCAGLAVACYAAAGFTFGVALPAAPPAILACNATFGSCQAACWAALFTPTP
>Cely_KAF1937296.1 hypothetical protein EJ02DRAFT_458854 [Clathrospora elynae]
MKFLINIITTALAILIFATTASAGPIGYAICQGGCAGVVMACYSAAGFTWGATLGATAPATVLACNAAYGTCQAACAAVLLVPLP
>Gtri_XP_009229506.1 hypothetical protein GGTG_13336 [Gaeumannomyces tritici R3-111a-1]
MRRFTSASIMLVTMAFTSPAFAGPAAYGVCQAGCAAVVMACYSAAGFTWGATLGVSAPPTIIACNTSFGTCQAACAAVLLSPTP
>Anig_XP_025460783.1 uncharacterized protein BO96DRAFT_407345 [Aspergillus niger CBS
101883]
MKNIYLAVFTILIFVSYASAGPAAYGICQAGCAAVVMACYSAAGYTWGATLGATAPPTIVACNSAFGVCYSSCAATLLAPTL
>Pver_XP_057021283.1 uncharacterized protein N7466_006287 [Penicillium verhagenii]
MRSHWAQFWLPLLLATNVSAGPAAYGVCQAGCAALVMACYSAAGFTWGVAMGATIPASIVTCNSAFGTCQAACASVLLAPTL
```

#### **Supplementary File S9.**
#### **Hexapoda HLP sequences found in NCBI protein database**

>Apis_XP_003248175.1 uncharacterized protein LOC100573341 [Acyrthosiphon pisum] Hemiptera
M*VAQKILSLMLVGLLIASSANA*GPIAAGICYAGCAGVTVACFTAAGFTFGTVPGAVIAATPALAACNAAFGICEASCVAALLLPTP
>Apis_XP_029343601.1 uncharacterized protein LOC100163777 [Acyrthosiphon pisum] Hemiptera
*MVGQKMWSLMLVGLLIASSANA*GPIAAGICYAGCAGVTVACFSAAGFTFGTVPGALIAATPALAACNAAFGVCEASCMAALFVPVP
>Apis_XP_016657431.1 uncharacterized protein LOC100164849 [Acyrthosiphon pisum] Hemiptera
*MVAQKFLSLMLAGLLIASSANA*GPIAAGICYAGCAGVTVACFAAAGFTFGTVPGAVIAATPALAACNAAFGICEASCVAALVVPVP
>Apis_XP_003247138.1 uncharacterized protein LOC100165615 [Acyrthosiphon pisum] Hemiptera
*MIAQKLWSLIFVGLLISSSANA*GPIAAGICYAGCAAVTVACFSAAGFTFGTVPGAVIAATPMLAACNAAFGICEASCVAALIVPVP
>Apis_XP_016657429.1 uncharacterized protein LOC107882876 [Acyrthosiphon pisum] Hemiptera
*MVAQKIWSLMLVGLLIASSANA*GPIAAGICYAGCAGVTVACFAAAGFTFGTVPGAVIAATPALAACNAAFGICEASCIAALVVPVP
>Agif_XP_044016122.1 uncharacterized protein LOC122857800 [Aphidius gifuensis]
MVSSKASFCMVVMMVILLSTHTSDAGPIGAGICYAGCAALVGACFAAAGFTFGTVPGAIIAATPALAGCNAAFAACEAACVAALIAPTP
>Acra_KAF0749826.1 Uncharacterized protein FWK35_00035468 [Aphis craccivora]
MTAQKMWTFVLVGLLMMTCVTEASRLNRFMSNVCYFGCLAERVACFSSTGAIFGTVPYGIIAVTPTLESCTVVFKICKASCIAILISSKI
>Acra_KAF0768549.1 Uncharacterized protein FWK35_00012841 [Aphis craccivora]
MVTHKMLSLVLVGLLLSSSAHAGPLAAGVCYAGCAAVTVACFSAAGFTFGTVPGAIIAATPALAACNAAFGVCEASCIAALVVPVP
>Agos_XP_027838196.1 uncharacterized protein LOC114120478 [Aphis gossypii]
MVAHKMLSLVLVGLLLSSTAHAGPLAAGVCYAGCAAVTVACFSAAGFTFGTVPGAIIAATPALAACNAAFGICEASCIAALVVPVP
>Aluc_KAF6202012.1 hypothetical protein GE061_004408 [Apolygus lucorum]
MAASTTTTVILAFLVASSLFVGSDGGPVAAGICYAGCASMVVACFAAAGFTFGTVPGAQIAAVPALAGCNTAFGVCEAACVAALVTPTP
>Aluc_KAF6202579.1 hypothetical protein GE061_002977 [Apolygus lucorum]
MAASTTTTIVLAFLIASSLFVGSDGGPIAAGICYAGCASIVVACFAAAGFTFGTVPGAQIAAVPALAGCNTAFGVCEAACVAALVTPVP
>Bkin_XP_033220284.1 uncharacterized protein LOC117174934 [Belonocnema kinseyi]
MDYKLYALFVLVFLANFGSAGPIAAGIGYAGCAALACACFAVAGFTFGTVTIATILASPALTACNVAFAKCEAAVMAMLVAPTP
>Btab_CAH0392446.1 unnamed protein product [Bemisia tabaci]
MKHLLSLTFAAMLLSTATAGPIGAGICYAGCAGVVVACFSAAGFTFGTVPGSQIAAVPALAACNSAFGSCMAACSAALALPIP
>Bger_PSN34680.1 hypothetical protein C0J52_22484 [Blattella germanica]
MLLLVNQVYSGPIAAGICYAGCAGVVVACFAAAGATFGTVPGSVIAATPALATCNTAFGTCEAACVAALLAFTP
>Bger_PSN33737.1 hypothetical protein C0J52_23715 [Blattella germanica]
MLLICQVCQVGSGRVAAGICYAGCAAVVVACFAAAGFTFGTVPGALIAATPALAACNGAFAVCERACIAALADPVP
>Cass_CAH1129794.1 unnamed protein product [Ceutorhynchus assimilis]
MMATTMVIKTALFTILLFYMCSTSAEAGPAAAGVCYAGCAAVTVACFAAAGFTFGTVPGAVIAATPALAACNAAFGVCEAACMAALFMPTP
>Cass_CAG9767903.1 unnamed protein product [Ceutorhynchus assimilis]
MATTMVTKAALFTILLFYMCSTSADAGPAAAGVCYAGCAAVTVACFAAAGFTFGTVPGALIAATPALAACNAAFGVCEAGCMAALFMPTP
>Cass_CAG9767865.1 unnamed protein product [Ceutorhynchus assimilis]
MVTKTALFAILLFYMCSISAEAGPAAAGVCYAGCAAVAVACFAAAGFTFGTVPGAVIVATPALAACNSAFGVCEAACMAALFMPTP
>Crip_CAH1729104.1 unnamed protein product [Chironomus riparius]
MMSYKVNFCAVLLLLLTVTNNVDCGVIAFGICEATCNNLATACYTATGFTIAPLTLGLGTPVAVLACNAAFAECMSSCVLAGLSPLL
>Cced_VVC36620.1 Hypothetical protein CINCED_3A000125 [Cinara cedri]
MFARKTIVLLTVLMMFGIAQAGPLAAGICYAGCASVTVACFAAAGFTFGTVPGAVIAATPALAACNTAFGICEAACVAALVAPTV
>Cmar_CRK86781.1 CLUMA_CG000612, isoform A [Clunio marinus]
MRNVIFICIVLLLSASSIKAGPLAYGICQTGCNAMVVACYAAAGFTFGTVTAGAGVPAAIIACNVALGTCMAACVAAGCAPTP
>Csep_XP_044751818.1 uncharacterized protein LOC123311792 [Coccinella septempunctata]
MLQVKMVLTYGICQAACAGLVVACFSAAGFTFGTVPGAIIAATPALAACNTAFASCSAACAPLLLSPI
>Cmon_KAL3282720.1 hypothetical protein HHI36_005892 [Cryptolaemus montrouzieri]
MPLLTYGLCQAACAGIVVACFSAAGFTFGTVPGAIIGATPALAACNSAFAACSSQCSWLLLSPV
>Cson_AAU06531.1 unknown salivary protein [Culicoides sonorensis]
MKNILIYMSILCLLSYPVVAGPAASSICYAGCAAVVVACFAAAGFTFGTVPGAQIAAVPALASCNAAFATCEAACMAAFFLPTP
>Dvit_XP_050533516.1 uncharacterized protein LOC126901216 [Daktulosphaira vitifoliae]
MATHKIPFILSILILTSSVVTAGPLGVGICYAGCAGVTVACFAAAGATFGTVPAAVIAASPALAACNSAFAGCYSACALALIAPTP
>Dcit_KAI5706695.1 hypothetical protein M8J75_010515 [Diaphorina citri]
MNTKVILLCFLLICSQYTEAGPIAMATAQAGCAAVVMACYAAAGATWGATLGATAPASVIACNSAFGVCMTAASSFLLAPIP
>Dsim_XP_046747924.1 uncharacterized protein LOC124412242 [Diprion similis]
MKISTSVVLILLLAGTINVEAGPLAAGICYAGCAAVVTACFAAAGFTFGTVPGAQIAAVPALVACNAGFATCEAACVAALLTPTP
>Dnox_XP_015363544.1 PREDICTED: uncharacterized protein LOC107161588 [Diuraphis noxia]
MVAQKMWSLILIGLLFSSSANAGPIXAGICYAGCAAVTVACFGAAGFTFGTVPGAVIAATPALAACNAAFGICEASCVAALVVPTP
>Fcan_OXA63722.1 hypothetical protein Fcan01_01313 [Folsomia candida]
MTTQFHTVFTLFMATLLVNSALGGPAAAGVCYAGCSAVVVACYAAAGFTFGTVTAGVGTPAAIIACNSAFGICESACVAALLMPTP
>Fari_XP_011307298.1 uncharacterized protein [Fopius arisanus]
MASTKVIIGMILLIIAGDTMAGPLAYGICQTGCNAVAVACYAGAGATFGVVTAGAGVAPAILACNAALGVCMTACVAAGFAPTP
>Ffus_KAK3929298.1 putative disease resistance protein [Frankliniella fusca]
MLQPFIMKLVLALLAVAALLNSATAGPAAYGICQSGCNALAVACYLAAGSVMGVGSVPACDAALGVCMAACIAAGAAPTP
>Focc_KAE8743865.1 hypothetical protein FOCC_FOCC009496 [Frankliniella occidentalis]
MKLILVLLAVAALLGSATGGPAAYGICQTGCNALAVACYASAGVVMGVGSVIPCNIALGTCMAACIAAGAAPTP
>Hvig_KAK9871193.1 hypothetical protein WA026_011474 [Henosepilachna vigintioctopunctata]
MVLETKITCLHIFIDIKMPLLTYGICQAACAAVVVACFSAAGVTFGTVPATLIAATPALAACNTAYASCYAACSPLILSPI
>Lbou_XP_051172728.1 uncharacterized protein LOC127289023 [Leptopilina boulardi]
MINRKLTILIVLALANLTTAGPLAAGICYGGCALVVCACFSAAGFTFGTVPGAQIAAVPALAACNSAFGSCMALCSAAVVAPTP
>Lhet_XP_043465910.1 uncharacterized protein LOC122500848 [Leptopilina heterotoma]
MVNQKLIIALVLALANVATAGPLAAGICYAGCAGVVCACFAAAGVVFGTVPLSVIAATPALAGCNSAFATCMSLCSAAVIAPTP

>Meup_CAI6362607.1 unnamed protein product [Macrosiphum euphorbiae] Hemiptera
*MVGRKMLSLMLVGLLIASSANA*GPIAAGICYAGCAGVTVACFSAAGFTFGTVPGAVIAATPALAACNAAFGICEASCMAALFVPVP
>Meup_CAI6365037.1 unnamed protein product [Macrosiphum euphorbiae] Hemiptera
*MVAQKIWALMLAGLLIASSANA*GPIAAGICYAGCAAVTVACFAAAGFTFGTVPGAVIAATPALAACNAAFGICEASCVAALIVPTP
>Meup_CAI6365297.1 unnamed protein product [Macrosiphum euphorbiae] Hemiptera
*MVAQKIWALMLAGLLVASSANA*GPIAAGICYAGCAAVTVACFGAAGFTFGTVPGAVIAATPALAACNAAFGICEASCVAALIVPTP
>Musi_KAJ1531914.1 hypothetical protein ONE63_000557 [Megalurothrips usitatus]
MRLLYLLVVAMLLLAAVAPPVQGGPAAYGVCQSGCNALAVACYLAAGSVMGVGSVPACNAALGVCMTACIAAGAAPTI
>Msac_XP_025194587.1 uncharacterized protein LOC112594148 [Melanaphis sacchari]
MVAQKMLSLVLVGLLLSSSAHAGPLTAGICYAGCAAVTVACFSAAGFTFGTVPGAVIAATPALAACNAAFGICEASCVAALVVPVP
>Mdir_XP_060877949.1 uncharacterized protein LOC132950475 [Metopolophium dirhodum]
MVAQKLWSLIFVGLLISSSANAGPIAAGICYAGCAAVTVACFSAAGFTFGTVPGAVIAATPMLAACNAAFGICEASCVAALIVPVP
>Mper_XP_022178996.1 uncharacterized protein LOC111039710 [Myzus persicae]
MVAQKMWSFLLVGLLLSSSASAGPIAAGICYAGCAAVTVACFSAAGFTFGTVPGAVIAATPVLAACNAAFGFCEASCVAALIVPTP
>Mper_XP_022178987.1 uncharacterized protein LOC111039701 [Myzus persicae]
MVAQKMWSLILVAILLSSSANAGPIAAGVCYAGCAAVTVACFSAAGFTFGTVPGAVIAATPVLAACNAAFGVCEASCVAALFVPVP
>Nfab_XP_046430417.1 uncharacterized protein LOC124184582 [Neodiprion fabricii]
MKISTSAVLMLLVAGIINVEAGPVAAGICYAGCGALVTACFAAAGFTFGTVPGAQIAAVPALVGCNTAFATCEAACMAALFTPTP
>Nlec_XP_015509762.1 uncharacterized protein LOC107216940 [Neodiprion lecontei]
MKISTSAVLMLLVAGIINVEAGPVAAGICYAGCGALVTACFAAAGFTFGTVPGAQIAAVPALVGCNTAFGTCEAACMAALFTPTP
>Odal_CAL8100433.1 unnamed protein product [Orchesella dallaii]
MMAGFKNFGTIKTFGVICLLLASMAVNEADAGLGLGALCSAGCATMAVACYSAAGAVFGTVTAGIGTAPAILACNAAFGTCMGNCAIATAAPTP
>Pcor_KAK7576216.1 hypothetical protein V9T40_012502 [Parthenolecanium corni]
MKQVSVFVFILLLSTNLVSAGPAASGICYAGCAGVTVACFAAAGFTFGTVPGAVIAATPALATCNAAFGACEAACMAAFFLPTP
>Rmai_XP_026822757.1 uncharacterized protein LOC113560848 [Rhopalosiphum maidis]
MVGQKMLSLLMVGLLLSPPALAGPVAAGICYAGCAAVTVACFTAAGFTFGTVPASVIAATPVLAACNTAFGVCEASCVAALVVPTP
>Rfus_KAK9499088.1 hypothetical protein O3M35_003600 [Rhynocoris fuscipes]
MKMINLTATIIIVLLMTGEINGGLEAAGVCHFGCAKLVTACFSAAGYVFGTVPLTEISSNPVLIACNTVFAACETASLAALANPVL
>Sher_KAL5242494.1 hypothetical protein ACI65C_009904 [Semiaphis heraclei]
MWSLILVGLLFSSSANAGPIAAGVCYAGCAAVTVACFAAAGFTFGTVPGAVIAATPALAACNAAFGICEASCVAALIVPVP
>Tkay_KAL3393024.1 hypothetical protein TKK_012302 [Trichogramma kaykai]
MQRHVLLALVLVVALLCQSSDAGLIGAALCYSGCSAVGVACFAAAGFGFTVPGAVIAATPALVACNAALAKCMSVCTVAVVAPTP
>Tkay_KAL3393019.1 hypothetical protein TKK_012297 [Trichogramma kaykai]
MPKHVFLALVLVAASLCQRSQAIGVSLCYSGCSAVGVACFAAAGFGFTVPGAVIAATPALVACNAALVKCMSRCTK
>Tpal_XP_034252786.1 uncharacterized protein LOC117652178 [Thrips palmi]
MRSSSFLLAVLVLAAVLGPVQVMGGPAAYGICQSGCNALAVACYLAAGSVMGVGSVPACNVALGVCMTACIAAGAAPTL
>Ufor_KAL4113046.1 hypothetical protein QTP88_016747 [Uroleucon formosanum]
MWSMLFVGLLISSSANAGPIAAGICYAGCAAVTVACFSAAGFTFGTVPGALIAATPALAACNAAFGICEASCVAALVVPVP

#### Fungi HLP sequences most similar to the above Hexapoda HLPs

>FUN_XP_007922500.1 uncharacterized protein MYCFIDRAFT_43367 [Pseudocercospora fijiensis CIRAD86]
MQFQKIFTLSMLATYVTAGPAAYGICQAGCAGVTVACYSAAGFVFGVALPAAPPAILACNAAFGSCQAACWAALIAPTP
>FUN_KUI64379.1 hypothetical protein VM1G_11180 [Cytospora mali]
MQPIKMLAVLTMATTATAGPIGYGICQAGCSAVVTACYAAAGVTFGTIAALAAPAAIVGCNTAFGTCQAACAAVLLTPTP
>FUN_KAG1470267.1 hypothetical protein G6F56_002783 [Rhizopus delemar]
MKIQVFVLLILSFLVCICQAGPISYAICQTGCNAVGVACYSAAGFVFGTITGGLGAPPAVIACNAGLGVCMAACVAAGCTPTP
>FUN_KAG1142548.1 hypothetical protein G6F38_007664 [Rhizopus arrhizus]
MKKLIVIYLITALLVSTSYAGLLAYGICQTGCNALAVACYSAAGFTFGTVTGGLAVPPAIACCNVALGTCMAGCVAAGAAPIP
>FUN_KAE8155953.1 hypothetical protein BDV40DRAFT_282490 [Aspergillus tamarii]
MKILYPALLILLSSITQVNGGPAAYGICQAGCAAVVTACYTAAGFTWGATLGATAPASIVACNTAFGTCQAACATALLAPTP
>FUN_RCI02276.1 hypothetical protein CU098_008242 [Rhizopus stolonifer]
MKYQVVALLIIVFLACSCYAGPLSYGLCQSGCNVVAVACYSAAGFTFGTVSAGLLAPPAIIGCNVALGTCMTACVAAGCAPVP
>FUN_KAK7545839.1 hypothetical protein IWX46DRAFT_92887 [Phyllosticta citricarpa]
MRLTNLMTSLAVVTSATAGPLGYGICQAGCSGVVVACYAAAGFTFGTVLAVAAPPAILVCNSAYGTCQAACAAVLLAPTL
>FUN_KXT04810.1 hypothetical protein AC578_9754 [Pseudocercospora eumusae]
MKFTKIALPILASFVPLVKAGPAAYGVCQAGCAGLAVACYAAAGFTFGVALPAAPPAILACNATFGSCQAACWAALFTPTP
>FUN_KAF9114376.1 hypothetical protein BGX27_011009 [Mortierella sp. AM989]
MSYLFTYLIVLLTILGLSSAGPLAYGICQTGCNGLAVACYTAAGFTFGTLTAGLGIPAVIVGCNTGLGTCMVACVVAGFAPTP
>FUN_KAL0144254.1 hypothetical protein V8B55DRAFT_1573037 [Mucor lusitanicus]
MLKTVVYCLVLLGFLSYVSAGPLAYGICQTGCNALAVACYSAAGFTFGTVTAGAGVPAVILACNAAQGFCMAGCVAAGCAPIP
>FUN_ORY12622.1 hypothetical protein BCR34DRAFT_289991 [Clohesyomyces aquaticus]
MRLSNLSIAAAALLMLPDFTSAGPAAYGVCQAGCAAVVVACYGAAGFTWGATLGASAPATIIACNTTFGTCQAACWAALFTPTP
>FUN_XP_066624638.1 uncharacterized protein IWZ02DRAFT_209063 [Phyllosticta citriasiana]
MRLTNLMTSLAVVTSATAGPLGYGICQAGCSGVVVACYAAAGSTFGTVLAVAAPPAILVCNSTYGACQAACAAVLLAPTL
>FUN_KAF7952734.1 hypothetical protein EAE96_005964 [Botrytis aclada]
MKPTSTSILLIAFLAGITTAGPVAYGVCQSGCAAVVMACYGAGGATWGATLGATAPATIVACNTAFGICSAKCAGLLVAPIP
>FUN_KAL2820147.1 hypothetical protein BDW59DRAFT_174604 [Aspergillus cavernicola]
MKRLSTSAVLSLLLVTSKVNAGPAAYGVCQAGCSAVVMACYAAAGFTWGATMGASAPASVIACNSAFGTCQAACAAVLLAPTL
>FUN_KAI1176761.1 hypothetical protein F4777DRAFT_545056 [Nemania sp. FL0916]
MKLNTHLVSALAFTSAASAGPIAYGLCQAGCAAVVTACYGAAGFTWGATMGASAPASIVACNTAFGACQAGCWAAAIAPTP
>FUN_KAI4224460.1 MAG: hypothetical protein LQ349_007235 [Xanthoria aureola]
MKLTNLLTTFASLTTLTSAGPAGYGVCQAGCSAVVVACYSGAGFTFGTVTAGAAVPAAIVACNSAYGTCQAACAAVLIAPTP
>FUN_KAI9238970.1 MAG: hypothetical protein BYD32DRAFT_248401 [Podila humilis]
MKFQAILYYVVLFLMIIGLTNAGPLAYGICQTGCNAVVVACYTGAGATFGTVTAGAGIPAAIIGCNSALGVCMAACVAAGFAPTL

```
>FUN_XP_051460734.1 uncharacterized protein EV154DRAFT_459735 [Mucor mucedo]
MVNAFLKLCLFVVVVSCLVGQSYAGPLAYGICQTGCNALVVTCYTAAGAVFGTVTAGAGVPAAILGCNAGLGLCMAGCIAAGFAPTP
>FUN_KAI8355641.1 hypothetical protein EDC96DRAFT_484348 [Choanephora cucurbitarum]
MLKLTFVLLIVCLLMGLSEAGPLAYGICQTGCNALAVACYAGAGFTFGTITAGAGVPAVILGCNAALGTCMAACVAAGLAPIP
>FUN_KAK5797245.1 hypothetical protein F5H01DRAFT_285529 [Linnemannia elongata]
MTSRYNLLYVVLFFMILGLANAGPLAYGICQSGCNSLVVACYAAAGVTFGTITAGAGIPAAVVACNTALGTCMVACVAAGCAPTL
>FUN_XP_009229506.1 hypothetical protein GGTG_13336 [Gaeumannomyces tritici R3-111a-1]
MRRFTSASIMLVTMAFTSPAFAGPAAYGVCQAGCAAVVMACYSAAGFTWGATLGVSAPPTIIACNTSFGTCQAACAAVLLSPTP
>FUN_RYP24059.1 hypothetical protein DL765_000787 [Monosporascus sp. GIB2]
MKLTTSILLAIVAAVPTIHAGPAAYGACQAGCSAVVQACYAAAGFTWGATLGATAPASIVACNNAYGACQAACWAALFSPTP
>FUN_KAF9925183.1 hypothetical protein FBU30_004990 [Linnemannia zychae]
MQVKLPLILSIVGFANAGPGLYGICQTGCNALVVACYSAAGATFGTVTAGVGVAPAIIACNAALGTCMAGCVAAGFSPTP
>FUN_RXW22619.1 hypothetical protein EST38_g3243 [Candolleomyces aberdarensis]
MRPSLLLIPVLAASTAQAGLIAYGICQTGCNAVTVACYAAAGFTFGTIAAPLAPPAIVACNAGLGTCSTACATVALLAPTP
>FUN_KAJ8594645.1 hypothetical protein M405DRAFT_808728 [Rhizopogon salebrosus TDB-379]
MNYKHTAILLVAAIASPAVVAGPLGYAICQTGCNALAVACYAGAGFTFGVALPAAPPVIIACNAGLGTCMAACAVVALGPTP
>FUN_KAK3831940.1 MAG: hypothetical protein J3R72DRAFT_454504 [Linnemannia gamsii]
MNSRVLLFILVLLSILGLTTAGPLAYGICQSGCNALAVACYGAAGVTFGTMTAGAGIPAVIVGCNTGLGTCMVACIAAGFAPTL
```

#### Supplementary File S10. HLPs in Arthropoda(excluding Hexapoda)

```
>UYV66577.1 hypothetical protein LAZ67_4002162 [Cordylochernes scorpioides]
MLKVAVLLALLASAHAGPLIAGTCYAGCAALAVACFSAAGFVFGTVPGAQIAAVPALVKCNLAFGACEAACVAALVAPTP
>KZS03801.1 Uncharacterized protein APZ42_033386 [Daphnia magna]
MKLPSTLFVLLGVLPLLTVAGPLAYGICQTGCNAVVVACYAAAGFTFGTVTAGAGVPAAIVACNAALGVCMAGCIAAGFAPTP
>KZS10595.1 Uncharacterized protein APZ42_024884 [Daphnia magna]
MRHTSLFVLGLLISFPLLAESGLVAYGVCQTGCNALAVACYAAAGFTFGVSTFGAGIPAAIVGCNGALGICMAACAATLFAPTP
>KAK4036940.1 hypothetical protein OUZ56_028988 [Daphnia magna]
MKFLLAVCVLMAFLPLSTVAGPLAYGICQTGCNAGAVACYAAAGFTFGAVTAGASTPLVIMGCNGALGICMAGCVAAGFTPTL
>XP_032792670.1 uncharacterized protein LOC116929498 [Daphnia magna]
MKFLLAVCVLMAFLPLSTVAGPLVYGICQTGCNAGAVACYAAAGFTFGAVTAGASTPLVIMGCNGALGICMAGCVAAGFTPTL
>KAK4010891.1 hypothetical protein OUZ56_020014 [Daphnia magna]
MKLPSTLFVLLGVLPLLTVAGPLAYGICQTGCNAVVVACYAAAGFTFGTVTAGAGVPAAIVACNAALGIFEPKQHVT
>EFX79605.1 hypothetical protein DAPPUDRAFT_319504 [Daphnia pulex]
MKPFLIVFILIGLFTFLSEAGPLAYGLCQSGCNALVVACYAGAGFTFGTVTAGVGIPAAIVGCNAALGVCMAACIAAGLAPTP
>EFX62814.1 hypothetical protein DAPPUDRAFT_309423 [Daphnia pulex]
MKLFSTLFLLFGLLPFATVAGPLAYGICQSGCNAVVVACYAAGGFTFGTVTAGAGIPAAIIACNAGLGVCMAACIAAGFAPTP
>EFX79607.1 hypothetical protein DAPPUDRAFT_319502 [Daphnia pulex]
MKLFSTLFFLFGLLPFVTVAGPLAYGIYQSGCNAVVVACYAAGGFTFGTVTAGAGIPAAIIACNAGLGVCMAACIAAGFAPTP
>EFX65051.1 hypothetical protein DAPPUDRAFT_231870 [Daphnia pulex]
MKSTVAVCYILVALLPLMAVAGPLAYGICQTGCNAAVVACYTAGGATFGTVTAGVGVPAVIMGCNAALGVCMAGCVAAGFAPTL
>EFX65052.1 hypothetical protein DAPPUDRAFT_65748 [Daphnia pulex]
MTTALPLLTEGGIISYGICQTGCNAVAVACYAAAGFTFGVPTWGATIPAVLASCNSALGICMGVCALVIPLPAP
>EFX79606.1 hypothetical protein DAPPUDRAFT_319503 [Daphnia pulex]
MGGVRPLFSTLFLLFGLLPFVTVAGPLAYGICQTGCNAVVVACYAAAGFTFGTVTAGAGIPAAVIACNAGLGVCMAACIAAGFAPTP
>KAI9558907.1 hypothetical protein GHT06_015696 [Daphnia sinensis]
MKLNSTLFVLLSVLPLLTVAGPLAYGICQTGCNAVVVACYAGAGFTFGTVTAGAGVPAAIIACNAALGVCMAGCIAAGFAPTP
>KAI9552354.1 hypothetical protein GHT06_022719 [Daphnia sinensis]
MKFLLAVCVLMALLPLSTVAGPLAYGVCQTGCNAGVVACYAAAGFTFGTVTAGASTPLVIMGCNAALGVCMAGCVAAGFAPTL
>RWR98541.1 hypothetical protein B4U79_05045, partial [Dinothrombium tinctorium]
CFGGPLAAGICYAGCAAIVVACFSAAGFTFGTVPGAVIAATPALAACNAAFSTCMASCSAAVIAPTP
>RWS02174.1 uncharacterized protein B4U79_12827 [Dinothrombium tinctorium]
MATPKLSISLLFLMMIASNCVCGPVAVGICYAGCAALACACFSAAGFTFGTVPISIIAATPALATCNSAFATCMAACSAALVAPTP
>KAI1290199.1 hypothetical protein HDE_08419 [Halotydeus destructor]
MVKSLIHGLLLALVLVSSADALFASYGVCQAGCAVAVTACYAAAGAVFGTVTAGVGTPAAILACNAGQAACYAGCAAALVMPIP
>KAI1289864.1 hypothetical protein HDE_08452 [Halotydeus destructor]
MQRPMLGLFILVLLASNAMAGPLLYGICQAGCAAVVVACYTAAGAVFGTVAAPAAPAAIVGCNSAFGTCSASCAAATILAPTP
>KAI1289865.1 hypothetical protein HDE_08453 [Halotydeus destructor]
MQRPMLGLFILVLLASNTMAGPLLYGICQAGCATVAVACYAAGGAVFGTVLAAAASPAIAGCNSAFGTCSAACAKVTLGAPTP
>OQV14256.1 hypothetical protein BV898_11493 [Hypsibius exemplaris]
MNTRTSTAICLVFLMAVAQVHSGLIGILAYGACQTGCNAGWVACCSAAGVTAGTVTAGLGVPAAVMACSALQGTCMAACAALALAPTP
>CAD7622705.1 unnamed protein product [Medioppia subpectinata]
MSTKLALLLIAVIVVILVSESSAGPVSWAACQTACNVGYVTCCVIAGGIAGTFTLVGAPAALAACSTVQGSCMAACTPLLLAPTP
>CAD7624846.1 unnamed protein product [Medioppia subpectinata]
MKIMVKLLIRIMVVLLTAHESMAGILGLLGGYGLCQTACNAAWVACLASAGVVAGTVTAGAGAPAAVLACNALQGVCMGSCAVTFLVAPTP
>GAV05447.1 hypothetical protein RvY_15580 [Ramazzottius varieornatus]
MMSRSIVIFLGLLLVTDASAGPLAVAAAQAGCAAVQVACYAGAGAVFGTVTAGVGTPAAILACNSAFGTCMAAAYWAFFLPTV
```

```
RWR98541.1     ----------------------CFGGPL----AAGICYAGCAAIVVACFSAAGFTFGTVPGAVIAATPALAACNAAFSTCMASCS-AAVIAPTP
RWS02174.1     ---MATPKLSISLLFLMMIASNCVCGPV----AVGICYAGCAALACACFSAAGFTFGTVPISIIAATPALATCNSAFATCMAACS-AALVAPTP
UYV66577.1     ---------MLKVAVLLALLASAHAGPL----IAGTCYAGCAALAVACFSAAGFVFGTVPGAQIAAVPALVKCNLAFGACEAACV-AALVAPTP
KAI1289864.1   -----MQRPMLGLFILVLLASNAMAGPL----LYGICQAGCAAVVVACYTAAGAVFGTV--AAPAAPAAIVGCNSAFGTCSASCAAATILAPTP
KAI1289865.1   -----MQRPMLGLFILVLLASNTMAGPL----LYGICQAGCATVAVACYAAGGAVFGTV--LAAAASPAIAGCNSAFGTCSAACAKVTLGAPTP
KAI1290199.1   ----MVKSLIHGLLLALVLVSSADALFA----SYGVCQAGCAVAVTACYAAAGAVFGTVT-AGVGTPAAILACNAGQAACYAGCAAALVMPIP-
GAV05447.1     -----MMSRSIVIFLGLLLVTDASAGPL----AVAAAQAGCAAVQVACYAGAGAVFGTVT-AGVGTPAAILACNSAFGTCMAAAYWAFFLPTV-
KZS03801.1     -----MKLPSTLFVLLGVLPLLTVAGPL----AYGICQTGCNAVVVACYAAAGFTFGTVT-AGAGVPAAIVACNAALGVCMAGCIAAGFAPTP-
KAI9558907.1   -----MKLNSTLFVLLSVLPLLTVAGPL----AYGICQTGCNAVVVACYAGAGFTFGTVT-AGAGVPAAIIACNAALGVCMAGCIAAGFAPTP-
KAK4010891.1   -----MKLPSTLFVLLGVLPLLTVAGPL----AYGICQTGCNAVVVACYAAAGFTFGTVT-AGAGVPAAIVACNAALGIFEPKQHVT-------
EFX62814.1     -----MKLFSTLFLLFGLLPFATVAGPL----AYGICQSGCNAVVVACYAAGGFTFGTVT-AGAGIPAAIIACNAGLGVCMAACIAAGFAPTP-
EFX79607.1     -----MKLFSTLFFLFGLLPFVTVAGPL----AYGIYQSGCNAVVVACYAAGGFTFGTVT-AGAGIPAAIIACNAGLGVCMAACIAAGFAPTP-
EFX79606.1     -MGGVRPLFSTLFLLFGLLPFVTVAGPL----AYGICQTGCNAVVVACYAAAGFTFGTVT-AGAGIPAAVIACNAGLGVCMAACIAAGFAPTP-
EFX79605.1     -----MKPFLIVFILIGLFTFLSEAGPL----AYGLCQSGCNALVVACYAGAGFTFGTVT-AGVGIPAAIVGCNAALGVCMAACIAAGLAPTP-
KAK4036940.1   -----MKFLLAVCVLMAFLPLSTVAGPL----AYGICQTGCNAGAVACYAAAGFTFGAVT-AGASTPLVIMGCNGALGICMAGCVAAGFTPTL-
XP_032792670.1 -----MKFLLAVCVLMAFLPLSTVAGPL----VYGICQTGCNAGAVACYAAAGFTFGAVT-AGASTPLVIMGCNGALGICMAGCVAAGFTPTL-
KAI9552354.1   -----MKFLLAVCVLMALLPLSTVAGPL----AYGVCQTGCNAGVVACYAAAGFTFGTVT-AGASTPLVIMGCNAALGVCMAGCVAAGFAPTL-
EFX65051.1     ----MKSTVAVCYILVALLPLMAVAGPL----AYGICQTGCNAAVVACYTAGGATFGTVT-AGVGVPAVIMGCNAALGVCMAGCVAAGFAPTL-
KZS10595.1     ----MRHTSLFVLGLLISFPLLAESGLV----AYGVCQTGCNALAVACYAAAGFTFGVST-FGAGIPAAIVGCNGALGICMAACAATLFAPTP-
EFX65052.1     --------------MTTALPLLTEGGII----SYGICQTGCNAVAVACYAAAGFTFGVPT-WGATIPAVLASCNSALGICMGVCALVIPLPAP-
OQV14256.1     MNTRTSTAICLVFLMAVAQVHSGLIGIL----AYGACQTGCNAGWVACCSAAGVTAGTVT-AGLGVPAAVMACSALQGTCMAACAA-LALAPTP
CAD7622705.1   --MSTKLALLLIAVIVVILVSESSAGPV----SWAACQTACNVGYVTCCVIAGGIAGTFT-L-VGAPAALAACSTVQGSCMAACTP-LLLAPTP
CAD7624846.1   --MKIMVKLLIRIMVVLLTAHESMAGILGLLGGYGLCQTACNAAWVACLASAGVVAGTVT-AGAGAPAAVLACNALQGVCMGSCAVTFLVAPTP
```

**Supplementary file S11. HLP sequences in other organisms (excluding Fungi, Arthropods)**

#### Nematodes

>NEM_KAH7680518.1 cysteine-rich protein, partial [Aphelenchoides avenae]
MKLRVILMLFAIFDLTGGVLAYGICQAGCAGLAAACYSAAGYVFGTVTVGAGTPAAILTCNKAFGVCSAKCALIALAP
>NEM_KAI6186997.1 hypothetical protein M3Y98_00193900 [Aphelenchoides besseyi]
MRYISLLMFLLFFDFASSGPIGYGVCQAGCAGVVMACYSAAGYTWGATLGASAPATIVACNTAFGSCQAACASVLLMPTP
>NEM_KAI1697871.1 hypothetical protein DdX_18235 [Ditylenchus destructor]
MSTSAILILLFIGSVTGGPAAYGICQAGCAALVAACYAAAGAVFGTVTAGLGTPAAIIACNSSFGTCQGACWAALFSPTP
>NEM_KAI3410909.1 hypothetical protein GPALN_002991 [Globodera pallida]
MSCSKLASVLLILLLVVHLSNGGPMAYGICQAGCAALVVTCYTAAGAVFGTVTAGVATAPALLGCNVAFGKCQAACWAALMLPTA
>NEM_KAL3070285.1 hypothetical protein niasHS_016112 [Heterodera schachtii]
MAQKFTAVFTVFLLLSLVANSHAGPAAYGVCQAGCAALVVTCYVAAGATFGTVTAGAGTPAIILGCNSAFGTCQAACWAALFAPTP
>NEM_KAL3070296.1 hypothetical protein niasHS_016123 [Heterodera schachtii]
MAKKITLSSALFAVLLMISLLVAHSHAGPAAYGVCQAGCAAVVVACYGAAGVVFGTITAGVGTPPAIMACNAAYGSCQAACWAALFSPTL
>NEM_KAL3122511.1 hypothetical protein niasHT_003047 [Heterodera trifolii]
MVKKISASSSSFITSFILLLLSLNPAIGGPAAYGVCQAGCSAVVVACYAAAGAVFGTVTAGIGTPHAILACNTAYGSCQSACWTALFSPTP
>NEM_KAL3110422.1 hypothetical protein niasHT_018252 [Heterodera trifolii]
MAHKITASSPSVVVIIILLLISIHSAISGPAAYGVCQAGCSAVVVACYAAAGAVFGTVTAGIGTPHAIVACNTAYGTCQSACWAALYSPTV
>NEM_KAL3111509.1 hypothetical protein niasHT_018284 [Heterodera trifolii]
MAQAQKITVFSAVFATILLLSLVDNSSAGPAAYGVCQAGCAALVVTCYVAAGATFGTVTAGAGTPAIILACNSAFGTCQAACWAALIAPTP
>NEM_KAL3086999.1 hypothetical protein niasHT_025523 [Heterodera trifolii]
MAQKFTAVFTVFLLLSLVANSHAGPAAYGVCQAGCAALVVTCYLAAGATFGTVTAGAGTPAIILGCNSAFGTCQAACWAALFAPTP
>NEM_KAL3122504.1 hypothetical protein niasHT_003040 [Heterodera trifolii]
MAQKFTAVFTVFLLLSLVANSHAGPAAYGVCQAGCAALVVTCYVAAGATFGTVTAGAGTPAIILGCNSAFGTCQAACWAALFAPTP
>NEM_KAL3087012.1 hypothetical protein niasHT_025536 [Heterodera trifolii]
MAKKITLSSALFSLILLLLSLVATSRAGPAAYGVCQAGCAAVVVACYGAAGVVFGTITAGVGTPPAIMACNAAYGSCQAACWAALFTPTL
>NEM_KAL3111496.1 hypothetical protein niasHT_018271 [Heterodera trifolii]
MAKKITLSSALFAVLLLLSLVATSRAGPAAYGVCQAGCAAVVVACYGAAGVVFGTITAGVGTPPAIMACNAAYGSCQAACWAALFTPTL
>NEM_KAL3122513.1 hypothetical protein niasHT_003049 [Heterodera trifolii]
MAKKITLSSALFALILLLSLVAHSRAGPAAYGVCQAGCAAVVVACYGAAGVVFGTITAGVGTPPAIMACNAAYGSCQAACWAALFTPTL
>NEM_KAL3087010.1 hypothetical protein niasHT_025534 [Heterodera trifolii]
MAHKITASSPSVVVITILFLISFHSAIAGPAAYGVCQAGCSAVVVACYAAAGAVFGTVTAGIGTPHAIVACNTAYGTCQSACWTALFSPTP
>NEM_KAL3070294.1 hypothetical protein niasHS_016121 [Heterodera schachtii]
MVKKITESSPYVAIITILLLLSLNPAIAGPAAYGICQAGCSAVVVACYAAAGAVFGTVTAGIGTPHAIVACNTAYGTCQSACWAALYSPTL
>NEM_VDM39105.1 unnamed protein product [Toxocara canis]
MRISMIYVFIILYSTVLITEGGPITYVACVAACNAAVVACYASLGFVFGTVTAGAGTPAAVLACNAAQSACWVGCSAGGVAPTP

#### Other organisms

>KAK2550313.1 hypothetical protein P5673_029003 [Acropora cervicornis]
MLLLLTRTARPGPLAYGICQTGCNTVWVACVAAAGGVAGVSTGGAGVPASILACNSAQGVCMAVCVAAGLLPTL
>KAK2554542.1 hypothetical protein P5673_024000 [Acropora cervicornis]
MNAVVRSFIVLALLTSTARSGPLAYGICQTGCNTVWVACVTAAGGVAGVSTGGAAVPASILACNAAQGVCMAACVAAGLSPTL
>CAF0773123.1 unnamed protein product [Adineta ricciae]
MKYTTILVFLLLLMSSCTKAGPVAYGMCQAGCASVVVACYSAAGAVFGTVTGGVGAPPALIACNLAFGKCSAVCAAIALTPTP
>CAF0915355.1 unnamed protein product [Adineta ricciae]
MKYTTILVFLLLLMTSCTKAGPLAYGMCQAGCASVVVACYSAAGAVFGTVTGGLGAPPAIIACNLAFGKCSAVCAAIALTPTP
>CAF1468521.1 unnamed protein product [Adineta steineri]
*MKFANQMKILLIFLFTLSTFVSG*GPLSYAACQTACNKGAMTCYGIAGVTFGVGAVPACGLAQGSCMAACTPLLVAPTP
>UJR08264.1 hypothetical protein I4U23_012537 [Adineta vaga]
MKLTALILSFILLLSSCVQGGPAVYGICQAAAAGAVFGTVTAGAGTVPAIVACNAAFGQCSAACAALVLTPTP
>CAF1048684.1 unnamed protein product [Adineta steineri]
MQFANQMKILLIFLCILSHSVSGGPISYGTCQTACNKGAMSCYGSAGVIFGVGAVLACGLAQGSCMAACTPF
>CAF0783321.1 unnamed protein product [Adineta steineri]
MQFANQMKILLIFLFILSHSASGGPLTYAACQTACNKGAMSCYGSAGVIFGVGAVLTCGLAQGSCMVACTPLLAPIIP
>CAF1409581.1 unnamed protein product [Adineta steineri]
MQFANQMKILLIFLFILSHSVSGGPLTYAACQTACNKGAMSCYGSAGVIFGVGAVLTCGLAQGSCMIACTPLLAPVTP
>CAF1047280.1 unnamed protein product [Adineta steineri]
MKLTVVLLCFIILTAPCIEGGPAAYGICQAGCATVTVACYAAAGAVFGTITAGVGTAPAILACNAAFGQCSLACIAAGCIPIP
>CAF1133829.1 unnamed protein product [Adineta steineri]
MKLAVILCFILLTATCIEGGPAAYGICQAGCAAVTVACYAAAGVVFGTITAGAATAPAILACNAAFGQCSAACIAAGFLPTP
>CAF1138014.1 unnamed protein product [Adineta steineri]
MKLAVILCFILLTATCIEGGPAAYGICQAGCAAVTVACYAAAGAVFGTITAGVATAPAILACNVAFGQCSAACVAAGLLPTP
>CAF1087552.1 unnamed protein product [Adineta steineri]
MMKFIVVFLCFGILTAPYAEAGLAAYGICQAGCCALAVACYGAAGAVFGTVTAGAATAPAILACNAAFGKCSAACIAAGFAPTP
>CAF1047241.1 unnamed protein product [Adineta steineri]
MMKFIVVFLCFGILTAPYAEAGLAAYGICQAGCCALAVACYGAAGAVFGTVTAGAGTAPAILACNAAFGKCSAACVAAGCAPTP
>UJR08263.1 hypothetical protein I4U23_012536 [Adineta vaga]
MKLATIFLSLILLTSTCVHSGPAAYGICQAGCAAMVVACYAAAGAVFGTVTAGAATAPAIIACNVAFGKCSAACIAAGFAPTP
>KAL6049488.1 Zygote-specific protein [Balamuthia mandrillaris]

*MKTLAVLFMTLIFTGLIVRSAEA*GPVAYGVCQAGCSALVVKCYAAAGVVFGTITAGAGTPAVILGCNAAFGLCSANCAIVALTAPTP
>CAF0985211.1 unnamed protein product [Brachionus calyciflorus]
MNQARLIFLLVFVSILMNAEAGPISGAICSACCAAGAVACYSAAGFVFGTVTAGAGTPAVLIGCNTAFGKCMAACAVAGVAPAP
>CAF1096744.1 unnamed protein product [Brachionus calyciflorus]
MEVKKIFSFLIIVSILINAEAGPISAAICASCCAAGVVACYTAAGFVFGTVTAGAGTPAVIIGCNTAFGKCMAACAAAAVSPIP
>CAF0844728.1 unnamed protein product [Brachionus calyciflorus]
MNPEAGPISAAICSACCAAGVVACYSTAGFVFGTVTAVAGTPAVIIGCNTAFGKCMAACAAAAVSPIP
>CAF1080100.1 unnamed protein product [Brachionus calyciflorus]
MKPTGLIILLVFVSILMNAEAGPISAAICSACCAGGVVACYSAAGFVFGTVTAGAGTPAVIIGCNTAFGKCMAACAVAGVAPAP
>CAF0844706.1 unnamed protein product [Brachionus calyciflorus]
MKPTRLIFLLVFVSILMNAEAGPISGAICSACCAAGVVACYSAAGFVFGTVTAGAGTPAVIIGCNTAFGKCMAACAVAGVAPAP
>XP_067822278.1 hypothetical protein CCR75_000535 [Bremia lactucae]
*MIKKLSTVLVLTAFVLAITKVNA*GPVAYGICQAGCNAIAAACYAAAGAKFGTVTAGVGTAPAIVGCNSALGACMVKCVAAGFLPIP
>KAG2439968.1 hypothetical protein HXX76_004087 [Chlamydomonas incerta]
MKGFVKSLLLLAVLAAACHATRAGPLAYGICQSGCNSVAVACYKAGGFIFGVPTWGAAVPATIGGCNAALGKCMAACVAAGASPTA
>KAG2440151.1 hypothetical protein HXX76_004264 [Chlamydomonas incerta]
MKGIAKSLLVLAILATAFTATHAGPFAYGICQTGCNAIAVACYAAAGFTFGVPTWGAAVPVAILACNAALGTCMAACVAAGLSPIP
>XP_042925655.1 uncharacterized protein CHLRE_03g151900v5 [Chlamydomonas reinhardtii]
MKGFAKSLLLALAILLATAFSATHAGPVAYGICQTGCNSVAVACYSAAGFTFGVPSWGAAAPSSIAACNVALGKCMSSCVAAGLWPI
>XP_001695656.1 uncharacterized protein CHLRE_03g151950v5 [Chlamydomonas reinhardtii]
MKGFAKSLLALAILATAFSATHAGPIAYGICQTGCNALAVACYAAAGFTFGVPSWGAAIPATVLACNAGLGTCMAACVAAGLSPIP
>PRW32660.1 cell envelope integrity [Chlorella sorokiniana]
*MKAAHVLLLLLLATMAAFAPGARA*GLLSAFAGYGACQTACNVAWVSCYAGTGLIAGTVTAGLGAPEMALACNAAQGLCMTACAGGLLLSPTP
>CAN0187930.1 unnamed protein product [Chordaria linearis]
MRGITVIALSALLAAMAAIAEAGPLAALTAYGSCQTGCNILVCACYASAGAVFGAATGGAAVPAAIIGCNAGLGSCMAACWAVTGAAAAAPTL
>CAM9621471.1 unnamed protein product [Choristocarpus tenellus]
MLLLIFLLASGTVLAGPLAAIAAYGSCQTGCNGVAVACYTAGGAVFGATTAGVALSPALLACNAALGTCMGTCAVVAGLAAVTPTF
>KAJ7387352.1 hypothetical protein OS493_004343 [Desmophyllum pertusum]
*MNNIVKLLIMVALFTNTAHS*GLLAYGICQTGCNAVWVACVTAAGGVAGVSTGGAAVPAAILACNFAQGVCMAACVAAGLSPTP
>KAJ7387358.1 hypothetical protein OS493_004349 [Desmophyllum pertusum]
*MNQAIVKFLIMLALFTSTANS*GPLAYGICQSGCNAVWVACCAAGGGVAGVSTGGAAVAPAFLACSAAQGVCMTACVAAGFAPTL
>KAJ7387350.1 hypothetical protein OS493_004341 [Desmophyllum pertusum]
*MNQAIVKVLIMLALFTNTAHS*GPLAYGICQTGCNFAWVTCVTAAGGVAGVSTGGVGVPAVILACNAAQGVCMAACIAAGFAPTL
>CAM9191507.1 unnamed protein product [Dictyota dichotoma]
MMAAIKVEAGPIAAIAAYGTCQSGCNALAVACYTAGGAIFGTLTAGLGIPAVIVGCNASLGTCMAACAAATAAAAVMPTP
>CAN0477318.1 unnamed protein product [Discosporangium mesarthrocarpum]
MKLTLLLVLLWIQVVVGGPFAAMVAYGLCQTGCNTVAVACYAAGGAVFGTVTAGVGVAPAIIGCNVALGVCMTSCATVAGIGVGAPTP
>CAM9369918.1 unnamed protein product [Ectocarpus fasciculatus]
*MLSALVVCLVLTASTVAG*GPAAALAAYGACQSGCNVLAVSCYAAAGFTFGVATGGAAVPAVIVSCNAALGTCMAACWGVTGAAAFAPTP
>CAM9769094.1 unnamed protein product [Ectocarpus sp. 4 AP-2014]
MKFFVSTLSALIVCLVLMASTVAGGPAAALAAYGSCQTGCNVVAVACYAAGGATFGVVTGGAGVPIAIAGCNTALGTCMAACWGVTGAAALAPTP
>CAM9742539.1 unnamed protein product [Ectocarpus sp. 1 AP-2014]
MKFFVSTLSALIVCLVLMASAVAGGPAAALAAYGSCQTGCNVVAVACYAAGGATFGVVTGGAGVPIAIAGCNTALGTCMAACWGVTGAAALAPTP
>CAM9614101.1 unnamed protein product [Ectocarpus sp. 9 AP-2014]
MKFFMSMLSALIVCLVFMATTVAGGPAEALAAYGSCQTGCNAVAVACYAAAGATFGVATGGAGVPIAIAGCNTALGTCMAACWGVTGAAALAPTP
>KAG2485775.1 hypothetical protein HYH03_015488 [Edaphochlamys debaryana]
MPRFGLKTLVCCLAIMAAISGADAGILSYGICQSGCNAVVVACYAAAGFTFGTVTAGAGIPAVLVGCNAGLGTCMAACVAAGLLPVP
>KAG2485774.1 hypothetical protein HYH03_015487 [Edaphochlamys debaryana]
MARISLALPLLLCLALGAALHGADAGVAAYGICQTGCNSLAAACYGAGGSVFGTVAAAFGAPASILGCNKALGACMSSCVAAGLTPSA
>KXJ27680.1 hypothetical protein AC249_AIPGENE16893 [Exaiptasia diaphana]
MNKAAALLLIMMVTAQQSQGGLLAYGICQTGCNTVWVACVAAAGGTAGVSTGGAAVPAAIVACNTAQGACMAACVAAGLTPTP
>GAX13526.1 hypothetical protein FisN_27Lh032 [Fistulifera solaris]
MNITAKTFLLLAVTTSSASAGLLSYGICQSGCNGMAVACYSSAGFVFGTVTAGAGIPAAIVGCNTALGCCMASCVVAGLSPVP
>GAX28313.1 hypothetical protein FisN_27Hh032 [Fistulifera solaris]
MKLSPKSLLLLAVTTSSASAGLLSYGICQSGCNGMAVACYSSAGFVFGAVTAGAGIPAAIVGCNTALGCCMASCVVAGISPVP
>GAX28312.1 hypothetical protein FisN_27Hu033 [Fistulifera solaris]
MYHESKSESPLSLGGRQHRVGGTLSYGICQTGCNAVAVACYASAGAVFGTVTAGAGVPAAIVGCNSALGVCMVSCIAAGCSPIP
>CAM9346533.1 unnamed protein product [Fucus serratus]
MKKSNVLTLLIFITVLGCLLPPVDGGPFAAMICYTGCNSLAVACYAAAGFTFGTVTAGVGTPAAILGCNAGLGTCMTACVAAGFTPTP
>KXZ52489.1 hypothetical protein GPECTOR_9g533 [Gonium pectorale]
MASKQLKSFIVVALLAIFVSSFKPVAAGPLAYGICQTGCNAMVVACYSAAGFTFGTVTAGTGIPAAIAACNAALGTCMAACVAAGCTPTP
>KXZ52487.1 hypothetical protein GPECTOR_9g531 [Gonium pectorale]
MVATLIKRFLLAGLIALLCVASFEPVAAGLFGYGVCQTGCNTLAGACYAAGGATFGTVTAGGGTPATITGCNPALGACMKSCVAAGLNPFG
>CAM9755229.1 unnamed protein product, partial [Himanthalia elongata]
TRCNILAVACYTAAGATFGTVTAGAGIPLVIVGCNAGLGTCMAACAVSTGLAPTP
>GFH17887.1 uncharacterized protein HaLaN_14609, partial [Haematococcus lacustris]
*MSISLTLRRIAASVAFAVLLLAACATQADA*GPLGALAAYGACQTGCNTVVCACYAALGVTFGAVGVAAIPACGAAQGVCMAACAQMTLAAVVAPP
>CAM9217243.1 unnamed protein product [Heribaudiella fluviatilis]
MRIKSIIVAALVPVVSLVNAGLFTAIAAYGVCQTGCNALAVACYTSAGFTFGTVTAGAALPAVIVGCNAALGTCMASCYVATAAAVAVPAP
>CAM9147544.1 unnamed protein product, partial [Heribaudiella fluviatilis]
SIIVAALVPVVPLVNAGLITAIAAYGVCQTGCNALAVACYTTAGFTFGTVTAGAALPAVIVGCNAALGTCMAGCYVATAAAVAVPAP
>GIQ91937.1 hypothetical protein KIPB_015417 [Kipferlia bialata]
MLSEASYGVCQVGCDAGAVACYSAAGFVFGTVTAGLGTPAAVLTCNSILSACMTTCAVRYLGEASAEGTGTICIY
>CAN0340601.1 unnamed protein product, partial [Laminaria digitata]
CNVVAVACYTAGGLTFGVPTGGAAVPAIALGCNSALGVCMAACWAVTGAAVFTPTP
>PAA85964.1 hypothetical protein BOX15_Mlig004286g3 [Macrostomum lignano]
MAIQFVTKVALLALLLIGGPQLQAEAGIIGAVASYGVCQTGCNTVWVACVAAAGGVAGVSTGGTAVPAAILACNAANGVCMAGCASVTLLAPTP

>PAA61898.1 hypothetical protein BOX15_Mlig019107g1 [Macrostomum lignano]
MATSFATAAVLTVSLLLTCGPLLQAEAGLFGALFGYGICQTGCNTVWVACVAAAGGVAGVSTGGTAVPAAILACNAAQGVCMAACASVTIAGPV
>PAA85965.1 hypothetical protein BOX15_Mlig019107g3 [Macrostomum lignano]
MATSFATAAVLTVSLLLTCGPLLQAEAGLFGALFGYGICQTGCNTVWVACVTAAGGVAGVSTGGTAVPAAILACNAAQGVCMAACASVTIAGPV
>XP_032218746.2 uncharacterized protein LOC116601956 [Nematostella vectensis]
*MAARKMLMLLALTMLIASSSA*GPLAYGICQTGCNAVWVTCVAAAGGVAGVSTGGAGVPAAVLACNAAQGVCMAACVAAGLSPTL
>KAF4681825.1 hypothetical protein FOZ60_011535 [Perkinsus olseni]
MPYIIIFVLAALELLLTEAGPLTYGICQAGCAAAWVSCNAACGVVAGTVTAGAATPACVLACNGAQGACYAACAAMVLTPTP
>CAH0481440.1 unnamed protein product [Peronospora belbahrii]
MLKTLTSVLVVLGLASTNQVTAGPFAYGICQTGCNAVAVACYAAGGAVFGTVTAGASTIPAIIACNAALGTCMAACVAAGLSPTP
>RQM11952.1 hypothetical protein DD237_007205 [Peronospora effusa]
MLKKSAIALVVLGLASTHKVTAGPLAYGICQTGCNAIVVACYAAAGAVFGTVTAGIGTMPAILACNVALGTCMASCVAAGCAPTP
>KAJ8610761.1 hypothetical protein MRB53_038312 [Persea americana]
*MHFRLIVTAVFTASTITSA*GPATYGLCQAGCSSLVMACYGAAGFTWGATLGATAPATVVACNFAFGTCQASCAAALLSPTP
>KAG6970997.1 hypothetical protein JG688_00004631 [Phytophthora aleatoria]
MLKKLIITLMAALASLPEVDAGILAYGICQSGCNAVAVACYSAAGAVFGTVTAGVGAIPAVIGCNGALGTCMAGCVAAGLSPTP
>KAG6970559.1 hypothetical protein JG688_00004803 [Phytophthora aleatoria]
MFKKLTAALAVLALASTSEVNAGPLAYGICQTGCSAVAAACYAAAGAVFGTVTGGIGTMPAIIGCNVALGTCMASCIAAGLTPTP
>KAG2763445.1 hypothetical protein Pcac1_g24884 [Phytophthora cactorum]
MFKKLTAALASTSEVNAGPLAYGICQTGCNAVAAACYAAAGAVFGTVTGGIGTMPAIIGCNVALGTCMASCIAAGLTPTP
>KAF1779425.1 hypothetical protein GQ600_19997 [Phytophthora cactorum]
*MLKKLIITLMAALASLPEVDA*GILAYGICQSGCNAVAVARYSAAGAVFGTVTAGVGAIPAVIGCNGALGTCMAGCVAAGLSPTP
>KAG1693246.1 hypothetical protein DVH05_023711 [Phytophthora capsici]
MNKLLTKLTAALVVLGLTATPEVSAGPLAYGICQTGCNAVAVACYAGSGAVFGTVTAGLGVAPAIIACNVALGTCMASCVAAGCTPTP
>KAK1943604.1 hypothetical protein P3T76_005000 [Phytophthora citrophthora]
MKFLNKLTATLVVLGLAATPEVNAGPLAYGICQSGCNAVAVACYAGAGAVFGTVTAGLGAAPAILGCNIGLGTCMAACVAAGCSPTP
>KAE8906787.1 hypothetical protein PF003_g9484 [Phytophthora fragariae]
MLKKIAAALVVLGLTASPEANAGLLAYGICQTGCNALAVACYTAAGAVFGTVTVGVGTLPAVIGCNVALGTCMASCVAAGLAPTP
>KAE8877033.1 hypothetical protein PF003_g38827 [Phytophthora fragariae]
MLKKLILALVVVFAPLHEVNAGPLAYGICQTGCNAVAVACYAAGGAVFGTMTAGVGTAPAVLACNAALGVCMAGCVAAGCTPTP
>KAG3113898.1 hypothetical protein PI125_g6907 [Phytophthora idaei]
MLKKLIITLMAALASLPEVDAGILAYGICQSGCNAVAVACYSAAGAVFGTVTAGVGAIPAVIGCNGALGTCMAGCVAAGLSPTP
>KAG3170974.1 hypothetical protein PI126_g2096 [Phytophthora idaei]
MFKKLTAALAVLALASTSEVNAGPLAYGICQTGCNAVAVACYAAAGAVFGTVTGGIGTMPAIIGCNVALGTCMASCIAAGLTPTP
>OWZ04856.1 hypothetical protein PHMEG_00023169 [Phytophthora megakarya]
MQVKLGGGVLLLALTTIPEAYAGPLAYGICQTGCNAVAVACYGAAGAVFGTVTAGVATLPAIISCNAALGVCMASCVAAGLTPTL
>OWY98340.1 hypothetical protein PHMEG_00030918 [Phytophthora megakarya]
MLKKLIFALLVVLTPLPESDAGPLAYGICQSGCNGVAVACYAAAGAVFGTVTAGVGAAPAVVGCNIALGACMTGCVAAGLAPTP
>ETI30167.1 hypothetical protein F443_22718 [Phytophthora nicotianae P1569]
MLKKLILALVAVFAPLPEVDAGILAYGICQTGCNAVVVACYSAAGAVFGTVTAGVGTIPAVVACNGALGTCMAGCVAAGLSPTP
>KAL3658320.1 hypothetical protein V7S43_016705 [Phytophthora oleae]
MLNKLAATLVVLGLAMSEVQGGPLAYAICQTGCNAVVAACYGGAGAVFGTVTAGAGVAPAVIACNAALGTCMAACVAAGCTPTP
>KAL3658321.1 hypothetical protein V7S43_016706 [Phytophthora oleae]
MSKKAIVAIVVVLGLASLPEVHGGPLAYGICQTGCNALVVACYGAAGAVFGTVTAGIGTLPAIITCNAALGTCMTGCVAAGCTPTP
>KAE8975738.1 hypothetical protein PR002_g25512 [Phytophthora rubi]
MLKKIAAALVVLGLTASPEANAGPLAYGICQTGCNALAVACYTAAGAVFGTVTVGVGTLPAVIGCNVALGTCMASCVAAGLAPTP
>KAE8975739.1 hypothetical protein PR002_g25513 [Phytophthora rubi]
MLMKFTAAFVVLAVAVSTKVNAGPLAFGICQAGCNAAAVACYSSAGVLFLPAAFGCAALSTCMAVCAAAGMAPTP
>XP_009530821.1 hypothetical protein PHYSODRAFT_514663 [Phytophthora sojae]
MKLAAVLVALGLAAAPEVSAGPLAYGICQTGCNAVAVACYSAAGVVFGTVTAGVGSIPAVIGCNAALGTCMAGCVAAGLSPTP
>XP_009530822.1 hypothetical protein PHYSODRAFT_514712 [Phytophthora sojae]
MLKKLTAALVVLGLAAAVNAGPAAYGICQTGCNVLAVACYAATGAVFGTAIPAIINCNKALGVCMASCVAAGLIPTP
>CAM9955383.1 unnamed protein product [Pleurocladia lacustris]
MIKFVALIMLMLSVTVDGGPLSALAAYGTCQTGCNAMAVACYAAARCTFGVATGGAAAPAVVIGCNSALGTCMAACWAATGSIFVTPTP
>CAH3111068.1 unnamed protein product [Porites lobata]
*MNAVTKIFILFTLFSTAQS*GPFAYAICQTGCNTVWVACVAAAGGTAGVSTGGAAVPAAILACNAAQGTCMAACVAAGLSPTP
>GLD92447.1 hypothetical protein PINS_up000980 [Pythium insidiosum]
MQQRSILFTLSLLFFMTTLAGQTAASLQLYGVCQAVCAAGVVAWYHQFELVFGTIVANADAPREALACNASYGACQRKCAYFFLAQ
>CAF1383500.1 unnamed protein product [Rotaria magnacalcarata]
MKSVAVFLCIIILTTSCVKAGPTAYGICQAGCAAVVVACYAAGGAVFGTVTAGAGASAAIIACNLAFGKCSAACAVAFFMPTP
>CAF3014777.1 unnamed protein product [Rotaria sp. Silwood2]
MKLSAIFLYIIILTASCSEAGPAAYGVCQAGCAALVVACYAAAGAVFGTVTAGAGITPAILACNIAFGKCSAACAVAVLMPTP
>CAF0862737.1 unnamed protein product [Rotaria sp. Silwood1]
MKLSGILLCIIILTASCIEGGPAAYALCQAGCAAVVVACYAAAGAVFGTVTAGAATAPAILACNIAFGKCSAVCAAIALTPTP
>CAF4273281.1 unnamed protein product [Rotaria sp. Silwood2]
MKLSAIFLYIIILTASCSEAGPAAYGVCQAGYAALVVACYAAAGAVFGTVTAGVGTTPAILACNIAFGKCSAACAVAALMPTP
>CAF3550138.1 unnamed protein product [Rotaria socialis]
MKSVALFLCIIILTTSCVKAGPTAYGICQAGCAAVVVACYAAAGAVFGTVTAGAGAPAAIIACNLAFGKCSAACAVAFFMPTP
>CAF1122499.1 unnamed protein product, partial [Rotaria sordida]
HASAGPVAYGICQAGCAALAVACYSAAGAVFGTVTAGIGTPPAIMACNSAFGVCSAKCALIALAPTP
>CAM9890062.1 unnamed protein product, partial [Scytosiphon promiscuus]
LPSKLVPHFAVAAVLFCFTSNFNRNPNLYPVDGICPIFFPRCNGVAVACYAAAGCTFGVATGGAAVPAVIVGCNSALGACMASCWAVTGAAAATPTL
>CAB9498995.1 expressed unknown protein [Seminavis robusta]
MMNFLLNANTCLRLFVALLLVGAVESGPLAYGICQTGCNGIVVACYGAAGFTFGTVTAGAGIPAALVACNTSLGACMAACVAAGCAPTP
>CAM9969753.1 unnamed protein product [Sphaerotrichia firma]
MKYIQVLLALAVIFAAPIPTTAGPLAALAAYGSCQTGCNVLACACYASAGAVFGAATGGAAVPAAVLGCNAGLGTCMAACWAVTGAAAATPTL
>PFX32892.1 hypothetical protein AWC38_SpisGene2161 [Stylophora pistillata]

*MVSFMKYVVVLAMFIQITQS*GPLAYGICQTGCNAVWVACVAAAGGVAGVSTGGIGVPAAILACNAAQGTCMAACVAAGLLPTL
>PFX32959.1 hypothetical protein AWC38_SpisGene2158 [Stylophora pistillata]
*MTHKNSEMAFVVKFIVVLAMFIQIAQS*GLQAYGICQTGCNTVWVDCVAAAGGTAGVSTGGIGVPGAILACNAAQGTCMVACVAAGAAPTP
>OLP78469.1 hypothetical protein AK812_SmicGene41345 [Symbiodinium microadriaticum]
MEMKVLLAVLLISQLSVADGGPLAYGICQTGCNAVAVACYAAAGATFGTVTAGVGTPAAIIACNSALGVCMAACVAAGCTPTP
>PNH04781.1 hypothetical protein TSOC_009012 [Tetrabaena socialis]
MSREPGAPAFRAADAGMLAYGICQCGCNVVTGACYAAGGFTFGTVTAGVGAPAVILACNIAQGACMTGCIAAGFAPTP
>PNH05266.1 hypothetical protein TSOC_008486 [Tetrabaena socialis]
MAFKPLRALLLAALLAAAASSVRAGPLAYAICQTGCNALVVACYGAAGYIFGTVTAGAGVPVAILTCNSKLGVW
>PNH03832.1 hypothetical protein TSOC_010074 [Tetrabaena socialis]
MPSTRKAALLLCALLLAATWCPTAVAGPLSYGICQTGCNALVVSCYTAAGFTFGTVTAGAGIPAVIIGCNAGLGFCMAGCVAAGCAPIP
>KAG5177616.1 hypothetical protein JKP88DRAFT_226309 [Tribonema minus]
MRQAVAALLGMCLALQGHTASAGPAAYAICQTGCNAGVVACYTAAGATFGTMTAGAGTPLVIIGCNTALGTCMAACVAAGLSPTL
>RDD42285.1 hypothetical protein TrispH2_006804 [Trichoplax sp. H2]
MKFRAQTIIVLILILQMQIFISEAGLLTYGLCQTACNGGWVTCYAFFGSVAGTVTAGVGTPGTLLACNSGQGYCMSVCAALFWAPTP
>XP_002949807.1 uncharacterized protein VOLCADRAFT_59734 [Volvox carteri f. nagariensis]
MRFLFVAKIVLFAAVLAAVTGSAAAGPIAYGICQSGCNAIAVACYASAGLTFGAVTAGRCFGAPAAAVACNLALGKCMACCIAAGFAPTF
>XP_002959848.1 uncharacterized protein VOLCADRAFT_127280 [Volvox carteri f. nagariensis]
MRFLFVAKIVLFAAVLAAVTGSAAAGPIAYGICQSGCNAIAVACYASAGLTFGAVTAGRCIGAPAAAIACNLALGKCMMCCIAAGFAPTF
>OQV14256.1 hypothetical protein BV898_11493 [Hypsibius exemplaris]
MGLKFRRYRPTVCVRPSVCYQTKVTLQYKTFRVSLRTSYTDHHLTLSHRKSPNWRTCRLGNLYYHEMNTRTSTAICLVFL
MAVAQVHSGLIGILAYGACQTGCNAGWVACCSAAGVTAGTVTAGLGVPAAVMACSALQGTCMAACAALALAPTP
>GAV05447.1 hypothetical protein RvY_15580 [Ramazzottius varieornatus]
MMSRSIVIFLGLLLVTDASAGPLAVAAAQAGCAAVQVACYAGAGAVFGTVTAGVGTPAAILACNSAFGTCMAAAYWAFFL
PTV
